# Cardiovascular Autonomic Neuropathy in Type 1 Diabetes is Associated with Several Metabolic Pathways – New Risk Markers on the Horizon

**DOI:** 10.1101/2021.05.18.444673

**Authors:** Christian S Hansen, Tommi Suvitaival, Simone Theilade, Ismo Mattila, Maria Lajer, Kajetan Trošt, Linda Ahonen, Tine W Hansen, Cristina Legido-Quigley, Peter Rossing, Tarunveer S Ahluwalia

**Author notes:** **Corresponding author**: Christian Stevns Hansen, MD, PhD, Steno Diabetes Center Copenhagen, Niels Steensens Vej 2-4, DK-2820 Gentofte, Denmark, ☎ (direct), ✉.

## Abstract

**Objective:** Cardiovascular autonomic neuropathy (CAN) in diabetes is associated with increased mortality and morbidity and is a non-treatable complication. We investigated associations between circulating metabolites and presence of CAN in persons with type 1 diabetes (T1D).

**Methods:** CAN was assessed by cardiovascular reflex tests (CARTs) in 302 persons with T1D as heart rate response to: deep breathing; lying-to-standing test; and the Valsalva manoeuvre. More than 1 pathological CART defined the CAN diagnosis.

Serum metabolomics and lipidomics profiles were analysed with two complementary non-targeted massspectrometry methods. Cross-sectional associations between single metabolites and CAN were assessed by linear regression. Models were fitted with and without adjustments for relevant confounders and multiple testing.

**Results:** Participants were mean (IQR) aged 55(49, 63) years, 50% males, with diabetes duration 39(32, 47) years, HbA1c 63(55,69) mmol/mol and 34% had the CAN diagnosis.

A total of 75 metabolites and 106 lipids were examined. In crude models, CAN diagnosis was associated with higher levels of hydroxy fatty acids (2,4- and 3,4-dihydroxybutanoic acids, 4–deoxytetronic acid), creatinine, sugar derivates (ribitol, ribonic acid, myo-inositol), citric acid, glycerol, phenols, phosphatidylcholines and lower levels of free fatty acids and amino acid methionine (p<0.05). Upon adjustment, positive associations with CAN were retained with hydroxy fatty acids, tricarboxylic acid (TCA) cycle-based sugar derivates, and citric acid and phenols (P_adjusted_<0.05).

**Conclusions:** Metabolic pathways, including the TCA cycle, hydroxy fatty acids, phosphatidylcholines and sugar derivatives, were associated with CAN in T1D. These novel metabolic pathways associated with CAN could prove to be future modifiable risk factors.

## Introduction

People with diabetes have are prone to complications a high prevalence of autonomic dysfunction e.g. the prevalence of cardiovascular autonomic neuropathy (CAN) ranges from 20% in unselected populations (1, 2) and between 35% to 65% in people with long-standing diabetes (3). CAN an independent predictor of cardiovascular mortality and morbidity (4–10) diabetic nephropathy(11–13). and bone metabolism(14). For decades strides have been taking to compose a treatment for CAN. Anti-oxidant treatment studies with alpha-lipoic acid have demonstrated conflicting results on efficacy (15, 16). Several drugs such angiotensin-converting enzyme inhibitors angiotensin II receptor blockers, aldose reductase inhibitors, β-adrenergic blockers have demonstrated beneficial effects on autonomic function in early CAN (17), but no benefits have been seen in patients with manifest CAN. Thus, despite the severity of CAN no disease modifying treatment for the complication exists (18). The exploration of novel risk factors is therefore warranted. Exploring this can be facilitated by the use of non-targeted investigations on the metabolome and lipidome. Indeed, several studies have revealed that intermediate metabolites are associated with the severity and progression of diabetic nephropathy (19–22) and retinopathy(23). Recently similar correlations have been found in regard to CAN. Autonomic dysfunction and components of the tricarboxylic acid (TCA) cycle(24) indicating the CAN is associated with changes in human energy metabolism. Our aim for the current study is to explore potential patho-metabolic pathways by investigating the association between circulating metabolites and presence of CAN in persons with type 1 diabetes (T1D).

## Methods

### Study population

Participants in this cross-sectional study were type 1 diabetes participants following the standard medical care at the outpatient clinic of Steno Diabetes Center Copenhagen, Denmark from 2009 to 2011 as described previously (25). Participants were recruited from a cohort of participants in a case-control study (conducted in 1993-2001) including 900 participants with either longstanding normoalbuminuria or diabetic nephropathy. 571 participants of the 900 participants were still alive for the cross-sectional study. 375 participants responded to the study invitation. A total of 73 were not eligible for inclusion; leaving 302 for analyses.

Only participants with at least one CAN measure or heart rate measures and omics profiles available were included. Forty-seven participants had missing values. Only participants in the age range of 20 to 80 years were included in the study due to non-existing reference values of CAN outside this age range. Twenty-six participants were outside the age range. Of the 302 participants, 258 had sufficient cardiovascular autonomic reflex tests (CARTs) performed to estimate the presence of the CAN diagnosis.

All participants gave written informed consent, the study conformed to the Declaration of Helsinki and the study protocol was approved by the local ethics committee; ClinicalTrials.gov ID NCT01171248.

### Assessment of autonomic function

Participants rested for 5 minutes in a supine position in a quiet room at room temperature (18-23 degrees Celsius) prior to assessment of autonomic function.

Autonomic function was assessed by measures of CAN by 2-minute resting heart rate variability (HRV) indices and cardiovascular autonomic reflex test (CARTs). The HRV measure the standard deviation of normal-to-normal intervals (SDNN) was derived from the resting heart. Subsequent to the HRV measurement, the three standard CARTs recommended for diagnosing CAN (26) were performed: the lying-to-standing test (30/15), the deep breathing test (E/I ratio) and the Valsalva maneuver. CARTs were performed in the mentioned.

CARTs and SDNN measures were analyzed as continuous variables. Age-dependent cut-off levels defined by Cardone et al. (27) were used to define pathological results of the CARTs. The CAN diagnosis was defined as the presence of two or three pathological CARTs as recommended by the American Diabetes Association (28) and classified for participants with more than one valid CART measure. Participants with one or no CARTs measures were classified as “no CAN estimation”. Higher values of the CAN measures applied for autonomic assessment in this study imply better autonomic function, whereas higher heart rate imply worse autonomic function.

Resting hear rate (HR), SDNN and CARTs were recorded by trained technicians using a Vagus^tm^ device (Medicus Engineering, Aarhus, Denmark).

### Metabolomics analyses. Sample quantification and Identification

The metabolomics and lipidomic analysis have been described in detail previously(29, 30). Serum samples were stored at −80°C until analysis by two different analytical methods. Metabolomics profiling of samples was performed using a two-dimensional gas chromatography with time of flight mass spectrometry. Peaks were identified from the raw data with ChromaTOF, and the resulting features aligned with Guineu.

Samples for lipidomics analysis were prepared using a modified Folch extraction procedure and analyzed by ultra-high-performance liquid chromatography quadruple time of flight mass spectrometry method. Raw acquired data was preprocessed with MZmine 2. A complete list of processed metabolites (including amino acids, free fatty acids, compounds from the energy metabolism pathways and polyols) is available in Tofte et al.(29) (30) were finally post processed in R software, as described previously (29). Lipid species have been defined by the number of carbon atoms (indicating total fatty acid chain length) and the number of double bonds for a specific species. These identities have been presented as “Class (number of carbon atoms:number of double-bonds)”.

The inclusion of metabolites and lipids (within the coverage of the two mass spectrometry platforms) in subsequent data analysis was solely based on the certainty of identification and the level of technical precision, thereby not restricting to any particular pathway or prior hypothesis.

### Baseline biochemical measures

HbA1c was measured by high-performance liquid chromatography (Variant, Bio-Rad Laboratories, Munich, Germany) and serum creatinine concentration by an enzymatic method (Hitachi 912; Roche Diagnostics, Mannheim, Germany). Urinary albumin excretion ratio was measured in 24-hour urine collections by an enzyme immunoassay. Chronic Kidney Disease Epidemiology Collaboration Equation was used to calculate the estimated glomerular filtration rate (eGFR) from p-creatinine.

### Anthropometric measures

Height and weight were measured with light indoor clothing, without shoes, using a fixed rigid stadiometer (Seca, Chino, USA) and an electronic scale (Mettler Toledo, Glostrup, Denmark), respectively.

### Blood pressure

Oscillometric (A&D Medical, UA787) office blood pressure was measured in a supine position after 15 minutes rest using an appropriate cuff size. Three measurements were obtained and averaged.

### Lifestyle measures

Lifestyle measures were obtained by questionnaires. Participants were classified as current smokers if using ≥ 1 cigarettes or cigars or pipes per day and all others were classified as non-smokers

### Statistical analyses

Continuous variables were reported as mean ± standard deviation (SD) for normally distributed data. Skewed data were reported as median (interquartile range, IQR) and were log2-transformed for analyses. Categorical variables were presented as total numbers with corresponding percentages. Comparisons of continuous and categorical variables between groups were performed using the analysis of variance (ANOVA) and X^2^-test, respectively. Data were imputed and auto-scaled prior to model-fitting. Associations between clinical characteristics of interest (the CAN diagnosis and CAN indices) and the levels of individual compounds were assessed with compound-specific linear regression models adjusted to clinical variables in a complete cases approach. Using the R package limma as described previously(30). All analyses were performed using three levels of confounder adjustment: Crude Model: no adjustment; Adjusted Model: adjusted age, sex, plasma glucose, HbA_1c_. body mass index, diabetes duration, smoking, statin use, total cholesterol and total triglycerides. To explore the effect of kidney function on associations a third model (fully adjusted Model) was applied with additional adjustment for eGFR. Due to 41 missing values of urinary albumin excretion rate (mg/24-hour) additional adjustments for albuminuria were done as a post-hoc analyses for models with significant findings. Results from regression analyses were visualized as forest plots, bipartite graphs and heatmaps using the lipidomeR package.

P-values for each analysis were corrected for multiple testing using the Benjamini–Hochberg method. Significant associations between clinical variables and molecule levels from crude models were integrated into a single visualization as a chord diagram using R-package circlize.

All data analyses were completed with R version 4.0.4 (The R Foundation for Statistical Computing, www.R-project.org).

## Results

Three-hundred-and-two participants were included in the final analyses. These were patients with any Cart or HRV measured. They were median (IQR) aged 55 (49;63) years, 50% males, with diabetes duration 39 (32;47) years, HbA1c 63 (55;69)mmol/mol, estimated glomerular filtration rate (eGFR) 83 (60;98)ml/min/1.73m2. A total of 258 had sufficient number of CARTs recorded to estimate the presence of the CAN diagnosis. A total of 88 (34%) persons were diagnosed with CAN. Group differences are displayed in Table 1.

**Table 1.** Baseline Characteristics

|  | No CAN, N = 170 | Definite CAN, N = 88 | No CAN estimation N=44 | p-value<br>(no CAN<br>vs. CAN) |
| --- | --- | --- | --- | --- |
| Sex (male), (n/%) | 76 (46%) | 45 (54%) | 24 (57%) | 0.3 |
| Age (years) | 55 (48, 62) | 55 (49, 60) | 60 (52, 66) | 0.7 |
| HbA1c (mmol/mol) | 62 (55, 67) | 64 (59, 73) | 61 (53, 69) | 0.002 |
| HbA1c (%) | 7.80 (7.20, 8.30) | 8.05 (7.57, 8.83) | 7.70 (7.03, 8.47) | 0.002 |
| Body mass index (kg/m<sup>2</sup>) | 24.2 (22.2, 26.9) | 24.0 (22.3, 26.2) | 23.8 (21.9, 26.1) | 0.7 |
| Systolic blood pressure (mmHg) | 128 (118, 141) | 131 (120, 147) | 130 (114, 137) | 0.10 |
| Diastolic blood pressure (mmHg) | 73 (68, 79) | 73 (68, 80) | 74 (68, 78) | >0.9 |
| Diabetes duration (years) | 37 (31, 45) | 40 (35, 48) | 43 (38, 51) | 0.026 |
| Smoking (n/%) | 26 (16%) | 20 (24%) | 10 (24%) | 0.2 |
| eGFR (ml/min/1.73 m<sup>2</sup>) | 87 (75, 101) | 62 (43, 84) | 83 (61, 104) | <0.001 |
| Urinary albumin excretion rate (mg/24-hour) | 10 (6, 23) | 56 (14, 432) | 11 (6, 54) | <0.001 |
| Total cholesterol (mmol/L) | 4.55 (4.18, 5.10) | 4.50 (4.00, 4.90) | 4.55 (4.12, 5.10) | 0.10 |
| HDL cholesterol (mmol/L) | 1.69 (1.39, 2.08) | 1.54 (1.27, 1.86) | 1.67 (1.39, 2.09) | 0.013 |
| LDL cholesterol (mmol/L) | 2.30 (1.90, 2.73) | 2.30 (1.90, 2.70) | 2.30 (2.00, 2.95) | 0.9 |
| Triglycerides (mmol/L) | 0.89 (0.68, 1.24) | 0.98 (0.80, 1.23) | 0.81 (0.67, 1.07) | 0.048 |
| Diuretic (n/%) | 65 (40%) | 68 (81%) | 22 (52%) | <0.001 |
| RAAS blocker (n/%) | 68 (42%) | 7 (8.3%) | 15 (36%) | <0.001 |
| Statins(n/%) | 90 (55%) | 67 (80%) | 21 (50%) | <0.001 |
| Pathological E/I ratio (n/%) |  |  |  |  |
| Pathological 30/15 ratio (n/%) |  |  |  |  |
| Pathological Valsalva (n/%) |  |  |  |  |
| E/I ratio | 1.17 (1.12, 1.28) | 1.04 (1.02, 1.07) | 1.10 (1.05, 1.16) | <0.001 |
| 30/15 ratio | 1.10 (1.06, 1.17) | 1.01 (1.00, 1.03) | 61 (53, 69) | <0.001 |
| Valsalva | 1.52 (1.36, 1.78) | 1.16 (1.11, 1.27) | 1.48 (1.32, 1.64) | <0.001 |
| SDNN (ms) | 30 (20, 42) | 10 (7, 15) | 20 (13, 33) | <0.001 |
| Heart rate (beats/minute) | 65 (58, 73) | 74 (68, 84) | 68 (59, 78) | <0.001 |

Data are in medians (IQR) or n (%). eGFR=Estimated glomerular filtration rate (ml/min/1.73m<sup>2</sup>), RAAS = Renin angiotensin aldosterone system, CAN=cardiovascular autonomic neuropathy. Wilcoxon rank sum test; Pearson's Chi-squared test; Fisher's exact test.

### Metabolomic analysis

A total of 75 metabolites were identified and passed quality control (Supplementary Table xx). Metabolites were part of the multivariate linear regression models.

In unadjusted models participants with CAN had higher level of hydroxy fatty acids (2,4- and 3,4-dihydroxybutanoic acids, 4–deoxytetronic acid), creatinine, sugar derivates (ribitol, ribonic acid, myoinositol), citric acid, glycerol, phenols, phosphatidylcholines and lower levels of free fatty acids and amino acid methionine participants with CAN (Figure 1 and 2). The majority of the associations to CARTs were seen for the E/I ration and the Valsalva maneuver. SDNN was not associated with any metabolites.

**Figure 1:**
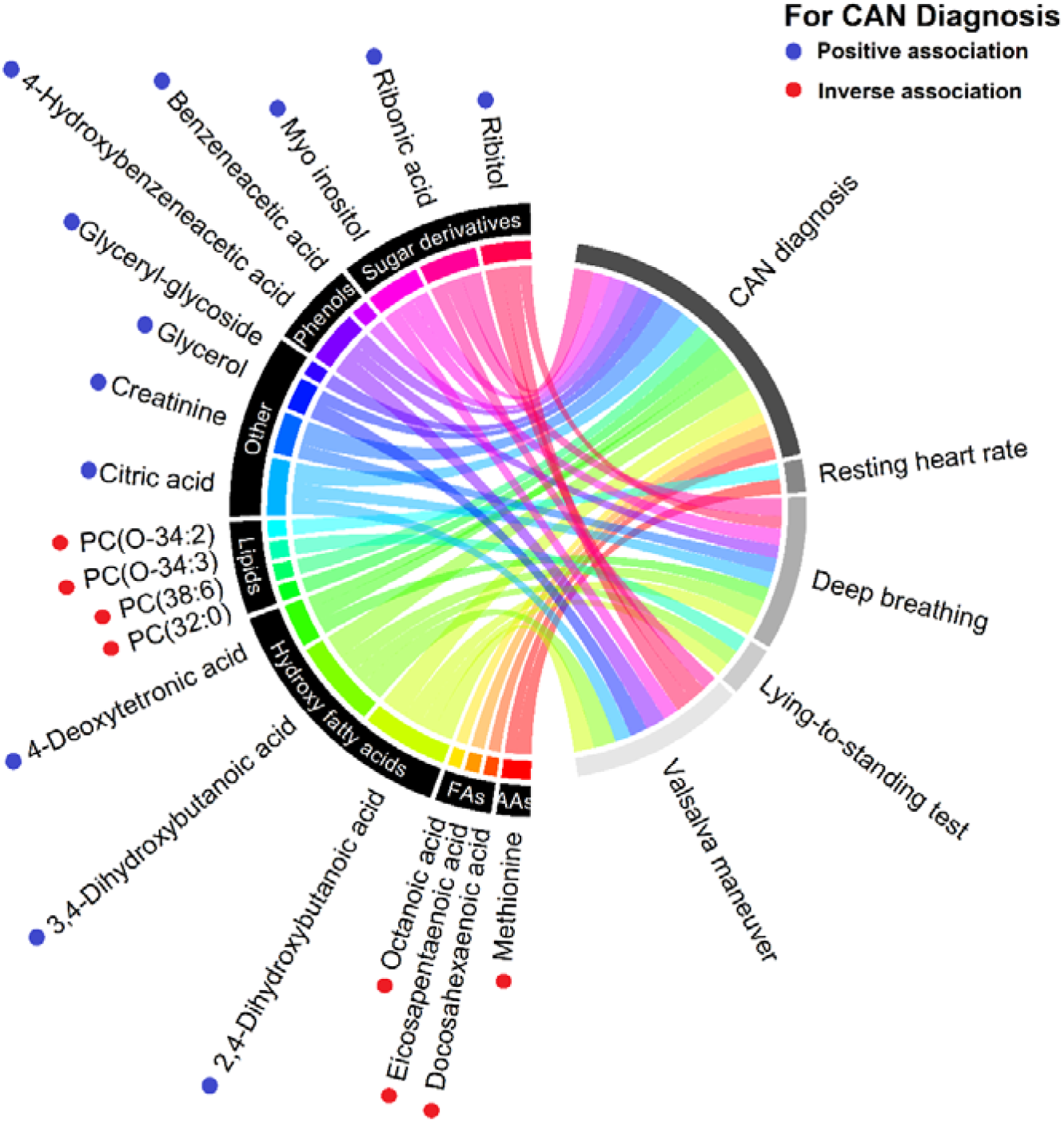
Chord diagram of the detected associations between metabolites(left) and the cardiovascular autonomic neuropathy (CAN) diagnosis and specific CAN measures (right) from the crude model. Metabolites are categorized into pathways and shown with unique colors. Line width indicate strenght of the respective assocation. PC: Phosphatidylcholines

**Figure 2:**
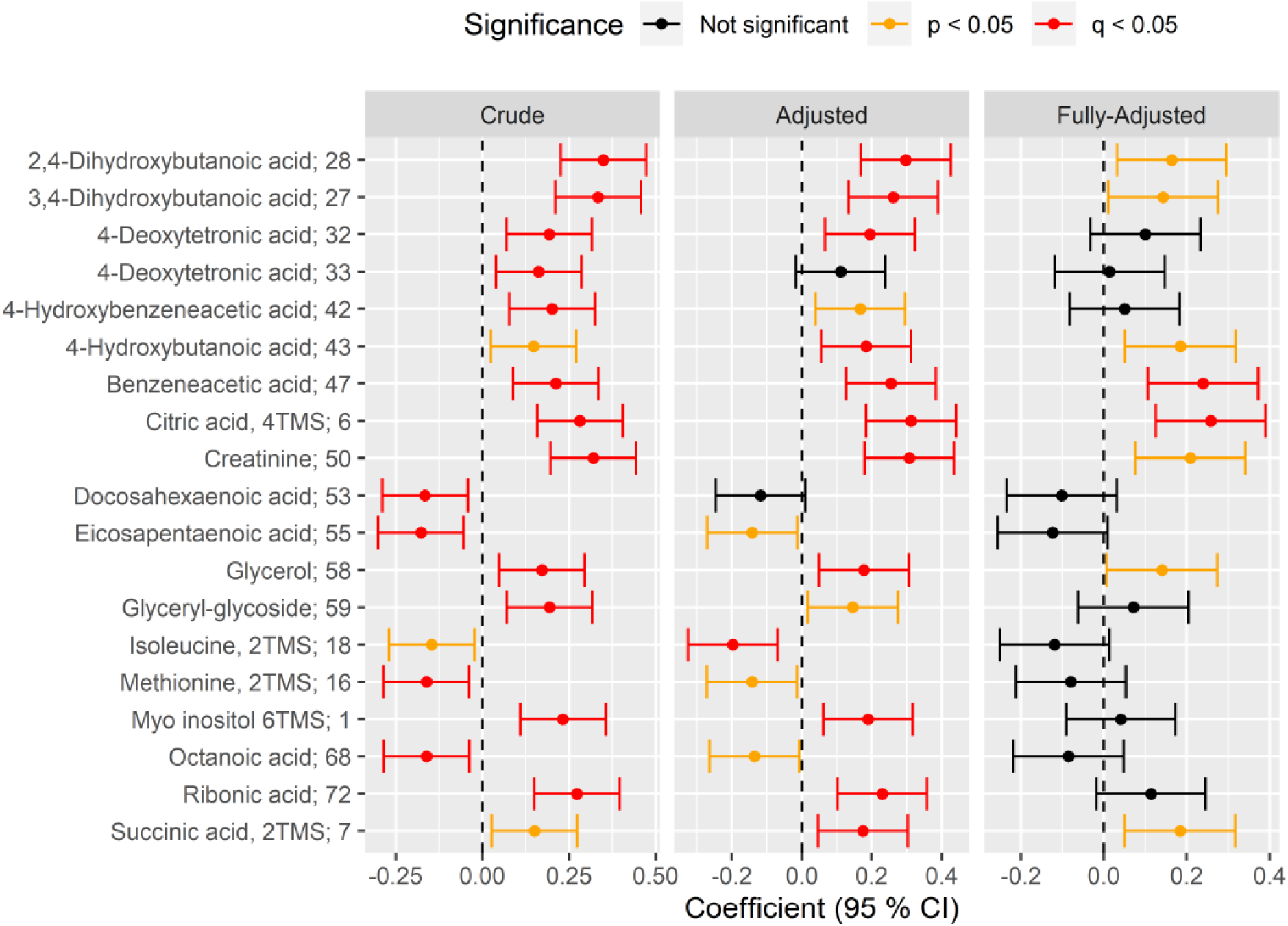
Forest plot of standardized association between CAN and metabolite level in crude (left), adjusted (middle) and fully-adjusted (right) metabolite-specific regression models (rows). Positive (negative) coefficient on the x-axis indicates positive (inverse) association between CAN and metabolite level. Statistical significance of the association is indicated by color of the confidence 95% interval (red: significant after correction to multiple testing; orange: significant nominal p-value; black: not significant.Results are shown for crude model (unadjusted; left), for models adjusted for adjusted age, sex, HbA1c, body mass index, diabetes duration, smoking, statin use, total cholesterol and total triglycerides (“Adjusted”; middle) and models further adjusted for eGFR (“Fully-Adjusted”, right).

In adjusted models, significant associations remained between the CAN diagnosis and CAN indices for several outcomes:

#### Hydroxy fatty acids

The CAN diagnosis was associated with higher levels of three hydroxy fatty acids in model 2: 2,4- and 3,4-dihydroxybutanoic acids, 4–deoxytetronic acid. Associations remained significant after additional adjustment for eGFR (Model 3).

Higher E/I ratio was associated with higher levels of all three above mentioned hydroxy fatty acids in unadjusted models. Higher levels of the Valsalva test were also associated with higher levels of 2,4- and 3,4–dihydroxybutanoic acids, but not 4–deoxytetronic acid. Associations were lost with further adjustments.

No other CAN indices were associated with hydroxy fatty acids (online appendix xxx).

#### TCA-cycle

In model 2 adjusted for adjusted age, sex, plasma glucose, HbA_1c_, body mass index, diabetes duration, smoking, statin use, beta blocker use, total cholesterol and total triglycerides we found the CAN diagnosis to be associated with higher levels of citrate, but no other metabolites of the TCA cycle. This association remained significant after additional adjustment for eGR (Table 2). Higher values of the E/I ratio and the Valsalva test were associated lower levels of citrate in unadjusted modes. The E/I ratio retained associations in model 2

No other associations to the TCA-cycle were found (Online supplementary material xxx).

#### Sugar derivatives

In unadjusted models the CAN diagnosis was associated with higher levels of myo inositol and ribonic acid. These findings remained consistent in model 2, but lost significance whit further adjustment for eGFR. (Table 2).

Results indicate that the above, mentioned associations were driven by the E/I-ratio and the Valsalva maneuver. Where higher values of both CARTs were associated with lower levels of the myo inositol in the unadjusted models. Higher values of the Valsalva maneuver were associated with lower levels of myo inositiol, ribonic acid and ribitol. The E/I ratio was inversely associated with ribitol in model 1 and 2. No other associations were found (Online supplementary material xxx).

#### Phenols

The CAN diagnosis was associated higher levels of bezeneacetic acid and 4-hydroxybezeneacetic acid in model 1 and 2. Only for bezeneacetic acid the associations remained significant in model 3 (Table 2). CARTs associated with phenols were the E/I ratio and the Vasalva maneuver. Where only higher values of the two CARTs were found to be associated with lower levels of 4-hydroxybezeneacetic acid and only in model 1(Online supplementary material xxx)

#### Amino acids

The CAN diagnosis was associated with two amino acid; methionine and isoleucine. Participant with CAN had lower levels of these amino acids in model 1 and 2. Associations were lost after further adjustment for eGFR.

Higher heart rate was associated with lower methionine, only in unadjusted models. No CARTs were shown to be associated with amino acids (Online supplementary material xxx).

### Lipidomic analysis

One-hundred-and-four lipid species from five major lipid classes were identified and underwent quality control. The 5 classes include diacyl-phosphatidylcholines (PCs), acyl-phosphatidylcholines, lyso-phosphatidylcholines (LPCs), triacylglycerols (TGs), free fatty acids (FFA) and sphingomyelins. The investigated lipids are listed under in online supplementary material xxx

#### Phosphatidylcholines

In unadjusted models the CAN diagnoses was associated with lower levels of phosphatidylcholine 38:6 and 32:0. And higher values of the 30/15 ratio was associated with higher values of the phosphatidylcholine 34:3. No other significant associations were found. (Online supplementary material xxx)

#### Triacylglycerols, Free fatty acids, Sphingomyelins

NO significant associations were found for any CAN measure and any triacylglycerols, free fatty acids or sphingomyelins

## Discussion

We investigated the blood metabolome in a cross-sectional cohort of 302 persons with type 1 diabetes with a diabetes duration of 39 years. We found that the CAN diagnosis was associated with several metabolic pathways including hydroxy fatty acids, the TCA cycle, sugar derivatives, phenols, amino acids and phosphatidylcholines. Below we have discussed the possible implications of separate categories of outcomes.

### Hydroxy fatty acids

In all models of adjustments then Can diagnosis was associated with 2,4- and 3,4-dihydroxybutanoic acids and 4–deoxytetronic acid. Analyses of CARTs indicate that both parasympathetic and sympathetic dysfunction may be eliciting changes in hydroxy fatty acid levels as lower (more detrimental) levels of the E/I ratio and the Valsalva test was associated with higher level of 2,4- and 3,4-dihydroxybutanoic acids in unadjusted models. A similar association was seen for the E/I-ratio with respect to 4–deoxytetronic acid. Associations between CAN and indices of CAN have not been described previously. Nor has the association be described in regard to other diabetic complications.

### TCA-cycle

In participants with CAN levels of citrate was higher compared to participants through all levels of adjustments. This association could be driven predominately by parasympathetic dysfunction was poorer results of the E/I ratio was associated with higher level of citrate in model 1 and 2, where the Valsalva test had the same associations but only in model 1.

Your results coincide with previous findings in type 1 diabetes (24) where metabolites of the TCA cycle have been associated with measures of CAN. Here, worse HRV indices (SDNN and RMSSD), was associated with higher levels of fumerate and citrate. The CAN diagnoses and CARTs were not found to associate to the TCA cycle. Our confirms that CAN may be related to disruptions in the TCA cycle and in a manner where more robust CAN measures are associated with these finding.

Citrate is an essential part of the TCA cycle which is the base of cellular energy metabolism. Diabetic animal studies have shown decreased mitochondrial dysfunction that are associated with microvascular diabetes complications (19). Nerve tissue specific diabetic animal studies have shown decreased levels of glycolytic and TCA intermediates in sural, sciatic, and dorsal root ganglion(31). Whether such changes are present in humans remains to be investigated. Taken together animal and human studies including the present study indicates that CAN is associated with changes in the TCA cycle. It remains however, to be investigated how serum measures of TCA intermediates in CAN may be related with tissue specific levels. In addition, the cross-sectional nature of our does not allow for conclusions off causal relations.

### Sugar derivatives

We found the CAN diagnose and both parasympathetic and sympathetic indices of the CAN diagnose to be associated with higher levels of myo inositol and ribonic acid. Such association have not been presented human studies previously. However, myo inositol is a pylol and may play a part in the pylol pathway, which is a molecular pathway associated with diabetic complications by creation of reactive oxygen species. The pylol pathway has been associated diabetic peripheral neuropathy(32) and nephropathy(33). Indeed, in the present study population we have shown higher levels of myo inositol to be predictive of future diabetic kidney disease(29). Thus, the CAN may be associated with higher level of sugar derivates where some are associated with the pylol pathway. Most plausible is that these sugar derivatives are risk factors to CAN and not vice versa. However, the present study design does not allow for conclusion of causality.

### Phenols

We found that patients with CAN had higher levels of bezeneacetic acid and 4-hydroxybezeneacetic and that both parasympathetic and sympathetic dysfunction was associated with higher levels of these phenols. These associations have not been presented previously. It is not known how and if these phenols represent a significant role in pathogenic mechanisms leading to diabetes complications.

### Amino acids

Patients with CAN had lower levels of methionine and isoleucine. In type 1 diabetes both increased and decreased levels of various amino acids have been found when compare to controls(34).However, the mechanisms and implications of theses imbalances are not understood. Worse levels of the HRV SDNN have been linked to lower levels of the amino acid glutamine in adjusted models(24). These finding underline that lower levels of some amin acids may be detrimental to the autonomic nervous system.

### Phosphatidylcholines

In crude analyses we found CAN to be associated with lower levels of phosphatidylcholine 38:6 and 32:0 and the E/I ratio the proportionally associated with phosphatidylcholine 34:3. Recently opposite associations were found in patient with recent onset type 2 diabetes(35), where more detrimental measures of autonomic dysfunction by HRV where associated with higher levels of several phosphatidylcholines including 32:0. The same study examined type 1 diabetes patients with recent onset of diabetes and found on associations between CAN and lipid compounds. To our knowledge no other studies haves investigated association between CAN and lipidomic profiles. Our study and the resent sited study indicate that phosphatidylcholines may play a part in the development of CAN, however potentially in differently in different durations of diabetes.

### Adjustment for eGFR

In models adjusted for eGFR several associations lost statistical significance. This may be due to confounding. However, we have not been able to elucidate pathological mechanisms which demonstrates that reduced kidney function is a reasonable explanation to this loss of associations. If indeed nephropathy plays a role in determination serum levels of the investigated metabolites it may be justified to reply heavily on the conclusion based on model 3. Such mechanisms remain to be explored.

## Conclusion

Metabolic pathways, including the TCA cycle, hydroxy fatty acids, phosphatidylcholines and sugar derivatives, were associated with CAN in T1D. These novel metabolic pathways associated with CAN could prove to be future modifiable risk factors.

## Strengths and limitations

The cross-sectional nature of the study does not allow for conclusions made on causality. Our findings may be caused by residual confounding Results may not be generalizable to a broad diabetes population as patients were not randomly recruited from the outpatient clinic.

## Supporting information

Lipidomics_supplementary_data

Metabolomics_supplementary_data

## Contributors

CSH interpreted the data and drafted the manuscript; TS analyzed and interpreted the data and made critical revision of the manuscript for key intellectual content. TSA conceived and designed the research, interpreted the data, made critical revision of the manuscript for key intellectual content and supervised the study. IM, KT, LA, ST and ML researched data and made critical revision of the manuscript for key intellectual content. TWH interpreted the data, and critically revised the manuscript for key intellectual content. PR conceived and designed the research, and interpreted the data, handled funding and supervision, made critical revision of the manuscript for key intellectual content and supervised the study. CSH is the guarantor of this work and, as such, had full access to all the data in the study and takes responsibility for the integrity of the data and the accuracy of the data analysis.

## Sources of funding

The work leading to this article received funding from the European Community’s Seventh Framework programme under grant agreement no. HEALTH-F2-2009-241544 (SysKID consortium). TSA was supported by Novo Nordisk Foundation (Steno Collaborative Grant) NNF18OC0052457.

## Declaration of interests

The authors declare that there is no duality of interest associated with this manuscript.

## Notes

### Competing Interest Statement

The authors have declared no competing interest.

