## Supplementary material for "Cardiovascular Autonomic Neuropathy in Type 1 Diabetes is Associated with Several Metabolic Pathways – New Risk Markers on the Horizon": Lipidomics_supplementary_data

### 0033_PROFIL_2017 Neuropathy – Lipidomics

Tommi Suvitaival,, Steno Diabetes Center Copenhagen

October 30, 2020

#### Contents

**1 Settings** **4**

**2 Load Data** **5**

**3 Filter** **6**

**4 Map Names** **7**

**5 CAN Stat** **8**

5.1 Crude Model . . . 8

5.1.1 Heatmap . . . 8

5.1.2 Tables of Model Coefficients . . . 10

5.1.3 Forest Plot of Model Coefficients . . . 13

5.2 Adjusted Model . . . 14

5.2.1 Heatmap . . . 14

5.2.2 Tables of Model Coefficients . . . 16

5.3 Fully-Adjusted Model . . . 19

5.3.1 Heatmap . . . 19

5.3.2 Tables of Model Coefficients . . . 21

**6 Vibration Sensation Threshold** **24**

6.1 Crude Model . . . 24

6.1.1 Heatmap . . . 24

6.1.2 Tables of Model Coefficients . . . 26

6.2 Adjusted Model . . . 29

6.2.1 Heatmap . . . 29

6.2.2 Tables of Model Coefficients . . . 31

6.3 Fully-Adjusted Model . . . 34

6.3.1 Heatmap . . . 34

6.3.2 Tables of Model Coefficients . . . 36

**7 Secondary Analyses** **39**
7.1 Resting HR Vagus . . . 39
7.1.1 Crude Model . . . 39
7.1.1.1 Heatmap . . . 39
7.1.1.2 Tables of Model Coefficients . . . 41
7.1.1.3 Forest Plot of Model Coefficients . . . 42
7.1.2 Adjusted Model . . . 43
7.1.2.1 Heatmap . . . 43
7.1.2.2 Tables of Model Coefficients . . . 45
7.1.3 Fully-Adjusted Model . . . 48
7.1.3.1 Heatmap . . . 48
7.1.3.2 Tables of Model Coefficients . . . 50
7.2 Deep Breathing (E_I) . . . 53
7.2.1 Crude Model . . . 53
7.2.1.1 Heatmap . . . 53
7.2.1.2 Tables of Model Coefficients . . . 55
7.2.2 Adjusted Model . . . 58
7.2.2.1 Heatmap . . . 58
7.2.2.2 Tables of Model Coefficients . . . 60
7.2.3 Fully-Adjusted Model . . . 63
7.2.3.1 Heatmap . . . 63
7.2.3.2 Tables of Model Coefficients . . . 65
7.3 Lying to Standing Test (lig_staa) . . . 68
7.3.1 Crude Model . . . 68
7.3.1.1 Heatmap . . . 69
7.3.1.2 Tables of Model Coefficients . . . 70
7.3.1.3 Forest Plot of Model Coefficients . . . 73
7.3.2 Adjusted Model . . . 74
7.3.2.1 Heatmap . . . 74
7.3.2.2 Tables of Model Coefficients . . . 76
7.3.3 Fully-Adjusted Model . . . 79
7.3.3.1 Heatmap . . . 79
7.3.3.2 Tables of Model Coefficients . . . 81
7.4 Valsalva Maneuver (Valsal) . . . 84
7.4.1 Crude Model . . . 84
7.4.1.1 Heatmap . . . 85

7.4.1.2 Tables of Model Coefficients . . . 86
7.4.2 Adjusted Model . . . 89
7.4.2.1 Heatmap . . . 89
7.4.2.2 Tables of Model Coefficients . . . 91
7.4.3 Fully-Adjusted Model . . . 94
7.4.3.1 Heatmap . . . 94
7.4.3.2 Tables of Model Coefficients . . . 96
7.5 Heart Rate Variablity (SDNN) . . . 99
7.5.1 Crude Model . . . 99
7.5.1.1 Heatmap . . . 99
7.5.1.2 Tables of Model Coefficients . . . 101
7.5.2 Adjusted Model . . . 104
7.5.2.1 Heatmap . . . 104
7.5.2.2 Tables of Model Coefficients . . . 106
7.5.3 Fully-Adjusted Model . . . 109
7.5.3.1 Heatmap . . . 109
7.5.3.2 Tables of Model Coefficients . . . 111
7.6 Neuropathy Questionnaire (mnsineuropat) . . . 114
7.6.1 Crude Model . . . 114
7.6.1.1 Heatmap . . . 114
7.6.1.2 Tables of Model Coefficients . . . 116
7.6.2 Adjusted Model . . . 119
7.6.2.1 Heatmap . . . 119
7.6.2.2 Tables of Model Coefficients . . . 121
7.6.3 Fully-Adjusted Model . . . 124
7.6.3.1 Heatmap . . . 124
7.6.3.2 Tables of Model Coefficients . . . 126

**8 Appendix** **129**

### 1 Settings

### 2 Load Data

### 3 Filter

```
## character(0)
```

### 4 Map Names

```
## [1] "map_lipid_names has been created by Tommi Suvitaival"
## [1] ""
## [1] "2019-05-06"
```

### 5 CAN Stat

#### 5.1 Crude Model

```
## [1] "Fitting models:"
## [1] "~ CAN_stat"
## [1] ""
```

##### 5.1.1 Heatmap

```
## [1] "heatmap_lipidome_from_limma was created by Tommi Suvitaival"
## [1] ""
## [1] "2019-05-21"
```

```
## Warning: Removed 102 rows containing missing values (geom_point).
```

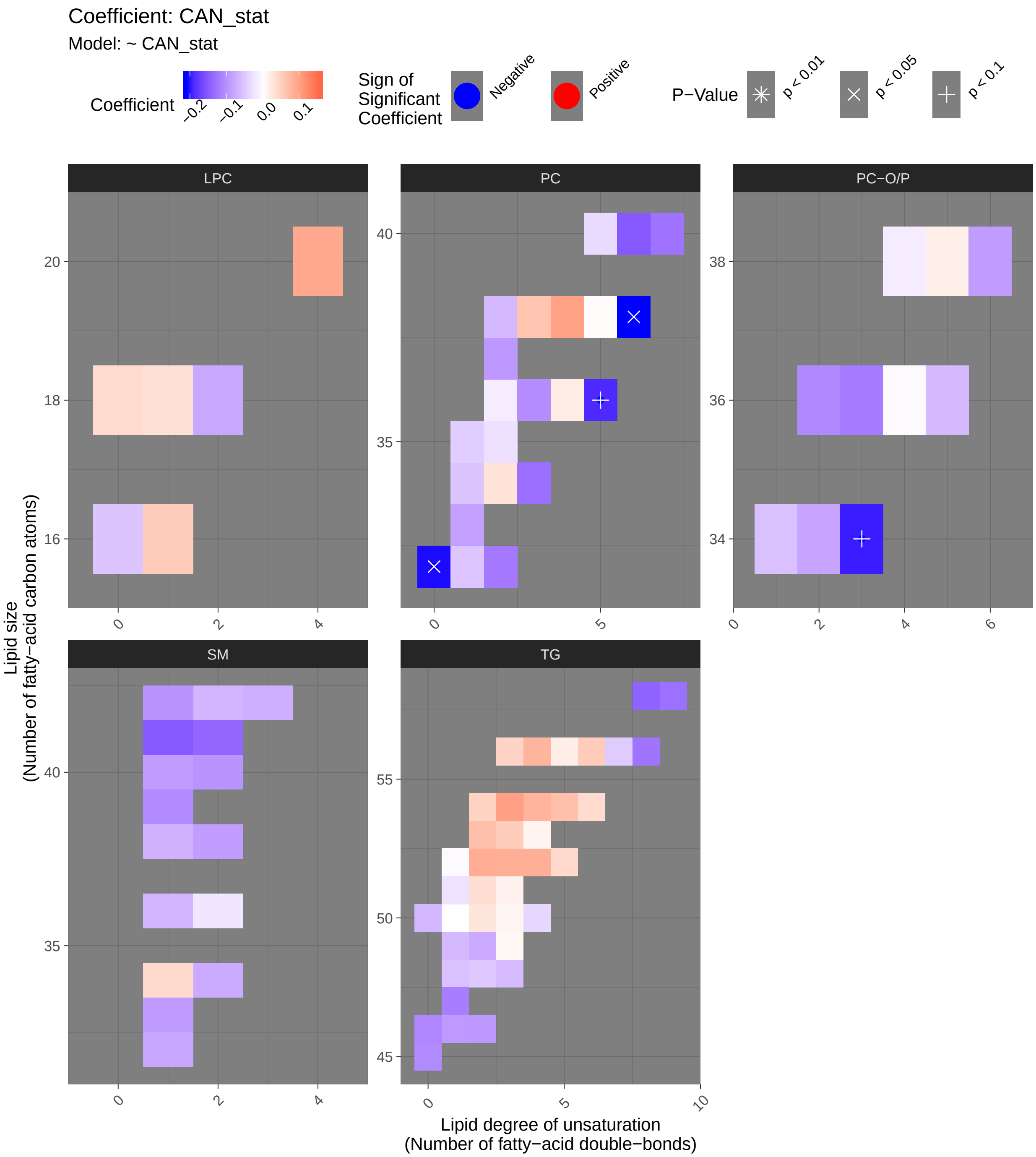
Coefficient: CAN_stat
Model: ~ CAN_stat
Coefficient
−0.2
−0.1
0.0
0.1
Sign of Significant Coefficient
Negative
Positive
P−Value
p < 0.01
p < 0.05
p < 0.1
LPC
PC
PC−O/P
SM
TG
Lipid size (Number of fatty−acid carbon atoms)
Lipid degree of unsaturation (Number of fatty−acid double−bonds)

##### 5.1.2 Tables of Model Coefficients

```
## [1] ""
## [1] "Table: CAN_stat"
## [1] "  (from model: "
## [1] "   ~ CAN_stat)"
## [1] ""
##                                Name Coefficient  P.Value adj.P.Val
## 1              PC(38:6)_[LVL2]; 8   -0.209000 0.000768    0.0488
## 2             PC(32:0)_[LVL2]; 96   -0.206000 0.000921    0.0488
## 3          PC(O-34:3)_[LVL2]; 140   -0.198000 0.001440    0.0509
## 4             PC(36:5)_[LVL2]; 23   -0.189000 0.002270    0.0603
## 5   TG(18:0/18:1/20:4)_[LVL2]; 141    0.156000 0.011800    0.2500
## 6             PC(40:6)_[LVL2]; 31   -0.150000 0.015500    0.2500
## 7           SM(d41:1)_[LVL2]; 102   -0.149000 0.016500    0.2500
## 8   TG(18:1/18:1/22:6)_[LVL2]; 147   -0.141000 0.022600    0.3000
## 9           SM(d41:2)_[LVL2]; 139   -0.138000 0.026200    0.3090
## 10           PC(34:3)_[LVL2]; 113   -0.130000 0.035800    0.3560
## 11           TG(58:9)_[LVL3]; 207   -0.128000 0.039200    0.3560
## 12           PC(40:7)_[LVL2]; 165   -0.126000 0.042200    0.3560
## 13  TG(16:0/18:2/22:6)_[LVL2]; 117   -0.125000 0.043700    0.3560
## 14           PC(32:2)_[LVL2]; 204   -0.121000 0.051600    0.3910
## 15         PC(O-36:3)_[LVL2]; 268   -0.119000 0.055400    0.3910
## 16           TG(47:1)_[LVL3]; 227   -0.116000 0.060700    0.4020
## 17   TG(18:2/22:5/16:0)_[LVL2]; 69   -0.113000 0.069300    0.4320
## 18           TG(46:0)_[LVL3]; 168   -0.109000 0.079800    0.4470
## 19         PC(O-36:2)_[LVL2]; 312   -0.106000 0.087700    0.4470
## 20            TG(45:0)_[LVL2]; 65   -0.106000 0.088000    0.4470
## 21          SM(d39:1)_[LVL2]; 179   -0.106000 0.088600    0.4470
## 22           TG(54:3)_[LVL3]; 124    0.103000 0.096700    0.4550
## 23            PC(36:3)_[LVL2]; 10   -0.102000 0.098900    0.4550
## 24             PC(38:4)_[LVL2]; 9    0.099800 0.108000    0.4550
## 25      SM(d18:1/24:0)_[LVL2]; 61   -0.097200 0.117000    0.4550
## 26   TG(18:1/18:1/18:1)_[LVL2]; 15    0.096800 0.119000    0.4550
## 27           SM(d40:2)_[LVL2]; 80   -0.096600 0.119000    0.4550
## 28          LPC(20:4)_[LVL2]; 120    0.093400 0.132000    0.4550
## 29           PC(37:2)_[LVL2]; 350   -0.092500 0.136000    0.4550
## 30           TG(46:2)_[LVL3]; 248   -0.092200 0.137000    0.4550
## 31           TG(46:1)_[LVL3]; 128   -0.091500 0.140000    0.4550
## 32          SM(d33:1)_[LVL2]; 166   -0.089900 0.147000    0.4550
## 33            TG(52:2)_[LVL3]; 97    0.089300 0.150000    0.4550
## 34         PC(O-38:6)_[LVL2]; 236   -0.089100 0.151000    0.4550
## 35           SM(d40:1)_[LVL2]; 39   -0.089000 0.151000    0.4550
## 36          SM(d38:2)_[LVL2]; 151   -0.087800 0.157000    0.4550
## 37           TG(52:3)_[LVL3]; 101    0.086300 0.164000    0.4550
## 38           TG(52:4)_[LVL3]; 157    0.086300 0.164000    0.4550
## 39           PC(33:1)_[LVL2]; 177   -0.085600 0.168000    0.4550
## 40         PC(O-34:2)_[LVL2]; 171   -0.081900 0.187000    0.4840
## 41           TG(54:4)_[LVL3]; 129    0.081800 0.187000    0.4840
## 42           TG(56:4)_[LVL3]; 278    0.080300 0.196000    0.4870
## 43          SM(d32:1)_[LVL2]; 105   -0.079900 0.198000    0.4870
## 44           LPC(18:2)_[LVL2]; 33   -0.077100 0.214000    0.5090
## 45           TG(49:2)_[LVL3]; 231   -0.076700 0.216000    0.5090
## 46  SM(d16:1/18:1) or SM(d18:2/16:   -0.075000 0.227000    0.5230
```

```
## 47       SM(d18:2/24:1)_[LVL2]; 40   -0.071400 0.250000    0.5580
## 48            SM(d38:1)_[LVL2]; 67   -0.070200 0.258000    0.5580
## 49            TG(54:5)_[LVL3]; 240    0.069400 0.263000    0.5580
## 50            TG(53:2)_[LVL2]; 234    0.069400 0.263000    0.5580
## 51             SM(42:2)_[LVL2]; 14   -0.066400 0.285000    0.5820
## 52            SM(d36:1)_[LVL2]; 55   -0.066300 0.286000    0.5820
## 53            TG(50:0)_[LVL2]; 159   -0.065100 0.294000    0.5840
## 54             PC(38:3)_[LVL2]; 29    0.063500 0.306000    0.5840
## 55            TG(49:1)_[LVL3]; 187   -0.062900 0.310000    0.5840
## 56          PC(O-36:5)_[LVL2]; 92   -0.062600 0.313000    0.5840
## 57            PC(38:2)_[LVL2]; 197   -0.062300 0.315000    0.5840
## 58            TG(48:3)_[LVL3]; 384   -0.061800 0.319000    0.5840
## 59   TG(18:2/18:1/18:1)_[LVL2]; 20    0.056800 0.360000    0.6290
## 60   TG(14:0/16:0/18:1)_[LVL2]; 54   -0.056400 0.363000    0.6290
## 61  PC(16:0e/18:1(9Z))_[LVL1]; 134   -0.056100 0.366000    0.6290
## 62            TG(56:6)_[LVL3]; 275    0.055600 0.370000    0.6290
## 63           LPC(16:1)_[LVL2]; 258    0.055100 0.374000    0.6290
## 64            TG(53:3)_[LVL3]; 239    0.054500 0.380000    0.6290
## 65              LPC(16:0)_[LVL1]; 5   -0.051500 0.406000    0.6450
## 66              PC(34:1)_[LVL2]; 2   -0.051500 0.406000    0.6450
## 67             PC(32:1)_[LVL2]; 44   -0.051400 0.407000    0.6450
## 68  TG(18:1/12:0/18:1) or TG(18:2/   -0.049500 0.425000    0.6620
## 69             TG(54:2)_[LVL3]; 52    0.047900 0.440000    0.6770
## 70            TG(56:3)_[LVL2]; 290    0.046400 0.454000    0.6790
## 71            TG(56:7)_[LVL3]; 309   -0.046400 0.455000    0.6790
## 72            PC(35:1)_[LVL2]; 178   -0.044900 0.469000    0.6810
## 73  TG(18:2/18:2/18:2) or TG(18:3/   -0.044900 0.469000    0.6810
## 74   TG(16:0/18:2/18:2)_[LVL2]; 27    0.044300 0.475000    0.6810
## 75            TG(52:5)_[LVL3]; 286    0.041800 0.500000    0.7030
## 76            SM(d34:1)_[LVL2]; 26    0.041400 0.504000    0.7030
## 77            LPC(18:0)_[LVL1]; 22    0.039600 0.524000    0.7210
## 78            TG(54:6)_[LVL3]; 316    0.038500 0.534000    0.7260
## 79  TG(14:0/18:2/18:2)_[LVL2]; 189   -0.035900 0.563000    0.7470
## 80  TG(16:0/22:5/18:1) or TG(20:4/    0.035800 0.564000    0.7470
## 81            TG(51:2)_[LVL2]; 123    0.035100 0.572000    0.7470
## 82    TG(18:1/18:1/16:0)_[LVL2]; 7    0.034100 0.582000    0.7470
## 83            LPC(18:1)_[LVL2]; 34    0.033900 0.585000    0.7470
## 84             PC(40:5)_[LVL2]; 95   -0.033100 0.593000    0.7490
## 85              PC(34:2)_[LVL2]; 4    0.031100 0.616000    0.7680
## 86            TG(50:2)_[LVL3]; 167    0.027600 0.656000    0.8000
## 87  TG(18:2/18:1/16:0)_[LVL2]; 500    0.027600 0.656000    0.8000
## 88            PC(35:2)_[LVL2]; 143   -0.026900 0.664000    0.8000
## 89            TG(51:1)_[LVL3]; 249   -0.025700 0.679000    0.8090
## 90   TG(14:0/18:1/18:1)_[LVL2]; 25    0.022800 0.714000    0.8410
## 91           SM(d36:2)_[LVL2]; 160   -0.021900 0.724000    0.8430
## 92              PC(36:4)_[LVL2]; 1    0.021000 0.735000    0.8470
## 93            TG(56:5)_[LVL2]; 230    0.019500 0.753000    0.8580
## 94          PC(O-38:5)_[LVL2]; 76    0.018000 0.772000    0.8650
## 95   TG(18:1/18:2/18:2)_[LVL2]; 57    0.017700 0.775000    0.8650
## 96         PC(O-38:4)_[LVL2]; 131   -0.016100 0.795000    0.8720
## 97              PC(36:2)_[LVL2]; 3   -0.015900 0.798000    0.8720
## 98            TG(51:3)_[LVL3]; 198    0.014000 0.822000    0.8890
## 99            TG(53:4)_[LVL3]; 314    0.012500 0.840000    0.8990
## 100            TG(50:3)_[LVL2]; 47    0.010600 0.864000    0.9160
```

```
## 101           TG(49:3)_[LVL3]; 218    0.007800 0.900000    0.9440
## 102 TG(16:0/18:2/18:3)_[LVL2]; 106    0.006660 0.915000    0.9500
## 103  TG(16:0/18:0/18:1)_[LVL2]; 51   -0.004720 0.939000    0.9630
## 104          PC(O-36:4)_[LVL2]; 71   -0.004310 0.945000    0.9630
## 105            PC(38:5)_[LVL2]; 24    0.003100 0.960000    0.9690
## 106            TG(50:1)_[LVL3]; 19   -0.000589 0.992000    0.9920
```

##### 5.1.3 Forest Plot of Model Coefficients

```
## Warning: Ignoring unknown aesthetics: x
```

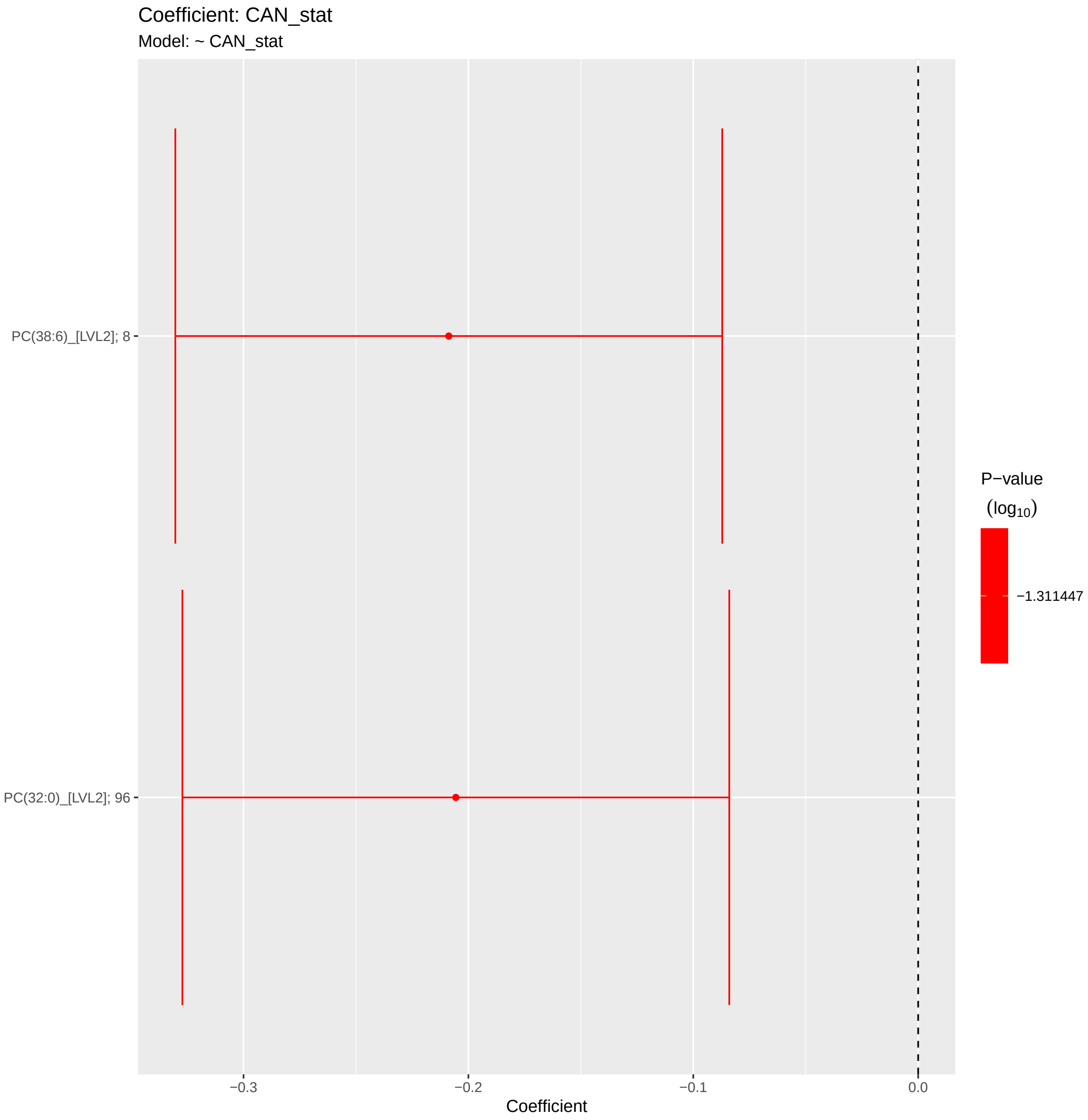

#### 5.2 Adjusted Model

```
## [1] "Fitting models:"
## [1] "~ CAN_stat + Age + bmi + Blood_glucose + Duration_DM + Gender + Hba1c_baseline + log_Blood_TGA +
## [1] ""
```

##### 5.2.1 Heatmap

```
## [1] "heatmap_lipidome_from_limma was created by Tommi Suvitaival"
## [1] ""
## [1] "2019-05-21"
```

```
## Warning: Removed 102 rows containing missing values (geom_point).
```

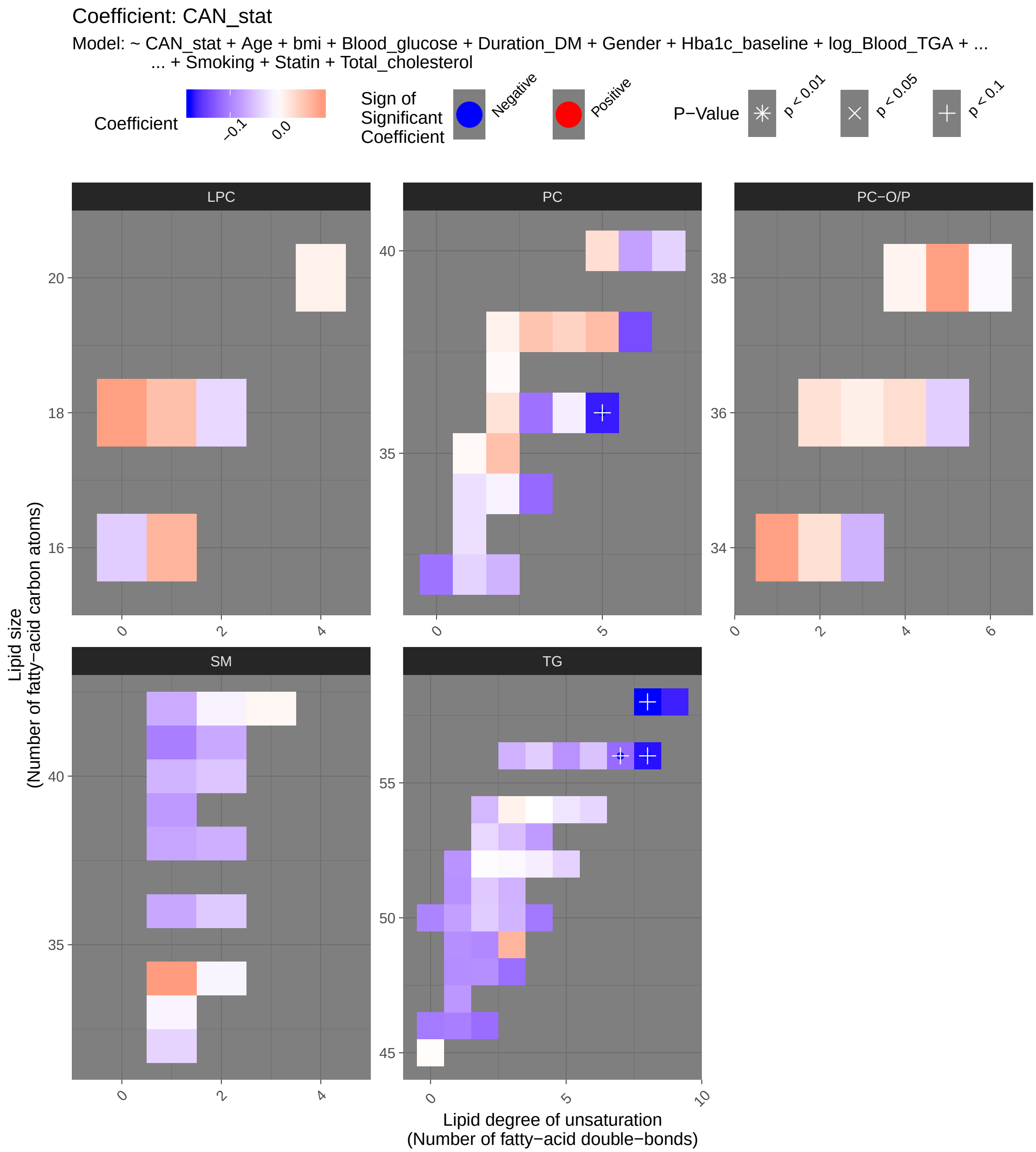
Coefficient: CAN_stat
Model: ~ CAN_stat + Age + bmi + Blood_glucose + Duration_DM + Gender + Hba1c_baseline + log_Blood_TGA + ...
... + Smoking + Statin + Total_cholesterol
Coefficient
−0.1
0.0
Sign of Significant Coefficient
Negative
Positive
P−Value
p < 0.01
p < 0.05
p < 0.1
LPC
PC
PC−O/P
SM
TG
Lipid size
(Number of fatty−acid carbon atoms)
Lipid degree of unsaturation
(Number of fatty−acid double−bonds)

##### 5.2.2 Tables of Model Coefficients

```
## [1] ""
## [1] "Table: CAN_stat"
## [1] "  (from model: "
## [1] "   ~ CAN_stat + Age + bmi + Blood_glucose + Duration_DM +"
## [1] "     Gender + Hba1c_baseline + log_Blood_TGA + Smoking + Statin +"
## [1] "     Total_cholesterol)"
## [1] ""
##                               Name Coefficient P.Value adj.P.Val
## 1              PC(36:5)_[LVL2]; 23   -0.175000 0.00118    0.0591
## 2   TG(18:1/18:1/22:6)_[LVL2]; 147   -0.185000 0.00149    0.0591
## 3   TG(16:0/18:2/22:6)_[LVL2]; 117   -0.180000 0.00167    0.0591
## 4    TG(18:2/22:5/16:0)_[LVL2]; 69   -0.161000 0.00328    0.0868
## 5             TG(58:9)_[LVL3]; 207   -0.173000 0.00475    0.1010
## 6               PC(38:6)_[LVL2]; 8   -0.143000 0.00717    0.1270
## 7   TG(14:0/18:2/18:2)_[LVL2]; 189   -0.107000 0.01500    0.2010
## 8   TG(18:2/18:2/18:2) or TG(18:3/   -0.125000 0.01610    0.2010
## 9             TG(48:3)_[LVL3]; 384   -0.116000 0.01710    0.2010
## 10            TG(46:2)_[LVL3]; 248   -0.116000 0.02730    0.2660
## 11            TG(56:7)_[LVL3]; 309   -0.119000 0.03040    0.2660
## 12  TG(18:1/12:0/18:1) or TG(18:2/   -0.090100 0.04340    0.2660
## 13            PC(34:3)_[LVL2]; 113   -0.120000 0.04480    0.2660
## 14             PC(32:0)_[LVL2]; 96   -0.112000 0.04550    0.2660
## 15   TG(14:0/16:0/18:1)_[LVL2]; 54   -0.091900 0.04860    0.2660
## 16   TG(16:0/18:0/18:1)_[LVL2]; 51   -0.086000 0.04930    0.2660
## 17             TG(50:1)_[LVL3]; 19   -0.075200 0.05080    0.2660
## 18            TG(51:1)_[LVL3]; 249   -0.087800 0.05110    0.2660
## 19            TG(46:0)_[LVL3]; 168   -0.107000 0.05560    0.2660
## 20            TG(49:2)_[LVL3]; 231   -0.094600 0.05660    0.2660
## 21             PC(36:3)_[LVL2]; 10   -0.113000 0.05680    0.2660
## 22            TG(46:1)_[LVL3]; 128   -0.102000 0.05760    0.2660
## 23          SM(d41:1)_[LVL2]; 102   -0.102000 0.05940    0.2660
## 24            TG(56:5)_[LVL2]; 230   -0.086500 0.06020    0.2660
## 25            TG(50:0)_[LVL2]; 159   -0.098300 0.07160    0.3030
## 26  TG(16:0/18:2/18:3)_[LVL2]; 106   -0.088000 0.07590    0.3050
## 27            TG(49:1)_[LVL3]; 187   -0.089000 0.07760    0.3050
## 28            TG(53:4)_[LVL3]; 314   -0.079600 0.09040    0.3400
## 29             TG(50:3)_[LVL2]; 47   -0.059200 0.09770    0.3400
## 30  TG(16:0/22:5/18:1) or TG(20:4/   -0.076700 0.09940    0.3400
## 31  PC(16:0e/18:1(9Z))_[LVL1]; 134    0.091000 0.09940    0.3400
## 32            SM(d34:1)_[LVL2]; 26    0.094700 0.11800    0.3830
## 33            TG(49:3)_[LVL3]; 218    0.071400 0.11900    0.3830
## 34            LPC(18:0)_[LVL1]; 22    0.090900 0.13500    0.4020
## 35   TG(18:1/18:2/18:2)_[LVL2]; 57   -0.075600 0.13900    0.4020
## 36            TG(51:3)_[LVL3]; 198   -0.061800 0.13900    0.4020
## 37          PC(O-38:5)_[LVL2]; 76    0.090700 0.14700    0.4020
## 38             TG(54:2)_[LVL3]; 52   -0.057200 0.14800    0.4020
## 39   TG(14:0/18:1/18:1)_[LVL2]; 25   -0.051700 0.15200    0.4020
## 40             PC(40:6)_[LVL2]; 31   -0.074500 0.15400    0.4020
## 41          SM(d39:1)_[LVL2]; 179   -0.081200 0.15600    0.4020
## 42            SM(d38:1)_[LVL2]; 67   -0.071500 0.18000    0.4500
## 43            TG(47:1)_[LVL3]; 227   -0.082800 0.18900    0.4500
## 44   TG(16:0/18:2/18:2)_[LVL2]; 27   -0.053200 0.19000    0.4500
```

```
## 45            SM(d36:1)_[LVL2]; 55   -0.070000 0.19200    0.4500
## 46           SM(d41:2)_[LVL2]; 139   -0.068700 0.19500    0.4500
## 47       SM(d18:1/24:0)_[LVL2]; 61   -0.066900 0.20300    0.4580
## 48            TG(56:3)_[LVL2]; 290   -0.061800 0.21400    0.4670
## 49            TG(53:3)_[LVL3]; 239   -0.051100 0.21600    0.4670
## 50           SM(d38:2)_[LVL2]; 151   -0.064100 0.24300    0.5030
## 51  TG(18:0/18:1/20:4)_[LVL2]; 141    0.068200 0.24500    0.5030
## 52            SM(d40:1)_[LVL2]; 39   -0.059300 0.24700    0.5030
## 53           LPC(16:1)_[LVL2]; 258    0.070900 0.26000    0.5030
## 54             PC(38:5)_[LVL2]; 24    0.063600 0.26100    0.5030
## 55            TG(51:2)_[LVL2]; 123   -0.042500 0.26500    0.5030
## 56          PC(O-34:3)_[LVL2]; 140   -0.060500 0.26600    0.5030
## 57            TG(50:2)_[LVL3]; 167   -0.039900 0.29400    0.5460
## 58            PC(32:2)_[LVL2]; 204   -0.060300 0.31500    0.5640
## 59            PC(35:2)_[LVL2]; 143    0.060400 0.31900    0.5640
## 60             PC(38:3)_[LVL2]; 29    0.057100 0.32100    0.5640
## 61            LPC(18:1)_[LVL2]; 34    0.060700 0.32500    0.5640
## 62            TG(56:6)_[LVL3]; 275   -0.048600 0.33000    0.5640
## 63            SM(d40:2)_[LVL2]; 80   -0.045700 0.36200    0.6090
## 64            TG(56:4)_[LVL3]; 278   -0.039100 0.39500    0.6550
## 65            TG(53:2)_[LVL2]; 234   -0.030800 0.43700    0.7090
## 66    TG(18:1/18:1/16:0)_[LVL2]; 7   -0.041700 0.44100    0.7090
## 67           SM(d36:2)_[LVL2]; 160   -0.041900 0.45100    0.7130
## 68            TG(52:5)_[LVL3]; 286   -0.035100 0.47600    0.7420
## 69              PC(38:4)_[LVL2]; 9    0.042800 0.48300    0.7420
## 70           SM(d32:1)_[LVL2]; 105   -0.033800 0.50800    0.7600
## 71              LPC(16:0)_[LVL1]; 5   -0.039500 0.51000    0.7600
## 72   TG(18:2/18:1/18:1)_[LVL2]; 20   -0.030100 0.52100    0.7600
## 73            PC(40:7)_[LVL2]; 165   -0.034700 0.52400    0.7600
## 74           PC(O-36:5)_[LVL2]; 92   -0.038400 0.53600    0.7670
## 75            TG(54:6)_[LVL3]; 316   -0.032400 0.54500    0.7670
## 76  TG(18:2/18:1/16:0)_[LVL2]; 500    0.035200 0.55700    0.7670
## 77             PC(32:1)_[LVL2]; 44   -0.034800 0.56400    0.7670
## 78             PC(40:5)_[LVL2]; 95    0.031000 0.56500    0.7670
## 79   TG(18:1/18:1/18:1)_[LVL2]; 15    0.025000 0.57400    0.7710
## 80            LPC(18:2)_[LVL2]; 33   -0.030900 0.60800    0.7900
## 81           PC(O-36:4)_[LVL2]; 71    0.032200 0.61000    0.7900
## 82          PC(O-36:2)_[LVL2]; 312    0.029000 0.61400    0.7900
## 83          PC(O-34:2)_[LVL2]; 171    0.030000 0.61900    0.7900
## 84              PC(36:2)_[LVL2]; 3    0.027900 0.66900    0.8430
## 85            PC(33:1)_[LVL2]; 177   -0.024000 0.67600    0.8430
## 86              PC(34:1)_[LVL2]; 2   -0.025000 0.70600    0.8670
## 87            TG(54:5)_[LVL3]; 240   -0.020000 0.71200    0.8670
## 88            TG(52:4)_[LVL3]; 157   -0.013000 0.76700    0.9120
## 89          PC(O-36:3)_[LVL2]; 268    0.015400 0.77200    0.9120
## 90            TG(54:3)_[LVL3]; 124    0.012700 0.77400    0.9120
## 91            PC(38:2)_[LVL2]; 197    0.013300 0.81600    0.9400
## 92           LPC(20:4)_[LVL2]; 120    0.013600 0.82500    0.9400
## 93              PC(36:4)_[LVL2]; 1   -0.013100 0.83900    0.9400
## 94             SM(42:2)_[LVL2]; 14   -0.009880 0.85600    0.9400
## 95  SM(d16:1/18:1) or SM(d18:2/16:   -0.008170 0.86100    0.9400
## 96          PC(O-38:4)_[LVL2]; 131    0.010900 0.86100    0.9400
## 97              PC(34:2)_[LVL2]; 4   -0.010200 0.87400    0.9400
## 98           SM(d33:1)_[LVL2]; 166   -0.008560 0.87600    0.9400
```

```
## 99        SM(d18:2/24:1)_[LVL2]; 40    0.007850 0.88400    0.9400
## 100            TG(52:3)_[LVL3]; 101   -0.005740 0.88700    0.9400
## 101            PC(35:1)_[LVL2]; 178    0.006380 0.91000    0.9550
## 102             TG(45:0)_[LVL2]; 65    0.003380 0.92800    0.9550
## 103          PC(O-38:6)_[LVL2]; 236   -0.004960 0.93100    0.9550
## 104            PC(37:2)_[LVL2]; 350    0.004520 0.93800    0.9550
## 105             TG(52:2)_[LVL3]; 97   -0.002410 0.94600    0.9550
## 106            TG(54:4)_[LVL3]; 129   -0.000159 0.99800    0.9980
```

#### 5.3 Fully-Adjusted Model

```
## [1] "Fitting models:"
## [1] "~ CAN_stat + Age + bmi + Blood_glucose + Duration_DM + Gender + Hba1c_baseline + log_Blood_TGA +
## [1] ""
```

##### 5.3.1 Heatmap

```
## [1] "heatmap_lipidome_from_limma was created by Tommi Suvitaival"
## [1] ""
## [1] "2019-05-21"
```

```
## Warning: Removed 106 rows containing missing values (geom_point).
```

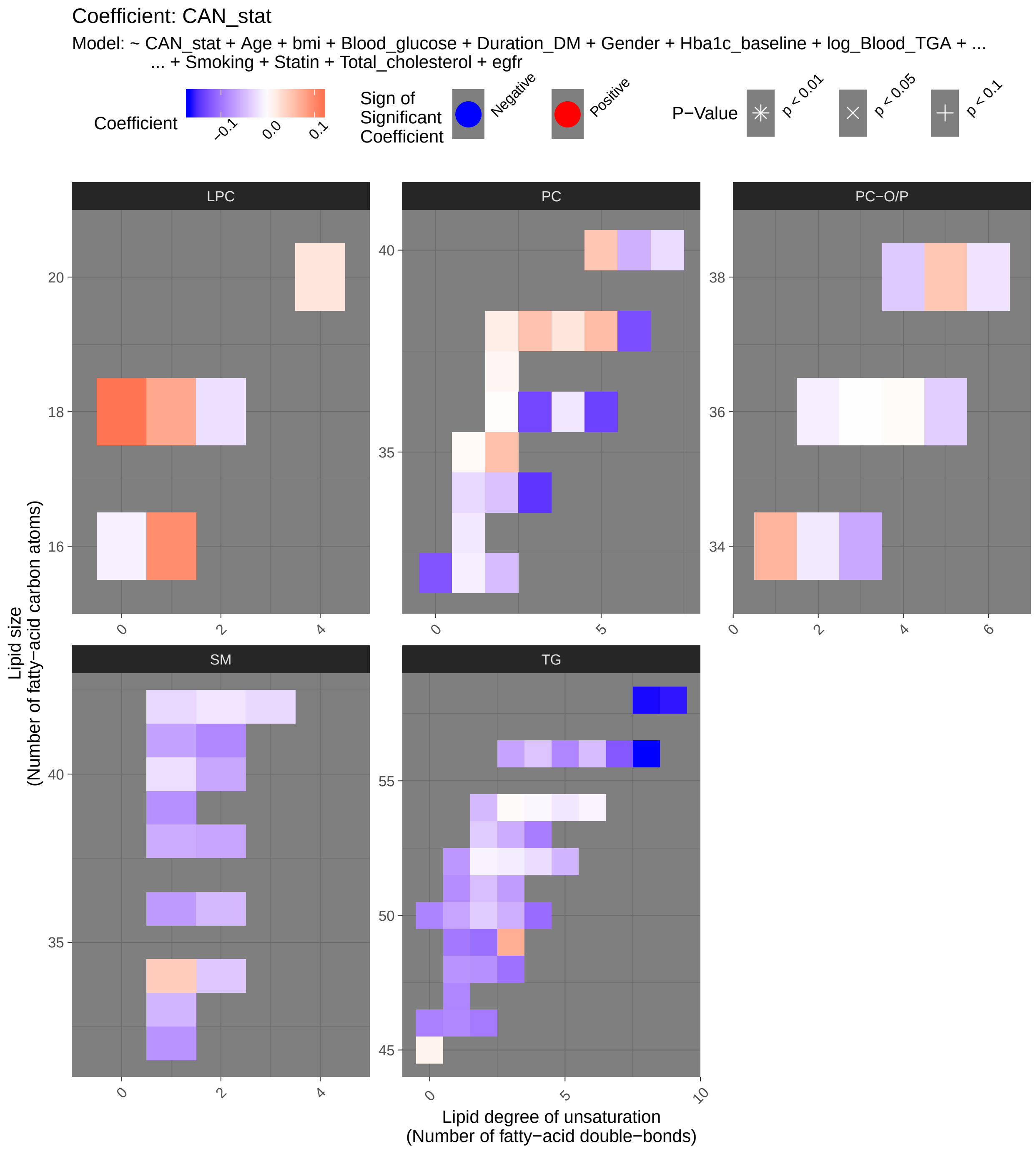
Coefficient: CAN_stat
Model: ~ CAN_stat + Age + bmi + Blood_glucose + Duration_DM + Gender + Hba1c_baseline + log_Blood_TGA + ...
... + Smoking + Statin + Total_cholesterol + egfr
Coefficient
−0.1
0.0
0.1
Sign of Significant Coefficient
Negative
Positive
P−Value
p < 0.01
p < 0.05
p < 0.1
LPC
PC
PC−O/P
SM
TG
Lipid size (Number of fatty−acid carbon atoms)
Lipid degree of unsaturation (Number of fatty−acid double−bonds)

##### 5.3.2 Tables of Model Coefficients

```
## [1] ""
## [1] "Table: CAN_stat"
## [1] "  (from model: "
## [1] "   ~ CAN_stat + Age + bmi + Blood_glucose + Duration_DM +"
## [1] "     Gender + Hba1c_baseline + log_Blood_TGA + Smoking + Statin +"
## [1] "     Total_cholesterol + egfr)"
## [1] ""
##                               Name Coefficient P.Value adj.P.Val
## 1   TG(16:0/18:2/22:6)_[LVL2]; 117   -0.166000 0.00552     0.306
## 2   TG(18:1/18:1/22:6)_[LVL2]; 147   -0.164000 0.00696     0.306
## 3    TG(18:2/22:5/16:0)_[LVL2]; 69   -0.150000 0.00939     0.306
## 4            TG(58:9)_[LVL3]; 207   -0.160000 0.01300     0.306
## 5             PC(36:5)_[LVL2]; 23   -0.136000 0.01530     0.306
## 6            PC(34:3)_[LVL2]; 113   -0.144000 0.02180     0.306
## 7              PC(38:6)_[LVL2]; 8   -0.127000 0.02300     0.306
## 8   TG(14:0/18:2/18:2)_[LVL2]; 189   -0.106000 0.02310     0.306
## 9             PC(36:3)_[LVL2]; 10   -0.132000 0.03330     0.356
## 10            PC(32:0)_[LVL2]; 96   -0.123000 0.03680     0.356
## 11           TG(56:7)_[LVL3]; 309   -0.120000 0.03700     0.356
## 12  TG(16:0/18:2/18:3)_[LVL2]; 106   -0.104000 0.04530     0.358
## 13           TG(49:2)_[LVL3]; 231   -0.104000 0.04590     0.358
## 14           TG(48:3)_[LVL3]; 384   -0.101000 0.04730     0.358
## 15  TG(18:2/18:2/18:2) or TG(18:3/   -0.103000 0.05830     0.396
## 16           TG(53:4)_[LVL3]; 314   -0.093100 0.05970     0.396
## 17           TG(49:1)_[LVL3]; 187   -0.096300 0.06930     0.403
## 18           LPC(18:0)_[LVL1]; 22    0.115000 0.07200     0.403
## 19           TG(56:5)_[LVL2]; 230   -0.087100 0.07230     0.403
## 20           TG(46:2)_[LVL3]; 248   -0.095400 0.08410     0.427
## 21           TG(51:1)_[LVL3]; 249   -0.081700 0.08460     0.427
## 22  TG(18:1/12:0/18:1) or TG(18:2/   -0.079500 0.08970     0.432
## 23   TG(16:0/18:0/18:1)_[LVL2]; 51   -0.074500 0.10500     0.444
## 24           TG(51:3)_[LVL3]; 198   -0.071200 0.10600     0.444
## 25            TG(50:1)_[LVL3]; 19   -0.065100 0.10700     0.444
## 26   TG(14:0/16:0/18:1)_[LVL2]; 54   -0.077200 0.11400     0.444
## 27           TG(46:0)_[LVL3]; 168   -0.091200 0.11800     0.444
## 28           TG(50:0)_[LVL2]; 159   -0.088800 0.12200     0.444
## 29          SM(d41:2)_[LVL2]; 139   -0.085300 0.12600     0.444
## 30            TG(50:3)_[LVL2]; 47   -0.056900 0.13100     0.444
## 31           TG(46:1)_[LVL3]; 128   -0.085200 0.13100     0.444
## 32  TG(16:0/22:5/18:1) or TG(20:4/   -0.072400 0.13900     0.444
## 33          LPC(16:1)_[LVL2]; 258    0.096900 0.14300     0.444
## 34          SM(d32:1)_[LVL2]; 105   -0.077200 0.14600     0.444
## 35           TG(49:3)_[LVL3]; 218    0.070000 0.14700     0.444
## 36           TG(53:3)_[LVL3]; 239   -0.059600 0.17000     0.501
## 37   TG(16:0/18:2/18:2)_[LVL2]; 27   -0.057500 0.17800     0.501
## 38   TG(18:1/18:2/18:2)_[LVL2]; 57   -0.072100 0.18000     0.501
## 39          SM(d39:1)_[LVL2]; 179   -0.079600 0.18600     0.505
## 40           TG(47:1)_[LVL3]; 227   -0.086700 0.19200     0.508
## 41           SM(d36:1)_[LVL2]; 55   -0.072700 0.19800     0.512
## 42           TG(56:3)_[LVL2]; 290   -0.065700 0.21000     0.519
## 43   TG(14:0/18:1/18:1)_[LVL2]; 25   -0.047500 0.21100     0.519
## 44            TG(54:2)_[LVL3]; 52   -0.049900 0.23000     0.541
```

```
## 45           SM(d41:1)_[LVL2]; 102   -0.067700 0.23200     0.541
## 46            SM(d40:2)_[LVL2]; 80   -0.062600 0.23500     0.541
## 47            TG(51:2)_[LVL2]; 123   -0.046400 0.24800     0.550
## 48           LPC(18:1)_[LVL2]; 34    0.074800 0.24900     0.550
## 49  PC(16:0e/18:1(9Z))_[LVL1]; 134    0.064100 0.26700     0.567
## 50           SM(d38:2)_[LVL2]; 151   -0.064100 0.26700     0.567
## 51          PC(O-34:3)_[LVL2]; 140   -0.062300 0.27600     0.573
## 52  TG(18:0/18:1/20:4)_[LVL2]; 141    0.066200 0.28400     0.579
## 53            SM(d38:1)_[LVL2]; 67   -0.059200 0.29000     0.580
## 54            TG(52:5)_[LVL3]; 286   -0.053300 0.30400     0.590
## 55             PC(40:6)_[LVL2]; 31   -0.056100 0.30600     0.590
## 56             PC(38:5)_[LVL2]; 24    0.057200 0.33700     0.628
## 57    TG(18:1/18:1/16:0)_[LVL2]; 7   -0.054600 0.33800     0.628
## 58           SM(d33:1)_[LVL2]; 166   -0.053100 0.35100     0.636
## 59            TG(50:2)_[LVL3]; 167   -0.035900 0.37000     0.636
## 60            TG(56:6)_[LVL3]; 275   -0.046800 0.37200     0.636
## 61            TG(53:2)_[LVL2]; 234   -0.037000 0.37500     0.636
## 62           SM(d36:2)_[LVL2]; 160   -0.051500 0.37800     0.636
## 63             PC(40:5)_[LVL2]; 95    0.049700 0.38000     0.636
## 64             PC(38:3)_[LVL2]; 29    0.052500 0.38500     0.636
## 65            PC(35:2)_[LVL2]; 143    0.054300 0.39500     0.636
## 66            TG(56:4)_[LVL3]; 278   -0.041100 0.39600     0.636
## 67  SM(d16:1/18:1) or SM(d18:2/16:   -0.039100 0.42300     0.669
## 68           PC(O-38:5)_[LVL2]; 76    0.048400 0.45800     0.705
## 69            PC(32:2)_[LVL2]; 204   -0.046700 0.45900     0.705
## 70            SM(d34:1)_[LVL2]; 26    0.044100 0.48300     0.731
## 71   TG(18:1/18:1/18:1)_[LVL2]; 15    0.031300 0.50500     0.754
## 72              PC(34:2)_[LVL2]; 4   -0.043700 0.51600     0.759
## 73  TG(18:2/18:1/16:0)_[LVL2]; 500    0.039100 0.53600     0.778
## 74          PC(O-38:4)_[LVL2]; 131   -0.037900 0.55900     0.801
## 75           PC(O-36:5)_[LVL2]; 92   -0.035100 0.59100     0.832
## 76            TG(52:4)_[LVL3]; 157   -0.024400 0.59800     0.832
## 77       SM(d18:1/24:0)_[LVL2]; 61   -0.028300 0.60400     0.832
## 78   TG(18:2/18:1/18:1)_[LVL2]; 20   -0.024900 0.61500     0.836
## 79       SM(d18:2/24:1)_[LVL2]; 40   -0.027100 0.62900     0.844
## 80            SM(d40:1)_[LVL2]; 39   -0.023100 0.66500     0.882
## 81            PC(40:7)_[LVL2]; 165   -0.024000 0.67500     0.883
## 82              PC(34:1)_[LVL2]; 2   -0.028300 0.68600     0.886
## 83            LPC(18:2)_[LVL2]; 33   -0.022800 0.72000     0.909
## 84           LPC(20:4)_[LVL2]; 120    0.022700 0.72600     0.909
## 85              PC(38:4)_[LVL2]; 9    0.021900 0.73200     0.909
## 86             SM(42:2)_[LVL2]; 14   -0.018900 0.74200     0.909
## 87          PC(O-38:6)_[LVL2]; 236   -0.019600 0.74600     0.909
## 88            TG(52:3)_[LVL3]; 101   -0.013200 0.75600     0.910
## 89            TG(54:5)_[LVL3]; 240   -0.016900 0.76700     0.911
## 90             TG(45:0)_[LVL2]; 65    0.010600 0.78700     0.911
## 91            PC(33:1)_[LVL2]; 177   -0.016200 0.78900     0.911
## 92              PC(36:4)_[LVL2]; 1   -0.017200 0.80000     0.911
## 93            PC(38:2)_[LVL2]; 197    0.015000 0.80300     0.911
## 94          PC(O-34:2)_[LVL2]; 171   -0.015200 0.80900     0.911
## 95             TG(52:2)_[LVL3]; 97   -0.008750 0.81700     0.911
## 96          PC(O-36:2)_[LVL2]; 312   -0.011400 0.84800     0.937
## 97             PC(32:1)_[LVL2]; 44   -0.011200 0.86000     0.940
## 98             LPC(16:0)_[LVL1]; 5   -0.009550 0.87900     0.950
```

```
## 99            TG(54:6)_[LVL3]; 316   -0.007950 0.88700     0.950
## 100           PC(37:2)_[LVL2]; 350    0.007890 0.89700     0.951
## 101           TG(54:4)_[LVL3]; 129   -0.005640 0.91600     0.962
## 102           PC(35:1)_[LVL2]; 178    0.004680 0.93700     0.970
## 103           TG(54:3)_[LVL3]; 124    0.003330 0.94300     0.970
## 104          PC(O-36:4)_[LVL2]; 71    0.003590 0.95700     0.975
## 105             PC(36:2)_[LVL2]; 3    0.002130 0.97500     0.984
## 106         PC(O-36:3)_[LVL2]; 268    0.000598 0.99200     0.992
```

### 6 Vibration Sensation Threshold

#### 6.1 Crude Model

```
## [1] "Fitting models:"
## [1] "~ Vib_pat"
## [1] ""
```

##### 6.1.1 Heatmap

```
## [1] "heatmap_lipidome_from_limma was created by Tommi Suvitaival"
## [1] ""
## [1] "2019-05-21"
```

```
## Warning: Removed 106 rows containing missing values (geom_point).
```

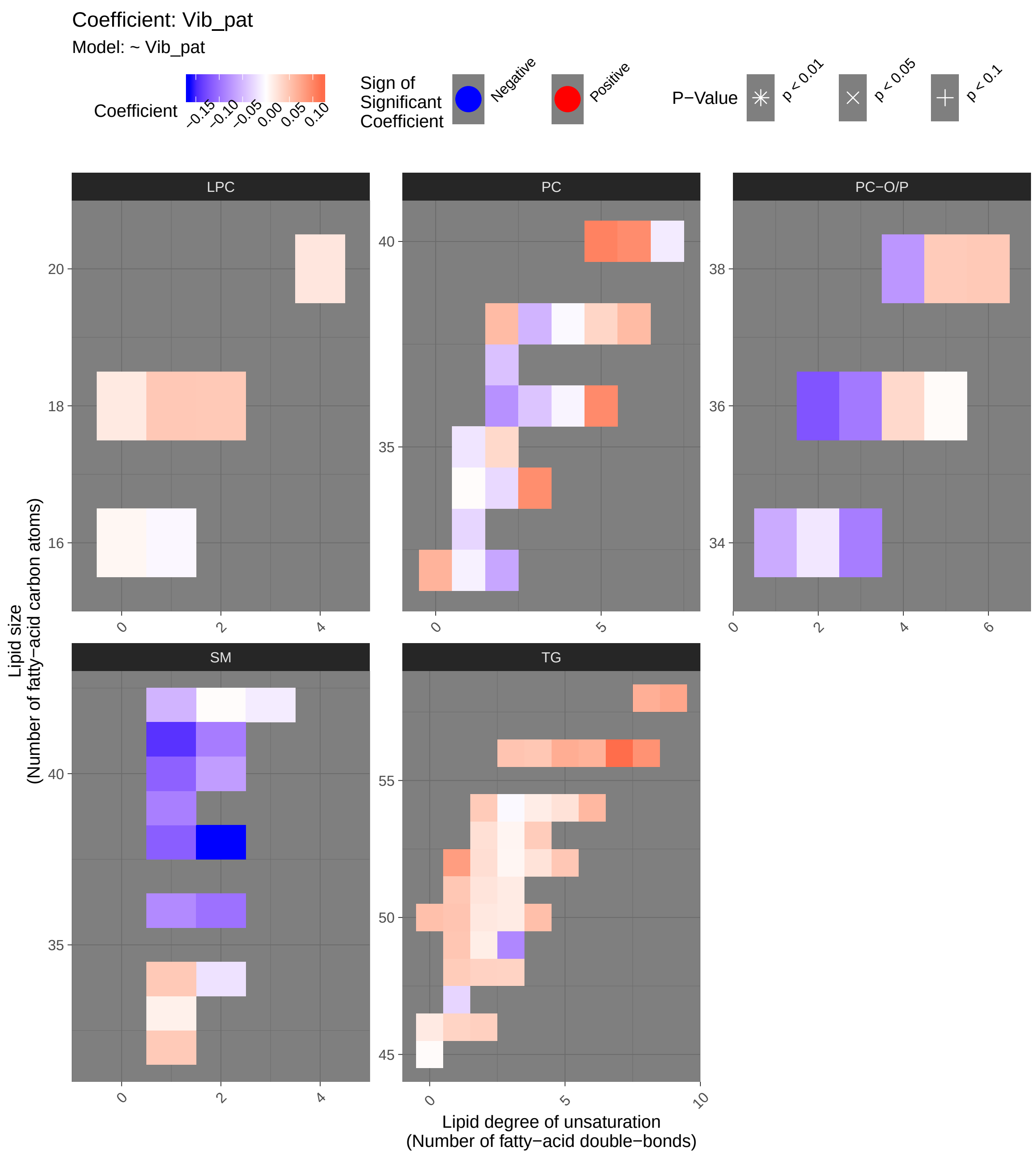
Coefficient: Vib_pat
Model: ~ Vib_pat
Coefficient
−0.15
−0.10
−0.05
0.00
0.05
0.10
Sign of Significant Coefficient
Negative
Positive
P−Value
p < 0.01
p < 0.05
p < 0.1
LPC
PC
PC−O/P
SM
TG
Lipid size (Number of fatty−acid carbon atoms)
Lipid degree of unsaturation (Number of fatty−acid double−bonds)

##### 6.1.2 Tables of Model Coefficients

```
## [1] ""
## [1] "Table: Vib_pat"
## [1] "  (from model: "
## [1] "    ~ Vib_pat)"
## [1] ""
##                                Name Coefficient P.Value adj.P.Val
## 1            SM(d38:2)_[LVL2]; 151    -0.16300 0.00458     0.485
## 2            SM(d41:1)_[LVL2]; 102    -0.14300 0.01310     0.694
## 3           PC(O-36:2)_[LVL2]; 312    -0.12100 0.03590     0.735
## 4              TG(56:7)_[LVL3]; 309     0.11900 0.03880     0.735
## 5             SM(d38:1)_[LVL2]; 67    -0.11400 0.04750     0.735
## 6             SM(d40:1)_[LVL2]; 39    -0.11200 0.05230     0.735
## 7    TG(18:2/22:5/16:0)_[LVL2]; 69     0.10600 0.06610     0.735
## 8              PC(40:5)_[LVL2]; 95     0.10300 0.07200     0.735
## 9            SM(d36:2)_[LVL2]; 160    -0.10000 0.08110     0.735
## 10             PC(36:5)_[LVL2]; 23     0.09690 0.09170     0.735
## 11             PC(40:6)_[LVL2]; 31     0.09530 0.09710     0.735
## 12          PC(O-36:3)_[LVL2]; 268    -0.09470 0.09920     0.735
## 13            PC(34:3)_[LVL2]; 113     0.09430 0.10100     0.735
## 14           SM(d41:2)_[LVL2]; 139    -0.09200 0.10900     0.735
## 15          PC(O-34:3)_[LVL2]; 140    -0.09170 0.11000     0.735
## 16  TG(16:0/18:2/22:6)_[LVL2]; 117     0.09110 0.11300     0.735
## 17           SM(d39:1)_[LVL2]; 179    -0.08980 0.11800     0.735
## 18            TG(49:3)_[LVL3]; 218    -0.08450 0.14100     0.810
## 19   TG(16:0/18:0/18:1)_[LVL2]; 51     0.08250 0.15100     0.810
## 20            SM(d36:1)_[LVL2]; 55    -0.08200 0.15300     0.810
## 21  TG(16:0/18:2/18:3)_[LVL2]; 106     0.08030 0.16200     0.810
## 22  TG(18:2/18:2/18:2) or TG(18:3/     0.07890 0.17000     0.810
## 23              PC(36:2)_[LVL2]; 3    -0.07720 0.17900     0.810
## 24  TG(16:0/22:5/18:1) or TG(20:4/     0.07540 0.19000     0.810
## 25            TG(58:9)_[LVL3]; 207     0.07510 0.19100     0.810
## 26          PC(O-38:4)_[LVL2]; 131    -0.07340 0.20100     0.821
## 27            TG(56:5)_[LVL2]; 230     0.06930 0.22800     0.859
## 28            SM(d40:2)_[LVL2]; 80    -0.06870 0.23200     0.859
## 29  TG(18:1/18:1/22:6)_[LVL2]; 147     0.06780 0.23800     0.859
## 30            TG(56:6)_[LVL3]; 275     0.06540 0.25500     0.859
## 31             PC(32:0)_[LVL2]; 96     0.06410 0.26400     0.859
## 32            PC(32:2)_[LVL2]; 204    -0.06230 0.27800     0.859
## 33            TG(54:6)_[LVL3]; 316     0.06000 0.29600     0.859
## 34  PC(16:0e/18:1(9Z))_[LVL1]; 134    -0.05890 0.30500     0.859
## 35              PC(38:6)_[LVL2]; 8     0.05790 0.31300     0.859
## 36            PC(38:2)_[LVL2]; 197     0.05750 0.31700     0.859
## 37  TG(14:0/18:2/18:2)_[LVL2]; 189     0.05440 0.34300     0.859
## 38            TG(50:0)_[LVL2]; 159     0.05340 0.35200     0.859
## 39             PC(38:3)_[LVL2]; 29    -0.05250 0.36100     0.859
## 40       SM(d18:1/24:0)_[LVL2]; 61    -0.05230 0.36300     0.859
## 41             TG(50:1)_[LVL3]; 19     0.04990 0.38500     0.859
## 42            TG(56:3)_[LVL2]; 290     0.04970 0.38700     0.859
## 43            TG(49:1)_[LVL3]; 187     0.04840 0.40000     0.859
## 44            TG(51:1)_[LVL3]; 249     0.04790 0.40400     0.859
## 45            TG(56:4)_[LVL3]; 278     0.04760 0.40700     0.859
## 46            TG(52:5)_[LVL3]; 286     0.04720 0.41100     0.859
```

```
## 47            LPC(18:1)_[LVL2]; 34     0.04640 0.41900     0.859
## 48          PC(O-38:6)_[LVL2]; 236     0.04600 0.42300     0.859
## 49            SM(d34:1)_[LVL2]; 26     0.04600 0.42300     0.859
## 50            LPC(18:2)_[LVL2]; 33     0.04570 0.42600     0.859
## 51           SM(d32:1)_[LVL2]; 105     0.04510 0.43200     0.859
## 52             TG(54:2)_[LVL3]; 52     0.04430 0.44100     0.859
## 53           PC(O-38:5)_[LVL2]; 76     0.04410 0.44300     0.859
## 54   TG(14:0/16:0/18:1)_[LVL2]; 54     0.04350 0.44800     0.859
## 55            PC(37:2)_[LVL2]; 350    -0.04320 0.45200     0.859
## 56            TG(53:4)_[LVL3]; 314     0.04310 0.45400     0.859
## 57             PC(36:3)_[LVL2]; 10    -0.04110 0.47500     0.876
## 58  TG(18:2/18:1/16:0)_[LVL2]; 500    -0.04060 0.47900     0.876
## 59            TG(46:2)_[LVL3]; 248     0.03980 0.48900     0.878
## 60  TG(18:1/12:0/18:1) or TG(18:2/     0.03810 0.50700     0.885
## 61            TG(48:3)_[LVL3]; 384     0.03730 0.51700     0.885
## 62            TG(46:1)_[LVL3]; 128     0.03670 0.52300     0.885
## 63   TG(18:1/18:2/18:2)_[LVL2]; 57     0.03640 0.52600     0.885
## 64             PC(38:5)_[LVL2]; 24     0.03530 0.53900     0.889
## 65  TG(18:0/18:1/20:4)_[LVL2]; 141     0.03480 0.54500     0.889
## 66            PC(35:2)_[LVL2]; 143     0.03270 0.57000     0.911
## 67           PC(O-36:4)_[LVL2]; 71     0.03220 0.57600     0.911
## 68   TG(18:1/18:1/18:1)_[LVL2]; 15    -0.03140 0.58400     0.911
## 69            TG(47:1)_[LVL3]; 227    -0.02900 0.61400     0.928
## 70            PC(33:1)_[LVL2]; 177    -0.02820 0.62400     0.928
## 71             TG(52:2)_[LVL3]; 97     0.02780 0.62900     0.928
## 72            TG(53:2)_[LVL2]; 234     0.02660 0.64300     0.928
## 73               PC(34:2)_[LVL2]; 4    -0.02650 0.64500     0.928
## 74   TG(16:0/18:2/18:2)_[LVL2]; 27     0.02630 0.64800     0.928
## 75    TG(18:1/18:1/16:0)_[LVL2]; 7     0.02480 0.66600     0.928
## 76            TG(54:5)_[LVL3]; 240     0.02450 0.67000     0.928
## 77            TG(52:4)_[LVL3]; 157     0.02400 0.67600     0.928
## 78   TG(14:0/18:1/18:1)_[LVL2]; 25     0.02290 0.69000     0.928
## 79            TG(51:2)_[LVL2]; 123     0.02280 0.69200     0.928
## 80           LPC(20:4)_[LVL2]; 120     0.02100 0.71500     0.936
## 81  SM(d16:1/18:1) or SM(d18:2/16:    -0.02020 0.72500     0.936
## 82            TG(50:2)_[LVL3]; 167     0.01940 0.73500     0.936
## 83            LPC(18:0)_[LVL1]; 22     0.01820 0.75100     0.936
## 84            PC(35:1)_[LVL2]; 178    -0.01820 0.75200     0.936
## 85            TG(46:0)_[LVL3]; 168     0.01760 0.75900     0.936
## 86            TG(51:3)_[LVL3]; 198     0.01710 0.76600     0.936
## 87             TG(50:3)_[LVL2]; 47     0.01680 0.77000     0.936
## 88          PC(O-34:2)_[LVL2]; 171    -0.01620 0.77700     0.936
## 89            TG(54:4)_[LVL3]; 129     0.01520 0.79100     0.936
## 90            TG(49:2)_[LVL3]; 231     0.01490 0.79500     0.936
## 91            PC(40:7)_[LVL2]; 165    -0.01400 0.80700     0.940
## 92       SM(d18:2/24:1)_[LVL2]; 40    -0.01340 0.81600     0.940
## 93           SM(d33:1)_[LVL2]; 166     0.01230 0.83000     0.947
## 94             PC(32:1)_[LVL2]; 44    -0.00963 0.86700     0.964
## 95            TG(53:3)_[LVL3]; 239     0.00831 0.88500     0.964
## 96            TG(52:3)_[LVL3]; 101     0.00776 0.89300     0.964
## 97               PC(36:4)_[LVL2]; 1    -0.00764 0.89400     0.964
## 98             LPC(16:0)_[LVL1]; 5     0.00717 0.90100     0.964
## 99           LPC(16:1)_[LVL2]; 258    -0.00555 0.92300     0.964
## 100  TG(18:2/18:1/18:1)_[LVL2]; 20    -0.00482 0.93300     0.964
```

```
## 101            TG(54:3)_[LVL3]; 124    -0.00441 0.93900     0.964
## 102              PC(38:4)_[LVL2]; 9    -0.00441 0.93900     0.964
## 103          PC(O-36:5)_[LVL2]; 92     0.00381 0.94700     0.964
## 104             TG(45:0)_[LVL2]; 65     0.00306 0.95700     0.964
## 105             SM(42:2)_[LVL2]; 14     0.00273 0.96200     0.964
## 106              PC(34:1)_[LVL2]; 2     0.00261 0.96400     0.964
```

#### 6.2 Adjusted Model

```
## [1] "Fitting models:"
## [1] "~ Vib_pat + Age + bmi + Blood_glucose + Duration_DM + Gender + Hba1c_baseline + log_Blood_TGA +
## [1] ""
```

##### 6.2.1 Heatmap

```
## [1] "heatmap_lipidome_from_limma was created by Tommi Suvitaival"
## [1] ""
## [1] "2019-05-21"
```

```
## Warning: Removed 106 rows containing missing values (geom_point).
```

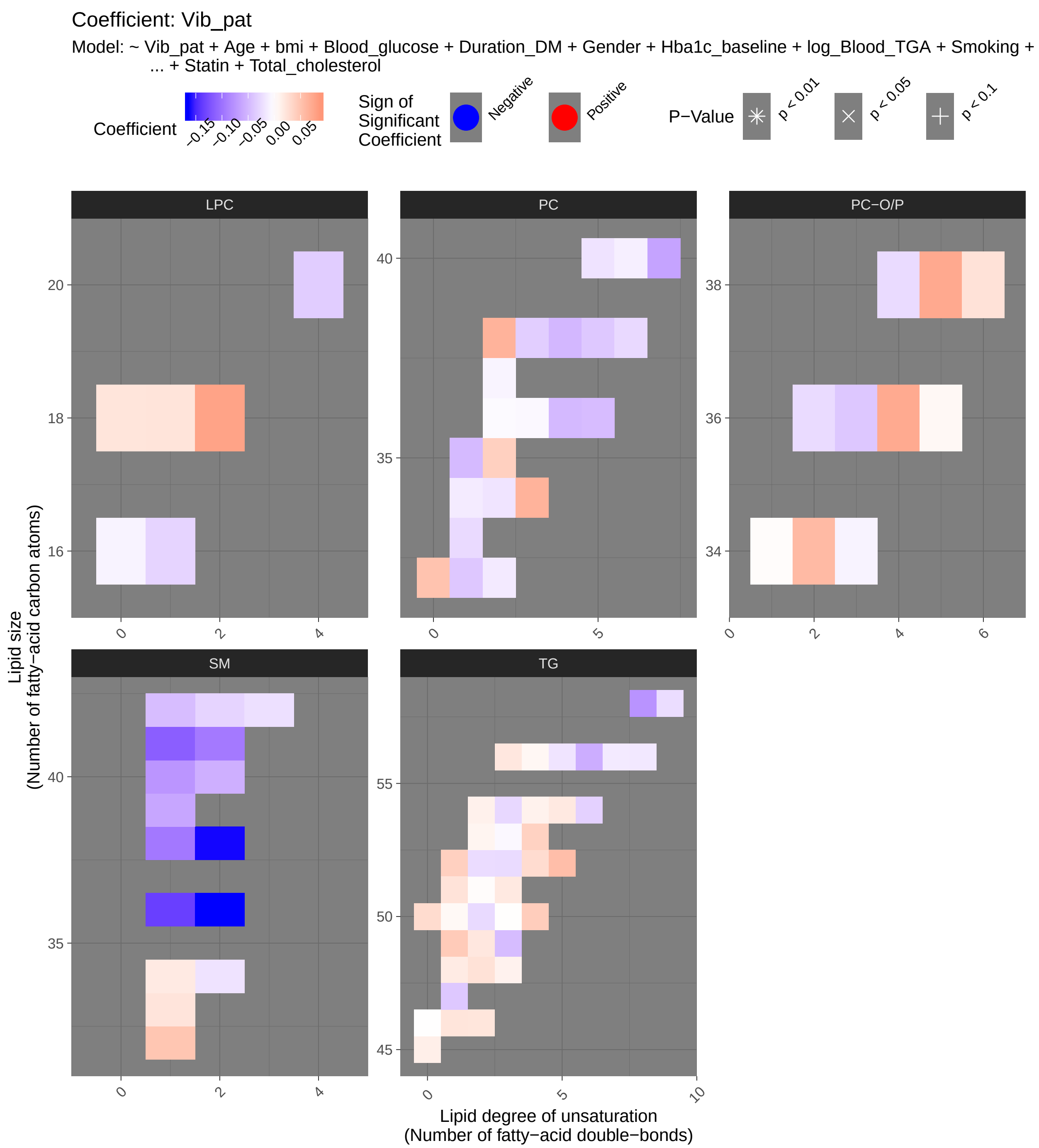
Coefficient: Vib_pat
Model: ~ Vib_pat + Age + bmi + Blood_glucose + Duration_DM + Gender + Hba1c_baseline + log_Blood_TGA + Smoking + ... + Statin + Total_cholesterol
Coefficient
−0.15 −0.10 −0.05 0.00 0.05
Sign of Significant Coefficient
Negative
Positive
P−Value
p < 0.01
p < 0.05
p < 0.1
LPC
PC
PC−O/P
SM
TG
Lipid size
(Number of fatty−acid carbon atoms)
Lipid degree of unsaturation
(Number of fatty−acid double−bonds)

##### 6.2.2 Tables of Model Coefficients

```
## [1] ""
## [1] "Table: Vib_pat"
## [1] "  (from model: "
## [1] "   ~ Vib_pat + Age + bmi + Blood_glucose + Duration_DM +"
## [1] "     Gender + Hba1c_baseline + log_Blood_TGA + Smoking + Statin +"
## [1] "     Total_cholesterol)"
## [1] ""
##                               Name Coefficient P.Value adj.P.Val
## 1           SM(d38:2)_[LVL2]; 151   -0.162000 0.00254     0.137
## 2           SM(d36:2)_[LVL2]; 160   -0.163000 0.00258     0.137
## 3            SM(d36:1)_[LVL2]; 55   -0.135000 0.01070     0.377
## 4           SM(d41:1)_[LVL2]; 102   -0.114000 0.03150     0.834
## 5            SM(d38:1)_[LVL2]; 67   -0.095200 0.06740     0.959
## 6   TG(18:2/18:1/16:0)_[LVL2]; 500   -0.106000 0.06840     0.959
## 7           SM(d41:2)_[LVL2]; 139   -0.094900 0.07200     0.959
## 8   TG(16:0/18:2/18:3)_[LVL2]; 106    0.087100 0.07240     0.959
## 9            SM(d40:1)_[LVL2]; 39   -0.074400 0.13300     0.959
## 10   TG(18:1/18:1/18:1)_[LVL2]; 15   -0.060900 0.16100     0.959
## 11  TG(18:1/18:1/22:6)_[LVL2]; 147   -0.075800 0.19600     0.959
## 12            LPC(18:2)_[LVL2]; 33    0.077600 0.20100     0.959
## 13            PC(40:7)_[LVL2]; 165   -0.064800 0.23200     0.959
## 14            TG(56:6)_[LVL3]; 275   -0.057600 0.24200     0.959
## 15          PC(O-38:5)_[LVL2]; 76    0.072300 0.24400     0.959
## 16            TG(52:5)_[LVL3]; 286    0.055400 0.25000     0.959
## 17          PC(O-36:4)_[LVL2]; 71    0.071700 0.25000     0.959
## 18            PC(38:2)_[LVL2]; 197    0.064400 0.25300     0.959
## 19           SM(d40:2)_[LVL2]; 80   -0.056000 0.26000     0.959
## 20          SM(d39:1)_[LVL2]; 179   -0.062500 0.26700     0.959
## 21            PC(34:3)_[LVL2]; 113    0.064300 0.26900     0.959
## 22            TG(49:3)_[LVL3]; 218   -0.046900 0.32000     0.959
## 23  TG(14:0/18:2/18:2)_[LVL2]; 189    0.042800 0.32400     0.959
## 24          SM(d32:1)_[LVL2]; 105    0.049000 0.32800     0.959
## 25         PC(O-34:2)_[LVL2]; 171    0.058500 0.33000     0.959
## 26   TG(16:0/18:0/18:1)_[LVL2]; 51    0.040100 0.35200     0.959
## 27       SM(d18:1/24:0)_[LVL2]; 61   -0.046400 0.35900     0.959
## 28             PC(32:0)_[LVL2]; 96    0.050800 0.36000     0.959
## 29            TG(49:1)_[LVL3]; 187    0.044200 0.38700     0.959
## 30            PC(35:1)_[LVL2]; 178   -0.048200 0.38700     0.959
## 31             PC(36:5)_[LVL2]; 23   -0.046500 0.39500     0.959
## 32              PC(38:4)_[LVL2]; 9   -0.050300 0.39500     0.959
## 33            TG(53:4)_[LVL3]; 314    0.038400 0.40600     0.959
## 34              PC(36:4)_[LVL2]; 1   -0.049000 0.43800     0.959
## 35         PC(O-36:3)_[LVL2]; 268   -0.039200 0.45700     0.959
## 36             TG(52:2)_[LVL3]; 97   -0.024400 0.48500     0.959
## 37            TG(52:4)_[LVL3]; 157    0.029700 0.48800     0.959
## 38  TG(18:0/18:1/20:4)_[LVL2]; 141   -0.039500 0.49000     0.959
## 39             PC(38:5)_[LVL2]; 24   -0.038400 0.49000     0.959
## 40            TG(50:2)_[LVL3]; 167   -0.025800 0.49000     0.959
## 41            PC(35:2)_[LVL2]; 143    0.039800 0.51000     0.959
## 42             PC(32:1)_[LVL2]; 44   -0.039200 0.51100     0.959
## 43            TG(52:3)_[LVL3]; 101   -0.025300 0.52400     0.959
## 44            TG(54:3)_[LVL3]; 124   -0.027300 0.52700     0.959
```

```
## 45           TG(47:1)_[LVL3]; 227   -0.038300 0.54000     0.959
## 46           TG(54:6)_[LVL3]; 316   -0.031700 0.54500     0.959
## 47            PC(38:3)_[LVL2]; 29   -0.034300 0.54900     0.959
## 48          LPC(20:4)_[LVL2]; 120   -0.035000 0.56300     0.959
## 49            SM(42:2)_[LVL2]; 14   -0.030100 0.57400     0.959
## 50           TG(50:0)_[LVL2]; 159    0.030200 0.57600     0.959
## 51  TG(16:0/22:5/18:1) or TG(20:4/   -0.025500 0.57600     0.959
## 52  TG(18:1/12:0/18:1) or TG(18:2/    0.024900 0.57800     0.959
## 53           TG(51:1)_[LVL3]; 249    0.023700 0.59800     0.959
## 54   TG(18:2/22:5/16:0)_[LVL2]; 69   -0.028600 0.59900     0.959
## 55             PC(38:6)_[LVL2]; 8   -0.026200 0.62500     0.959
## 56          LPC(16:1)_[LVL2]; 258   -0.030000 0.62900     0.959
## 57           TG(51:3)_[LVL3]; 198    0.018800 0.65000     0.959
## 58           PC(33:1)_[LVL2]; 177   -0.025500 0.65100     0.959
## 59         PC(O-36:2)_[LVL2]; 312   -0.025200 0.65800     0.959
## 60   TG(14:0/18:1/18:1)_[LVL2]; 25   -0.015600 0.66100     0.959
## 61         PC(O-38:6)_[LVL2]; 236    0.024600 0.66200     0.959
## 62  SM(d16:1/18:1) or SM(d18:2/16:   -0.019800 0.66800     0.959
## 63          SM(d33:1)_[LVL2]; 166    0.022700 0.67500     0.959
## 64           TG(46:1)_[LVL3]; 128    0.022500 0.67500     0.959
## 65           TG(56:3)_[LVL2]; 290    0.020300 0.67800     0.959
## 66           TG(56:5)_[LVL2]; 230   -0.018700 0.68000     0.959
## 67           TG(46:2)_[LVL3]; 248    0.021800 0.68000     0.959
## 68           TG(49:2)_[LVL3]; 231    0.020300 0.68500     0.959
## 69       SM(d18:2/24:1)_[LVL2]; 40   -0.021500 0.68500     0.959
## 70         PC(O-38:4)_[LVL2]; 131   -0.024800 0.68700     0.959
## 71           LPC(18:1)_[LVL2]; 34    0.023400 0.69900     0.959
## 72           TG(58:9)_[LVL3]; 207   -0.023500 0.70000     0.959
## 73           LPC(18:0)_[LVL1]; 22    0.022600 0.70500     0.959
## 74            TG(45:0)_[LVL2]; 65    0.013900 0.70600     0.959
## 75   TG(14:0/16:0/18:1)_[LVL2]; 54    0.017000 0.71200     0.959
## 76   TG(18:2/18:1/18:1)_[LVL2]; 20   -0.017100 0.71600     0.959
## 77            PC(40:5)_[LVL2]; 95   -0.019100 0.71800     0.959
## 78           TG(54:5)_[LVL3]; 240    0.018900 0.72100     0.959
## 79   TG(16:0/18:2/18:2)_[LVL2]; 27    0.013700 0.73200     0.959
## 80            TG(54:2)_[LVL3]; 52    0.011700 0.76200     0.959
## 81             PC(34:2)_[LVL2]; 4   -0.018700 0.76600     0.959
## 82           SM(d34:1)_[LVL2]; 26    0.017600 0.76900     0.959
## 83   TG(18:1/18:2/18:2)_[LVL2]; 57    0.014600 0.77100     0.959
## 84  TG(16:0/18:2/22:6)_[LVL2]; 117   -0.016100 0.77800     0.959
## 85  TG(18:2/18:2/18:2) or TG(18:3/   -0.013400 0.79200     0.959
## 86           TG(56:7)_[LVL3]; 309   -0.014600 0.79400     0.959
## 87           PC(32:2)_[LVL2]; 204   -0.014400 0.80700     0.959
## 88           TG(48:3)_[LVL3]; 384    0.010800 0.82200     0.959
## 89           TG(53:2)_[LVL2]; 234    0.008630 0.82400     0.959
## 90           TG(54:4)_[LVL3]; 129    0.011000 0.82500     0.959
## 91            PC(40:6)_[LVL2]; 31   -0.011200 0.83100     0.959
## 92             PC(34:1)_[LVL2]; 2   -0.013800 0.83200     0.959
## 93         PC(O-34:3)_[LVL2]; 140   -0.008170 0.87900     0.973
## 94           TG(56:4)_[LVL3]; 278    0.006900 0.88000     0.973
## 95            LPC(16:0)_[LVL1]; 5   -0.008230 0.88800     0.973
## 96            TG(50:1)_[LVL3]; 19    0.005330 0.88800     0.973
## 97           PC(37:2)_[LVL2]; 350   -0.007940 0.89000     0.973
## 98           TG(53:3)_[LVL3]; 239   -0.004930 0.90300     0.975
```

```
## 99    TG(18:1/18:1/16:0)_[LVL2]; 7    0.005730 0.91300     0.975
## 100          PC(O-36:5)_[LVL2]; 92    0.006100 0.92000     0.975
## 101            PC(36:3)_[LVL2]; 10   -0.004670 0.93600     0.979
## 102           TG(51:2)_[LVL2]; 123    0.002750 0.94200     0.979
## 103             PC(36:2)_[LVL2]; 3   -0.003390 0.95800     0.981
## 104 PC(16:0e/18:1(9Z))_[LVL1]; 134    0.002610 0.96200     0.981
## 105            TG(50:3)_[LVL2]; 47    0.001040 0.97600     0.986
## 106           TG(46:0)_[LVL3]; 168    0.000523 0.99300     0.993
```

#### 6.3 Fully-Adjusted Model

```
## [1] "Fitting models:"
## [1] "~ Vib_pat + Age + bmi + Blood_glucose + Duration_DM + Gender + Hba1c_baseline + log_Blood_TGA +
## [1] ""
```

##### 6.3.1 Heatmap

```
## [1] "heatmap_lipidome_from_limma was created by Tommi Suvitaival"
## [1] ""
## [1] "2019-05-21"
```

```
## Warning: Removed 106 rows containing missing values (geom_point).
```

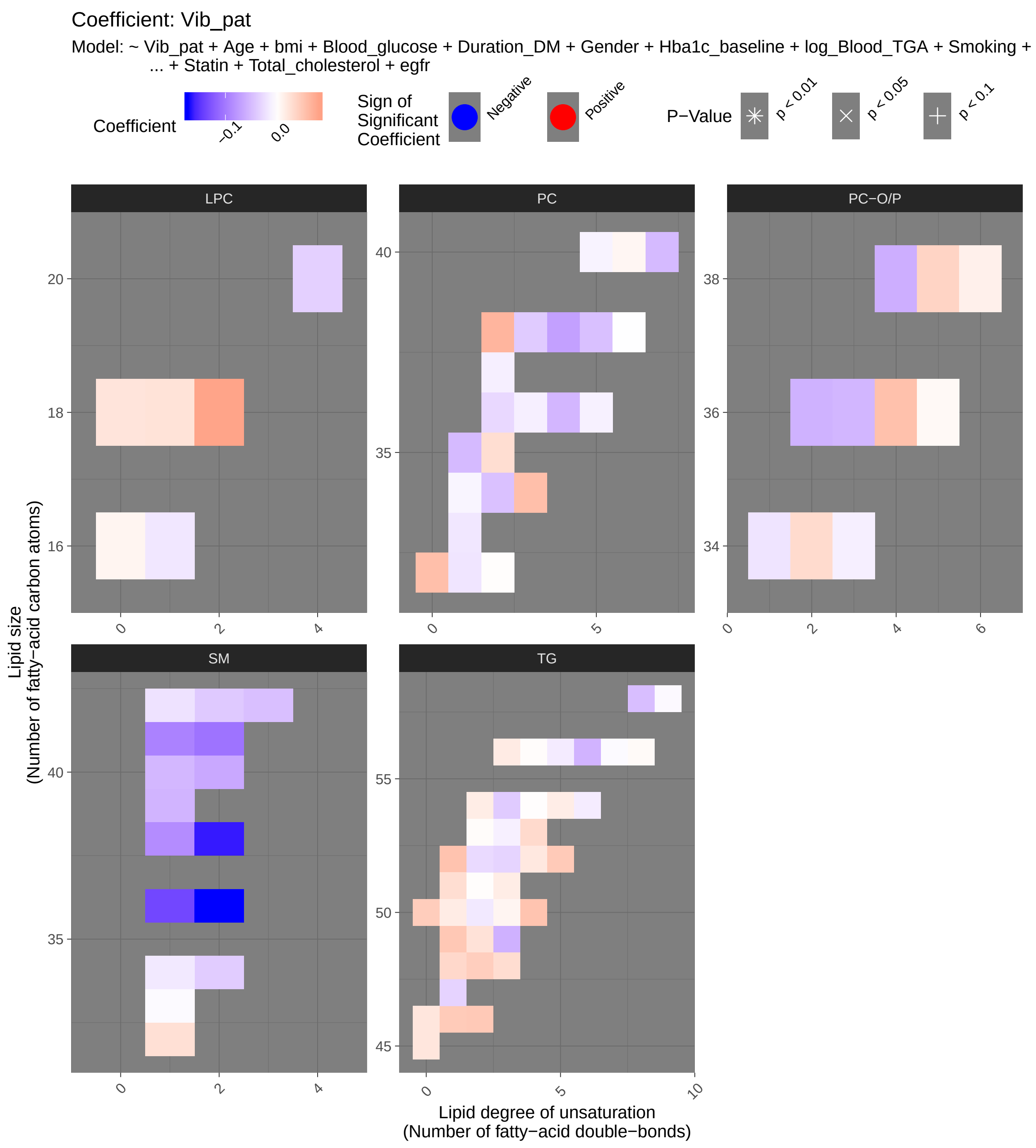

Coefficient: Vib_pat
Model: ~ Vib_pat + Age + bmi + Blood_glucose + Duration_DM + Gender + Hba1c_baseline + log_Blood_TGA + Smoking + ... + Statin + Total_cholesterol + egfr
Coefficient
−0.1
0.0
Sign of Significant Coefficient
Negative
Positive
P−Value
p < 0.01
p < 0.05
p < 0.1
LPC
PC
PC−O/P
SM
TG
Lipid size (Number of fatty−acid carbon atoms)
Lipid degree of unsaturation (Number of fatty−acid double−bonds)

##### 6.3.2 Tables of Model Coefficients

```
## [1] ""
## [1] "Table: Vib_pat"
## [1] "  (from model: "
## [1] "    ~ Vib_pat + Age + bmi + Blood_glucose + Duration_DM +"
## [1] "      Gender + Hba1c_baseline + log_Blood_TGA + Smoking + Statin +"
## [1] "      Total_cholesterol + egfr)"
## [1] ""
##                               Name Coefficient P.Value adj.P.Val
## 1            SM(d36:2)_[LVL2]; 160    -0.17300 0.00182     0.142
## 2            SM(d38:2)_[LVL2]; 151    -0.16500 0.00267     0.142
## 3             SM(d36:1)_[LVL2]; 55    -0.13700 0.01100     0.390
## 4            SM(d41:2)_[LVL2]; 139    -0.10500 0.05200     0.987
## 5   TG(18:2/18:1/16:0)_[LVL2]; 500    -0.10900 0.06520     0.987
## 6            SM(d41:1)_[LVL2]; 102    -0.09320 0.08220     0.987
## 7   TG(16:0/18:2/18:3)_[LVL2]; 106     0.08300 0.09390     0.987
## 8             SM(d38:1)_[LVL2]; 67    -0.08540 0.10800     0.987
## 9    TG(18:1/18:1/18:1)_[LVL2]; 15    -0.06510 0.14300     0.987
## 10            LPC(18:2)_[LVL2]; 33     0.08110 0.19100     0.987
## 11            SM(d40:2)_[LVL2]; 80    -0.06460 0.20300     0.987
## 12   TG(16:0/18:0/18:1)_[LVL2]; 51     0.05410 0.21800     0.987
## 13            TG(49:3)_[LVL3]; 218    -0.05780 0.22900     0.987
## 14  TG(14:0/18:2/18:2)_[LVL2]; 189     0.05310 0.23100     0.987
## 15              PC(38:4)_[LVL2]; 9    -0.07030 0.24300     0.987
## 16            PC(38:2)_[LVL2]; 197     0.06610 0.25100     0.987
## 17            TG(56:6)_[LVL3]; 275    -0.05590 0.26700     0.987
## 18            SM(d40:1)_[LVL2]; 39    -0.05300 0.29200     0.987
## 19             PC(32:0)_[LVL2]; 96     0.05750 0.31000     0.987
## 20          PC(O-36:3)_[LVL2]; 268    -0.05420 0.31300     0.987
## 21          PC(O-36:2)_[LVL2]; 312    -0.05740 0.31900     0.987
## 22            TG(52:5)_[LVL3]; 286     0.04810 0.32700     0.987
## 23          PC(O-38:4)_[LVL2]; 131    -0.06020 0.33300     0.987
## 24  TG(18:1/12:0/18:1) or TG(18:2/     0.04390 0.33500     0.987
## 25            PC(34:3)_[LVL2]; 113     0.05720 0.33500     0.987
## 26           SM(d39:1)_[LVL2]; 179    -0.05530 0.33600     0.987
## 27            TG(49:1)_[LVL3]; 187     0.04990 0.33900     0.987
## 28            PC(40:7)_[LVL2]; 165    -0.05120 0.35400     0.987
## 29            TG(46:2)_[LVL3]; 248     0.04830 0.36600     0.987
## 30            PC(35:1)_[LVL2]; 178    -0.05130 0.36900     0.987
## 31           PC(O-36:4)_[LVL2]; 71     0.05590 0.37900     0.987
## 32       SM(d18:2/24:1)_[LVL2]; 40    -0.04710 0.38100     0.987
## 33            TG(46:1)_[LVL3]; 128     0.04730 0.38600     0.987
## 34            TG(54:3)_[LVL3]; 124    -0.03800 0.38900     0.987
## 35              PC(36:4)_[LVL2]; 1    -0.05380 0.40400     0.987
## 36            TG(50:0)_[LVL2]; 159     0.04530 0.41000     0.987
## 37             PC(38:5)_[LVL2]; 24    -0.04620 0.41600     0.987
## 38  TG(18:1/18:1/22:6)_[LVL2]; 147    -0.04820 0.41700     0.987
## 39  SM(d16:1/18:1) or SM(d18:2/16:    -0.03720 0.42800     0.987
## 40            TG(52:3)_[LVL3]; 101    -0.03170 0.43500     0.987
## 41             TG(52:2)_[LVL3]; 97    -0.02710 0.44700     0.987
## 42  TG(18:0/18:1/20:4)_[LVL2]; 141    -0.04370 0.45500     0.987
## 43   TG(14:0/16:0/18:1)_[LVL2]; 54     0.03490 0.45700     0.987
## 44             SM(42:2)_[LVL2]; 14    -0.04000 0.46400     0.987
```

```
## 45              PC(34:2)_[LVL2]; 4     -0.04550 0.47600      0.987
## 46            TG(53:4)_[LVL3]; 314      0.03310 0.48300      0.987
## 47            TG(51:1)_[LVL3]; 249      0.03030 0.51100      0.987
## 48             PC(38:3)_[LVL2]; 29     -0.03840 0.51100      0.987
## 49            TG(48:3)_[LVL3]; 384      0.03070 0.52900      0.987
## 50           PC(O-38:5)_[LVL2]; 76      0.03880 0.53600      0.987
## 51             TG(45:0)_[LVL2]; 65      0.02280 0.54600      0.987
## 52           SM(d32:1)_[LVL2]; 105      0.02820 0.57900      0.987
## 53           LPC(20:4)_[LVL2]; 120     -0.03420 0.57900      0.987
## 54          PC(O-34:2)_[LVL2]; 171      0.03270 0.59200      0.987
## 55            TG(49:2)_[LVL3]; 231      0.02620 0.60900      0.987
## 56            TG(47:1)_[LVL3]; 227     -0.03200 0.61600      0.987
## 57   TG(18:2/18:1/18:1)_[LVL2]; 20     -0.02390 0.61900      0.987
## 58            PC(35:2)_[LVL2]; 143      0.02970 0.63000      0.987
## 59            TG(52:4)_[LVL3]; 157      0.02070 0.63600      0.987
## 60             TG(50:1)_[LVL3]; 19      0.01720 0.65700      0.987
## 61  TG(16:0/22:5/18:1) or TG(20:4/     -0.02060 0.65800      0.987
## 62              PC(36:2)_[LVL2]; 3     -0.02830 0.66500      0.987
## 63            TG(50:2)_[LVL3]; 167     -0.01620 0.67100      0.987
## 64             TG(54:2)_[LVL3]; 52      0.01650 0.67600      0.987
## 65       SM(d18:1/24:0)_[LVL2]; 61     -0.02140 0.67600      0.987
## 66            LPC(18:1)_[LVL2]; 34      0.02580 0.67700      0.987
## 67            TG(46:0)_[LVL3]; 168      0.02260 0.69000      0.987
## 68            TG(51:3)_[LVL3]; 198      0.01670 0.69200      0.987
## 69            LPC(18:0)_[LVL1]; 22      0.02400 0.69300      0.987
## 70            TG(56:3)_[LVL2]; 290      0.01800 0.71900      0.987
## 71  PC(16:0e/18:1(9Z))_[LVL1]; 134     -0.02000 0.72000      0.987
## 72   TG(18:1/18:2/18:2)_[LVL2]; 57      0.01670 0.74500      0.987
## 73             PC(32:1)_[LVL2]; 44     -0.01930 0.75100      0.987
## 74            TG(56:5)_[LVL2]; 230     -0.01430 0.75700      0.987
## 75            PC(33:1)_[LVL2]; 177     -0.01760 0.75900      0.987
## 76            TG(54:5)_[LVL3]; 240      0.01600 0.76700      0.987
## 77           LPC(16:1)_[LVL2]; 258     -0.01790 0.77800      0.987
## 78   TG(16:0/18:2/18:2)_[LVL2]; 27      0.01120 0.78400      0.987
## 79            SM(d34:1)_[LVL2]; 26     -0.01630 0.78800      0.987
## 80            TG(53:3)_[LVL3]; 239     -0.01070 0.79600      0.987
## 81            TG(54:6)_[LVL3]; 316     -0.01360 0.79900      0.987
## 82             TG(50:3)_[LVL2]; 47      0.00884 0.80400      0.987
## 83          PC(O-38:6)_[LVL2]; 236      0.01350 0.81500      0.987
## 84   TG(18:2/22:5/16:0)_[LVL2]; 69     -0.01250 0.82100      0.987
## 85          PC(O-34:3)_[LVL2]; 140     -0.01160 0.83200      0.987
## 86            PC(37:2)_[LVL2]; 350     -0.01180 0.84000      0.987
## 87             PC(36:3)_[LVL2]; 10     -0.01170 0.84500      0.987
## 88             PC(36:5)_[LVL2]; 23     -0.01010 0.85400      0.987
## 89  TG(18:2/18:2/18:2) or TG(18:3/      0.00883 0.86400      0.987
## 90             PC(40:5)_[LVL2]; 95     -0.00908 0.86700      0.987
## 91             LPC(16:0)_[LVL1]; 5      0.00952 0.87300      0.987
## 92   TG(14:0/18:1/18:1)_[LVL2]; 25     -0.00567 0.87600      0.987
## 93             PC(40:6)_[LVL2]; 31      0.00817 0.87800      0.987
## 94              PC(34:1)_[LVL2]; 2     -0.00726 0.91300      0.987
## 95           PC(O-36:5)_[LVL2]; 92      0.00577 0.92600      0.987
## 96  TG(16:0/18:2/22:6)_[LVL2]; 117      0.00479 0.93400      0.987
## 97            TG(53:2)_[LVL2]; 234      0.00299 0.94000      0.987
## 98            TG(58:9)_[LVL3]; 207     -0.00442 0.94300      0.987
```

```
## 99           SM(d33:1)_[LVL2]; 166    -0.00357 0.94800     0.987
## 100           TG(56:7)_[LVL3]; 309    -0.00358 0.95000     0.987
## 101           TG(56:4)_[LVL3]; 278     0.00271 0.95400     0.987
## 102           TG(51:2)_[LVL2]; 123     0.00219 0.95500     0.987
## 103           TG(54:4)_[LVL3]; 129     0.00162 0.97500     0.987
## 104   TG(18:1/18:1/16:0)_[LVL2]; 7     0.00153 0.97700     0.987
## 105           PC(32:2)_[LVL2]; 204     0.00171 0.97700     0.987
## 106             PC(38:6)_[LVL2]; 8    -0.00071 0.99000     0.990
```

### 7 Secondary Analyses

#### 7.1 Resting HR Vagus

##### 7.1.1 Crude Model

```
## [1] "Fitting models:"
## [1] "~ rest_HR_vag"
## [1] ""
```

###### 7.1.1.1 Heatmap

```
## [1] "heatmap_lipidome_from_limma was created by Tommi Suvitaival"
## [1] ""
## [1] "2019-05-21"
```

```
## Warning: Removed 105 rows containing missing values (geom_point).
```

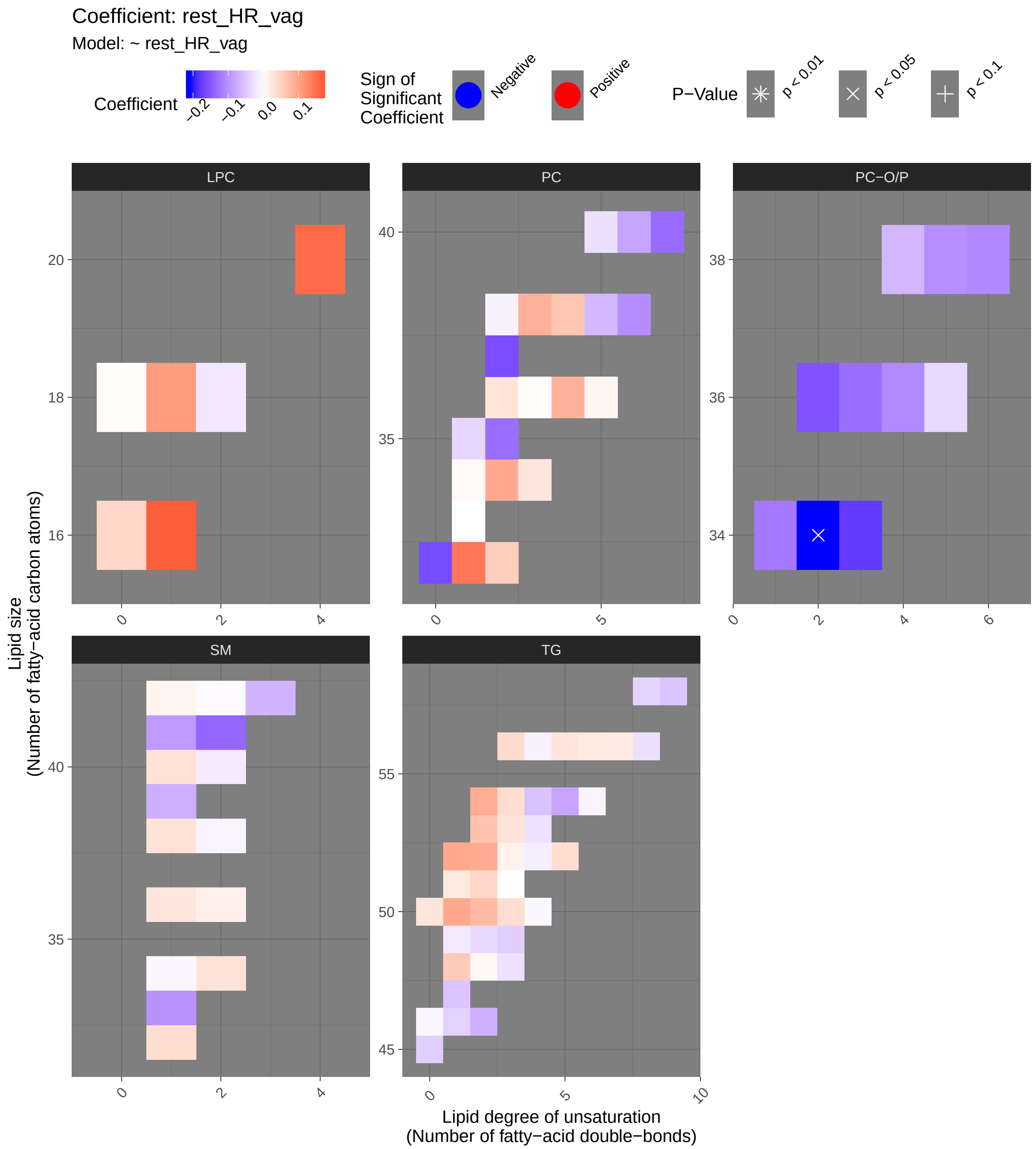
Coefficient: rest_HR_vag
Model: ~ rest_HR_vag
Coefficient
−0.2
−0.1
0.0
0.1
Sign of Significant Coefficient
Negative
Positive
P−Value
p < 0.01
p < 0.05
p < 0.1
LPC
PC
PC−O/P
SM
TG
Lipid size (Number of fatty−acid carbon atoms)
Lipid degree of unsaturation (Number of fatty−acid double−bonds)

###### 7.1.1.2 Tables of Model Coefficients

```
## [1] ""
## [1] "Table: rest_HR_vag"
## [1] "  (from model: "
## [1] "   ~ rest_HR_vag)"
## [1] ""
##                       Name Coefficient  P.Value adj.P.Val
## 1 PC(O-34:2)_[LVL2]; 171      -0.209 0.000337    0.0357
```

###### 7.1.1.3 Forest Plot of Model Coefficients

```
## Warning: Ignoring unknown aesthetics: x
```

```
## NULL
```

###### 7.1.2 Adjusted Model

```
## [1] "Fitting models:"
## [1] "~ rest_HR_vag + Age + bmi + Blood_glucose + Duration_DM + Gender + Hba1c_baseline + log_Blood_T
## [1] ""
```

###### 7.1.2.1 Heatmap

```
## [1] "heatmap_lipidome_from_limma was created by Tommi Suvitaival"
## [1] ""
## [1] "2019-05-21"
```

```
## Warning: Removed 92 rows containing missing values (geom_point).
```

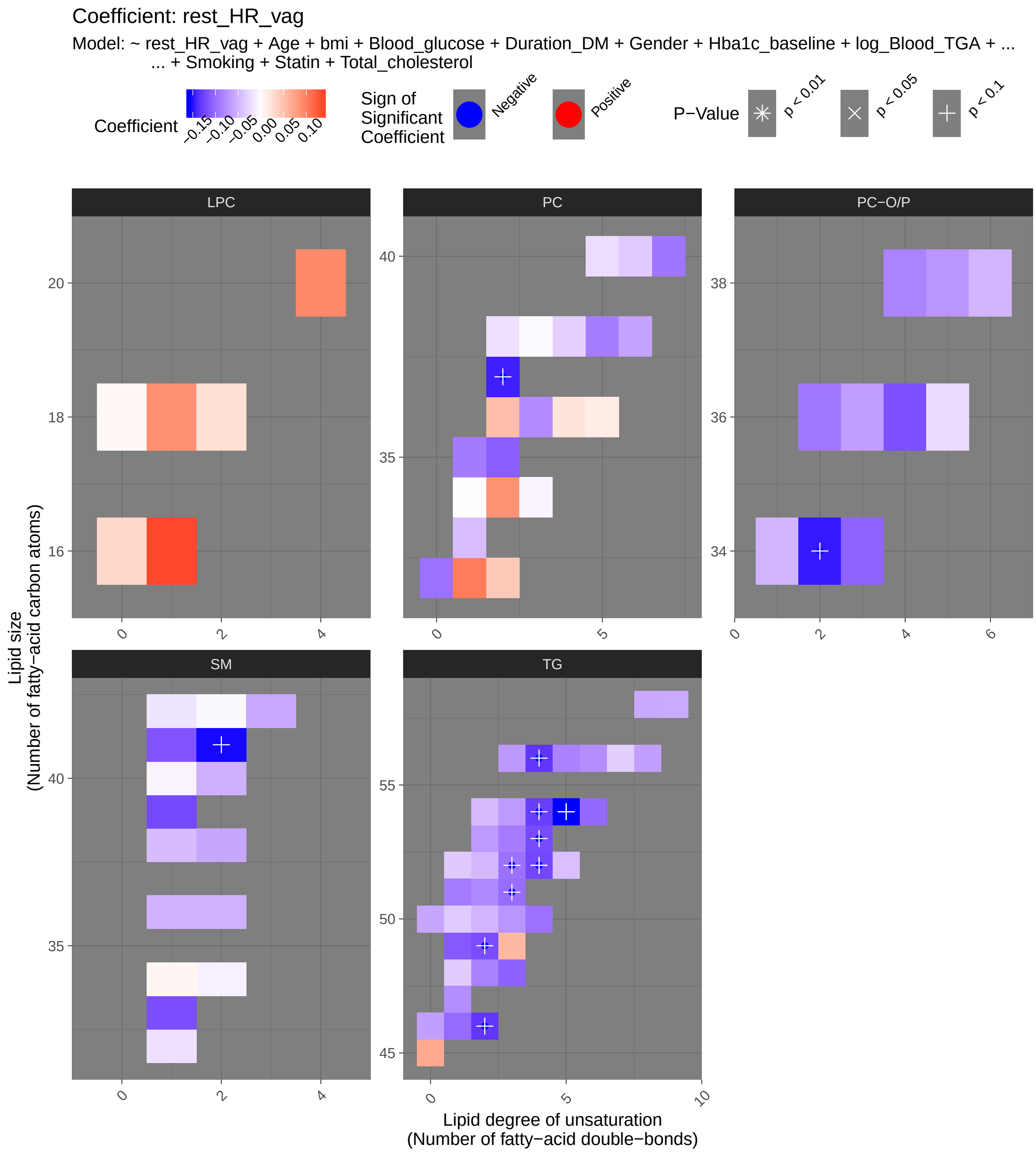

Coefficient: rest_HR_vag
Model: ~ rest_HR_vag + Age + bmi + Blood_glucose + Duration_DM + Gender + Hba1c_baseline + log_Blood_TGA + ...
... + Smoking + Statin + Total_cholesterol
Coefficient
−0.15 −0.10 −0.05 0.00 0.05 0.10
Sign of Significant Coefficient
Negative
Positive
P−Value
p < 0.01
p < 0.05
p < 0.1
LPC
PC
PC−O/P
SM
TG
Lipid size
(Number of fatty−acid carbon atoms)
Lipid degree of unsaturation
(Number of fatty−acid double−bonds)

###### 7.1.2.2 Tables of Model Coefficients

```
## [1] ""
## [1] "Table: rest_HR_vag"
## [1] "  (from model: "
## [1] "    ~ rest_HR_vag + Age + bmi + Blood_glucose + Duration_DM +"
## [1] "      Gender + Hba1c_baseline + log_Blood_TGA + Smoking + Statin +"
## [1] "      Total_cholesterol)"
## [1] ""
##                               Name Coefficient P.Value adj.P.Val
## 1             TG(54:5)_[LVL3]; 240    -0.15600 0.00178    0.0648
## 2             TG(56:4)_[LVL3]; 278    -0.13400 0.00198    0.0648
## 3           SM(d41:2)_[LVL2]; 139    -0.15400 0.00215    0.0648
## 4             TG(52:4)_[LVL3]; 157    -0.12300 0.00244    0.0648
## 5    TG(18:1/18:2/18:2)_[LVL2]; 57    -0.13600 0.00418    0.0885
## 6             TG(53:4)_[LVL3]; 314    -0.12100 0.00577    0.0908
## 7             TG(54:4)_[LVL3]; 129    -0.12900 0.00635    0.0908
## 8             TG(46:2)_[LVL3]; 248    -0.13400 0.00749    0.0908
## 9             PC(37:2)_[LVL2]; 350    -0.14600 0.00781    0.0908
## 10          PC(O-34:2)_[LVL2]; 171    -0.14900 0.00949    0.0908
## 11            TG(51:3)_[LVL3]; 198    -0.09920 0.01120    0.0908
## 12            TG(49:2)_[LVL3]; 231    -0.12000 0.01130    0.0908
## 13            TG(52:3)_[LVL3]; 101    -0.09580 0.01140    0.0908
## 14   TG(16:0/18:2/18:2)_[LVL2]; 27    -0.09580 0.01200    0.0908
## 15  TG(14:0/18:2/18:2)_[LVL2]; 189    -0.09600 0.02010    0.1110
## 16            TG(48:3)_[LVL3]; 384    -0.10600 0.02030    0.1110
## 17           SM(d33:1)_[LVL2]; 166    -0.11900 0.02050    0.1110
## 18            TG(53:3)_[LVL3]; 239    -0.08880 0.02060    0.1110
## 19            TG(49:1)_[LVL3]; 187    -0.11200 0.02090    0.1110
## 20           SM(d39:1)_[LVL2]; 179    -0.12300 0.02200    0.1110
## 21           SM(d41:1)_[LVL2]; 102    -0.11500 0.02210    0.1110
## 22           LPC(16:1)_[LVL2]; 258     0.13500 0.02300    0.1110
## 23            TG(51:2)_[LVL2]; 123    -0.07990 0.02510    0.1160
## 24             TG(50:3)_[LVL2]; 47    -0.07160 0.03110    0.1370
## 25          PC(O-34:3)_[LVL2]; 140    -0.10600 0.03870    0.1530
## 26  TG(16:0/18:2/18:3)_[LVL2]; 106    -0.09510 0.04000    0.1530
## 27            TG(51:1)_[LVL3]; 249    -0.08810 0.04000    0.1530
## 28            TG(54:6)_[LVL3]; 316    -0.10100 0.04160    0.1530
## 29   TG(18:2/18:1/18:1)_[LVL2]; 20    -0.08920 0.04590    0.1530
## 30  TG(18:1/12:0/18:1) or TG(18:2/    -0.08510 0.04600    0.1530
## 31           PC(O-36:4)_[LVL2]; 71    -0.11800 0.04720    0.1530
## 32            TG(46:1)_[LVL3]; 128    -0.10100 0.04800    0.1530
## 33  TG(16:0/22:5/18:1) or TG(20:4/    -0.08560 0.04870    0.1530
## 34            TG(56:5)_[LVL2]; 230    -0.08450 0.05020    0.1530
## 35             TG(45:0)_[LVL2]; 65     0.06890 0.05040    0.1530
## 36            PC(35:2)_[LVL2]; 143    -0.10900 0.05820    0.1710
## 37            TG(53:2)_[LVL2]; 234    -0.06840 0.06470    0.1850
## 38             PC(32:0)_[LVL2]; 96    -0.09600 0.06860    0.1910
## 39            PC(40:7)_[LVL2]; 165    -0.09290 0.07220    0.1930
## 40             PC(32:1)_[LVL2]; 44     0.10300 0.07270    0.1930
## 41          PC(O-36:2)_[LVL2]; 312    -0.09150 0.09320    0.2310
## 42            PC(35:1)_[LVL2]; 178    -0.08970 0.09330    0.2310
## 43             PC(38:5)_[LVL2]; 24    -0.08830 0.09450    0.2310
## 44            TG(54:3)_[LVL3]; 124    -0.06780 0.09570    0.2310
```

```
## 45           TG(56:6)_[LVL3]; 275    -0.07640 0.10300    0.2440
## 46          LPC(20:4)_[LVL2]; 120     0.09260 0.10700    0.2470
## 47           LPC(18:1)_[LVL2]; 34     0.08870 0.12500    0.2830
## 48           TG(56:3)_[LVL2]; 290    -0.06910 0.13800    0.3040
## 49             PC(34:2)_[LVL2]; 4     0.08720 0.14800    0.3200
## 50         PC(O-38:4)_[LVL2]; 131    -0.08420 0.15100    0.3200
## 51            TG(52:2)_[LVL3]; 97    -0.04710 0.15600    0.3230
## 52            PC(36:3)_[LVL2]; 10    -0.07810 0.16000    0.3270
## 53           TG(50:2)_[LVL3]; 167    -0.04970 0.16500    0.3270
## 54   TG(14:0/18:1/18:1)_[LVL2]; 25    -0.04700 0.16600    0.3270
## 55         PC(O-36:3)_[LVL2]; 268    -0.06560 0.19300    0.3680
## 56           TG(47:1)_[LVL3]; 227    -0.07690 0.19400    0.3680
## 57           TG(49:3)_[LVL3]; 218     0.05650 0.20700    0.3830
## 58            TG(54:2)_[LVL3]; 52    -0.04590 0.20900    0.3830
## 59             PC(38:6)_[LVL2]; 8    -0.06250 0.22000    0.3960
## 60           TG(46:0)_[LVL3]; 168    -0.06440 0.22500    0.3980
## 61          PC(O-38:5)_[LVL2]; 76    -0.07130 0.22900    0.3980
## 62  TG(16:0/18:2/22:6)_[LVL2]; 117    -0.06420 0.23700    0.4040
## 63           TG(50:0)_[LVL2]; 159    -0.05960 0.24900    0.4110
## 64          SM(d38:2)_[LVL2]; 151    -0.05980 0.25000    0.4110
## 65      SM(d18:2/24:1)_[LVL2]; 40    -0.05800 0.25200    0.4110
## 66           SM(d40:2)_[LVL2]; 80    -0.05270 0.26700    0.4290
## 67  TG(18:1/18:1/22:6)_[LVL2]; 147    -0.05780 0.30200    0.4780
## 68           SM(d36:1)_[LVL2]; 55    -0.05120 0.31600    0.4930
## 69          SM(d36:2)_[LVL2]; 160    -0.05180 0.32500    0.5000
## 70           TG(58:9)_[LVL3]; 207    -0.05660 0.33000    0.5000
## 71            TG(50:1)_[LVL3]; 19    -0.03440 0.34200    0.5110
## 72  PC(16:0e/18:1(9Z))_[LVL1]; 134    -0.04970 0.34700    0.5110
## 73           TG(52:5)_[LVL3]; 286    -0.04250 0.35600    0.5140
## 74         PC(O-38:6)_[LVL2]; 236    -0.04930 0.35900    0.5140
## 75           SM(d38:1)_[LVL2]; 67    -0.04460 0.37300    0.5270
## 76   TG(16:0/18:0/18:1)_[LVL2]; 51    -0.03540 0.39000    0.5440
## 77             PC(36:2)_[LVL2]; 3     0.05230 0.39500    0.5440
## 78           PC(33:1)_[LVL2]; 177    -0.04450 0.40900    0.5560
## 79   TG(14:0/16:0/18:1)_[LVL2]; 54    -0.03450 0.43400    0.5780
## 80           PC(32:2)_[LVL2]; 204     0.04390 0.43600    0.5780
## 81    TG(18:1/18:1/16:0)_[LVL2]; 7    -0.03760 0.45000    0.5890
## 82  TG(18:0/18:1/20:4)_[LVL2]; 141    -0.03970 0.46900    0.6060
## 83            PC(40:6)_[LVL2]; 31    -0.03490 0.48500    0.6190
## 84  TG(18:2/18:2/18:2) or TG(18:3/    -0.03280 0.50000    0.6310
## 85           TG(56:7)_[LVL3]; 309    -0.03200 0.54500    0.6790
## 86            LPC(16:0)_[LVL1]; 5     0.03200 0.56700    0.6980
## 87   TG(18:2/22:5/16:0)_[LVL2]; 69    -0.02930 0.57300    0.6980
## 88             PC(38:4)_[LVL2]; 9    -0.03100 0.58500    0.7040
## 89            PC(40:5)_[LVL2]; 95    -0.02230 0.65700    0.7800
## 90           LPC(18:2)_[LVL2]; 33     0.02550 0.66200    0.7800
## 91          SM(d32:1)_[LVL2]; 105    -0.01960 0.68100    0.7930
## 92          PC(O-36:5)_[LVL2]; 92    -0.02300 0.69000    0.7960
## 93           PC(38:2)_[LVL2]; 197    -0.02000 0.71000    0.8050
## 94             PC(36:4)_[LVL2]; 1     0.02180 0.71900    0.8050
## 95      SM(d18:1/24:0)_[LVL2]; 61    -0.01720 0.72100    0.8050
## 96            PC(36:5)_[LVL2]; 23     0.01550 0.76600    0.8450
## 97  SM(d16:1/18:1) or SM(d18:2/16:    -0.00954 0.82800    0.9050
## 98   TG(18:1/18:1/18:1)_[LVL2]; 15    -0.00739 0.85900    0.9290
```

```
## 99            SM(d40:1)_[LVL2]; 39    -0.00773 0.87100    0.9310
## 100 TG(18:2/18:1/16:0)_[LVL2]; 500     0.00853 0.87800    0.9310
## 101           SM(d34:1)_[LVL2]; 26     0.00740 0.89800    0.9340
## 102           PC(34:3)_[LVL2]; 113    -0.00703 0.89900    0.9340
## 103           LPC(18:0)_[LVL1]; 22     0.00559 0.92200    0.9420
## 104            SM(42:2)_[LVL2]; 14    -0.00489 0.92400    0.9420
## 105            PC(38:3)_[LVL2]; 29    -0.00329 0.95200    0.9610
## 106             PC(34:1)_[LVL2]; 2    -0.00110 0.98600    0.9860
```

###### 7.1.3 Fully-Adjusted Model

```
## [1] "Fitting models:"
## [1] "~ rest_HR_vag + Age + bmi + Blood_glucose + Duration_DM + Gender + Hba1c_baseline + log_Blood_T
## [1] ""
```

###### 7.1.3.1 Heatmap

```
## [1] "heatmap_lipidome_from_limma was created by Tommi Suvitaival"
## [1] ""
## [1] "2019-05-21"
```

```
## Warning: Removed 91 rows containing missing values (geom_point).
```

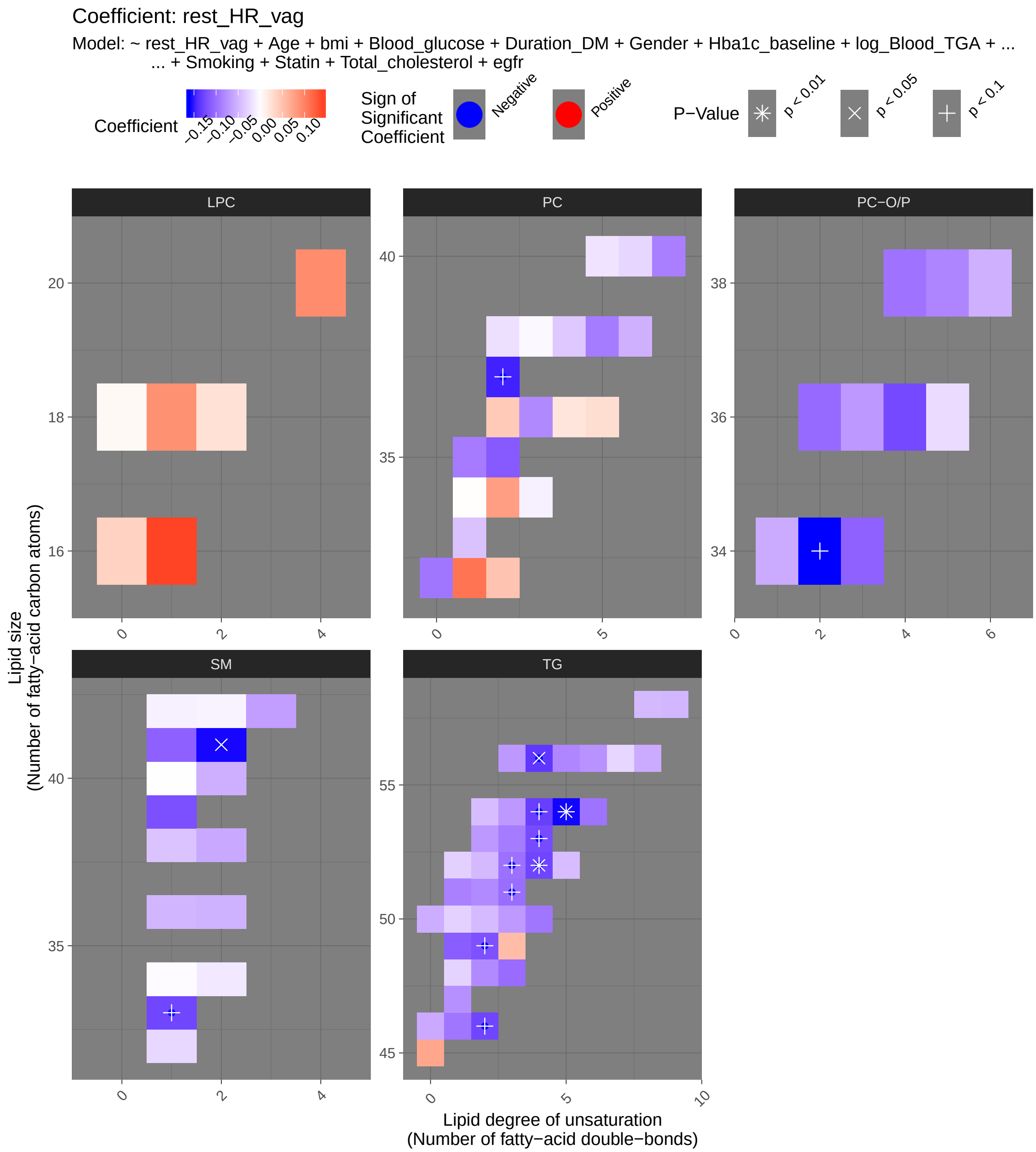
Coefficient: rest_HR_vag
Model: ~ rest_HR_vag + Age + bmi + Blood_glucose + Duration_DM + Gender + Hba1c_baseline + log_Blood_TGA + ...
... + Smoking + Statin + Total_cholesterol + egfr
Coefficient
−0.15 −0.10 −0.05 0.00 0.05 0.10
Sign of Significant Coefficient
Negative
Positive
P−Value
p < 0.01
p < 0.05
p < 0.1
LPC
PC
PC−O/P
SM
TG
Lipid size
(Number of fatty−acid carbon atoms)
Lipid degree of unsaturation
(Number of fatty−acid double−bonds)

###### 7.1.3.2 Tables of Model Coefficients

```
## [1] ""
## [1] "Table: rest_HR_vag"
## [1] "  (from model: "
## [1] "   ~ rest_HR_vag + Age + bmi + Blood_glucose + Duration_DM +"
## [1] "     Gender + Hba1c_baseline + log_Blood_TGA + Smoking + Statin +"
## [1] "     Total_cholesterol + egfr)"
## [1] ""
##                               Name Coefficient P.Value adj.P.Val
## 1              TG(54:5)_[LVL3]; 240   -0.157000 0.00164    0.0481
## 2              TG(56:4)_[LVL3]; 278   -0.136000 0.00177    0.0481
## 3              TG(52:4)_[LVL3]; 157   -0.127000 0.00180    0.0481
## 4             SM(d41:2)_[LVL2]; 139   -0.157000 0.00181    0.0481
## 5    TG(18:1/18:2/18:2)_[LVL2]; 57   -0.136000 0.00436    0.0699
## 6              TG(53:4)_[LVL3]; 314   -0.124000 0.00493    0.0699
## 7           PC(O-34:2)_[LVL2]; 171   -0.159000 0.00525    0.0699
## 8              TG(54:4)_[LVL3]; 129   -0.132000 0.00528    0.0699
## 9              PC(37:2)_[LVL2]; 350   -0.148000 0.00718    0.0846
## 10             TG(52:3)_[LVL3]; 101   -0.097800 0.01000    0.0905
## 11             TG(51:3)_[LVL3]; 198   -0.100000 0.01070    0.0905
## 12             TG(46:2)_[LVL3]; 248   -0.127000 0.01070    0.0905
## 13   TG(16:0/18:2/18:2)_[LVL2]; 27   -0.097200 0.01110    0.0905
## 14             TG(49:2)_[LVL3]; 231   -0.119000 0.01200    0.0905
## 15            SM(d33:1)_[LVL2]; 166   -0.127000 0.01310    0.0926
## 16             TG(53:3)_[LVL3]; 239   -0.090800 0.01840    0.1170
## 17            LPC(16:1)_[LVL2]; 258    0.140000 0.01880    0.1170
## 18             TG(49:1)_[LVL3]; 187   -0.112000 0.02160    0.1260
## 19  TG(14:0/18:2/18:2)_[LVL2]; 189   -0.094000 0.02330    0.1260
## 20             TG(51:2)_[LVL2]; 123   -0.080300 0.02490    0.1260
## 21            SM(d39:1)_[LVL2]; 179   -0.121000 0.02500    0.1260
## 22             TG(48:3)_[LVL3]; 384   -0.101000 0.02690    0.1300
## 23            SM(d41:1)_[LVL2]; 102   -0.109000 0.03020    0.1390
## 24  TG(16:0/18:2/18:3)_[LVL2]; 106   -0.098000 0.03480    0.1420
## 25            PC(O-36:4)_[LVL2]; 71   -0.125000 0.03560    0.1420
## 26          PC(O-34:3)_[LVL2]; 140   -0.108000 0.03560    0.1420
## 27              TG(50:3)_[LVL2]; 47   -0.069800 0.03630    0.1420
## 28              TG(45:0)_[LVL2]; 65    0.071900 0.04160    0.1540
## 29   TG(18:2/18:1/18:1)_[LVL2]; 20   -0.091000 0.04240    0.1540
## 30             TG(51:1)_[LVL3]; 249   -0.086900 0.04360    0.1540
## 31             PC(35:2)_[LVL2]; 143   -0.113000 0.04990    0.1650
## 32             TG(54:6)_[LVL3]; 316   -0.096400 0.05290    0.1650
## 33              PC(32:1)_[LVL2]; 44    0.110000 0.05410    0.1650
## 34  TG(16:0/22:5/18:1) or TG(20:4/   -0.083800 0.05420    0.1650
## 35             TG(56:5)_[LVL2]; 230   -0.082900 0.05560    0.1650
## 36          PC(O-36:2)_[LVL2]; 312   -0.103000 0.05680    0.1650
## 37             TG(53:2)_[LVL2]; 234   -0.070400 0.05810    0.1650
## 38  TG(18:1/12:0/18:1) or TG(18:2/   -0.080400 0.05900    0.1650
## 39             TG(46:1)_[LVL3]; 128   -0.095000 0.06250    0.1700
## 40              PC(32:0)_[LVL2]; 96   -0.093800 0.07590    0.2010
## 41             TG(54:3)_[LVL3]; 124   -0.070600 0.08340    0.2160
## 42              PC(38:5)_[LVL2]; 24   -0.090900 0.08610    0.2170
## 43             PC(40:7)_[LVL2]; 165   -0.087900 0.08890    0.2190
## 44             PC(35:1)_[LVL2]; 178   -0.090600 0.09140    0.2200
```

```
## 45          PC(O-38:4)_[LVL2]; 131   -0.095900 0.09850    0.2320
## 46           LPC(20:4)_[LVL2]; 120    0.092800 0.10800    0.2490
## 47            TG(56:6)_[LVL3]; 275   -0.074900 0.11200    0.2520
## 48            LPC(18:1)_[LVL2]; 34    0.089000 0.12600    0.2770
## 49            TG(56:3)_[LVL2]; 290   -0.069900 0.13500    0.2920
## 50             PC(36:3)_[LVL2]; 10   -0.081100 0.14600    0.3090
## 51             TG(52:2)_[LVL3]; 97   -0.047800 0.15100    0.3110
## 52           PC(O-38:5)_[LVL2]; 76   -0.083700 0.15300    0.3110
## 53          PC(O-36:3)_[LVL2]; 268   -0.070500 0.16300    0.3250
## 54              PC(34:2)_[LVL2]; 4    0.079100 0.18800    0.3630
## 55       SM(d18:2/24:1)_[LVL2]; 40   -0.066200 0.18900    0.3630
## 56            TG(50:2)_[LVL3]; 167   -0.046700 0.19200    0.3630
## 57   TG(14:0/18:1/18:1)_[LVL2]; 25   -0.044000 0.19500    0.3630
## 58            TG(47:1)_[LVL3]; 227   -0.075600 0.20300    0.3720
## 59            TG(49:3)_[LVL3]; 218    0.054800 0.22200    0.3990
## 60             TG(54:2)_[LVL3]; 52   -0.044500 0.22600    0.3990
## 61            SM(d40:2)_[LVL2]; 80   -0.055100 0.24700    0.4290
## 62           SM(d38:2)_[LVL2]; 151   -0.058900 0.25900    0.4420
## 63            TG(46:0)_[LVL3]; 168   -0.058400 0.27100    0.4540
## 64  PC(16:0e/18:1(9Z))_[LVL1]; 134   -0.057500 0.27400    0.4540
## 65            TG(50:0)_[LVL2]; 159   -0.055600 0.28300    0.4600
## 66              PC(38:6)_[LVL2]; 8   -0.053500 0.29100    0.4600
## 67  TG(16:0/18:2/22:6)_[LVL2]; 117   -0.057200 0.29100    0.4600
## 68           SM(d36:2)_[LVL2]; 160   -0.052700 0.31900    0.4910
## 69          PC(O-38:6)_[LVL2]; 236   -0.053600 0.32000    0.4910
## 70            TG(52:5)_[LVL3]; 286   -0.045400 0.32600    0.4910
## 71            SM(d36:1)_[LVL2]; 55   -0.050100 0.32900    0.4910
## 72            PC(32:2)_[LVL2]; 204    0.048600 0.38900    0.5550
## 73            TG(58:9)_[LVL3]; 207   -0.049800 0.39000    0.5550
## 74             TG(50:1)_[LVL3]; 19   -0.031000 0.39200    0.5550
## 75  TG(18:1/18:1/22:6)_[LVL2]; 147   -0.047500 0.39200    0.5550
## 76            SM(d38:1)_[LVL2]; 67   -0.040700 0.41700    0.5820
## 77    TG(18:1/18:1/16:0)_[LVL2]; 7   -0.038900 0.43600    0.5960
## 78   TG(16:0/18:0/18:1)_[LVL2]; 51   -0.031700 0.44200    0.5960
## 79            PC(33:1)_[LVL2]; 177   -0.041400 0.44400    0.5960
## 80  TG(18:0/18:1/20:4)_[LVL2]; 141   -0.041000 0.45600    0.6040
## 81              PC(36:2)_[LVL2]; 3    0.044200 0.47200    0.6170
## 82             LPC(16:0)_[LVL1]; 5    0.037400 0.50300    0.6430
## 83   TG(14:0/16:0/18:1)_[LVL2]; 54   -0.029500 0.50300    0.6430
## 84              PC(38:4)_[LVL2]; 9   -0.037400 0.50900    0.6430
## 85             PC(40:6)_[LVL2]; 31   -0.027800 0.57700    0.7140
## 86           SM(d32:1)_[LVL2]; 105   -0.026400 0.57900    0.7140
## 87             PC(36:5)_[LVL2]; 23    0.027600 0.59000    0.7160
## 88  TG(18:2/18:2/18:2) or TG(18:3/   -0.025700 0.59500    0.7160
## 89            TG(56:7)_[LVL3]; 309   -0.027700 0.60100    0.7160
## 90   TG(18:2/22:5/16:0)_[LVL2]; 69   -0.023100 0.65500    0.7720
## 91            LPC(18:2)_[LVL2]; 33    0.025000 0.66900    0.7800
## 92           PC(O-36:5)_[LVL2]; 92   -0.023700 0.68200    0.7860
## 93            PC(38:2)_[LVL2]; 197   -0.020900 0.70000    0.7970
## 94             PC(40:5)_[LVL2]; 95   -0.018700 0.71100    0.8020
## 95              PC(36:4)_[LVL2]; 1    0.021900 0.72000    0.8030
## 96  SM(d16:1/18:1) or SM(d18:2/16:   -0.015200 0.73000    0.8060
## 97       SM(d18:1/24:0)_[LVL2]; 61   -0.009510 0.84300    0.9190
## 98   TG(18:1/18:1/18:1)_[LVL2]; 15   -0.007370 0.86000    0.9190
```

```
## 99            PC(34:3)_[LVL2]; 113   -0.009650 0.86200    0.9190
## 100 TG(18:2/18:1/16:0)_[LVL2]; 500    0.009380 0.86700    0.9190
## 101            SM(42:2)_[LVL2]; 14   -0.007850 0.87800    0.9220
## 102           LPC(18:0)_[LVL1]; 22    0.005390 0.92500    0.9600
## 103            PC(38:3)_[LVL2]; 29   -0.004620 0.93200    0.9600
## 104           SM(d34:1)_[LVL2]; 26   -0.002760 0.96100    0.9800
## 105             PC(34:1)_[LVL2]; 2    0.001120 0.98600    0.9890
## 106           SM(d40:1)_[LVL2]; 39   -0.000645 0.98900    0.9890
```

#### 7.2 Deep Breathing (E_I)

##### 7.2.1 Crude Model

```
## [1] "Fitting models:"
## [1] "~ E_I"
## [1] ""
```

###### 7.2.1.1 Heatmap

```
## [1] "heatmap_lipidome_from_limma was created by Tommi Suvitaival"
## [1] ""
## [1] "2019-05-21"
```

```
## Warning: Removed 106 rows containing missing values (geom_point).
```

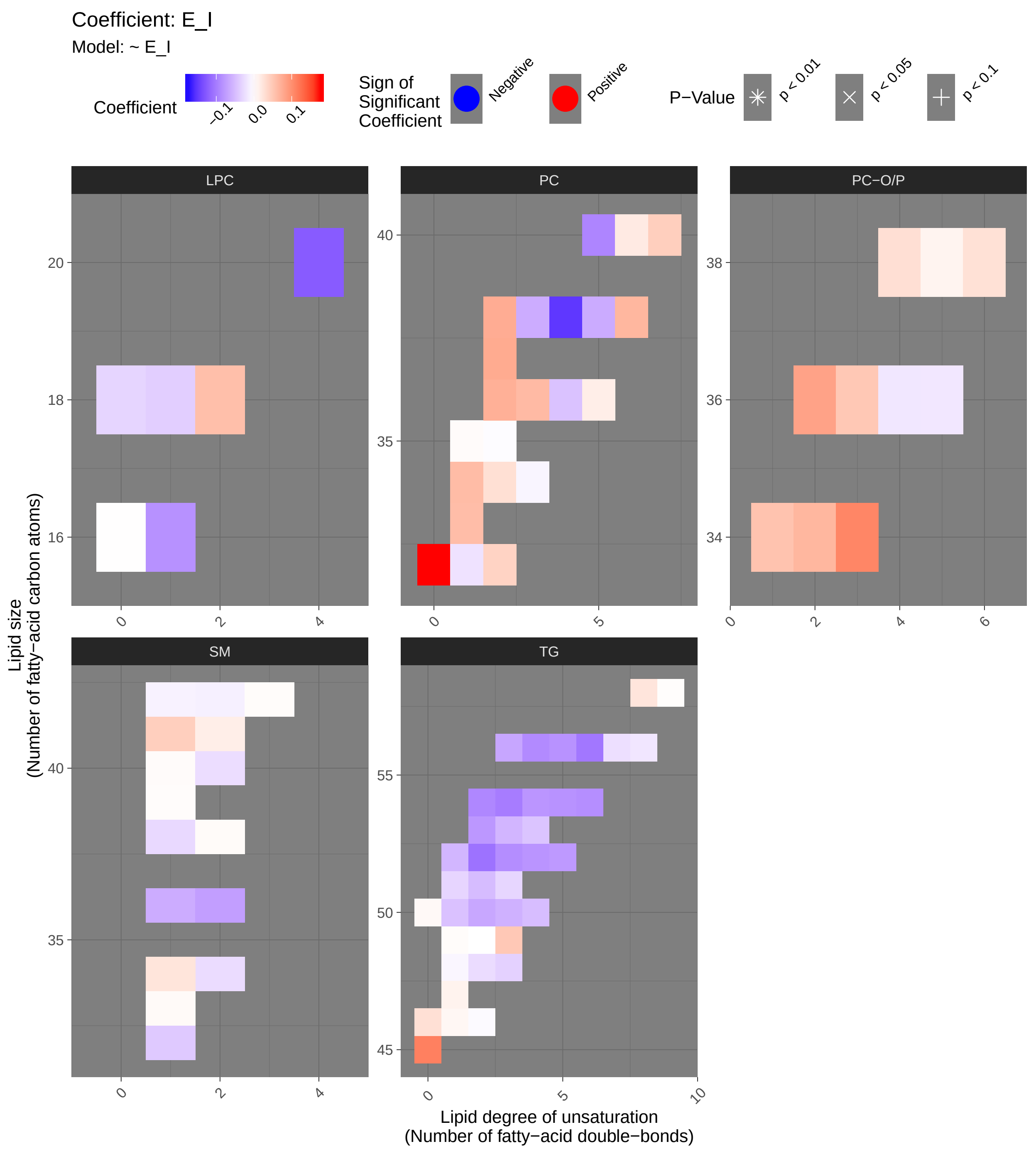

Coefficient: E_I
Model: ~ E_I
Coefficient
-0.1
0.0
0.1
Sign of Significant Coefficient
Negative
Positive
P−Value
p < 0.01
p < 0.05
p < 0.1
LPC
PC
PC−O/P
SM
TG
Lipid size (Number of fatty−acid carbon atoms)
Lipid degree of unsaturation (Number of fatty−acid double−bonds)

###### 7.2.1.2 Tables of Model Coefficients

```
## [1] ""
## [1] "Table: E_I"
## [1] "  (from model: "
## [1] "   ~ E_I)"
## [1] ""
##                              Name Coefficient P.Value adj.P.Val
## 1              PC(32:0)_[LVL2]; 96    1.76e-01 0.00265     0.172
## 2   TG(18:0/18:1/20:4)_[LVL2]; 141   -1.72e-01 0.00325     0.172
## 3               PC(38:4)_[LVL2]; 9   -1.51e-01 0.00989     0.350
## 4            LPC(20:4)_[LVL2]; 120   -1.25e-01 0.03260     0.686
## 5              TG(45:0)_[LVL2]; 65    1.13e-01 0.05380     0.686
## 6    TG(18:1/18:1/18:1)_[LVL2]; 15   -1.13e-01 0.05400     0.686
## 7           PC(O-34:3)_[LVL2]; 140    1.08e-01 0.06520     0.686
## 8              TG(52:2)_[LVL3]; 97   -1.08e-01 0.06570     0.686
## 9             TG(56:6)_[LVL3]; 275   -1.04e-01 0.07590     0.686
## 10            TG(54:3)_[LVL3]; 124   -9.94e-02 0.08940     0.686
## 11  TG(16:0/22:5/18:1) or TG(20:4/   -9.89e-02 0.09100     0.686
## 12             PC(40:5)_[LVL2]; 95   -9.29e-02 0.11200     0.686
## 13             TG(54:2)_[LVL3]; 52   -9.15e-02 0.11800     0.686
## 14            TG(56:4)_[LVL3]; 278   -8.91e-02 0.12800     0.686
## 15            TG(52:3)_[LVL3]; 101   -8.67e-02 0.13800     0.686
## 16            TG(54:6)_[LVL3]; 316   -8.58e-02 0.14300     0.686
## 17   TG(16:0/18:2/18:2)_[LVL2]; 27   -8.55e-02 0.14400     0.686
## 18          PC(O-36:2)_[LVL2]; 312    8.40e-02 0.15100     0.686
## 19           LPC(16:1)_[LVL2]; 258   -8.37e-02 0.15200     0.686
## 20            TG(54:5)_[LVL3]; 240   -8.30e-02 0.15600     0.686
## 21            TG(56:5)_[LVL2]; 230   -8.29e-02 0.15600     0.686
## 22   TG(18:2/18:1/18:1)_[LVL2]; 20   -8.23e-02 0.15900     0.686
## 23            TG(52:4)_[LVL3]; 157   -8.13e-02 0.16500     0.686
## 24            TG(54:4)_[LVL3]; 129   -8.00e-02 0.17100     0.686
## 25            TG(53:2)_[LVL2]; 234   -7.90e-02 0.17700     0.686
## 26            TG(52:5)_[LVL3]; 286   -7.71e-02 0.18800     0.686
## 27            PC(37:2)_[LVL2]; 350    7.68e-02 0.18900     0.686
## 28            PC(38:2)_[LVL2]; 197    7.53e-02 0.19800     0.686
## 29   TG(14:0/18:1/18:1)_[LVL2]; 25   -7.48e-02 0.20100     0.686
## 30   TG(18:1/18:2/18:2)_[LVL2]; 57   -7.47e-02 0.20100     0.686
## 31           SM(d36:2)_[LVL2]; 160   -7.31e-02 0.21100     0.686
## 32              PC(36:2)_[LVL2]; 3    7.22e-02 0.21700     0.686
## 33  TG(16:0/18:2/18:3)_[LVL2]; 106   -7.20e-02 0.21900     0.686
## 34  TG(18:2/18:1/16:0)_[LVL2]; 500   -7.07e-02 0.22700     0.686
## 35  TG(18:2/18:2/18:2) or TG(18:3/   -6.95e-02 0.23500     0.686
## 36            TG(56:3)_[LVL2]; 290   -6.72e-02 0.25100     0.686
## 37    TG(18:1/18:1/16:0)_[LVL2]; 7   -6.68e-02 0.25400     0.686
## 38            TG(50:2)_[LVL3]; 167   -6.63e-02 0.25700     0.686
## 39              PC(38:6)_[LVL2]; 8    6.62e-02 0.25800     0.686
## 40          PC(O-34:2)_[LVL2]; 171    6.60e-02 0.25900     0.686
## 41             PC(38:5)_[LVL2]; 24   -6.32e-02 0.28000     0.690
## 42             PC(36:3)_[LVL2]; 10    6.27e-02 0.28400     0.690
## 43            SM(d36:1)_[LVL2]; 55   -6.25e-02 0.28600     0.690
## 44             PC(38:3)_[LVL2]; 29   -6.23e-02 0.28700     0.690
## 45              PC(34:1)_[LVL2]; 2    6.12e-02 0.29500     0.696
## 46            PC(33:1)_[LVL2]; 177    6.02e-02 0.30300     0.699
```

```
## 47             TG(50:3)_[LVL2]; 47   -5.87e-02 0.31500     0.699
## 48            LPC(18:2)_[LVL2]; 33    5.86e-02 0.31700     0.699
## 49            TG(53:3)_[LVL3]; 239   -5.58e-02 0.34100     0.723
## 50   TG(16:0/18:0/18:1)_[LVL2]; 51   -5.51e-02 0.34700     0.723
## 51  PC(16:0e/18:1(9Z))_[LVL1]; 134    5.49e-02 0.34800     0.723
## 52            TG(51:2)_[LVL2]; 123   -5.06e-02 0.38700     0.766
## 53          PC(O-36:3)_[LVL2]; 268    5.05e-02 0.38800     0.766
## 54            TG(49:3)_[LVL3]; 218    5.00e-02 0.39200     0.766
## 55  TG(14:0/18:2/18:2)_[LVL2]; 189   -4.95e-02 0.39700     0.766
## 56             TG(50:1)_[LVL3]; 19   -4.66e-02 0.42600     0.795
## 57              PC(36:4)_[LVL2]; 1   -4.61e-02 0.43100     0.795
## 58            TG(53:4)_[LVL3]; 314   -4.46e-02 0.44600     0.795
## 59           SM(d41:1)_[LVL2]; 102    4.45e-02 0.44700     0.795
## 60            PC(40:7)_[LVL2]; 165    4.42e-02 0.45000     0.795
## 61           SM(d32:1)_[LVL2]; 105   -4.05e-02 0.48800     0.836
## 62            PC(32:2)_[LVL2]; 204    4.05e-02 0.48900     0.836
## 63            LPC(18:1)_[LVL2]; 34   -3.68e-02 0.52900     0.890
## 64            TG(48:3)_[LVL3]; 384   -3.41e-02 0.55900     0.927
## 65            LPC(18:0)_[LVL1]; 22   -3.17e-02 0.58800     0.932
## 66            TG(51:1)_[LVL3]; 249   -3.11e-02 0.59500     0.932
## 67            TG(51:3)_[LVL3]; 198   -3.08e-02 0.59800     0.932
## 68          PC(O-38:4)_[LVL2]; 131    2.91e-02 0.61900     0.932
## 69              PC(34:2)_[LVL2]; 4    2.90e-02 0.62100     0.932
## 70            SM(d38:1)_[LVL2]; 67   -2.86e-02 0.62500     0.932
## 71            TG(46:0)_[LVL3]; 168    2.83e-02 0.62800     0.932
## 72          PC(O-38:6)_[LVL2]; 236    2.76e-02 0.63700     0.932
## 73  TG(18:1/12:0/18:1) or TG(18:2/   -2.64e-02 0.65100     0.932
## 74  SM(d16:1/18:1) or SM(d18:2/16:   -2.59e-02 0.65900     0.932
## 75            SM(d40:2)_[LVL2]; 80   -2.52e-02 0.66700     0.932
## 76            SM(d34:1)_[LVL2]; 26    2.42e-02 0.67900     0.932
## 77            TG(56:7)_[LVL3]; 309   -2.37e-02 0.68500     0.932
## 78  TG(18:1/18:1/22:6)_[LVL2]; 147    2.37e-02 0.68600     0.932
## 79             PC(32:1)_[LVL2]; 44   -2.14e-02 0.71400     0.959
## 80   TG(18:2/22:5/16:0)_[LVL2]; 69   -1.94e-02 0.74000     0.963
## 81             PC(40:6)_[LVL2]; 31    1.92e-02 0.74200     0.963
## 82  TG(16:0/18:2/22:6)_[LVL2]; 117   -1.84e-02 0.75300     0.963
## 83           PC(O-36:4)_[LVL2]; 71   -1.79e-02 0.76000     0.963
## 84           PC(O-36:5)_[LVL2]; 92   -1.76e-02 0.76300     0.963
## 85           SM(d41:2)_[LVL2]; 139    1.57e-02 0.78800     0.976
## 86             PC(36:5)_[LVL2]; 23    1.54e-02 0.79200     0.976
## 87             SM(42:2)_[LVL2]; 14   -1.13e-02 0.84700     0.988
## 88            TG(47:1)_[LVL3]; 227    1.12e-02 0.84800     0.988
## 89           PC(O-38:5)_[LVL2]; 76    9.93e-03 0.86500     0.988
## 90       SM(d18:1/24:0)_[LVL2]; 61   -9.74e-03 0.86800     0.988
## 91            TG(46:1)_[LVL3]; 128    7.63e-03 0.89600     0.988
## 92            PC(34:3)_[LVL2]; 113   -7.56e-03 0.89700     0.988
## 93   TG(14:0/16:0/18:1)_[LVL2]; 54   -6.95e-03 0.90500     0.988
## 94            TG(50:0)_[LVL2]; 159    5.39e-03 0.92700     0.988
## 95           SM(d33:1)_[LVL2]; 166    4.60e-03 0.93700     0.988
## 96            TG(46:2)_[LVL3]; 248   -3.93e-03 0.94600     0.988
## 97           SM(d38:2)_[LVL2]; 151    3.77e-03 0.94900     0.988
## 98            SM(d40:1)_[LVL2]; 39    3.68e-03 0.95000     0.988
## 99            PC(35:1)_[LVL2]; 178    3.63e-03 0.95000     0.988
## 100      SM(d18:2/24:1)_[LVL2]; 40    3.23e-03 0.95600     0.988
```

```
## 101           TG(49:1)_[LVL3]; 187    3.14e-03 0.95700     0.988
## 102          SM(d39:1)_[LVL2]; 179    2.98e-03 0.95900     0.988
## 103           PC(35:2)_[LVL2]; 143   -2.51e-03 0.96600     0.988
## 104           TG(58:9)_[LVL3]; 207    2.25e-03 0.96900     0.988
## 105            LPC(16:0)_[LVL1]; 5    9.47e-04 0.98700     0.996
## 106           TG(49:2)_[LVL3]; 231   -7.23e-06 1.00000     1.000
```

###### 7.2.2 Adjusted Model

```
## [1] "Fitting models:"
## [1] "~ E_I + Age + bmi + Blood_glucose + Duration_DM + Gender + Hba1c_baseline + log_Blood_TGA + Smol
## [1] ""
```

###### 7.2.2.1 Heatmap

```
## [1] "heatmap_lipidome_from_limma was created by Tommi Suvitaival"
## [1] ""
## [1] "2019-05-21"
```

```
## Warning: Removed 106 rows containing missing values (geom_point).
```

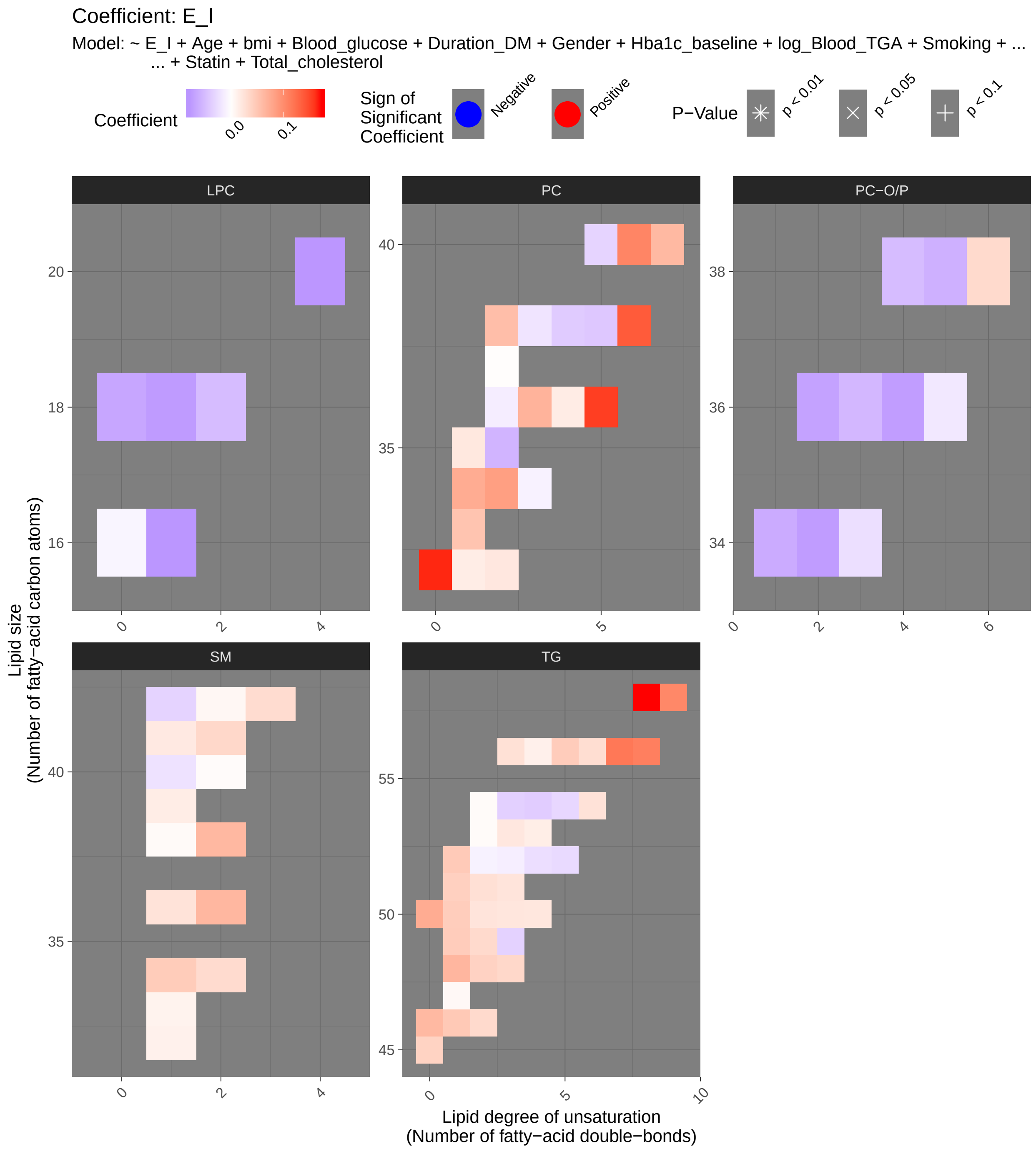
Coefficient: E_I
Model: ~ E_I + Age + bmi + Blood_glucose + Duration_DM + Gender + Hba1c_baseline + log_Blood_TGA + Smoking + ...
... + Statin + Total_cholesterol
Coefficient
0.0
0.1
Sign of Significant Coefficient
Negative
Positive
P−Value
p < 0.01
p < 0.05
p < 0.1
LPC
PC
PC−O/P
SM
TG
Lipid size
(Number of fatty−acid carbon atoms)
Lipid degree of unsaturation
(Number of fatty−acid double−bonds)

###### 7.2.2.2 Tables of Model Coefficients

```
## [1] ""
## [1] "Table: E_I"
## [1] "  (from model: "
## [1] "   ~ E_I + Age + bmi + Blood_glucose + Duration_DM + Gender"
## [1] "     + Hba1c_baseline + log_Blood_TGA + Smoking + Statin +"
## [1] "     Total_cholesterol)"
## [1] ""
##                               Name Coefficient P.Value adj.P.Val
## 1             PC(32:0)_[LVL2]; 96     0.16600 0.00267     0.141
## 2   TG(18:1/18:1/22:6)_[LVL2]; 147     0.17300 0.00276     0.141
## 3             PC(36:5)_[LVL2]; 23     0.15600 0.00398     0.141
## 4              PC(38:6)_[LVL2]; 8     0.13900 0.00886     0.235
## 5    TG(18:2/22:5/16:0)_[LVL2]; 69     0.13200 0.01520     0.323
## 6            TG(56:7)_[LVL3]; 309     0.11800 0.03310     0.585
## 7             PC(40:6)_[LVL2]; 31     0.10800 0.03950     0.597
## 8   TG(16:0/18:2/22:6)_[LVL2]; 117     0.11200 0.04840     0.641
## 9            TG(58:9)_[LVL3]; 207     0.10500 0.08450     0.857
## 10  TG(18:2/18:2/18:2) or TG(18:3/     0.07480 0.14300     0.857
## 11   TG(14:0/16:0/18:1)_[LVL2]; 54     0.06580 0.15300     0.857
## 12           TG(50:0)_[LVL2]; 159     0.07460 0.16500     0.857
## 13             PC(34:2)_[LVL2]; 4     0.08610 0.16800     0.857
## 14          LPC(20:4)_[LVL2]; 120    -0.07810 0.19400     0.857
## 15          LPC(16:1)_[LVL2]; 258    -0.07840 0.20300     0.857
## 16           LPC(18:1)_[LVL2]; 34    -0.07500 0.21400     0.857
## 17         PC(O-34:2)_[LVL2]; 171    -0.07420 0.21400     0.857
## 18   TG(18:1/18:1/18:1)_[LVL2]; 15    -0.05470 0.21500     0.857
## 19         PC(O-36:2)_[LVL2]; 312    -0.06950 0.21600     0.857
## 20          SM(d38:2)_[LVL2]; 151     0.06410 0.23400     0.857
## 21           PC(40:7)_[LVL2]; 165     0.06350 0.23500     0.857
## 22            PC(36:3)_[LVL2]; 10     0.06860 0.23800     0.857
## 23          PC(O-36:4)_[LVL2]; 71    -0.07300 0.23800     0.857
## 24          SM(d36:2)_[LVL2]; 160     0.06450 0.23800     0.857
## 25            TG(50:1)_[LVL3]; 19     0.04490 0.24000     0.857
## 26             PC(34:1)_[LVL2]; 2     0.07470 0.24900     0.857
## 27  PC(16:0e/18:1(9Z))_[LVL1]; 134    -0.06240 0.25100     0.857
## 28           TG(46:0)_[LVL3]; 168     0.06350 0.25200     0.857
## 29           LPC(18:0)_[LVL1]; 22    -0.06630 0.26500     0.857
## 30   TG(16:0/18:0/18:1)_[LVL2]; 51     0.04760 0.26800     0.857
## 31            TG(45:0)_[LVL2]; 65     0.04020 0.27900     0.857
## 32           PC(38:2)_[LVL2]; 197     0.05870 0.29500     0.857
## 33         PC(O-36:3)_[LVL2]; 268    -0.05320 0.30800     0.857
## 34           TG(56:5)_[LVL2]; 230     0.04610 0.31600     0.857
## 35          PC(O-38:5)_[LVL2]; 76    -0.05910 0.33700     0.857
## 36           PC(33:1)_[LVL2]; 177     0.05350 0.34000     0.857
## 37           TG(51:1)_[LVL3]; 249     0.04250 0.34100     0.857
## 38           PC(35:2)_[LVL2]; 143    -0.05580 0.35200     0.857
## 39           TG(46:1)_[LVL3]; 128     0.04910 0.36100     0.857
## 40  TG(18:1/12:0/18:1) or TG(18:2/     0.04070 0.36400     0.857
## 41           TG(49:1)_[LVL3]; 187     0.04580 0.37100     0.857
## 42           LPC(18:2)_[LVL2]; 33    -0.04980 0.40700     0.857
## 43         PC(O-38:4)_[LVL2]; 131    -0.05000 0.41300     0.857
## 44           TG(54:3)_[LVL3]; 124    -0.03490 0.42400     0.857
```

```
## 45             SM(d34:1)_[LVL2]; 26     0.04660 0.43200      0.857
## 46  TG(16:0/22:5/18:1) or TG(20:4/     0.03560 0.44200      0.857
## 47            TG(51:2)_[LVL2]; 123     0.02820 0.44800      0.857
## 48            TG(48:3)_[LVL3]; 384     0.03570 0.45200      0.857
## 49             PC(38:5)_[LVL2]; 24    -0.04140 0.45600      0.857
## 50            TG(54:4)_[LVL3]; 129    -0.03750 0.45700      0.857
## 51  TG(18:2/18:1/16:0)_[LVL2]; 500    -0.04130 0.46800      0.857
## 52            TG(49:3)_[LVL3]; 218    -0.03350 0.47700      0.857
## 53  SM(d16:1/18:1) or SM(d18:2/16:     0.03230 0.48300      0.857
## 54           SM(d41:2)_[LVL2]; 139     0.03550 0.50100      0.857
## 55            TG(49:2)_[LVL3]; 231     0.03350 0.50400      0.857
## 56            TG(50:2)_[LVL3]; 167     0.02420 0.51800      0.857
## 57              PC(38:4)_[LVL2]; 9    -0.03820 0.51900      0.857
## 58             TG(50:3)_[LVL2]; 47     0.02260 0.52000      0.857
## 59       SM(d18:1/24:0)_[LVL2]; 61    -0.03220 0.52200      0.857
## 60            TG(46:2)_[LVL3]; 248     0.03350 0.52400      0.857
## 61   TG(14:0/18:1/18:1)_[LVL2]; 25     0.02230 0.53200      0.857
## 62            TG(56:6)_[LVL3]; 275     0.03020 0.54400      0.857
## 63             PC(40:5)_[LVL2]; 95    -0.03200 0.54700      0.857
## 64       SM(d18:2/24:1)_[LVL2]; 40     0.03160 0.54900      0.857
## 65  TG(18:0/18:1/20:4)_[LVL2]; 141    -0.03380 0.55200      0.857
## 66          PC(O-38:6)_[LVL2]; 236     0.03320 0.55400      0.857
## 67            TG(51:3)_[LVL3]; 198     0.02410 0.56100      0.857
## 68            TG(52:4)_[LVL3]; 157    -0.02450 0.56900      0.857
## 69            TG(52:5)_[LVL3]; 286    -0.02750 0.57000      0.857
## 70            TG(56:3)_[LVL2]; 290     0.02770 0.57300      0.857
## 71   TG(18:2/18:1/18:1)_[LVL2]; 20    -0.02610 0.58000      0.857
## 72            TG(54:5)_[LVL3]; 240    -0.02950 0.58200      0.857
## 73            TG(53:3)_[LVL3]; 239     0.02170 0.59300      0.861
## 74  TG(14:0/18:2/18:2)_[LVL2]; 189     0.02190 0.60800      0.871
## 75            TG(54:6)_[LVL3]; 316     0.02590 0.62500      0.874
## 76   TG(16:0/18:2/18:2)_[LVL2]; 27    -0.01930 0.63300      0.874
## 77            SM(d36:1)_[LVL2]; 55     0.02510 0.63500      0.874
## 78          PC(O-34:3)_[LVL2]; 140    -0.02390 0.65400      0.889
## 79            SM(d40:1)_[LVL2]; 39    -0.02140 0.66600      0.894
## 80            PC(35:1)_[LVL2]; 178     0.02110 0.70400      0.910
## 81            PC(32:2)_[LVL2]; 204     0.02170 0.71200      0.910
## 82           SM(d41:1)_[LVL2]; 102     0.01960 0.71300      0.910
## 83             PC(38:3)_[LVL2]; 29    -0.01990 0.72700      0.910
## 84            TG(53:4)_[LVL3]; 314     0.01590 0.73500      0.910
## 85            TG(52:3)_[LVL3]; 101    -0.01230 0.75600      0.910
## 86           SM(d39:1)_[LVL2]; 179     0.01630 0.77200      0.910
## 87            TG(56:4)_[LVL3]; 278     0.01330 0.77300      0.910
## 88             PC(32:1)_[LVL2]; 44     0.01680 0.77700      0.910
## 89             TG(52:2)_[LVL3]; 97    -0.00986 0.77800      0.910
## 90           PC(O-36:5)_[LVL2]; 92    -0.01680 0.78100      0.910
## 91              PC(36:4)_[LVL2]; 1     0.01750 0.78100      0.910
## 92    TG(18:1/18:1/16:0)_[LVL2]; 7    -0.01330 0.80000      0.915
## 93           SM(d32:1)_[LVL2]; 105     0.01250 0.80300      0.915
## 94           SM(d33:1)_[LVL2]; 166     0.01120 0.83700      0.936
## 95              PC(36:2)_[LVL2]; 3    -0.01290 0.83900      0.936
## 96            PC(34:3)_[LVL2]; 113    -0.00942 0.87000      0.949
## 97  TG(16:0/18:2/18:3)_[LVL2]; 106    -0.00791 0.87300      0.949
## 98             SM(42:2)_[LVL2]; 14     0.00730 0.89100      0.949
```

```
## 99             LPC(16:0)_[LVL1]; 5    -0.00765 0.89600     0.949
## 100  TG(18:1/18:2/18:2)_[LVL2]; 57    -0.00609 0.90500     0.949
## 101           TG(53:2)_[LVL2]; 234     0.00395 0.91800     0.949
## 102           TG(47:1)_[LVL3]; 227     0.00618 0.92100     0.949
## 103            TG(54:2)_[LVL3]; 52     0.00372 0.92300     0.949
## 104           SM(d38:1)_[LVL2]; 67     0.00450 0.93100     0.949
## 105           SM(d40:2)_[LVL2]; 80     0.00330 0.94700     0.956
## 106           PC(37:2)_[LVL2]; 350     0.00173 0.97600     0.976
```

###### 7.2.3 Fully-Adjusted Model

```
## [1] "Fitting models:"
## [1] "~ E_I + Age + bmi + Blood_glucose + Duration_DM + Gender + Hba1c_baseline + log_Blood_TGA + Smo
## [1] ""
```

###### 7.2.3.1 Heatmap

```
## [1] "heatmap_lipidome_from_limma was created by Tommi Suvitaival"
## [1] ""
## [1] "2019-05-21"
```

```
## Warning: Removed 106 rows containing missing values (geom_point).
```

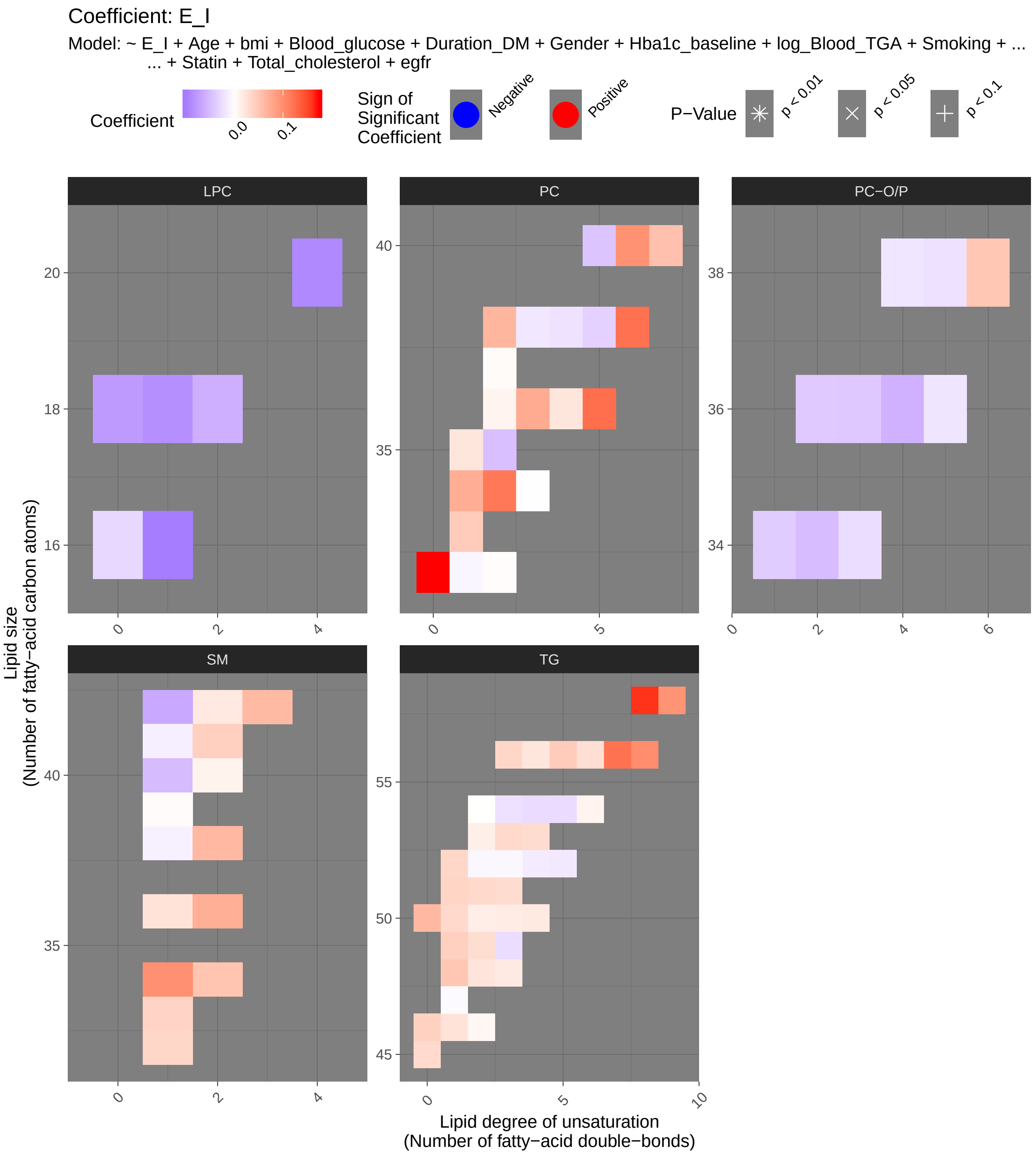
Coefficient: E_I
Model: ~ E_I + Age + bmi + Blood_glucose + Duration_DM + Gender + Hba1c_baseline + log_Blood_TGA + Smoking + ...
... + Statin + Total_cholesterol + egfr
Coefficient
0.0
0.1
Sign of Significant Coefficient
Negative
Positive
P−Value
p < 0.01
p < 0.05
p < 0.1
LPC
PC
PC−O/P
SM
TG
Lipid size
(Number of fatty−acid carbon atoms)
Lipid degree of unsaturation
(Number of fatty−acid double−bonds)

###### 7.2.3.2 Tables of Model Coefficients

```
## [1] ""
## [1] "Table: E_I"
## [1] "  (from model: "
## [1] "   ~ E_I + Age + bmi + Blood_glucose + Duration_DM + Gender"
## [1] "     + Hba1c_baseline + log_Blood_TGA + Smoking + Statin +"
## [1] "     Total_cholesterol + egfr)"
## [1] ""
##                               Name Coefficient P.Value adj.P.Val
## 1             PC(32:0)_[LVL2]; 96    0.173000 0.00229     0.243
## 2   TG(18:1/18:1/22:6)_[LVL2]; 147    0.160000 0.00687     0.364
## 3             PC(36:5)_[LVL2]; 23    0.124000 0.02360     0.564
## 4              PC(38:6)_[LVL2]; 8    0.123000 0.02380     0.564
## 5    TG(18:2/22:5/16:0)_[LVL2]; 69    0.123000 0.02760     0.564
## 6             TG(56:7)_[LVL3]; 309    0.122000 0.03190     0.564
## 7              PC(34:2)_[LVL2]; 4    0.117000 0.06690     0.926
## 8             PC(40:6)_[LVL2]; 31    0.096600 0.07150     0.926
## 9   TG(16:0/18:2/22:6)_[LVL2]; 117    0.100000 0.08440     0.926
## 10            SM(d34:1)_[LVL2]; 26    0.097800 0.10100     0.926
## 11           LPC(16:1)_[LVL2]; 258   -0.097300 0.12400     0.926
## 12            TG(58:9)_[LVL3]; 207    0.095200 0.12800     0.926
## 13           LPC(20:4)_[LVL2]; 120   -0.089200 0.14900     0.926
## 14            LPC(18:1)_[LVL2]; 34   -0.083800 0.17700     0.926
## 15           SM(d36:2)_[LVL2]; 160    0.071800 0.20100     0.926
## 16            PC(36:3)_[LVL2]; 10    0.075500 0.20700     0.926
## 17       SM(d18:1/24:0)_[LVL2]; 61   -0.064000 0.21100     0.926
## 18            LPC(18:0)_[LVL1]; 22   -0.075200 0.21900     0.926
## 19       SM(d18:2/24:1)_[LVL2]; 40    0.062100 0.24900     0.926
## 20           SM(d38:2)_[LVL2]; 151    0.063500 0.25200     0.926
## 21            TG(50:0)_[LVL2]; 159    0.062900 0.25400     0.926
## 22   TG(18:1/18:1/18:1)_[LVL2]; 15   -0.051600 0.25600     0.926
## 23            PC(38:2)_[LVL2]; 197    0.065300 0.25700     0.926
## 24  SM(d16:1/18:1) or SM(d18:2/16:    0.052600 0.26300     0.926
## 25              PC(34:1)_[LVL2]; 2    0.073400 0.27100     0.926
## 26   TG(14:0/16:0/18:1)_[LVL2]; 54    0.050700 0.28200     0.926
## 27  TG(18:2/18:2/18:2) or TG(18:3/    0.056000 0.28400     0.926
## 28            PC(40:7)_[LVL2]; 165    0.055600 0.31200     0.926
## 29            SM(d40:1)_[LVL2]; 39   -0.050500 0.31800     0.926
## 30            LPC(18:2)_[LVL2]; 33   -0.060200 0.33000     0.926
## 31            TG(56:5)_[LVL2]; 230    0.045600 0.33400     0.926
## 32           PC(O-36:4)_[LVL2]; 71   -0.057800 0.36200     0.926
## 33            TG(51:2)_[LVL2]; 123    0.034500 0.36700     0.926
## 34          PC(O-38:6)_[LVL2]; 236    0.050700 0.37800     0.926
## 35             TG(45:0)_[LVL2]; 65    0.033300 0.38200     0.926
## 36             TG(50:1)_[LVL3]; 19    0.034300 0.38200     0.926
## 37            TG(51:1)_[LVL3]; 249    0.038300 0.40300     0.926
## 38            TG(53:3)_[LVL3]; 239    0.034500 0.40600     0.926
## 39   TG(16:0/18:0/18:1)_[LVL2]; 51    0.036400 0.40900     0.926
## 40            TG(49:1)_[LVL3]; 187    0.043200 0.41100     0.926
## 41            PC(33:1)_[LVL2]; 177    0.046800 0.41600     0.926
## 42          PC(O-34:2)_[LVL2]; 171   -0.048900 0.42300     0.926
## 43             PC(40:5)_[LVL2]; 95   -0.043600 0.42500     0.926
## 44  TG(18:2/18:1/16:0)_[LVL2]; 500   -0.045600 0.43600     0.926
```

```
## 45          SM(d41:2)_[LVL2]; 139    0.042200 0.43700      0.926
## 46           PC(35:2)_[LVL2]; 143   -0.047100 0.44500      0.926
## 47         PC(O-36:3)_[LVL2]; 268   -0.040400 0.45000      0.926
## 48           TG(51:3)_[LVL3]; 198    0.031600 0.45800      0.926
## 49           TG(46:0)_[LVL3]; 168    0.041400 0.46500      0.926
## 50           TG(56:3)_[LVL2]; 290    0.036600 0.46800      0.926
## 51  TG(16:0/22:5/18:1) or TG(20:4/    0.033800 0.47700      0.926
## 52          SM(d32:1)_[LVL2]; 105    0.036400 0.47700      0.926
## 53         PC(O-36:2)_[LVL2]; 312   -0.040100 0.48400      0.926
## 54          SM(d33:1)_[LVL2]; 166    0.038500 0.48600      0.926
## 55  PC(16:0e/18:1(9Z))_[LVL1]; 134   -0.037900 0.49500      0.926
## 56           TG(53:4)_[LVL3]; 314    0.031500 0.51300      0.926
## 57            PC(38:5)_[LVL2]; 24   -0.033800 0.55400      0.926
## 58           TG(49:2)_[LVL3]; 231    0.030000 0.56100      0.926
## 59           TG(56:6)_[LVL3]; 275    0.029600 0.56200      0.926
## 60  TG(18:1/12:0/18:1) or TG(18:2/    0.024200 0.59800      0.926
## 61           TG(49:3)_[LVL3]; 218   -0.025200 0.60300      0.926
## 62           TG(54:4)_[LVL3]; 129   -0.026700 0.60600      0.926
## 63  TG(18:0/18:1/20:4)_[LVL2]; 141   -0.029900 0.60900      0.926
## 64           TG(54:3)_[LVL3]; 124   -0.022000 0.62300      0.926
## 65            LPC(16:0)_[LVL1]; 5   -0.028900 0.63000      0.926
## 66           TG(56:4)_[LVL3]; 278    0.022800 0.63000      0.926
## 67            TG(50:3)_[LVL2]; 47    0.017000 0.63800      0.926
## 68           SM(d36:1)_[LVL2]; 55    0.025200 0.64200      0.926
## 69           TG(54:5)_[LVL3]; 240   -0.025500 0.64400      0.926
## 70   TG(18:2/18:1/18:1)_[LVL2]; 20   -0.022400 0.64400      0.926
## 71         PC(O-34:3)_[LVL2]; 140   -0.025200 0.64600      0.926
## 72           TG(46:1)_[LVL3]; 128    0.025000 0.64800      0.926
## 73  TG(14:0/18:2/18:2)_[LVL2]; 189    0.019000 0.66500      0.926
## 74           TG(50:2)_[LVL3]; 167    0.015500 0.68700      0.926
## 75           PC(35:1)_[LVL2]; 178    0.023000 0.68800      0.926
## 76   TG(16:0/18:2/18:2)_[LVL2]; 27   -0.016100 0.69800      0.926
## 77           TG(48:3)_[LVL3]; 384    0.018800 0.69800      0.926
## 78           TG(53:2)_[LVL2]; 234    0.014900 0.70600      0.926
## 79            SM(42:2)_[LVL2]; 14    0.020300 0.71100      0.926
## 80   TG(14:0/18:1/18:1)_[LVL2]; 25    0.013300 0.71600      0.926
## 81          PC(O-38:5)_[LVL2]; 76   -0.022100 0.72300      0.926
## 82             PC(36:4)_[LVL2]; 1    0.022700 0.72600      0.926
## 83             PC(38:4)_[LVL2]; 9   -0.021200 0.72700      0.926
## 84           TG(52:4)_[LVL3]; 157   -0.014700 0.73900      0.926
## 85           TG(52:5)_[LVL3]; 286   -0.016300 0.74300      0.926
## 86            PC(38:3)_[LVL2]; 29   -0.017800 0.76100      0.927
## 87          PC(O-36:5)_[LVL2]; 92   -0.018800 0.76200      0.927
## 88         PC(O-38:4)_[LVL2]; 131   -0.018200 0.77000      0.927
## 89           SM(d40:2)_[LVL2]; 80    0.011100 0.82800      0.979
## 90          SM(d41:1)_[LVL2]; 102   -0.011400 0.83300      0.979
## 91           SM(d38:1)_[LVL2]; 67   -0.010500 0.84500      0.979
## 92           TG(54:6)_[LVL3]; 316    0.010300 0.85000      0.979
## 93            TG(52:2)_[LVL3]; 97   -0.005960 0.86800      0.980
## 94             PC(36:2)_[LVL2]; 3    0.010400 0.87300      0.980
## 95           TG(46:2)_[LVL3]; 248    0.007290 0.89200      0.980
## 96           TG(52:3)_[LVL3]; 101   -0.005490 0.89300      0.980
## 97            PC(32:1)_[LVL2]; 44   -0.006370 0.91600      0.980
## 98    TG(18:1/18:1/16:0)_[LVL2]; 7   -0.005380 0.92100      0.980
```

```
## 99   TG(18:1/18:2/18:2)_[LVL2]; 57   -0.004240 0.93600     0.980
## 100 TG(16:0/18:2/18:3)_[LVL2]; 106    0.003670 0.94200     0.980
## 101           PC(37:2)_[LVL2]; 350    0.003970 0.94600     0.980
## 102          SM(d39:1)_[LVL2]; 179    0.003520 0.95100     0.980
## 103           TG(47:1)_[LVL3]; 227   -0.003780 0.95300     0.980
## 104           PC(32:2)_[LVL2]; 204    0.002500 0.96700     0.985
## 105           PC(34:3)_[LVL2]; 113   -0.001340 0.98200     0.991
## 106            TG(54:2)_[LVL3]; 52    0.000371 0.99200     0.992
```

#### 7.3 Lying to Standing Test (lig_staa)

##### 7.3.1 Crude Model

```
## [1] "Fitting models:"
## [1] "~ lig_staa"
## [1] ""
```

###### 7.3.1.1 Heatmap

```
## [1] "heatmap_lipidome_from_limma was created by Tommi Suvitaival"
## [1] ""
## [1] "2019-05-21"

## Warning: Removed 105 rows containing missing values (geom_point).
```

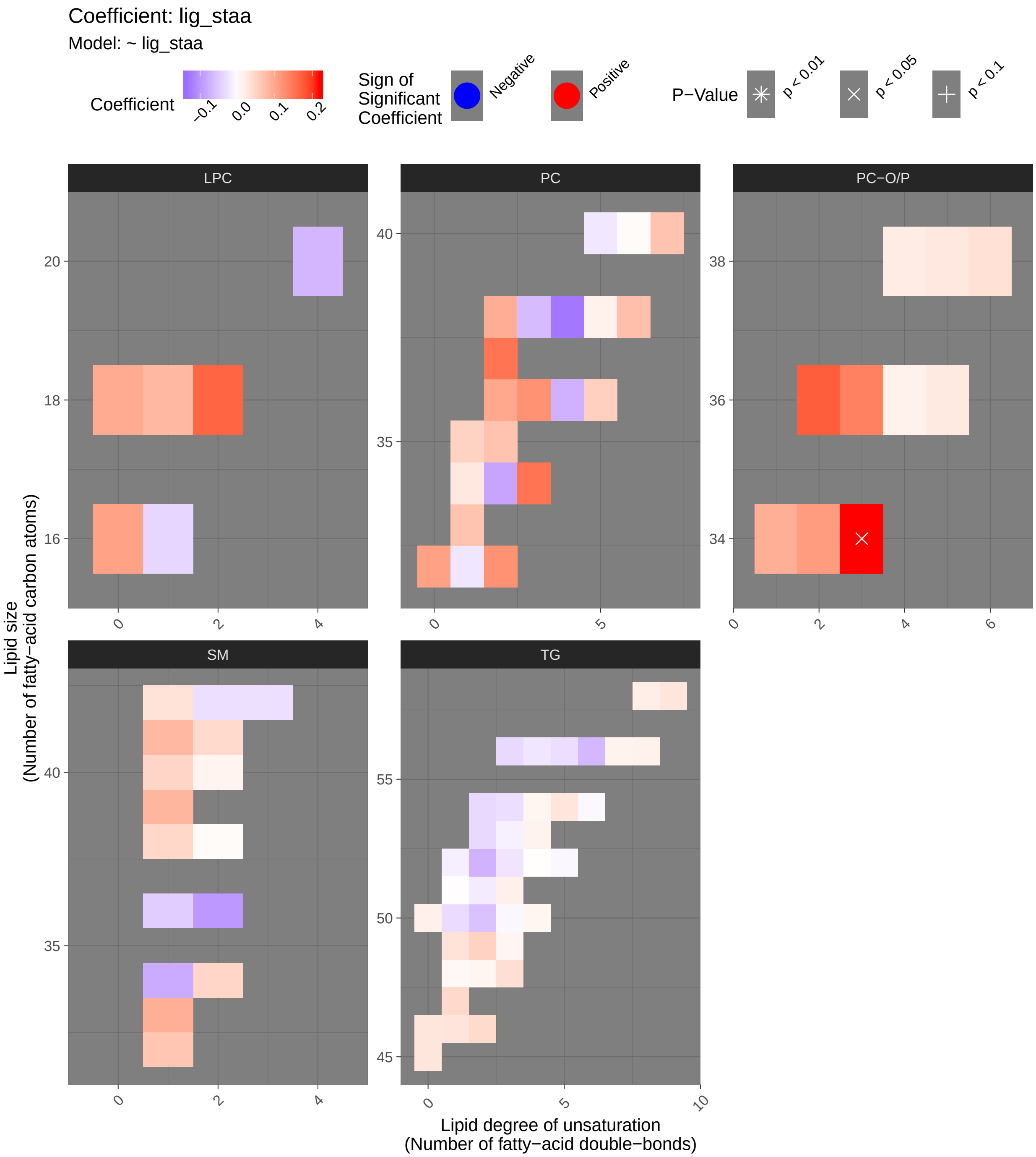

###### 7.3.1.2 Tables of Model Coefficients

```
## [1] ""
## [1] "Table: lig_staa"
## [1] "  (from model: "
## [1] "   ~ lig_staa)"
## [1] ""
##                               Name Coefficient P.Value adj.P.Val
## 1          PC(O-34:3)_[LVL2]; 140    0.219000 0.00021    0.0223
## 2          PC(O-36:2)_[LVL2]; 312    0.172000 0.00354    0.1620
## 3             LPC(18:2)_[LVL2]; 33    0.168000 0.00458    0.1620
## 4             PC(37:2)_[LVL2]; 350    0.153000 0.00963    0.2080
## 5             PC(34:3)_[LVL2]; 113    0.153000 0.00983    0.2080
## 6          PC(O-36:3)_[LVL2]; 268    0.139000 0.01830    0.3240
## 7   TG(18:0/18:1/20:4)_[LVL2]; 141   -0.136000 0.02150    0.3260
## 8               PC(38:4)_[LVL2]; 9   -0.128000 0.03050    0.4040
## 9              PC(36:3)_[LVL2]; 10    0.123000 0.03810    0.4100
## 10            PC(32:2)_[LVL2]; 204    0.122000 0.03870    0.4100
## 11          PC(O-34:2)_[LVL2]; 171    0.113000 0.05640    0.5430
## 12             PC(32:0)_[LVL2]; 96    0.106000 0.07320    0.6180
## 13             LPC(16:0)_[LVL1]; 5    0.105000 0.07570    0.6180
## 14              PC(36:2)_[LVL2]; 3    0.099000 0.09400    0.6590
## 15    TG(18:1/18:1/16:0)_[LVL2]; 7   -0.097600 0.09880    0.6590
## 16           SM(d36:2)_[LVL2]; 160   -0.097400 0.09940    0.6590
## 17            LPC(18:0)_[LVL1]; 22    0.095000 0.10800    0.6650
## 18            PC(38:2)_[LVL2]; 197    0.092000 0.12000    0.6650
## 19           SM(d33:1)_[LVL2]; 166    0.090600 0.12500    0.6650
## 20  PC(16:0e/18:1(9Z))_[LVL1]; 134    0.090500 0.12600    0.6650
## 21              PC(34:2)_[LVL2]; 4   -0.084300 0.15400    0.7720
## 22           SM(d39:1)_[LVL2]; 179    0.082700 0.16200    0.7720
## 23            LPC(18:1)_[LVL2]; 34    0.080800 0.17200    0.7720
## 24           SM(d41:1)_[LVL2]; 102    0.079300 0.18000    0.7720
## 25            SM(d34:1)_[LVL2]; 26   -0.078900 0.18200    0.7720
## 26              PC(36:4)_[LVL2]; 1   -0.073700 0.21200    0.8190
## 27              PC(38:6)_[LVL2]; 8    0.072700 0.21900    0.8190
## 28             TG(52:2)_[LVL3]; 97   -0.071900 0.22400    0.8190
## 29            PC(40:7)_[LVL2]; 165    0.068900 0.24400    0.8190
## 30            PC(35:2)_[LVL2]; 143    0.068300 0.24800    0.8190
## 31           LPC(20:4)_[LVL2]; 120   -0.068300 0.24800    0.8190
## 32            TG(56:6)_[LVL3]; 275   -0.067300 0.25500    0.8190
## 33            PC(33:1)_[LVL2]; 177    0.067300 0.25500    0.8190
## 34           SM(d32:1)_[LVL2]; 105    0.065300 0.26900    0.8380
## 35             PC(38:3)_[LVL2]; 29   -0.063400 0.28300    0.8580
## 36            TG(50:2)_[LVL3]; 167   -0.058200 0.32500    0.9570
## 37  TG(18:2/18:1/16:0)_[LVL2]; 500   -0.055400 0.34900    0.9580
## 38             PC(36:5)_[LVL2]; 23    0.054400 0.35700    0.9580
## 39   TG(18:1/18:1/18:1)_[LVL2]; 15   -0.053000 0.37000    0.9580
## 40            TG(49:2)_[LVL3]; 231    0.051400 0.38500    0.9580
## 41            PC(35:1)_[LVL2]; 178    0.050800 0.39000    0.9580
## 42   TG(14:0/18:1/18:1)_[LVL2]; 25   -0.049500 0.40200    0.9580
## 43            SM(d40:1)_[LVL2]; 39    0.048000 0.41700    0.9580
## 44            SM(d36:1)_[LVL2]; 55   -0.047100 0.42600    0.9580
## 45  SM(d16:1/18:1) or SM(d18:2/16:    0.046600 0.43000    0.9580
## 46            SM(d38:1)_[LVL2]; 67    0.045000 0.44600    0.9580
```

```
## 47            TG(47:1)_[LVL3]; 227    0.044100 0.45500    0.9580
## 48  TG(16:0/22:5/18:1) or TG(20:4/   -0.043200 0.46500    0.9580
## 49           SM(d41:2)_[LVL2]; 139    0.042200 0.47500    0.9580
## 50            TG(46:2)_[LVL3]; 248    0.041100 0.48600    0.9580
## 51           LPC(16:1)_[LVL2]; 258   -0.038500 0.51500    0.9580
## 52            TG(48:3)_[LVL3]; 384    0.037500 0.52600    0.9580
## 53            TG(53:2)_[LVL2]; 234   -0.035800 0.54400    0.9580
## 54             TG(54:2)_[LVL3]; 52   -0.035600 0.54700    0.9580
## 55            TG(56:3)_[LVL2]; 290   -0.035100 0.55200    0.9580
## 56          PC(O-38:6)_[LVL2]; 236    0.034300 0.56200    0.9580
## 57            TG(49:1)_[LVL3]; 187    0.033200 0.57400    0.9580
## 58       SM(d18:1/24:0)_[LVL2]; 61    0.032700 0.58100    0.9580
## 59             TG(50:1)_[LVL3]; 19   -0.032500 0.58300    0.9580
## 60            TG(46:1)_[LVL3]; 128    0.030700 0.60300    0.9580
## 61            TG(56:5)_[LVL2]; 230   -0.030700 0.60400    0.9580
## 62            TG(54:3)_[LVL3]; 124   -0.030300 0.60900    0.9580
## 63             SM(42:2)_[LVL2]; 14   -0.029500 0.61800    0.9580
## 64            TG(46:0)_[LVL3]; 168    0.028700 0.62700    0.9580
## 65            TG(54:5)_[LVL3]; 240    0.028500 0.62900    0.9580
## 66             TG(45:0)_[LVL2]; 65    0.028500 0.63000    0.9580
## 67       SM(d18:2/24:1)_[LVL2]; 40   -0.028500 0.63000    0.9580
## 68            TG(58:9)_[LVL3]; 207    0.028400 0.63100    0.9580
## 69           PC(O-38:5)_[LVL2]; 76    0.027000 0.64800    0.9580
## 70              PC(34:1)_[LVL2]; 2    0.026600 0.65200    0.9580
## 71   TG(18:1/18:2/18:2)_[LVL2]; 57    0.026000 0.66000    0.9580
## 72            TG(52:3)_[LVL3]; 101   -0.025100 0.67200    0.9580
## 73           PC(O-36:5)_[LVL2]; 92    0.024200 0.68200    0.9580
## 74            TG(56:4)_[LVL3]; 278   -0.023700 0.68900    0.9580
## 75             PC(32:1)_[LVL2]; 44   -0.023000 0.69700    0.9580
## 76             PC(40:5)_[LVL2]; 95   -0.022500 0.70300    0.9580
## 77          PC(O-38:4)_[LVL2]; 131    0.022100 0.70900    0.9580
## 78  TG(18:1/18:1/22:6)_[LVL2]; 147    0.019500 0.74100    0.9580
## 79            TG(51:2)_[LVL2]; 123   -0.018700 0.75200    0.9580
## 80            TG(50:0)_[LVL2]; 159    0.016800 0.77600    0.9580
## 81            TG(51:3)_[LVL3]; 198    0.016700 0.77700    0.9580
## 82             PC(38:5)_[LVL2]; 24    0.016400 0.78200    0.9580
## 83           PC(O-36:4)_[LVL2]; 71    0.016100 0.78600    0.9580
## 84  TG(16:0/18:2/22:6)_[LVL2]; 117    0.015400 0.79500    0.9580
## 85   TG(16:0/18:0/18:1)_[LVL2]; 51   -0.014900 0.80100    0.9580
## 86            TG(56:7)_[LVL3]; 309    0.014000 0.81300    0.9580
## 87            TG(53:3)_[LVL3]; 239   -0.014000 0.81300    0.9580
## 88            TG(53:4)_[LVL3]; 314    0.013500 0.81900    0.9580
## 89   TG(18:2/18:1/18:1)_[LVL2]; 20    0.013400 0.82100    0.9580
## 90            SM(d40:2)_[LVL2]; 80    0.013100 0.82400    0.9580
## 91            TG(54:4)_[LVL3]; 129    0.011900 0.84100    0.9580
## 92  TG(18:1/12:0/18:1) or TG(18:2/    0.011500 0.84600    0.9580
## 93  TG(14:0/18:2/18:2)_[LVL2]; 189    0.011400 0.84600    0.9580
## 94            TG(49:3)_[LVL3]; 218    0.011200 0.85000    0.9580
## 95  TG(16:0/18:2/18:3)_[LVL2]; 106    0.008940 0.88000    0.9620
## 96   TG(14:0/16:0/18:1)_[LVL2]; 54    0.007500 0.89900    0.9620
## 97            TG(52:5)_[LVL3]; 286   -0.007060 0.90500    0.9620
## 98             TG(50:3)_[LVL2]; 47   -0.007050 0.90500    0.9620
## 99            TG(54:6)_[LVL3]; 316   -0.006430 0.91300    0.9620
## 100  TG(16:0/18:2/18:2)_[LVL2]; 27    0.005290 0.92900    0.9620
```

```
## 101          SM(d38:2)_[LVL2]; 151    0.005060 0.93200    0.9620
## 102            PC(40:6)_[LVL2]; 31    0.004870 0.93400    0.9620
## 103  TG(18:2/22:5/16:0)_[LVL2]; 69   -0.004850 0.93500    0.9620
## 104           TG(51:1)_[LVL3]; 249    0.001730 0.97700    0.9900
## 105           TG(52:4)_[LVL3]; 157    0.001360 0.98200    0.9900
## 106 TG(18:2/18:2/18:2) or TG(18:3/   -0.000754 0.99000    0.9900
```

###### 7.3.1.3 Forest Plot of Model Coefficients

```
## Warning: Ignoring unknown aesthetics: x
```

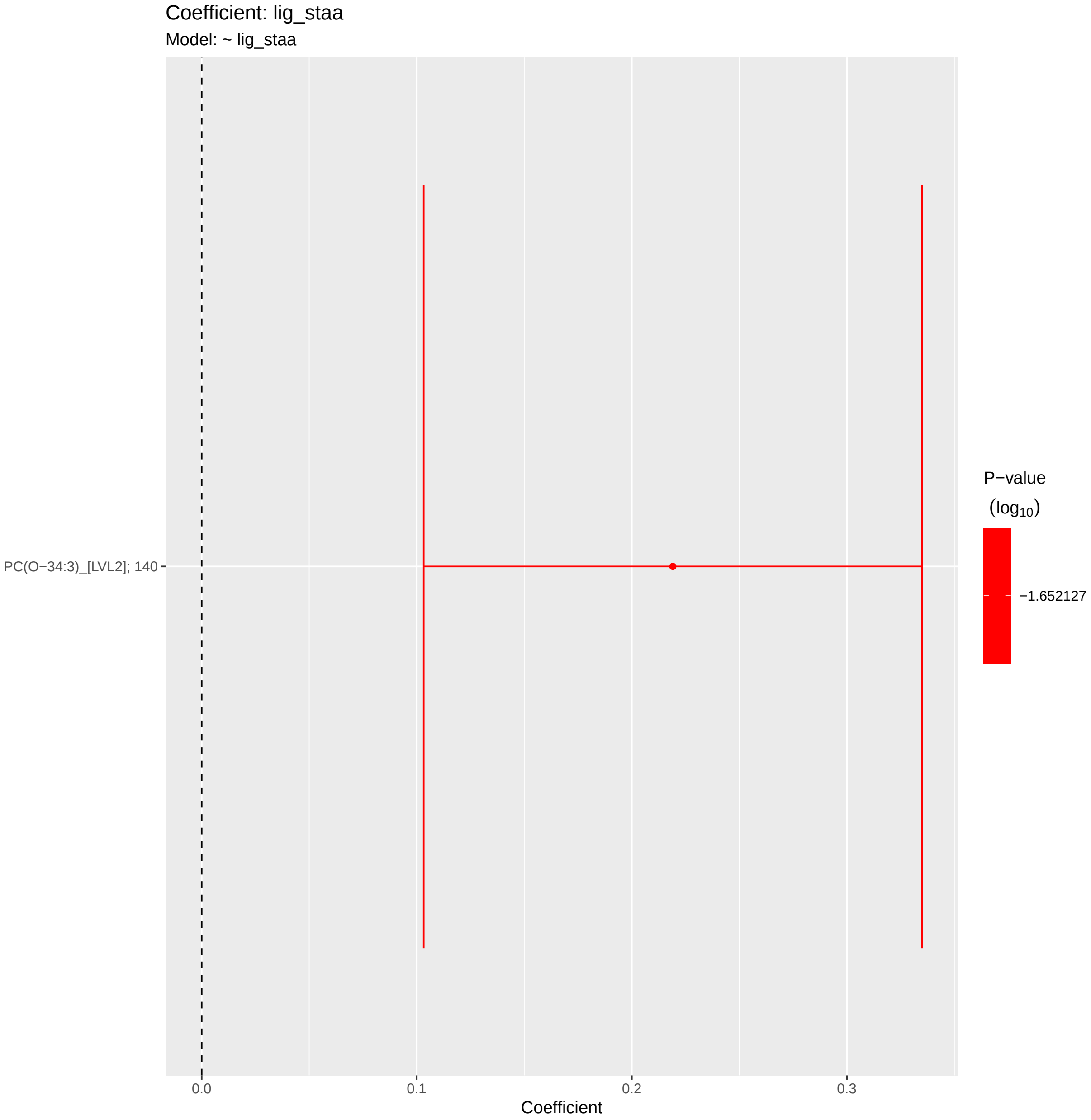

##### 7.3.2 Adjusted Model

```
## [1] "Fitting models:"
## [1] "~ lig_staa + Age + bmi + Blood_glucose + Duration_DM + Gender + Hba1c_baseline + log_Blood_TGA +
## [1] ""
```

###### 7.3.2.1 Heatmap

```
## [1] "heatmap_lipidome_from_limma was created by Tommi Suvitaival"
## [1] ""
## [1] "2019-05-21"

## Warning: Removed 106 rows containing missing values (geom_point).
```

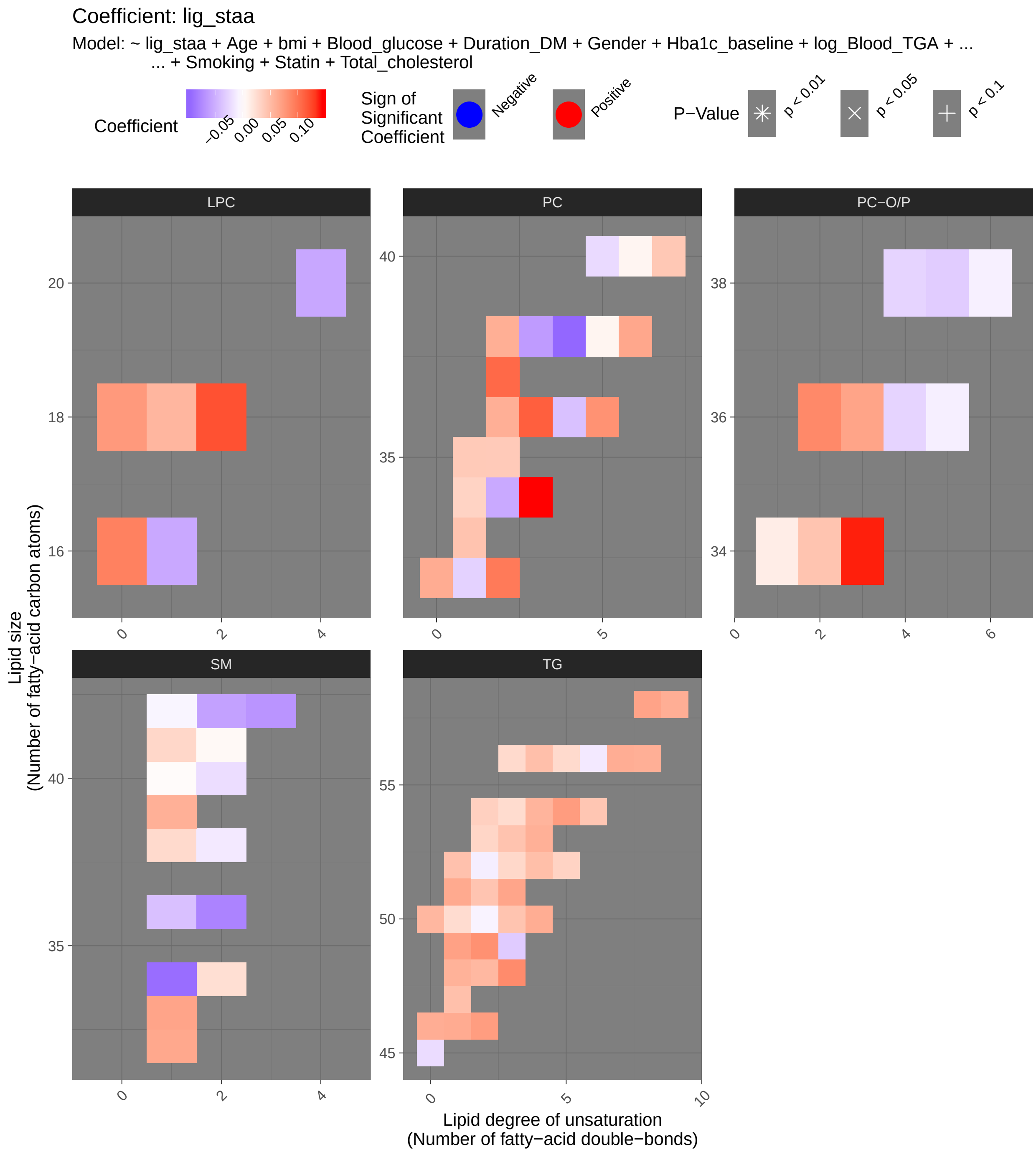

Coefficient: lig_staa
Model: ~ lig_staa + Age + bmi + Blood_glucose + Duration_DM + Gender + Hba1c_baseline + log_Blood_TGA + ...
... + Smoking + Statin + Total_cholesterol
Coefficient
−0.05
0.00
0.05
0.10
Sign of Significant Coefficient
Negative
Positive
P−Value
p < 0.01
p < 0.05
p < 0.1
LPC
PC
PC−O/P
SM
TG
Lipid size
(Number of fatty−acid carbon atoms)
Lipid degree of unsaturation
(Number of fatty−acid double−bonds)

###### 7.3.2.2 Tables of Model Coefficients

```
## [1] ""
## [1] "Table: lig_staa"
## [1] "  (from model: "
## [1] "   ~ lig_staa + Age + bmi + Blood_glucose + Duration_DM +"
## [1] "     Gender + Hba1c_baseline + log_Blood_TGA + Smoking + Statin +"
## [1] "     Total_cholesterol)"
## [1] ""
##                               Name Coefficient P.Value adj.P.Val
## 1          PC(O-34:3)_[LVL2]; 140     0.13900 0.00503     0.387
## 2            PC(34:3)_[LVL2]; 113     0.14400 0.00731     0.387
## 3            LPC(18:2)_[LVL2]; 33     0.12000 0.03050     0.550
## 4             PC(36:3)_[LVL2]; 10     0.11300 0.03700     0.550
## 5            PC(37:2)_[LVL2]; 350     0.10700 0.04620     0.550
## 6            TG(48:3)_[LVL3]; 384     0.08490 0.05600     0.550
## 7            PC(32:2)_[LVL2]; 204     0.09630 0.07880     0.550
## 8            TG(49:2)_[LVL3]; 231     0.08110 0.08060     0.550
## 9            TG(51:3)_[LVL3]; 198     0.06700 0.08220     0.550
## 10  TG(18:1/18:2/18:2)_[LVL2]; 57     0.08080 0.08500     0.550
## 11             PC(38:4)_[LVL2]; 9    -0.09490 0.08560     0.550
## 12  TG(18:2/18:1/18:1)_[LVL2]; 20     0.07450 0.08830     0.550
## 13             LPC(16:0)_[LVL1]; 5     0.09210 0.09190     0.550
## 14         PC(O-36:2)_[LVL2]; 312     0.08680 0.10200     0.550
## 15           SM(d34:1)_[LVL2]; 26    -0.09110 0.10400     0.550
## 16            PC(36:5)_[LVL2]; 23     0.08030 0.11200     0.550
## 17  TG(16:0/18:2/18:2)_[LVL2]; 27     0.05940 0.11200     0.550
## 18           TG(51:1)_[LVL3]; 249     0.06360 0.12700     0.550
## 19 TG(14:0/18:2/18:2)_[LVL2]; 189     0.06150 0.12800     0.550
## 20          SM(d36:2)_[LVL2]; 160    -0.07710 0.13200     0.550
## 21           TG(46:2)_[LVL3]; 248     0.07270 0.13700     0.550
## 22           TG(54:5)_[LVL3]; 240     0.07310 0.14000     0.550
## 23           TG(49:1)_[LVL3]; 187     0.06980 0.14200     0.550
## 24          SM(d32:1)_[LVL2]; 105     0.06470 0.16700     0.550
## 25         PC(O-36:3)_[LVL2]; 268     0.06800 0.16800     0.550
## 26            TG(50:3)_[LVL2]; 47     0.04460 0.16900     0.550
## 27           TG(53:4)_[LVL3]; 314     0.05880 0.17300     0.550
## 28           LPC(18:0)_[LVL1]; 22     0.07520 0.17800     0.550
## 29  TG(14:0/16:0/18:1)_[LVL2]; 54     0.05760 0.17900     0.550
## 30      SM(d18:2/24:1)_[LVL2]; 40    -0.06650 0.17900     0.550
## 31          SM(d33:1)_[LVL2]; 166     0.06760 0.18300     0.550
## 32             PC(38:6)_[LVL2]; 8     0.06530 0.18700     0.550
## 33 TG(18:1/12:0/18:1) or TG(18:2/     0.05330 0.20200     0.550
## 34           TG(51:2)_[LVL2]; 123     0.04410 0.20500     0.550
## 35           TG(46:1)_[LVL3]; 128     0.06270 0.20900     0.550
## 36 TG(18:1/18:1/22:6)_[LVL2]; 147     0.06820 0.21100     0.550
## 37 TG(18:0/18:1/20:4)_[LVL2]; 141    -0.06610 0.22000     0.550
## 38           TG(54:4)_[LVL3]; 129     0.05640 0.22600     0.550
## 39           TG(52:4)_[LVL3]; 157     0.04820 0.23000     0.550
## 40           TG(56:7)_[LVL3]; 309     0.06100 0.23100     0.550
## 41           TG(53:3)_[LVL3]; 239     0.04500 0.23300     0.550
## 42            PC(32:0)_[LVL2]; 96     0.06190 0.23400     0.550
## 43           TG(46:0)_[LVL3]; 168     0.06100 0.23600     0.550
## 44            SM(42:2)_[LVL2]; 14    -0.05840 0.24000     0.550
```

```
## 45            PC(38:2)_[LVL2]; 197     0.06000 0.24700      0.550
## 46   TG(16:0/18:0/18:1)_[LVL2]; 51     0.04630 0.24800      0.550
## 47             PC(38:3)_[LVL2]; 29    -0.06170 0.24800      0.550
## 48            TG(56:4)_[LVL3]; 278     0.04820 0.25800      0.550
## 49  TG(16:0/18:2/18:3)_[LVL2]; 106     0.05110 0.26000      0.550
## 50  TG(16:0/18:2/22:6)_[LVL2]; 117     0.05950 0.26100      0.550
## 51           SM(d39:1)_[LVL2]; 179     0.05920 0.26500      0.550
## 52    TG(18:1/18:1/16:0)_[LVL2]; 7    -0.05300 0.28100      0.565
## 53            TG(50:0)_[LVL2]; 159     0.05400 0.28200      0.565
## 54            TG(58:9)_[LVL3]; 207     0.06040 0.28800      0.565
## 55  TG(18:2/18:2/18:2) or TG(18:3/     0.04920 0.30000      0.579
## 56              PC(36:2)_[LVL2]; 3     0.05990 0.31700      0.599
## 57             TG(54:2)_[LVL3]; 52     0.03530 0.32200      0.599
## 58            LPC(18:1)_[LVL2]; 34     0.05450 0.33400      0.603
## 59           LPC(20:4)_[LVL2]; 120    -0.05430 0.33600      0.603
## 60           LPC(16:1)_[LVL2]; 258    -0.05370 0.35600      0.628
## 61              PC(34:2)_[LVL2]; 4    -0.05310 0.36600      0.637
## 62            TG(54:6)_[LVL3]; 316     0.04280 0.38200      0.652
## 63            TG(53:2)_[LVL2]; 234     0.03110 0.39200      0.653
## 64            PC(33:1)_[LVL2]; 177     0.04490 0.39400      0.653
## 65            PC(40:7)_[LVL2]; 165     0.04160 0.40800      0.662
## 66            TG(47:1)_[LVL3]; 227     0.04760 0.41300      0.662
## 67            TG(52:3)_[LVL3]; 101     0.02960 0.42400      0.662
## 68          PC(O-34:2)_[LVL2]; 171     0.04430 0.42900      0.662
## 69            SM(d36:1)_[LVL2]; 55    -0.03910 0.43200      0.662
## 70   TG(18:2/22:5/16:0)_[LVL2]; 69     0.03880 0.44200      0.662
## 71            PC(35:1)_[LVL2]; 178     0.04000 0.44300      0.662
## 72            TG(49:3)_[LVL3]; 218    -0.03170 0.45900      0.663
## 73             TG(50:1)_[LVL3]; 19     0.02610 0.46000      0.663
## 74            TG(52:5)_[LVL3]; 286     0.03300 0.46300      0.663
## 75            PC(35:2)_[LVL2]; 143     0.03950 0.48500      0.686
## 76            TG(54:3)_[LVL3]; 124     0.02650 0.51100      0.701
## 77            TG(56:5)_[LVL2]; 230     0.02780 0.51200      0.701
## 78              PC(36:4)_[LVL2]; 1    -0.03840 0.51600      0.701
## 79             TG(45:0)_[LVL2]; 65    -0.02150 0.53300      0.714
## 80            TG(56:3)_[LVL2]; 290     0.02800 0.53900      0.714
## 81           SM(d41:1)_[LVL2]; 102     0.03000 0.54900      0.719
## 82  TG(18:2/18:1/16:0)_[LVL2]; 500    -0.03090 0.57100      0.724
## 83            SM(d38:1)_[LVL2]; 67     0.02780 0.57300      0.724
## 84  SM(d16:1/18:1) or SM(d18:2/16:     0.02420 0.57300      0.724
## 85              PC(34:1)_[LVL2]; 2     0.03320 0.58600      0.727
## 86           PC(O-38:5)_[LVL2]; 76    -0.03120 0.59000      0.727
## 87  TG(16:0/22:5/18:1) or TG(20:4/     0.02240 0.59900      0.730
## 88             PC(32:1)_[LVL2]; 44    -0.02780 0.61500      0.741
## 89          PC(O-38:4)_[LVL2]; 131    -0.02660 0.64300      0.760
## 90             PC(40:5)_[LVL2]; 95    -0.02260 0.64500      0.760
## 91           PC(O-36:4)_[LVL2]; 71    -0.02610 0.65400      0.762
## 92            SM(d40:2)_[LVL2]; 80    -0.02050 0.66200      0.763
## 93             TG(52:2)_[LVL3]; 97    -0.01030 0.75200      0.858
## 94            TG(56:6)_[LVL3]; 275    -0.01340 0.76900      0.867
## 95           SM(d38:2)_[LVL2]; 151    -0.01340 0.79100      0.879
## 96  PC(16:0e/18:1(9Z))_[LVL1]; 134     0.01330 0.79600      0.879
## 97   TG(18:1/18:1/18:1)_[LVL2]; 15     0.00933 0.81800      0.894
## 98   TG(14:0/18:1/18:1)_[LVL2]; 25     0.00717 0.82800      0.896
```

```
## 99            TG(50:2)_[LVL3]; 167    -0.00706 0.83900     0.899
## 100          PC(O-36:5)_[LVL2]; 92    -0.00987 0.86200     0.907
## 101         PC(O-38:6)_[LVL2]; 236    -0.00907 0.86400     0.907
## 102            PC(40:6)_[LVL2]; 31     0.00736 0.88000     0.910
## 103            PC(38:5)_[LVL2]; 24     0.00753 0.88500     0.910
## 104      SM(d18:1/24:0)_[LVL2]; 61    -0.00614 0.89700     0.915
## 105          SM(d41:2)_[LVL2]; 139     0.00488 0.92200     0.931
## 106           SM(d40:1)_[LVL2]; 39     0.00292 0.95100     0.951
```

##### 7.3.3 Fully-Adjusted Model

```
## [1] "Fitting models:"
## [1] "~ lig_staa + Age + bmi + Blood_glucose + Duration_DM + Gender + Hba1c_baseline + log_Blood_TGA +
## [1] ""
```

###### 7.3.3.1 Heatmap

```
## [1] "heatmap_lipidome_from_limma was created by Tommi Suvitaival"
## [1] ""
## [1] "2019-05-21"

## Warning: Removed 106 rows containing missing values (geom_point).
```

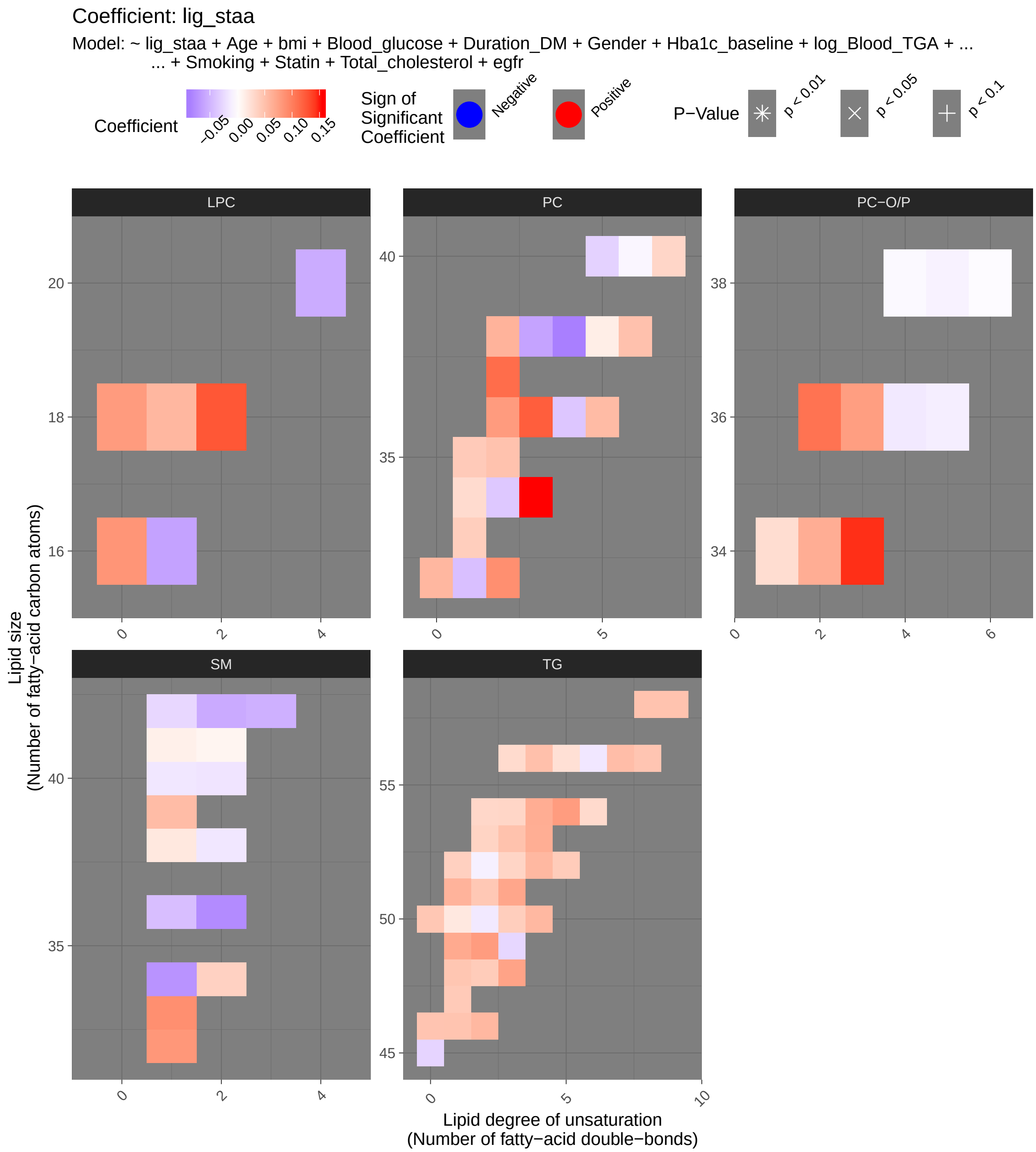

Coefficient: lig_staa
Model: ~ lig_staa + Age + bmi + Blood_glucose + Duration_DM + Gender + Hba1c_baseline + log_Blood_TGA + ...
... + Smoking + Statin + Total_cholesterol + egfr
Coefficient
−0.05 0.00 0.05 0.10 0.15
Sign of Significant Coefficient
Negative
Positive
P−Value
p < 0.01
p < 0.05
p < 0.1
LPC
PC
PC−O/P
SM
TG
Lipid size (Number of fatty−acid carbon atoms)
Lipid degree of unsaturation (Number of fatty−acid double−bonds)

###### 7.3.3.2 Tables of Model Coefficients

```
## [1] ""
## [1] "Table: lig_staa"
## [1] "  (from model: "
## [1] "   ~ lig_staa + Age + bmi + Blood_glucose + Duration_DM +"
## [1] "     Gender + Hba1c_baseline + log_Blood_TGA + Smoking + Statin +"
## [1] "     Total_cholesterol + egfr)"
## [1] ""
##                               Name Coefficient P.Value adj.P.Val
## 1          PC(O-34:3)_[LVL2]; 140    0.145000 0.00390     0.226
## 2            PC(34:3)_[LVL2]; 113    0.155000 0.00426     0.226
## 3            LPC(18:2)_[LVL2]; 33    0.127000 0.02460     0.638
## 4             PC(36:3)_[LVL2]; 10    0.122000 0.02580     0.638
## 5            PC(37:2)_[LVL2]; 350    0.113000 0.03740     0.638
## 6          PC(O-36:2)_[LVL2]; 312    0.109000 0.04090     0.638
## 7    TG(18:2/18:1/18:1)_[LVL2]; 20    0.081600 0.06500     0.638
## 8            TG(51:3)_[LVL3]; 198    0.071200 0.06820     0.638
## 9           SM(d32:1)_[LVL2]; 105    0.082600 0.07920     0.638
## 10   TG(18:1/18:2/18:2)_[LVL2]; 57    0.083200 0.08020     0.638
## 11          SM(d33:1)_[LVL2]; 166    0.088500 0.08170     0.638
## 12   TG(16:0/18:2/18:2)_[LVL2]; 27    0.063600 0.09280     0.638
## 13           TG(49:2)_[LVL3]; 231    0.078800 0.09380     0.638
## 14           TG(48:3)_[LVL3]; 384    0.073800 0.09950     0.638
## 15           PC(32:2)_[LVL2]; 204    0.088900 0.10900     0.638
## 16           TG(54:5)_[LVL3]; 240    0.078900 0.11600     0.638
## 17         PC(O-36:3)_[LVL2]; 268    0.077400 0.12100     0.638
## 18             PC(38:4)_[LVL2]; 9   -0.086000 0.12300     0.638
## 19            LPC(16:0)_[LVL1]; 5    0.084200 0.12800     0.638
## 20          SM(d36:2)_[LVL2]; 160   -0.077700 0.13400     0.638
## 21           TG(53:4)_[LVL3]; 314    0.065300 0.13500     0.638
## 22           TG(51:1)_[LVL3]; 249    0.060900 0.14900     0.638
## 23           TG(49:1)_[LVL3]; 187    0.068600 0.15400     0.638
## 24           TG(52:4)_[LVL3]; 157    0.057500 0.15700     0.638
## 25           LPC(18:0)_[LVL1]; 22    0.079400 0.16000     0.638
## 26           TG(54:4)_[LVL3]; 129    0.065900 0.16100     0.638
## 27  TG(14:0/18:2/18:2)_[LVL2]; 189    0.057000 0.16300     0.638
## 28             PC(36:2)_[LVL2]; 3    0.079900 0.18500     0.638
## 29           TG(53:3)_[LVL3]; 239    0.049700 0.19400     0.638
## 30           SM(d34:1)_[LVL2]; 26   -0.072400 0.19900     0.638
## 31  TG(16:0/18:2/18:3)_[LVL2]; 106    0.058500 0.20300     0.638
## 32           TG(51:2)_[LVL2]; 123    0.044700 0.20500     0.638
## 33  TG(18:0/18:1/20:4)_[LVL2]; 141   -0.066800 0.22100     0.638
## 34            TG(50:3)_[LVL2]; 47    0.039900 0.22500     0.638
## 35           TG(56:4)_[LVL3]; 278    0.051000 0.23800     0.638
## 36         PC(O-34:2)_[LVL2]; 171    0.065600 0.24200     0.638
## 37           PC(38:2)_[LVL2]; 197    0.061000 0.24500     0.638
## 38           TG(46:2)_[LVL3]; 248    0.057000 0.24600     0.638
## 39            PC(38:3)_[LVL2]; 29   -0.061400 0.25700     0.638
## 40            SM(42:2)_[LVL2]; 14   -0.056600 0.26100     0.638
## 41            PC(32:0)_[LVL2]; 96    0.058100 0.27000     0.638
## 42            PC(36:5)_[LVL2]; 23    0.055000 0.27300     0.638
## 43   TG(14:0/16:0/18:1)_[LVL2]; 54    0.046500 0.28100     0.638
## 44          LPC(16:1)_[LVL2]; 258   -0.062300 0.29000     0.638
```

```
## 45       SM(d18:2/24:1)_[LVL2]; 40   -0.052300 0.29300     0.638
## 46   TG(18:1/18:1/16:0)_[LVL2]; 7   -0.051800 0.29900     0.638
## 47           TG(56:7)_[LVL3]; 309    0.053100 0.30300     0.638
## 48           LPC(18:1)_[LVL2]; 34    0.057800 0.31200     0.638
## 49          SM(d39:1)_[LVL2]; 179    0.054100 0.31400     0.638
## 50             PC(38:6)_[LVL2]; 8    0.049800 0.31800     0.638
## 51  TG(18:1/12:0/18:1) or TG(18:2/    0.041700 0.32100     0.638
## 52          LPC(20:4)_[LVL2]; 120   -0.055400 0.33100     0.638
## 53           TG(53:2)_[LVL2]; 234    0.035300 0.33700     0.638
## 54           TG(46:1)_[LVL3]; 128    0.048100 0.33900     0.638
## 55   TG(16:0/18:0/18:1)_[LVL2]; 51    0.038200 0.34600     0.638
## 56           TG(52:5)_[LVL3]; 286    0.041600 0.36000     0.638
## 57           TG(46:0)_[LVL3]; 168    0.047600 0.36000     0.638
## 58           TG(52:3)_[LVL3]; 101    0.034200 0.36300     0.638
## 59            TG(54:2)_[LVL3]; 52    0.032500 0.36800     0.638
## 60           TG(50:0)_[LVL2]; 159    0.045500 0.37000     0.638
## 61  TG(18:1/18:1/22:6)_[LVL2]; 147    0.048400 0.37600     0.638
## 62  TG(16:0/18:2/22:6)_[LVL2]; 117    0.046600 0.38300     0.638
## 63           SM(d36:1)_[LVL2]; 55   -0.043200 0.39100     0.638
## 64           PC(35:2)_[LVL2]; 143    0.049000 0.39200     0.638
## 65  SM(d16:1/18:1) or SM(d18:2/16:    0.036900 0.39400     0.638
## 66           TG(58:9)_[LVL3]; 207    0.048600 0.39700     0.638
## 67           TG(54:3)_[LVL3]; 124    0.033600 0.41000     0.646
## 68            TG(45:0)_[LVL2]; 65   -0.028200 0.41900     0.646
## 69           PC(35:1)_[LVL2]; 178    0.042600 0.42100     0.646
## 70            PC(32:1)_[LVL2]; 44   -0.042200 0.45000     0.676
## 71           PC(33:1)_[LVL2]; 177    0.040000 0.45300     0.676
## 72  TG(18:2/18:2/18:2) or TG(18:3/    0.035100 0.46200     0.676
## 73           TG(47:1)_[LVL3]; 227    0.043000 0.46500     0.676
## 74           PC(40:7)_[LVL2]; 165    0.033100 0.51500     0.720
## 75           TG(56:3)_[LVL2]; 290    0.029500 0.52300     0.720
## 76             PC(36:4)_[LVL2]; 1   -0.038100 0.52500     0.720
## 77             PC(34:2)_[LVL2]; 4   -0.036400 0.53800     0.720
## 78           TG(49:3)_[LVL3]; 218   -0.026600 0.54000     0.720
## 79           TG(54:6)_[LVL3]; 316    0.030200 0.54000     0.720
## 80            PC(40:5)_[LVL2]; 95   -0.030000 0.54700     0.720
## 81  TG(18:2/18:1/16:0)_[LVL2]; 500   -0.032700 0.55500     0.720
## 82           TG(56:5)_[LVL2]; 230    0.025200 0.55700     0.720
## 83       SM(d18:1/24:0)_[LVL2]; 61   -0.026200 0.58300     0.739
## 84   TG(18:2/22:5/16:0)_[LVL2]; 69    0.027700 0.58600     0.739
## 85  PC(16:0e/18:1(9Z))_[LVL1]; 134    0.027700 0.59200     0.739
## 86            TG(50:1)_[LVL3]; 19    0.018300 0.60800     0.749
## 87             PC(34:1)_[LVL2]; 2    0.028900 0.64000     0.780
## 88  TG(16:0/22:5/18:1) or TG(20:4/    0.018600 0.66600     0.802
## 89           TG(50:2)_[LVL3]; 167   -0.014700 0.67500     0.804
## 90           SM(d38:1)_[LVL2]; 67    0.019000 0.70300     0.827
## 91           SM(d40:2)_[LVL2]; 80   -0.017700 0.71000     0.827
## 92           TG(56:6)_[LVL3]; 275   -0.016000 0.72900     0.840
## 93           SM(d40:1)_[LVL2]; 39   -0.015600 0.74000     0.844
## 94          SM(d38:2)_[LVL2]; 151   -0.016000 0.75500     0.850
## 95            TG(52:2)_[LVL3]; 97   -0.010100 0.76100     0.850
## 96            PC(38:5)_[LVL2]; 24    0.014200 0.78700     0.862
## 97   TG(18:1/18:1/18:1)_[LVL2]; 15    0.010300 0.80300     0.862
## 98          SM(d41:1)_[LVL2]; 102    0.012500 0.80400     0.862
```

```
## 99           PC(O-36:4)_[LVL2]; 71   -0.014500 0.80500     0.862
## 100          PC(O-36:5)_[LVL2]; 92   -0.011100 0.84700     0.898
## 101          SM(d41:2)_[LVL2]; 139    0.008450 0.86700     0.910
## 102          PC(O-38:5)_[LVL2]; 76   -0.008560 0.88300     0.917
## 103            PC(40:6)_[LVL2]; 31   -0.005720 0.90700     0.934
## 104         PC(O-38:4)_[LVL2]; 131   -0.004230 0.94100     0.960
## 105         PC(O-38:6)_[LVL2]; 236   -0.002770 0.95900     0.968
## 106  TG(14:0/18:1/18:1)_[LVL2]; 25   -0.000383 0.99100     0.991
```

#### 7.4 Valsalva Maneuver (Valsal)

##### 7.4.1 Crude Model

```
## [1] "Fitting models:"
## [1] "~ Valsal"
## [1] ""
```

###### 7.4.1.1 Heatmap

```
## [1] "heatmap_lipidome_from_limma was created by Tommi Suvitaival"
## [1] ""
## [1] "2019-05-21"
```

```
## Warning: Removed 104 rows containing missing values (geom_point).
```

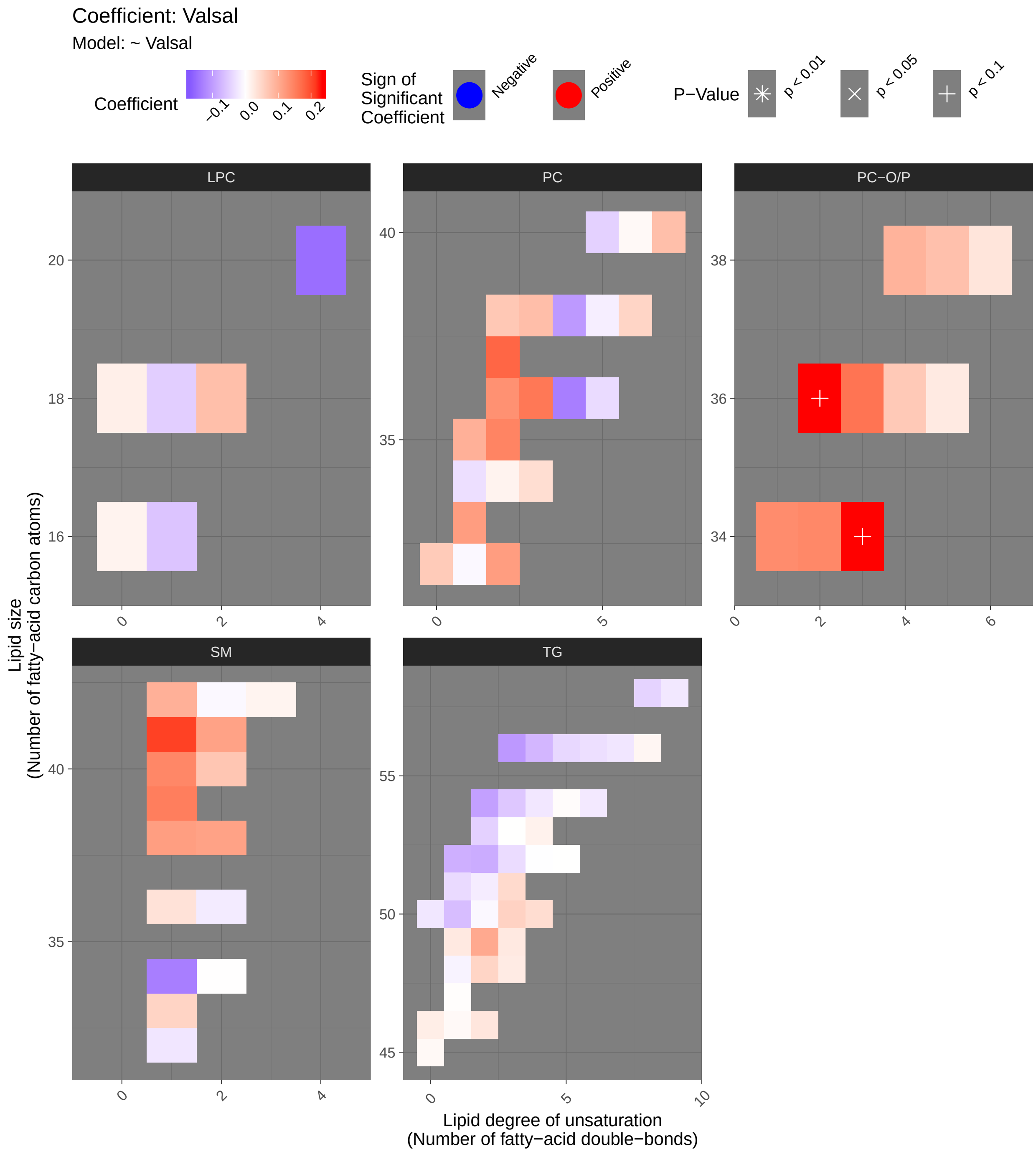

###### 7.4.1.2 Tables of Model Coefficients

```
## [1] ""
## [1] "Table: Valsal"
## [1] "  (from model: "
## [1] "   ~ Valsal)"
## [1] ""
##                               Name Coefficient P.Value adj.P.Val
## 1           PC(O-36:2)_[LVL2]; 312    0.234000 0.00107    0.0572
## 2           PC(O-34:3)_[LVL2]; 140    0.234000 0.00108    0.0572
## 3            SM(d41:1)_[LVL2]; 102    0.208000 0.00362    0.1280
## 4             PC(37:2)_[LVL2]; 350    0.177000 0.01330    0.3510
## 5   TG(18:0/18:1/20:4)_[LVL2]; 141   -0.168000 0.01870    0.3920
## 6           PC(O-36:3)_[LVL2]; 268    0.163000 0.02270    0.3920
## 7              PC(36:3)_[LVL2]; 10    0.159000 0.02590    0.3920
## 8            SM(d39:1)_[LVL2]; 179    0.153000 0.03300    0.4130
## 9            LPC(20:4)_[LVL2]; 120   -0.147000 0.04010    0.4130
## 10            PC(35:2)_[LVL2]; 143    0.147000 0.04040    0.4130
## 11            SM(d40:1)_[LVL2]; 39    0.143000 0.04540    0.4130
## 12          PC(O-34:2)_[LVL2]; 171    0.142000 0.04680    0.4130
## 13  PC(16:0e/18:1(9Z))_[LVL1]; 134    0.137000 0.05570    0.4490
## 14              PC(36:2)_[LVL2]; 3    0.132000 0.06580    0.4490
## 15            SM(d34:1)_[LVL2]; 26   -0.131000 0.06700    0.4490
## 16              PC(36:4)_[LVL2]; 1   -0.131000 0.06780    0.4490
## 17            PC(32:2)_[LVL2]; 204    0.119000 0.09750    0.5690
## 18            PC(33:1)_[LVL2]; 177    0.118000 0.09780    0.5690
## 19            SM(d38:1)_[LVL2]; 67    0.117000 0.10200    0.5690
## 20           SM(d41:2)_[LVL2]; 139    0.112000 0.11700    0.5930
## 21           SM(d38:2)_[LVL2]; 151    0.112000 0.11800    0.5930
## 22            TG(49:2)_[LVL3]; 231    0.104000 0.14600    0.6620
## 23            TG(56:3)_[LVL2]; 290   -0.104000 0.14600    0.6620
## 24              PC(38:4)_[LVL2]; 9   -0.103000 0.15000    0.6620
## 25       SM(d18:1/24:0)_[LVL2]; 61    0.095500 0.18200    0.7180
## 26            PC(35:1)_[LVL2]; 178    0.095400 0.18200    0.7180
## 27             TG(54:2)_[LVL3]; 52   -0.095300 0.18300    0.7180
## 28          PC(O-38:4)_[LVL2]; 131    0.092400 0.19600    0.7440
## 29             TG(52:2)_[LVL3]; 97   -0.083900 0.24100    0.8660
## 30   TG(16:0/18:0/18:1)_[LVL2]; 51   -0.080400 0.26100    0.8660
## 31             PC(38:3)_[LVL2]; 29    0.079100 0.26900    0.8660
## 32            LPC(18:2)_[LVL2]; 33    0.078000 0.27600    0.8660
## 33    TG(18:1/18:1/16:0)_[LVL2]; 7   -0.077600 0.27800    0.8660
## 34            PC(40:7)_[LVL2]; 165    0.077500 0.27900    0.8660
## 35           PC(O-38:5)_[LVL2]; 76    0.076400 0.28600    0.8660
## 36   TG(18:1/18:1/18:1)_[LVL2]; 15   -0.074600 0.29700    0.8740
## 37            TG(56:4)_[LVL3]; 278   -0.072800 0.30900    0.8850
## 38            SM(d40:2)_[LVL2]; 80    0.069800 0.33000    0.9060
## 39            PC(38:2)_[LVL2]; 197    0.067400 0.34700    0.9060
## 40  TG(16:0/22:5/18:1) or TG(20:4/   -0.066700 0.35100    0.9060
## 41           PC(O-36:4)_[LVL2]; 71    0.065900 0.35700    0.9060
## 42             TG(50:1)_[LVL3]; 19   -0.065600 0.35900    0.9060
## 43             PC(32:0)_[LVL2]; 96    0.063400 0.37600    0.9260
## 44           LPC(16:1)_[LVL2]; 258   -0.058800 0.41100    0.9760
## 45            TG(54:3)_[LVL3]; 124   -0.055600 0.43700    0.9760
## 46             TG(50:3)_[LVL2]; 47    0.054600 0.44500    0.9760
```

```
## 47          SM(d33:1)_[LVL2]; 166    0.052600 0.46200    0.9760
## 48  TG(18:1/12:0/18:1) or TG(18:2/    0.051300 0.47300    0.9760
## 49             PC(38:6)_[LVL2]; 8    0.051300 0.47300    0.9760
## 50           LPC(18:1)_[LVL2]; 34   -0.048700 0.49600    0.9760
## 51   TG(18:2/18:1/18:1)_[LVL2]; 20   -0.047400 0.50800    0.9760
## 52            PC(40:5)_[LVL2]; 95   -0.046200 0.51900    0.9760
## 53           TG(53:2)_[LVL2]; 234   -0.045500 0.52500    0.9760
## 54           TG(51:3)_[LVL3]; 198    0.045100 0.52900    0.9760
## 55  TG(18:1/18:1/22:6)_[LVL2]; 147   -0.043900 0.53900    0.9760
## 56   TG(18:2/22:5/16:0)_[LVL2]; 69   -0.043500 0.54400    0.9760
## 57  TG(14:0/18:2/18:2)_[LVL2]; 189    0.041700 0.56000    0.9760
## 58           PC(34:3)_[LVL2]; 113    0.040300 0.57400    0.9760
## 59           TG(56:5)_[LVL2]; 230   -0.038600 0.58900    0.9760
## 60  TG(18:2/18:2/18:2) or TG(18:3/   -0.038000 0.59500    0.9760
## 61           TG(51:1)_[LVL3]; 249   -0.036600 0.60900    0.9760
## 62            PC(36:5)_[LVL2]; 23   -0.035900 0.61600    0.9760
## 63  TG(18:2/18:1/16:0)_[LVL2]; 500   -0.035900 0.61600    0.9760
## 64           SM(d36:1)_[LVL2]; 55    0.035500 0.62000    0.9760
## 65           TG(52:3)_[LVL3]; 101   -0.034900 0.62600    0.9760
## 66         PC(O-38:6)_[LVL2]; 236    0.032500 0.64900    0.9760
## 67           TG(56:6)_[LVL3]; 275   -0.031900 0.65600    0.9760
## 68             PC(34:1)_[LVL2]; 2   -0.031900 0.65600    0.9760
## 69           TG(46:2)_[LVL3]; 248    0.030900 0.66600    0.9760
## 70           TG(49:1)_[LVL3]; 187    0.027400 0.70100    0.9760
## 71           TG(49:3)_[LVL3]; 218    0.026800 0.70700    0.9760
## 72          PC(O-36:5)_[LVL2]; 92    0.025600 0.72000    0.9760
## 73           TG(56:7)_[LVL3]; 309   -0.025100 0.72600    0.9760
## 74          SM(d32:1)_[LVL2]; 105   -0.024800 0.72800    0.9760
## 75           TG(48:3)_[LVL3]; 384    0.024400 0.73300    0.9760
## 76           TG(50:0)_[LVL2]; 159   -0.024200 0.73500    0.9760
## 77           TG(54:4)_[LVL3]; 129   -0.023500 0.74200    0.9760
## 78           TG(58:9)_[LVL3]; 207   -0.022500 0.75300    0.9760
## 79           TG(54:6)_[LVL3]; 316   -0.021700 0.76200    0.9760
## 80           TG(46:0)_[LVL3]; 168    0.021300 0.76500    0.9760
## 81   TG(18:1/18:2/18:2)_[LVL2]; 57   -0.020300 0.77600    0.9760
## 82          SM(d36:2)_[LVL2]; 160   -0.020200 0.77800    0.9760
## 83           LPC(18:0)_[LVL1]; 22    0.020000 0.77900    0.9760
## 84           TG(51:2)_[LVL2]; 123   -0.018400 0.79700    0.9760
## 85            PC(38:5)_[LVL2]; 24   -0.016700 0.81600    0.9760
## 86           TG(53:4)_[LVL3]; 314    0.016400 0.81800    0.9760
## 87             PC(34:2)_[LVL2]; 4    0.014700 0.83700    0.9760
## 88            LPC(16:0)_[LVL1]; 5    0.014200 0.84300    0.9760
## 89      SM(d18:2/24:1)_[LVL2]; 40    0.013700 0.84800    0.9760
## 90  TG(16:0/18:2/18:3)_[LVL2]; 106    0.013600 0.84900    0.9760
## 91   TG(14:0/16:0/18:1)_[LVL2]; 54   -0.012200 0.86500    0.9760
## 92  TG(16:0/18:2/22:6)_[LVL2]; 117    0.010600 0.88200    0.9760
## 93   TG(16:0/18:2/18:2)_[LVL2]; 27   -0.009410 0.89500    0.9760
## 94            TG(45:0)_[LVL2]; 65    0.007790 0.91300    0.9760
## 95            PC(40:6)_[LVL2]; 31    0.007560 0.91600    0.9760
## 96   TG(14:0/18:1/18:1)_[LVL2]; 25   -0.007400 0.91800    0.9760
## 97            PC(32:1)_[LVL2]; 44   -0.007350 0.91800    0.9760
## 98           TG(46:1)_[LVL3]; 128    0.007170 0.92000    0.9760
## 99           TG(50:2)_[LVL3]; 167   -0.007140 0.92000    0.9760
## 100           SM(42:2)_[LVL2]; 14   -0.007120 0.92100    0.9760
```

```
## 101           TG(54:5)_[LVL3]; 240    0.003780 0.95800    0.9940
## 102           TG(47:1)_[LVL3]; 227    0.002310 0.97400    0.9940
## 103           TG(52:4)_[LVL3]; 157   -0.001270 0.98600    0.9940
## 104 SM(d16:1/18:1) or SM(d18:2/16:    0.000925 0.99000    0.9940
## 105           TG(53:3)_[LVL3]; 239    0.000612 0.99300    0.9940
## 106           TG(52:5)_[LVL3]; 286    0.000520 0.99400    0.9940
```

##### 7.4.2 Adjusted Model

```
## [1] "Fitting models:"
## [1] "~ Valsal + Age + bmi + Blood_glucose + Duration_DM + Gender + Hba1c_baseline + log_Blood_TGA + [illegible]
## [1] ""
```

###### 7.4.2.1 Heatmap

```
## [1] "heatmap_lipidome_from_limma was created by Tommi Suvitaival"
## [1] ""
## [1] "2019-05-21"
```

```
## Warning: Removed 106 rows containing missing values (geom_point).
```

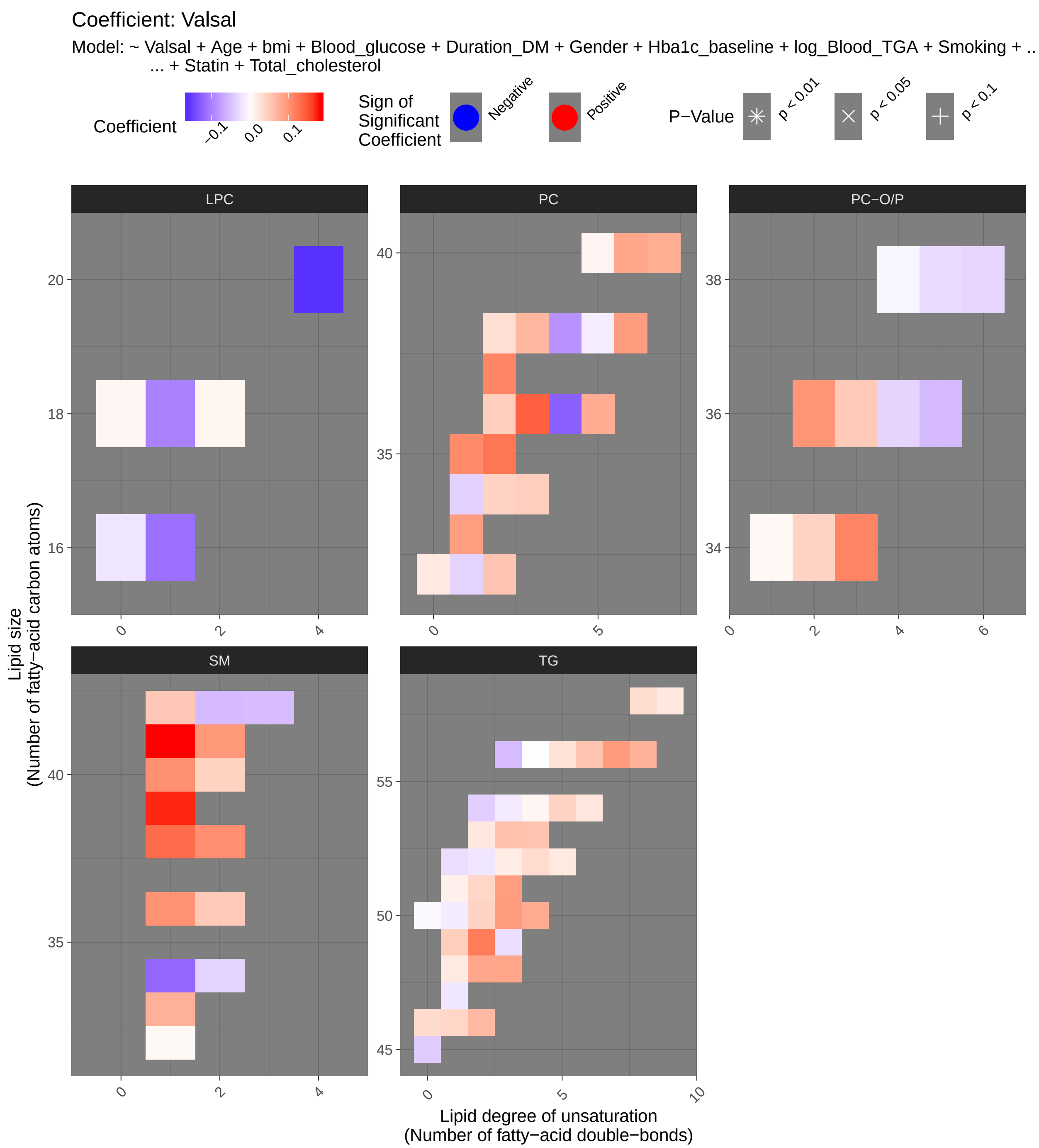

Coefficient: Valsal
Model: ~ Valsal + Age + bmi + Blood_glucose + Duration_DM + Gender + Hba1c_baseline + log_Blood_TGA + Smoking + ..
... + Statin + Total_cholesterol
Coefficient
−0.1
0.0
0.1
Sign of Significant Coefficient
Negative
Positive
P−Value
p < 0.01
p < 0.05
p < 0.1
LPC
PC
PC−O/P
SM
TG
Lipid size
(Number of fatty−acid carbon atoms)
Lipid degree of unsaturation
(Number of fatty−acid double−bonds)

###### 7.4.2.2 Tables of Model Coefficients

```
## [1] ""
## [1] "Table: Valsal"
## [1] "  (from model: "
## [1] "   ~ Valsal + Age + bmi + Blood_glucose + Duration_DM + Gender"
## [1] "     + Hba1c_baseline + log_Blood_TGA + Smoking + Statin +"
## [1] "     Total_cholesterol)"
## [1] ""
##                               Name Coefficient P.Value adj.P.Val
## 1           SM(d41:1)_[LVL2]; 102    0.181000 0.00649     0.688
## 2           SM(d39:1)_[LVL2]; 179    0.173000 0.01430     0.692
## 3           LPC(20:4)_[LVL2]; 120   -0.158000 0.02920     0.692
## 4             TG(50:3)_[LVL2]; 47    0.093200 0.03330     0.692
## 5            TG(49:2)_[LVL3]; 231    0.119000 0.04480     0.692
## 6            SM(d38:1)_[LVL2]; 67    0.132000 0.04790     0.692
## 7             PC(36:3)_[LVL2]; 10    0.140000 0.05230     0.692
## 8            TG(51:3)_[LVL3]; 198    0.091100 0.07810     0.692
## 9          PC(O-34:3)_[LVL2]; 140    0.113000 0.07940     0.692
## 10           PC(35:2)_[LVL2]; 143    0.125000 0.09660     0.692
## 11           SM(d34:1)_[LVL2]; 26   -0.120000 0.10100     0.692
## 12             PC(36:4)_[LVL2]; 1   -0.126000 0.10600     0.692
## 13           PC(37:2)_[LVL2]; 350    0.112000 0.11100     0.692
## 14           SM(d40:1)_[LVL2]; 39    0.102000 0.11400     0.692
## 15          SM(d38:2)_[LVL2]; 151    0.103000 0.11400     0.692
## 16           SM(d36:1)_[LVL2]; 55    0.100000 0.12000     0.692
## 17           PC(35:1)_[LVL2]; 178    0.107000 0.12000     0.692
## 18          LPC(16:1)_[LVL2]; 258   -0.113000 0.13000     0.692
## 19  TG(18:1/12:0/18:1) or TG(18:2/    0.082200 0.13200     0.692
## 20          SM(d41:2)_[LVL2]; 139    0.096800 0.14100     0.692
## 21             PC(38:6)_[LVL2]; 8    0.093300 0.14200     0.692
## 22  TG(14:0/18:2/18:2)_[LVL2]; 189    0.078700 0.15100     0.692
## 23           TG(48:3)_[LVL3]; 384    0.084100 0.15600     0.692
## 24         PC(O-36:2)_[LVL2]; 312    0.098700 0.15700     0.692
## 25           TG(56:7)_[LVL3]; 309    0.094300 0.17200     0.717
## 26            PC(40:6)_[LVL2]; 31    0.083100 0.18800     0.717
## 27           PC(33:1)_[LVL2]; 177    0.090800 0.18900     0.717
## 28           LPC(18:1)_[LVL2]; 34   -0.098800 0.18900     0.717
## 29            PC(36:5)_[LVL2]; 23    0.078200 0.23500     0.819
## 30  TG(18:0/18:1/20:4)_[LVL2]; 141   -0.086300 0.24000     0.819
## 31             PC(38:4)_[LVL2]; 9   -0.085300 0.24900     0.819
## 32           TG(53:3)_[LVL3]; 239    0.058900 0.25300     0.819
## 33           PC(40:7)_[LVL2]; 165    0.076400 0.25500     0.819
## 34          SM(d33:1)_[LVL2]; 166    0.073800 0.27500     0.857
## 35           TG(46:2)_[LVL3]; 248    0.066100 0.28900     0.875
## 36  TG(16:0/18:2/22:6)_[LVL2]; 117    0.072000 0.31400     0.902
## 37   TG(14:0/18:1/18:1)_[LVL2]; 25    0.045000 0.31700     0.902
## 38           TG(53:4)_[LVL3]; 314    0.055300 0.34000     0.902
## 39            PC(38:3)_[LVL2]; 29    0.067800 0.34300     0.902
## 40           TG(50:2)_[LVL3]; 167    0.041700 0.37800     0.902
## 41            TG(45:0)_[LVL2]; 65   -0.040100 0.38300     0.902
## 42           TG(56:6)_[LVL3]; 275    0.055000 0.38400     0.902
## 43   TG(18:1/18:1/18:1)_[LVL2]; 15   -0.046300 0.38700     0.902
## 44           TG(56:3)_[LVL2]; 290   -0.052600 0.39500     0.902
```

```
## 45             SM(42:2)_[LVL2]; 14   -0.053100 0.41800      0.902
## 46         PC(O-36:3)_[LVL2]; 268    0.050500 0.42000      0.902
## 47       SM(d18:2/24:1)_[LVL2]; 40   -0.051400 0.42100      0.902
## 48            TG(51:2)_[LVL2]; 123    0.038400 0.42500      0.902
## 49       SM(d18:1/24:0)_[LVL2]; 61    0.051600 0.42500      0.902
## 50            PC(32:2)_[LVL2]; 204    0.055500 0.43900      0.902
## 51           SM(d36:2)_[LVL2]; 160    0.049900 0.44000      0.902
## 52           PC(O-36:5)_[LVL2]; 92   -0.054200 0.44400      0.902
## 53            TG(49:1)_[LVL3]; 187    0.045800 0.45200      0.902
## 54  TG(16:0/18:2/18:3)_[LVL2]; 106    0.043700 0.46100      0.902
## 55             TG(54:2)_[LVL3]; 52   -0.037000 0.46800      0.902
## 56            SM(d40:2)_[LVL2]; 80    0.042800 0.49000      0.902
## 57            TG(54:5)_[LVL3]; 240    0.041900 0.50700      0.902
## 58            TG(52:4)_[LVL3]; 157    0.033600 0.52800      0.902
## 59            PC(34:3)_[LVL2]; 113    0.046000 0.53900      0.902
## 60  SM(d16:1/18:1) or SM(d18:2/16:   -0.034200 0.54800      0.902
## 61            TG(46:1)_[LVL3]; 128    0.038400 0.54900      0.902
## 62              PC(36:2)_[LVL2]; 3    0.046400 0.55000      0.902
## 63         PC(O-34:2)_[LVL2]; 171    0.041400 0.57300      0.902
## 64              PC(34:2)_[LVL2]; 4    0.042300 0.59300      0.902
## 65   TG(16:0/18:2/18:2)_[LVL2]; 27    0.026700 0.59500      0.902
## 66   TG(18:2/22:5/16:0)_[LVL2]; 69    0.035100 0.61200      0.902
## 67    TG(18:1/18:1/16:0)_[LVL2]; 7   -0.031900 0.61300      0.902
## 68            TG(46:0)_[LVL3]; 168    0.034500 0.61700      0.902
## 69         PC(O-38:6)_[LVL2]; 236   -0.032300 0.62100      0.902
## 70             PC(32:1)_[LVL2]; 44   -0.033700 0.63800      0.902
## 71   TG(16:0/18:0/18:1)_[LVL2]; 51   -0.025900 0.64400      0.902
## 72            TG(56:5)_[LVL2]; 230    0.026900 0.64600      0.902
## 73  TG(18:1/18:1/22:6)_[LVL2]; 147    0.033000 0.65100      0.902
## 74           PC(O-36:4)_[LVL2]; 71   -0.032900 0.65500      0.902
## 75            TG(49:3)_[LVL3]; 218   -0.025600 0.66100      0.902
## 76              PC(34:1)_[LVL2]; 2   -0.035200 0.66300      0.902
## 77             TG(52:2)_[LVL3]; 97   -0.019500 0.66400      0.902
## 78            TG(53:2)_[LVL2]; 234    0.021400 0.66700      0.902
## 79            PC(38:2)_[LVL2]; 197    0.030000 0.67600      0.902
## 80           PC(O-38:5)_[LVL2]; 76   -0.028800 0.69600      0.902
## 81            TG(54:6)_[LVL3]; 316    0.023300 0.71800      0.902
## 82   TG(14:0/16:0/18:1)_[LVL2]; 54    0.019900 0.73400      0.902
## 83            TG(52:3)_[LVL3]; 101    0.017300 0.73500      0.902
## 84            TG(52:5)_[LVL3]; 286    0.020100 0.73500      0.902
## 85             TG(50:1)_[LVL3]; 19   -0.015400 0.75400      0.902
## 86             PC(32:0)_[LVL2]; 96    0.020600 0.75900      0.902
## 87            TG(54:3)_[LVL3]; 124   -0.016500 0.76100      0.902
## 88            TG(58:9)_[LVL3]; 207    0.022100 0.76900      0.902
## 89             LPC(16:0)_[LVL1]; 5   -0.019600 0.78400      0.902
## 90   TG(18:2/18:1/18:1)_[LVL2]; 20   -0.014500 0.78900      0.902
## 91  TG(18:2/18:2/18:2) or TG(18:3/    0.017200 0.79100      0.902
## 92            TG(47:1)_[LVL3]; 227   -0.019400 0.79700      0.902
## 93            TG(51:1)_[LVL3]; 249    0.014100 0.79900      0.902
## 94   TG(18:1/18:2/18:2)_[LVL2]; 57    0.014900 0.80000      0.902
## 95  TG(18:2/18:1/16:0)_[LVL2]; 500   -0.017100 0.81300      0.907
## 96             PC(38:5)_[LVL2]; 24   -0.013800 0.84200      0.930
## 97            TG(54:4)_[LVL3]; 129    0.009320 0.87500      0.940
## 98             PC(40:5)_[LVL2]; 95    0.009990 0.87900      0.940
```

```
## 99  TG(16:0/22:5/18:1) or TG(20:4/    0.007910 0.89300     0.940
## 100           LPC(18:2)_[LVL2]; 33    0.009720 0.89600     0.940
## 101           LPC(18:0)_[LVL1]; 22    0.009300 0.90200     0.940
## 102 PC(16:0e/18:1(9Z))_[LVL1]; 134    0.006920 0.91500     0.940
## 103         PC(O-38:4)_[LVL2]; 131   -0.007590 0.91900     0.940
## 104          SM(d32:1)_[LVL2]; 105    0.006120 0.92200     0.940
## 105           TG(50:0)_[LVL2]; 159   -0.005290 0.93800     0.947
## 106           TG(56:4)_[LVL3]; 278    0.000851 0.98800     0.988
```

##### 7.4.3 Fully-Adjusted Model

```
## [1] "Fitting models:"
## [1] "~ Valsal + Age + bmi + Blood_glucose + Duration_DM + Gender + Hba1c_baseline + log_Blood_TGA + S
## [1] ""
```

###### 7.4.3.1 Heatmap

```
## [1] "heatmap_lipidome_from_limma was created by Tommi Suvitaival"
## [1] ""
## [1] "2019-05-21"
```

```
## Warning: Removed 106 rows containing missing values (geom_point).
```

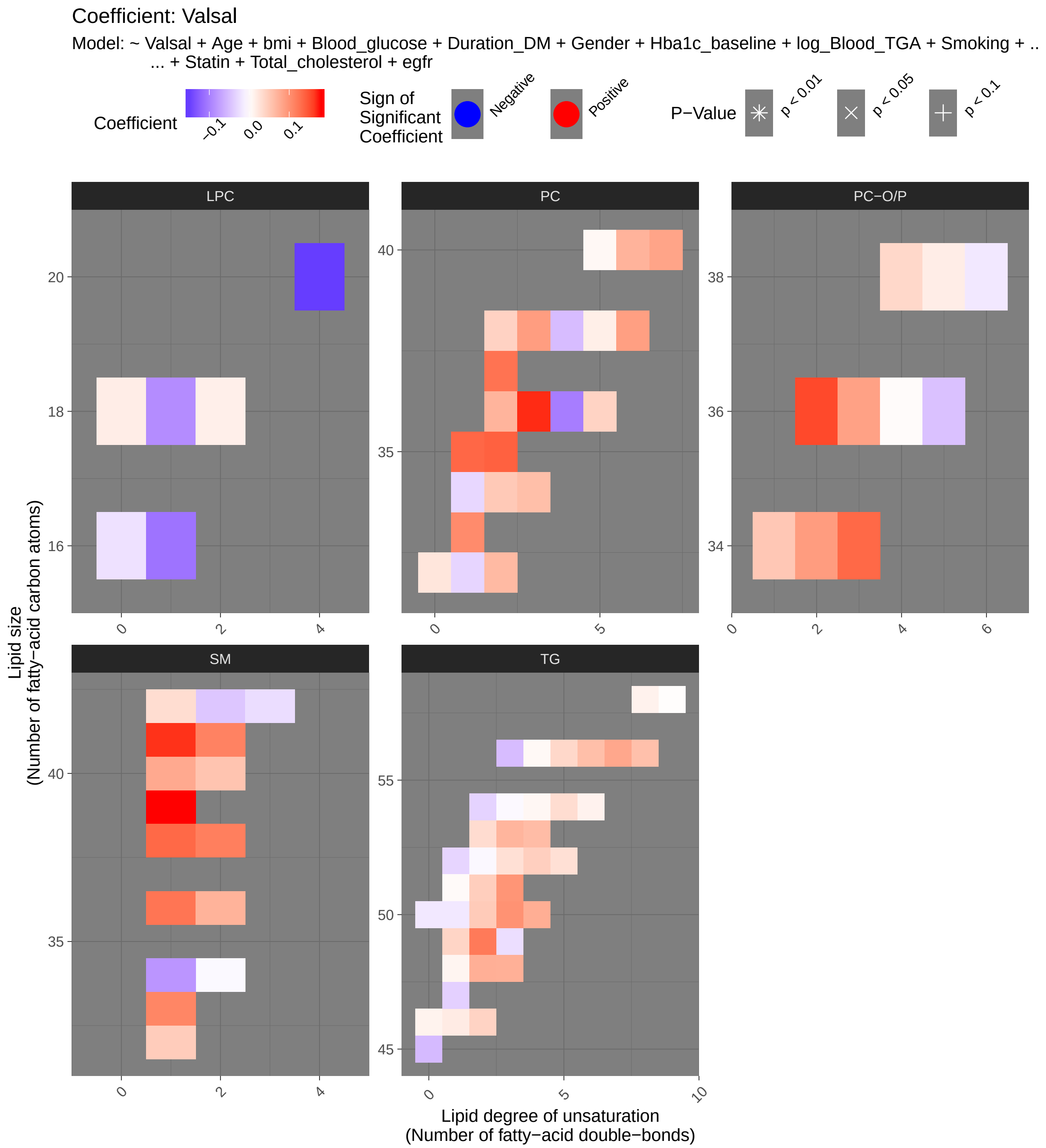
Coefficient: Valsal
Model: ~ Valsal + Age + bmi + Blood_glucose + Duration_DM + Gender + Hba1c_baseline + log_Blood_TGA + Smoking + ..
... + Statin + Total_cholesterol + egfr
Coefficient
−0.1
0.0
0.1
Sign of Significant Coefficient
Negative
Positive
P−Value
p < 0.01
p < 0.05
p < 0.1
LPC
PC
PC−O/P
SM
TG
Lipid size
(Number of fatty−acid carbon atoms)
Lipid degree of unsaturation
(Number of fatty−acid double−bonds)

###### 7.4.3.2 Tables of Model Coefficients

```
## [1] ""
## [1] "Table: Valsal"
## [1] "  (from model: "
## [1] "   ~ Valsal + Age + bmi + Blood_glucose + Duration_DM + Gender"
## [1] "     + Hba1c_baseline + log_Blood_TGA + Smoking + Statin +"
## [1] "     Total_cholesterol + egfr)"
## [1] ""
##                               Name Coefficient P.Value adj.P.Val
## 1           SM(d39:1)_[LVL2]; 179    0.179000  0.0136     0.565
## 2           SM(d41:1)_[LVL2]; 102    0.167000  0.0143     0.565
## 3             PC(36:3)_[LVL2]; 10    0.170000  0.0220     0.565
## 4             TG(50:3)_[LVL2]; 47    0.100000  0.0265     0.565
## 5          PC(O-36:2)_[LVL2]; 312    0.155000  0.0267     0.565
## 6          PC(O-34:3)_[LVL2]; 140    0.134000  0.0433     0.580
## 7           LPC(20:4)_[LVL2]; 120   -0.150000  0.0443     0.580
## 8            TG(49:2)_[LVL3]; 231    0.121000  0.0491     0.580
## 9            SM(d38:1)_[LVL2]; 67    0.134000  0.0519     0.580
## 10           PC(35:1)_[LVL2]; 178    0.135000  0.0557     0.580
## 11           SM(d36:1)_[LVL2]; 55    0.124000  0.0614     0.580
## 12           TG(51:3)_[LVL3]; 198    0.097800  0.0664     0.580
## 13           PC(35:2)_[LVL2]; 143    0.139000  0.0711     0.580
## 14           PC(37:2)_[LVL2]; 350    0.126000  0.0829     0.595
## 15          SM(d38:2)_[LVL2]; 151    0.116000  0.0843     0.595
## 16          SM(d41:2)_[LVL2]; 139    0.114000  0.0930     0.616
## 17          SM(d33:1)_[LVL2]; 166    0.111000  0.1080     0.672
## 18           PC(33:1)_[LVL2]; 177    0.106000  0.1370     0.807
## 19          LPC(16:1)_[LVL2]; 258   -0.109000  0.1550     0.809
## 20             PC(38:6)_[LVL2]; 8    0.088900  0.1740     0.809
## 21         PC(O-36:3)_[LVL2]; 268    0.086600  0.1740     0.809
## 22  TG(14:0/18:2/18:2)_[LVL2]; 189    0.075500  0.1820     0.809
## 23  TG(18:1/12:0/18:1) or TG(18:2/    0.074400  0.1850     0.809
## 24           TG(53:3)_[LVL3]; 239    0.069200  0.1920     0.809
## 25             PC(36:4)_[LVL2]; 1   -0.101000  0.2060     0.809
## 26            PC(38:3)_[LVL2]; 29    0.090700  0.2160     0.809
## 27         PC(O-34:2)_[LVL2]; 171    0.091700  0.2160     0.809
## 28           PC(40:7)_[LVL2]; 165    0.084700  0.2200     0.809
## 29           SM(d40:1)_[LVL2]; 39    0.079400  0.2270     0.809
## 30           TG(48:3)_[LVL3]; 384    0.073400  0.2290     0.809
## 31           LPC(18:1)_[LVL2]; 34   -0.089000  0.2510     0.810
## 32           TG(56:7)_[LVL3]; 309    0.081400  0.2510     0.810
## 33   TG(14:0/18:1/18:1)_[LVL2]; 25    0.052000  0.2620     0.810
## 34            TG(45:0)_[LVL2]; 65   -0.052900  0.2630     0.810
## 35           SM(d34:1)_[LVL2]; 26   -0.082000  0.2700     0.810
## 36            PC(40:6)_[LVL2]; 31    0.070600  0.2760     0.810
## 37          SM(d36:2)_[LVL2]; 160    0.071100  0.2830     0.810
## 38           TG(53:4)_[LVL3]; 314    0.062800  0.2920     0.816
## 39           TG(50:2)_[LVL3]; 167    0.049200  0.3130     0.852
## 40           TG(51:2)_[LVL2]; 123    0.046200  0.3520     0.894
## 41           TG(56:6)_[LVL3]; 275    0.060200  0.3550     0.894
## 42  TG(18:0/18:1/20:4)_[LVL2]; 141   -0.069600  0.3560     0.894
## 43             PC(36:2)_[LVL2]; 3    0.070300  0.3780     0.894
## 44           PC(32:2)_[LVL2]; 204    0.064700  0.3800     0.894
```

```
## 45            SM(d40:2)_[LVL2]; 80    0.055300  0.3860     0.894
## 46            TG(52:4)_[LVL3]; 157    0.044900  0.4130     0.894
## 47            TG(56:3)_[LVL2]; 290   -0.051200  0.4220     0.894
## 48  TG(16:0/18:2/22:6)_[LVL2]; 117    0.059100  0.4220     0.894
## 49  PC(16:0e/18:1(9Z))_[LVL1]; 134    0.052000  0.4260     0.894
## 50            PC(34:3)_[LVL2]; 113    0.060200  0.4340     0.894
## 51           SM(d32:1)_[LVL2]; 105    0.047800  0.4480     0.894
## 52  TG(16:0/18:2/18:3)_[LVL2]; 106    0.044900  0.4620     0.894
## 53   TG(18:1/18:1/18:1)_[LVL2]; 15   -0.039800  0.4710     0.894
## 54              PC(38:4)_[LVL2]; 9   -0.051300  0.4970     0.894
## 55          PC(O-36:5)_[LVL2]; 92   -0.047400  0.5150     0.894
## 56             SM(42:2)_[LVL2]; 14   -0.043700  0.5170     0.894
## 57             TG(54:2)_[LVL3]; 52   -0.033800  0.5200     0.894
## 58   TG(16:0/18:2/18:2)_[LVL2]; 27    0.033200  0.5210     0.894
## 59            TG(46:2)_[LVL3]; 248    0.040900  0.5210     0.894
## 60            TG(53:2)_[LVL2]; 234    0.032600  0.5240     0.894
## 61            TG(49:1)_[LVL3]; 187    0.039700  0.5260     0.894
## 62              PC(34:2)_[LVL2]; 4    0.050400  0.5360     0.894
## 63             PC(36:5)_[LVL2]; 23    0.041000  0.5400     0.894
## 64            TG(56:5)_[LVL2]; 230    0.036900  0.5400     0.894
## 65            PC(38:2)_[LVL2]; 197    0.042000  0.5700     0.922
## 66   TG(16:0/18:0/18:1)_[LVL2]; 51   -0.031800  0.5820     0.922
## 67            TG(52:3)_[LVL3]; 101    0.028800  0.5830     0.922
## 68            TG(54:5)_[LVL3]; 240    0.031800  0.6240     0.944
## 69          PC(O-38:4)_[LVL2]; 131    0.036700  0.6280     0.944
## 70            TG(52:5)_[LVL3]; 286    0.029300  0.6310     0.944
## 71       SM(d18:1/24:0)_[LVL2]; 61    0.031800  0.6320     0.944
## 72            TG(47:1)_[LVL3]; 227   -0.035700  0.6450     0.950
## 73             PC(32:1)_[LVL2]; 44   -0.031900  0.6650     0.966
## 74            TG(49:3)_[LVL3]; 218   -0.025000  0.6770     0.970
## 75       SM(d18:2/24:1)_[LVL2]; 40   -0.026100  0.6890     0.974
## 76              PC(34:1)_[LVL2]; 2   -0.030400  0.7140     0.980
## 77             PC(32:0)_[LVL2]; 96    0.023900  0.7300     0.980
## 78             TG(50:1)_[LVL3]; 19   -0.017200  0.7350     0.980
## 79             LPC(16:0)_[LVL1]; 5   -0.023100  0.7540     0.980
## 80            TG(46:1)_[LVL3]; 128    0.018800  0.7750     0.980
## 81   TG(18:2/18:1/18:1)_[LVL2]; 20   -0.015300  0.7840     0.980
## 82          PC(O-38:6)_[LVL2]; 236   -0.017400  0.7950     0.980
## 83    TG(18:1/18:1/16:0)_[LVL2]; 7   -0.016600  0.7980     0.980
## 84            TG(50:0)_[LVL2]; 159   -0.017500  0.8030     0.980
## 85   TG(18:2/22:5/16:0)_[LVL2]; 69    0.017700  0.8040     0.980
## 86  TG(16:0/22:5/18:1) or TG(20:4/    0.014600  0.8090     0.980
## 87          PC(O-38:5)_[LVL2]; 76    0.016700  0.8230     0.980
## 88             PC(38:5)_[LVL2]; 24    0.015500  0.8270     0.980
## 89            LPC(18:0)_[LVL1]; 22    0.016700  0.8290     0.980
## 90            LPC(18:2)_[LVL2]; 33    0.014800  0.8460     0.980
## 91            TG(54:6)_[LVL3]; 316    0.011900  0.8580     0.980
## 92  TG(18:1/18:1/22:6)_[LVL2]; 147    0.012500  0.8670     0.980
## 93            TG(46:0)_[LVL3]; 168    0.011500  0.8710     0.980
## 94   TG(14:0/16:0/18:1)_[LVL2]; 54    0.009580  0.8740     0.980
## 95            TG(54:4)_[LVL3]; 129    0.007610  0.9010     0.980
## 96             TG(52:2)_[LVL3]; 97   -0.004950  0.9150     0.980
## 97            TG(56:4)_[LVL3]; 278    0.005890  0.9210     0.980
## 98             PC(40:5)_[LVL2]; 95    0.006630  0.9220     0.980
```

```
## 99  SM(d16:1/18:1) or SM(d18:2/16:   -0.004820  0.9340     0.980
## 100           TG(54:3)_[LVL3]; 124   -0.004560  0.9350     0.980
## 101           TG(51:1)_[LVL3]; 249    0.004240  0.9410     0.980
## 102  TG(18:1/18:2/18:2)_[LVL2]; 57    0.003190  0.9580     0.980
## 103          PC(O-36:4)_[LVL2]; 71    0.003730  0.9600     0.980
## 104 TG(18:2/18:2/18:2) or TG(18:3/    0.003240  0.9610     0.980
## 105           TG(58:9)_[LVL3]; 207    0.001860  0.9810     0.990
## 106 TG(18:2/18:1/16:0)_[LVL2]; 500    0.000932  0.9900     0.990
```

#### 7.5 Heart Rate Variablity (SDNN)

##### 7.5.1 Crude Model

```
## [1] "Fitting models:"
## [1] "~ SDNN + rest_HR_vag"
## [1] ""
```

###### 7.5.1.1 Heatmap

```
## [1] "heatmap_lipidome_from_limma was created by Tommi Suvitaival"
## [1] ""
## [1] "2019-05-21"
```

```
## Warning: Removed 106 rows containing missing values (geom_point).
```

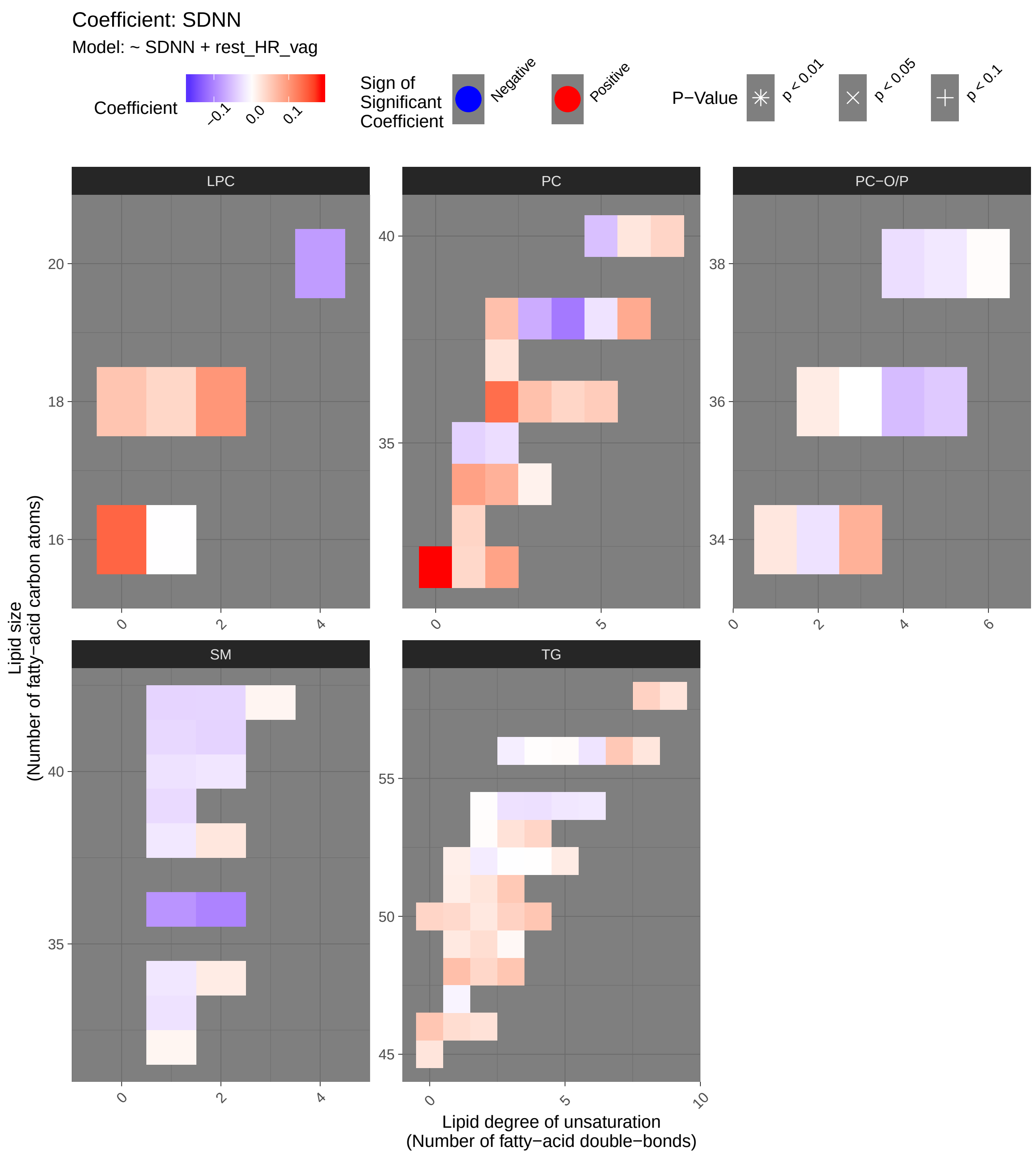

Coefficient: SDNN
Model: ~ SDNN + rest_HR_vag
Coefficient
−0.1
0.0
0.1
Sign of Significant Coefficient
Negative
Positive
P−Value
p < 0.01
p < 0.05
p < 0.1
LPC
PC
PC−O/P
SM
TG
Lipid size
(Number of fatty−acid carbon atoms)
Lipid degree of unsaturation
(Number of fatty−acid double−bonds)

###### 7.5.1.2 Tables of Model Coefficients

```
## [1] ""
## [1] "Table: SDNN"
## [1] "  (from model: "
## [1] "   ~ SDNN + rest_HR_vag)"
## [1] ""
##                                Name Coefficient P.Value adj.P.Val
## 1              PC(32:0)_[LVL2]; 96    0.188000 0.00298     0.316
## 2   TG(18:0/18:1/20:4)_[LVL2]; 141   -0.165000 0.00925     0.490
## 3              LPC(16:0)_[LVL1]; 5    0.143000 0.02380     0.833
## 4               PC(36:2)_[LVL2]; 3    0.136000 0.03140     0.833
## 5               PC(38:4)_[LVL2]; 9   -0.109000 0.08520     0.974
## 6             LPC(18:2)_[LVL2]; 33    0.102000 0.10800     0.974
## 7            SM(d36:2)_[LVL2]; 160   -0.100000 0.11300     0.974
## 8               PC(34:1)_[LVL2]; 2    0.091100 0.15000     0.974
## 9             PC(32:2)_[LVL2]; 204    0.089200 0.15900     0.974
## 10            SM(d36:1)_[LVL2]; 55   -0.086400 0.17200     0.974
## 11              PC(38:6)_[LVL2]; 8    0.082600 0.19200     0.974
## 12           LPC(20:4)_[LVL2]; 120   -0.079800 0.20700     0.974
## 13         PC(O-34:3)_[LVL2]; 140    0.076300 0.22800     0.974
## 14              PC(34:2)_[LVL2]; 4    0.075600 0.23200     0.974
## 15             PC(38:3)_[LVL2]; 29   -0.066900 0.29100     0.974
## 16   TG(14:0/16:0/18:1)_[LVL2]; 54    0.062900 0.32000     0.974
## 17            PC(38:2)_[LVL2]; 197    0.061300 0.33300     0.974
## 18             PC(36:3)_[LVL2]; 10    0.061100 0.33400     0.974
## 19            LPC(18:0)_[LVL1]; 22    0.057000 0.36700     0.974
## 20            TG(48:3)_[LVL3]; 384    0.056300 0.37300     0.974
## 21            TG(46:0)_[LVL3]; 168    0.056000 0.37600     0.974
## 22  TG(14:0/18:2/18:2)_[LVL2]; 189    0.055900 0.37700     0.974
## 23           PC(O-36:4)_[LVL2]; 71   -0.053900 0.39500     0.974
## 24            TG(56:7)_[LVL3]; 309    0.053500 0.39800     0.974
## 25            TG(51:3)_[LVL3]; 198    0.053200 0.40000     0.974
## 26             PC(40:5)_[LVL2]; 95   -0.051000 0.42000     0.974
## 27             PC(36:5)_[LVL2]; 23    0.049800 0.43200     0.974
## 28  TG(18:1/18:1/22:6)_[LVL2]; 147    0.044100 0.48600     0.974
## 29             TG(50:3)_[LVL2]; 47    0.043900 0.48800     0.974
## 30           PC(O-36:5)_[LVL2]; 92   -0.043700 0.49000     0.974
## 31            PC(33:1)_[LVL2]; 177    0.041200 0.51500     0.974
## 32            TG(53:4)_[LVL3]; 314    0.041000 0.51700     0.974
## 33            TG(50:0)_[LVL2]; 159    0.041000 0.51700     0.974
## 34            PC(40:7)_[LVL2]; 165    0.041000 0.51700     0.974
## 35              PC(36:4)_[LVL2]; 1    0.040600 0.52100     0.974
## 36            LPC(18:1)_[LVL2]; 34    0.039700 0.53000     0.974
## 37  TG(18:1/12:0/18:1) or TG(18:2/    0.039400 0.53300     0.974
## 38             PC(32:1)_[LVL2]; 44    0.038400 0.54400     0.974
## 39  TG(18:2/18:2/18:2) or TG(18:3/    0.037700 0.55200     0.974
## 40             TG(50:1)_[LVL3]; 19    0.037300 0.55600     0.974
## 41            PC(35:1)_[LVL2]; 178   -0.036400 0.56500     0.974
## 42   TG(16:0/18:2/18:2)_[LVL2]; 27    0.035100 0.57900     0.974
## 43           SM(d41:2)_[LVL2]; 139   -0.035000 0.58100     0.974
## 44      SM(d18:1/24:0)_[LVL2]; 61   -0.034200 0.58800     0.974
## 45             SM(42:2)_[LVL2]; 14   -0.033800 0.59300     0.974
## 46            TG(46:1)_[LVL3]; 128    0.033500 0.59600     0.974
```

```
## 47           TG(49:2)_[LVL3]; 231    0.032800 0.60500      0.974
## 48   TG(18:1/18:2/18:2)_[LVL2]; 57    0.031800 0.61500      0.974
## 49          SM(d41:1)_[LVL2]; 102   -0.031300 0.62100      0.974
## 50          SM(d39:1)_[LVL2]; 179   -0.029300 0.64400      0.974
## 51           TG(53:3)_[LVL3]; 239    0.028600 0.65100      0.974
## 52           TG(46:2)_[LVL3]; 248    0.028500 0.65300      0.974
## 53   TG(18:2/18:1/18:1)_[LVL2]; 20    0.027700 0.66100      0.974
## 54           PC(37:2)_[LVL2]; 350    0.027400 0.66500      0.974
## 55           PC(35:2)_[LVL2]; 143   -0.026900 0.67000      0.974
## 56         PC(O-38:4)_[LVL2]; 131   -0.026700 0.67300      0.974
## 57           TG(58:9)_[LVL3]; 207    0.026300 0.67800      0.974
## 58           TG(51:2)_[LVL2]; 123    0.026100 0.68000      0.974
## 59            TG(45:0)_[LVL2]; 65    0.025700 0.68500      0.974
## 60           TG(54:4)_[LVL3]; 129   -0.025000 0.69300      0.974
## 61  TG(16:0/18:2/22:6)_[LVL2]; 117    0.024900 0.69400      0.974
## 62            PC(40:6)_[LVL2]; 31    0.024800 0.69500      0.974
## 63           TG(54:3)_[LVL3]; 124   -0.024200 0.70200      0.974
## 64          SM(d38:2)_[LVL2]; 151    0.023900 0.70600      0.974
## 65  PC(16:0e/18:1(9Z))_[LVL1]; 134    0.023500 0.71100      0.974
## 66         PC(O-34:2)_[LVL2]; 171   -0.023200 0.71400      0.974
## 67           SM(d40:1)_[LVL2]; 39   -0.023100 0.71500      0.974
## 68          SM(d33:1)_[LVL2]; 166   -0.023100 0.71600      0.974
## 69           TG(50:2)_[LVL3]; 167    0.022300 0.72500      0.974
## 70            PC(38:5)_[LVL2]; 24   -0.022200 0.72600      0.974
## 71           TG(49:1)_[LVL3]; 187    0.021900 0.72900      0.974
## 72           TG(56:6)_[LVL3]; 275   -0.021800 0.73000      0.974
## 73   TG(18:2/22:5/16:0)_[LVL2]; 69    0.021200 0.73800      0.974
## 74  TG(16:0/18:2/18:3)_[LVL2]; 106    0.019900 0.75300      0.974
## 75           SM(d40:2)_[LVL2]; 80   -0.019900 0.75400      0.974
## 76   TG(14:0/18:1/18:1)_[LVL2]; 25    0.019600 0.75700      0.974
## 77           TG(54:5)_[LVL3]; 240   -0.019300 0.76000      0.974
## 78           SM(d34:1)_[LVL2]; 26   -0.019300 0.76100      0.974
## 79         PC(O-36:2)_[LVL2]; 312    0.019000 0.76400      0.974
## 80           TG(52:5)_[LVL3]; 286    0.018800 0.76700      0.974
## 81           SM(d38:1)_[LVL2]; 67   -0.018700 0.76800      0.974
## 82  SM(d16:1/18:1) or SM(d18:2/16:    0.018600 0.76800      0.974
## 83          PC(O-38:5)_[LVL2]; 76   -0.018300 0.77300      0.974
## 84           TG(54:6)_[LVL3]; 316   -0.017900 0.77700      0.974
## 85    TG(18:1/18:1/16:0)_[LVL2]; 7   -0.017100 0.78700      0.974
## 86           TG(51:1)_[LVL3]; 249    0.016400 0.79500      0.974
## 87   TG(16:0/18:0/18:1)_[LVL2]; 51    0.015500 0.80700      0.974
## 88            TG(52:2)_[LVL3]; 97   -0.015400 0.80800      0.974
## 89           TG(56:3)_[LVL2]; 290   -0.013600 0.83000      0.984
## 90           PC(34:3)_[LVL2]; 113    0.013100 0.83600      0.984
## 91       SM(d18:2/24:1)_[LVL2]; 40    0.009530 0.88000      0.996
## 92          SM(d32:1)_[LVL2]; 105    0.009360 0.88200      0.996
## 93           TG(47:1)_[LVL3]; 227   -0.008960 0.88700      0.996
## 94           TG(49:3)_[LVL3]; 218    0.006550 0.91800      0.996
## 95  TG(18:2/18:1/16:0)_[LVL2]; 500   -0.005020 0.93700      0.996
## 96  TG(16:0/22:5/18:1) or TG(20:4/   -0.004430 0.94400      0.996
## 97           TG(56:5)_[LVL2]; 230    0.003900 0.95100      0.996
## 98         PC(O-38:6)_[LVL2]; 236    0.003010 0.96200      0.996
## 99           TG(53:2)_[LVL2]; 234    0.002820 0.96400      0.996
## 100  TG(18:1/18:1/18:1)_[LVL2]; 15    0.002180 0.97200      0.996
```

```
## 101           TG(56:4)_[LVL3]; 278    0.001670 0.97900     0.996
## 102            TG(54:2)_[LVL3]; 52    0.001590 0.98000     0.996
## 103           TG(52:3)_[LVL3]; 101   -0.000857 0.98900     0.996
## 104          LPC(16:1)_[LVL2]; 258   -0.000672 0.99200     0.996
## 105           TG(52:4)_[LVL3]; 157    0.000514 0.99400     0.996
## 106         PC(O-36:3)_[LVL2]; 268   -0.000300 0.99600     0.996
```

##### 7.5.2 Adjusted Model

```
## [1] "Fitting models:"
## [1] "~ SDNN + rest_HR_vag + Age + bmi + Blood_glucose + Duration_DM + Gender + Hba1c_baseline + log_
## [1] ""
```

###### 7.5.2.1 Heatmap

```
## [1] "heatmap_lipidome_from_limma was created by Tommi Suvitaival"
## [1] ""
## [1] "2019-05-21"
```

```
## Warning: Removed 105 rows containing missing values (geom_point).
```

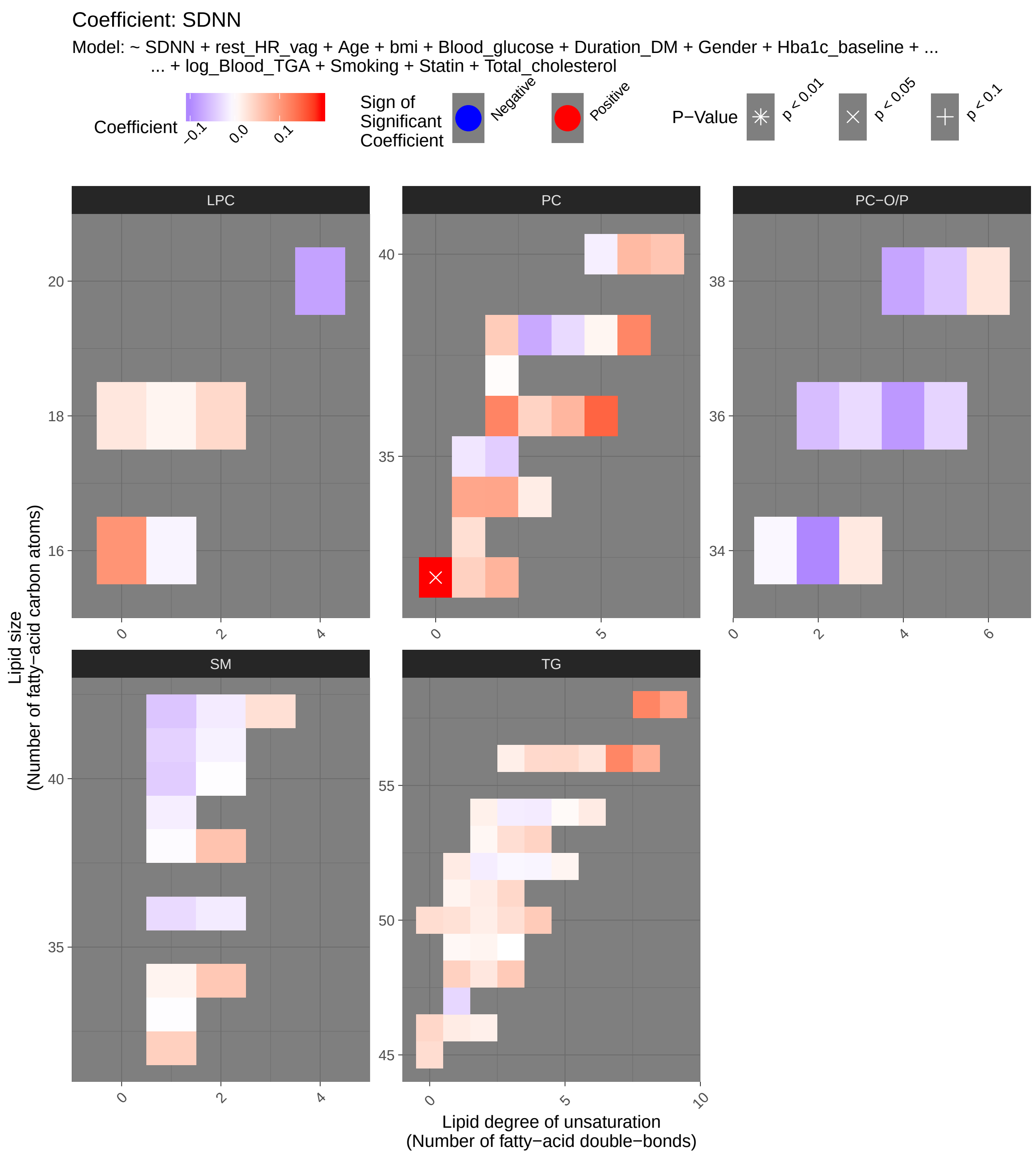

Coefficient: SDNN
Model: ~ SDNN + rest_HR_vag + Age + bmi + Blood_glucose + Duration_DM + Gender + Hba1c_baseline + ...
... + log_Blood_TGA + Smoking + Statin + Total_cholesterol
Coefficient
−0.1
0.0
0.1
Sign of Significant Coefficient
Negative
Positive
P−Value
p < 0.01
p < 0.05
p < 0.1
LPC
PC
PC−O/P
SM
TG
Lipid size (Number of fatty−acid carbon atoms)
Lipid degree of unsaturation (Number of fatty−acid double−bonds)

###### 7.5.2.2 Tables of Model Coefficients

```
## [1] ""
## [1] "Table: SDNN"
## [1] "  (from model: "
## [1] "   ~ SDNN + rest_HR_vag + Age + bmi + Blood_glucose +"
## [1] "     Duration_DM + Gender + Hba1c_baseline + log_Blood_TGA +"
## [1] "     Smoking + Statin + Total_cholesterol)"
## [1] ""
##                               Name Coefficient  P.Value adj.P.Val
## 1             PC(32:0)_[LVL2]; 96    0.194000 0.000457    0.0484
## 2             PC(36:5)_[LVL2]; 23    0.149000 0.006990    0.3700
## 3              PC(38:6)_[LVL2]; 8    0.119000 0.027200    0.8640
## 4            TG(56:7)_[LVL3]; 309    0.120000 0.032600    0.8640
## 5   TG(18:1/18:1/22:6)_[LVL2]; 147    0.121000 0.042100    0.8930
## 6              PC(36:2)_[LVL2]; 3    0.121000 0.064100    0.9500
## 7   TG(18:2/18:2/18:2) or TG(18:3/    0.095100 0.065100    0.9500
## 8             LPC(16:0)_[LVL1]; 5    0.107000 0.071700    0.9500
## 9          PC(O-34:2)_[LVL2]; 171   -0.101000 0.095000    0.9580
## 10  TG(18:0/18:1/20:4)_[LVL2]; 141   -0.092200 0.113000    0.9580
## 11   TG(18:2/22:5/16:0)_[LVL2]; 69    0.082300 0.135000    0.9580
## 12           TG(58:9)_[LVL3]; 207    0.091700 0.136000    0.9580
## 13             PC(34:2)_[LVL2]; 4    0.090300 0.158000    0.9580
## 14  TG(16:0/18:2/22:6)_[LVL2]; 117    0.080200 0.163000    0.9580
## 15          PC(O-36:4)_[LVL2]; 71   -0.086300 0.169000    0.9580
## 16             PC(34:1)_[LVL2]; 2    0.088700 0.182000    0.9580
## 17            PC(40:6)_[LVL2]; 31    0.070100 0.187000    0.9580
## 18          LPC(20:4)_[LVL2]; 120   -0.077900 0.201000    0.9580
## 19           PC(32:2)_[LVL2]; 204    0.075200 0.209000    0.9580
## 20            PC(38:3)_[LVL2]; 29   -0.071800 0.214000    0.9580
## 21  TG(14:0/18:2/18:2)_[LVL2]; 189    0.053100 0.224000    0.9580
## 22         PC(O-38:4)_[LVL2]; 131   -0.075100 0.227000    0.9580
## 23  SM(d16:1/18:1) or SM(d18:2/16:    0.055700 0.234000    0.9580
## 24   TG(18:1/18:2/18:2)_[LVL2]; 57    0.059200 0.237000    0.9580
## 25             PC(36:4)_[LVL2]; 1    0.073600 0.254000    0.9580
## 26           TG(48:3)_[LVL3]; 384    0.053500 0.267000    0.9580
## 27          SM(d38:2)_[LVL2]; 151    0.060300 0.274000    0.9580
## 28           PC(40:7)_[LVL2]; 165    0.058300 0.287000    0.9580
## 29   TG(14:0/16:0/18:1)_[LVL2]; 54    0.046900 0.316000    0.9580
## 30           TG(51:3)_[LVL3]; 198    0.040100 0.332000    0.9580
## 31           TG(53:4)_[LVL3]; 314    0.044400 0.337000    0.9580
## 32         PC(O-36:2)_[LVL2]; 312   -0.054800 0.343000    0.9580
## 33          SM(d32:1)_[LVL2]; 105    0.048000 0.344000    0.9580
## 34      SM(d18:1/24:0)_[LVL2]; 61   -0.047900 0.350000    0.9580
## 35            TG(45:0)_[LVL2]; 65    0.034700 0.352000    0.9580
## 36            TG(50:3)_[LVL2]; 47    0.032500 0.355000    0.9580
## 37           PC(38:2)_[LVL2]; 197    0.052100 0.363000    0.9580
## 38           TG(56:5)_[LVL2]; 230    0.038800 0.396000    0.9580
## 39           TG(56:4)_[LVL3]; 278    0.038500 0.399000    0.9580
## 40           SM(d40:1)_[LVL2]; 39   -0.042200 0.403000    0.9580
## 41           TG(53:3)_[LVL3]; 239    0.033200 0.414000    0.9580
## 42            TG(50:1)_[LVL3]; 19    0.030600 0.426000    0.9580
## 43  TG(16:0/22:5/18:1) or TG(20:4/    0.035500 0.440000    0.9580
## 44          PC(O-38:5)_[LVL2]; 76   -0.048200 0.443000    0.9580
```

```
## 45             PC(32:1)_[LVL2]; 44    0.046500 0.443000    0.9580
## 46             PC(36:3)_[LVL2]; 10    0.044800 0.448000    0.9580
## 47   TG(18:2/18:1/18:1)_[LVL2]; 20    0.034700 0.464000    0.9580
## 48            TG(46:0)_[LVL3]; 168    0.040600 0.471000    0.9580
## 49           SM(d41:1)_[LVL2]; 102   -0.038000 0.477000    0.9580
## 50   TG(16:0/18:2/18:2)_[LVL2]; 27    0.028100 0.487000    0.9580
## 51            PC(35:2)_[LVL2]; 143   -0.041100 0.501000    0.9580
## 52            LPC(18:2)_[LVL2]; 33    0.039000 0.529000    0.9580
## 53            TG(50:0)_[LVL2]; 159    0.034400 0.531000    0.9580
## 54  TG(16:0/18:2/18:3)_[LVL2]; 106    0.030300 0.536000    0.9580
## 55       SM(d18:2/24:1)_[LVL2]; 40    0.031500 0.559000    0.9580
## 56           PC(O-36:5)_[LVL2]; 92   -0.035100 0.567000    0.9580
## 57            PC(33:1)_[LVL2]; 177    0.032600 0.570000    0.9580
## 58          PC(O-36:3)_[LVL2]; 268   -0.030200 0.573000    0.9580
## 59            SM(d36:1)_[LVL2]; 55   -0.030100 0.579000    0.9580
## 60            TG(56:6)_[LVL3]; 275    0.027500 0.581000    0.9580
## 61  TG(18:1/12:0/18:1) or TG(18:2/    0.024000 0.595000    0.9580
## 62            TG(47:1)_[LVL3]; 227   -0.033300 0.597000    0.9580
## 63               PC(38:4)_[LVL2]; 9   -0.030700 0.610000    0.9580
## 64            TG(51:2)_[LVL2]; 123    0.018900 0.616000    0.9580
## 65   TG(16:0/18:0/18:1)_[LVL2]; 51    0.020500 0.640000    0.9580
## 66   TG(14:0/18:1/18:1)_[LVL2]; 25    0.016800 0.642000    0.9580
## 67          PC(O-38:6)_[LVL2]; 236    0.026100 0.648000    0.9580
## 68            TG(50:2)_[LVL3]; 167    0.017100 0.652000    0.9580
## 69             TG(52:2)_[LVL3]; 97   -0.015100 0.668000    0.9580
## 70          PC(O-34:3)_[LVL2]; 140    0.022000 0.684000    0.9580
## 71            LPC(18:0)_[LVL1]; 22    0.024300 0.688000    0.9580
## 72            TG(54:6)_[LVL3]; 316    0.020100 0.703000    0.9580
## 73             TG(54:2)_[LVL3]; 52    0.014500 0.708000    0.9580
## 74    TG(18:1/18:1/16:0)_[LVL2]; 7   -0.019500 0.712000    0.9580
## 75            PC(35:1)_[LVL2]; 178   -0.020600 0.716000    0.9580
## 76            TG(46:1)_[LVL3]; 128    0.019000 0.725000    0.9580
## 77            TG(54:3)_[LVL3]; 124   -0.014700 0.733000    0.9580
## 78            TG(54:4)_[LVL3]; 129   -0.016300 0.743000    0.9580
## 79            TG(56:3)_[LVL2]; 290    0.016200 0.743000    0.9580
## 80            PC(34:3)_[LVL2]; 113    0.018200 0.757000    0.9580
## 81           SM(d36:2)_[LVL2]; 160   -0.016700 0.765000    0.9580
## 82             SM(42:2)_[LVL2]; 14   -0.016000 0.769000    0.9580
## 83            TG(46:2)_[LVL3]; 248    0.015100 0.775000    0.9580
## 84  TG(18:2/18:1/16:0)_[LVL2]; 500   -0.016100 0.785000    0.9580
## 85            TG(51:1)_[LVL3]; 249    0.011400 0.802000    0.9580
## 86           SM(d39:1)_[LVL2]; 179   -0.013800 0.808000    0.9580
## 87             PC(40:5)_[LVL2]; 95   -0.012800 0.810000    0.9580
## 88            TG(49:2)_[LVL3]; 231    0.010700 0.831000    0.9580
## 89           SM(d41:2)_[LVL2]; 139   -0.011000 0.836000    0.9580
## 90            TG(53:2)_[LVL2]; 234    0.007780 0.843000    0.9580
## 91            TG(52:5)_[LVL3]; 286    0.009680 0.843000    0.9580
## 92            TG(52:4)_[LVL3]; 157   -0.008090 0.851000    0.9580
## 93            SM(d34:1)_[LVL2]; 26    0.011000 0.858000    0.9580
## 94            LPC(18:1)_[LVL2]; 34    0.010600 0.863000    0.9580
## 95             PC(38:5)_[LVL2]; 24    0.009570 0.864000    0.9580
## 96            TG(52:3)_[LVL3]; 101   -0.006690 0.867000    0.9580
## 97           LPC(16:1)_[LVL2]; 258   -0.009360 0.882000    0.9640
## 98            TG(49:1)_[LVL3]; 187    0.007010 0.891000    0.9640
```

```
## 99  PC(16:0e/18:1(9Z))_[LVL1]; 134   -0.006920 0.902000    0.9640
## 100           TG(54:5)_[LVL3]; 240    0.005470 0.917000    0.9640
## 101  TG(18:1/18:1/18:1)_[LVL2]; 15   -0.004520 0.919000    0.9640
## 102           SM(d38:1)_[LVL2]; 67   -0.003270 0.951000    0.9870
## 103           PC(37:2)_[LVL2]; 350    0.002700 0.963000    0.9870
## 104          SM(d33:1)_[LVL2]; 166   -0.001650 0.976000    0.9870
## 105           SM(d40:2)_[LVL2]; 80   -0.001400 0.978000    0.9870
## 106           TG(49:3)_[LVL3]; 218    0.000106 0.998000    0.9980
```

##### 7.5.3 Fully-Adjusted Model

```
## [1] "Fitting models:"
## [1] "~ SDNN + rest_HR_vag + Age + bmi + Blood_glucose + Duration_DM + Gender + Hba1c_baseline + log_[illegible]
## [1] ""
```

###### 7.5.3.1 Heatmap

```
## [1] "heatmap_lipidome_from_limma was created by Tommi Suvitaival"
## [1] ""
## [1] "2019-05-21"

## Warning: Removed 105 rows containing missing values (geom_point).
```

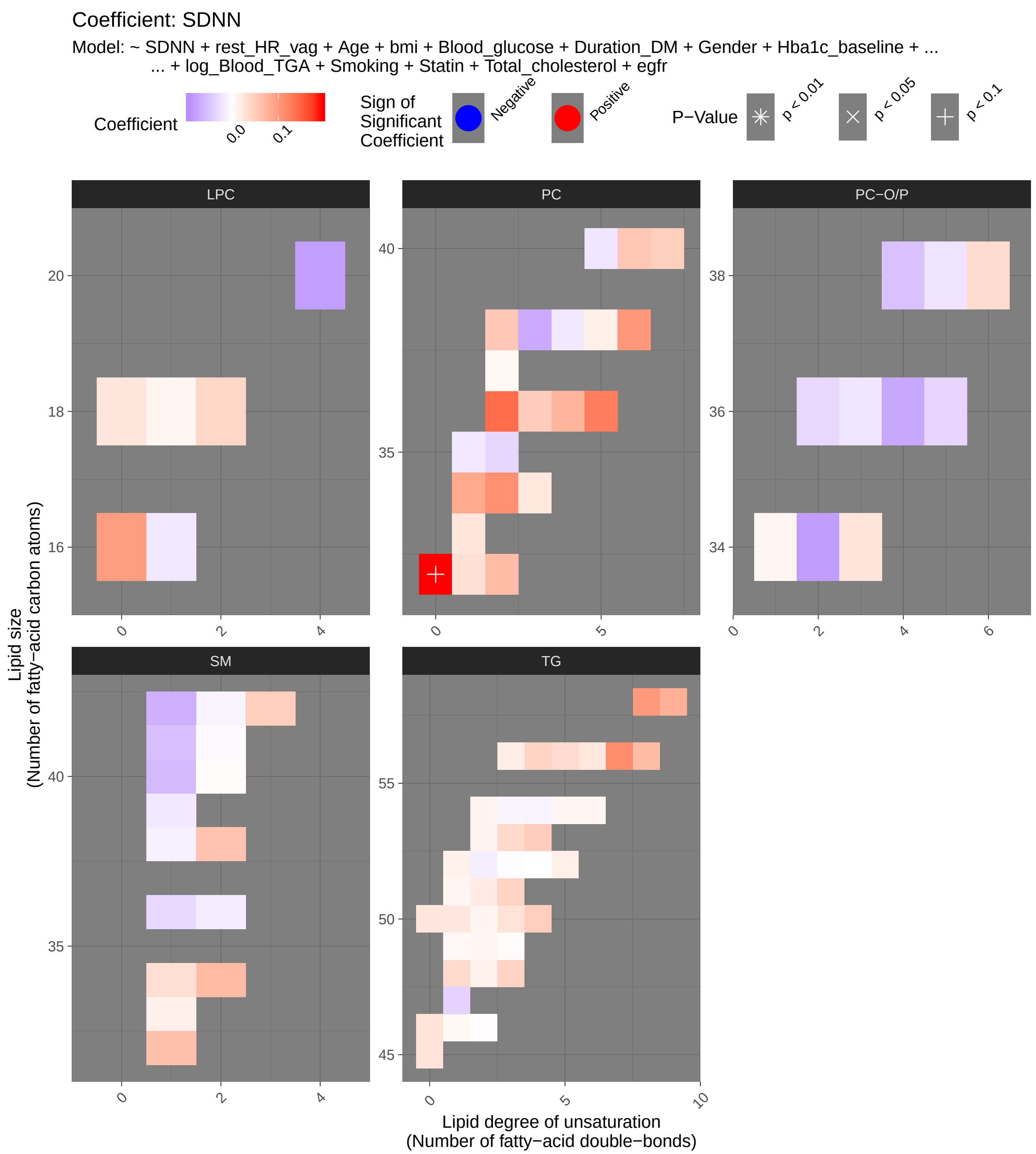
Coefficient: SDNN
Model: ~ SDNN + rest_HR_vag + Age + bmi + Blood_glucose + Duration_DM + Gender + Hba1c_baseline + ...
... + log_Blood_TGA + Smoking + Statin + Total_cholesterol + egfr
Coefficient
0.0
0.1
Sign of Significant Coefficient
Negative
Positive
P−Value
p < 0.01
p < 0.05
p < 0.1
LPC
PC
PC−O/P
SM
TG
Lipid size
(Number of fatty−acid carbon atoms)
Lipid degree of unsaturation
(Number of fatty−acid double−bonds)

###### 7.5.3.2 Tables of Model Coefficients

```
## [1] ""
## [1] "Table: SDNN"
## [1] "  (from model: "
## [1] "   ~ SDNN + rest_HR_vag + Age + bmi + Blood_glucose +"
## [1] "     Duration_DM + Gender + Hba1c_baseline + log_Blood_TGA +"
## [1] "     Smoking + Statin + Total_cholesterol + egfr)"
## [1] ""
##                               Name Coefficient  P.Value adj.P.Val
## 1             PC(32:0)_[LVL2]; 96    1.93e-01 0.000561    0.0595
## 2             PC(36:5)_[LVL2]; 23    1.26e-01 0.021500    0.9600
## 3              PC(36:2)_[LVL2]; 3    1.40e-01 0.032100    0.9600
## 4            TG(56:7)_[LVL3]; 309    1.12e-01 0.046300    0.9600
## 5              PC(38:6)_[LVL2]; 8    1.02e-01 0.058600    0.9600
## 6              PC(34:2)_[LVL2]; 4    1.09e-01 0.088700    0.9600
## 7   TG(18:1/18:1/22:6)_[LVL2]; 147    1.01e-01 0.089000    0.9600
## 8             LPC(16:0)_[LVL1]; 5    9.71e-02 0.104000    0.9600
## 9   TG(18:2/18:2/18:2) or TG(18:3/    8.17e-02 0.114000    0.9600
## 10  TG(18:0/18:1/20:4)_[LVL2]; 141   -9.11e-02 0.121000    0.9600
## 11  SM(d16:1/18:1) or SM(d18:2/16:    6.86e-02 0.144000    0.9600
## 12         PC(O-34:2)_[LVL2]; 171   -8.11e-02 0.179000    0.9600
## 13          LPC(20:4)_[LVL2]; 120   -7.97e-02 0.196000    0.9600
## 14           TG(58:9)_[LVL3]; 207    7.90e-02 0.202000    0.9600
## 15   TG(18:2/22:5/16:0)_[LVL2]; 69    7.07e-02 0.202000    0.9600
## 16             PC(34:1)_[LVL2]; 2    8.56e-02 0.202000    0.9600
## 17      SM(d18:1/24:0)_[LVL2]; 61   -6.52e-02 0.204000    0.9600
## 18          SM(d32:1)_[LVL2]; 105    6.32e-02 0.214000    0.9600
## 19            PC(38:3)_[LVL2]; 29   -7.03e-02 0.228000    0.9600
## 20   TG(18:1/18:2/18:2)_[LVL2]; 57    5.99e-02 0.236000    0.9600
## 21          PC(O-36:4)_[LVL2]; 71   -7.33e-02 0.245000    0.9600
## 22  TG(16:0/18:2/22:6)_[LVL2]; 117    6.66e-02 0.249000    0.9600
## 23             PC(36:4)_[LVL2]; 1    7.49e-02 0.251000    0.9600
## 24           SM(d40:1)_[LVL2]; 39   -5.80e-02 0.251000    0.9600
## 25  TG(14:0/18:2/18:2)_[LVL2]; 189    4.97e-02 0.260000    0.9600
## 26           PC(32:2)_[LVL2]; 204    6.65e-02 0.270000    0.9600
## 27           TG(53:4)_[LVL3]; 314    5.07e-02 0.277000    0.9600
## 28          SM(d38:2)_[LVL2]; 151    5.95e-02 0.286000    0.9600
## 29            PC(40:6)_[LVL2]; 31    5.62e-02 0.292000    0.9600
## 30           TG(51:3)_[LVL3]; 198    4.30e-02 0.303000    0.9600
## 31          SM(d41:1)_[LVL2]; 102   -5.24e-02 0.329000    0.9600
## 32           PC(38:2)_[LVL2]; 197    5.49e-02 0.343000    0.9600
## 33           TG(56:4)_[LVL3]; 278    4.31e-02 0.349000    0.9600
## 34           TG(53:3)_[LVL3]; 239    3.78e-02 0.356000    0.9600
## 35      SM(d18:2/24:1)_[LVL2]; 40    4.94e-02 0.358000    0.9600
## 36           TG(48:3)_[LVL3]; 384    4.38e-02 0.367000    0.9600
## 37           PC(40:7)_[LVL2]; 165    4.87e-02 0.377000    0.9600
## 38            PC(36:3)_[LVL2]; 10    5.21e-02 0.382000    0.9600
## 39         PC(O-38:4)_[LVL2]; 131   -5.17e-02 0.404000    0.9600
## 40            TG(50:3)_[LVL2]; 47    2.92e-02 0.412000    0.9600
## 41   TG(18:2/18:1/18:1)_[LVL2]; 20    3.92e-02 0.413000    0.9600
## 42   TG(14:0/16:0/18:1)_[LVL2]; 54    3.71e-02 0.430000    0.9600
## 43           TG(56:5)_[LVL2]; 230    3.61e-02 0.435000    0.9600
## 44   TG(16:0/18:2/18:2)_[LVL2]; 27    3.14e-02 0.440000    0.9600
```

```
## 45             TG(45:0)_[LVL2]; 65    2.90e-02 0.441000    0.9600
## 46  TG(16:0/18:2/18:3)_[LVL2]; 106    3.70e-02 0.455000    0.9600
## 47  TG(16:0/22:5/18:1) or TG(20:4/    3.24e-02 0.485000    0.9600
## 48            LPC(18:2)_[LVL2]; 33    4.07e-02 0.516000    0.9600
## 49          PC(O-38:6)_[LVL2]; 236    3.55e-02 0.537000    0.9600
## 50             TG(50:1)_[LVL3]; 19    2.39e-02 0.537000    0.9600
## 51            SM(d36:1)_[LVL2]; 55   -3.31e-02 0.547000    0.9600
## 52            TG(47:1)_[LVL3]; 227   -3.66e-02 0.565000    0.9600
## 53          PC(O-36:2)_[LVL2]; 312   -3.21e-02 0.577000    0.9600
## 54           PC(O-36:5)_[LVL2]; 92   -3.41e-02 0.582000    0.9600
## 55            PC(35:2)_[LVL2]; 143   -3.32e-02 0.590000    0.9600
## 56            SM(d34:1)_[LVL2]; 26    3.28e-02 0.592000    0.9600
## 57            TG(51:2)_[LVL2]; 123    2.02e-02 0.597000    0.9600
## 58             PC(32:1)_[LVL2]; 44    3.18e-02 0.602000    0.9600
## 59            TG(46:0)_[LVL3]; 168    2.86e-02 0.614000    0.9600
## 60            TG(56:6)_[LVL3]; 275    2.48e-02 0.622000    0.9600
## 61          PC(O-34:3)_[LVL2]; 140    2.68e-02 0.623000    0.9600
## 62            TG(50:0)_[LVL2]; 159    2.64e-02 0.634000    0.9600
## 63            PC(33:1)_[LVL2]; 177    2.65e-02 0.647000    0.9600
## 64            LPC(18:0)_[LVL1]; 22    2.52e-02 0.681000    0.9600
## 65            PC(34:3)_[LVL2]; 113    2.41e-02 0.686000    0.9600
## 66             TG(52:2)_[LVL3]; 97   -1.39e-02 0.697000    0.9600
## 67             PC(40:5)_[LVL2]; 95   -2.08e-02 0.700000    0.9600
## 68          PC(O-36:3)_[LVL2]; 268   -2.05e-02 0.704000    0.9600
## 69           PC(O-38:5)_[LVL2]; 76   -2.27e-02 0.717000    0.9600
## 70            TG(56:3)_[LVL2]; 290    1.81e-02 0.718000    0.9600
## 71            PC(35:1)_[LVL2]; 178   -1.92e-02 0.738000    0.9600
## 72           SM(d39:1)_[LVL2]; 179   -1.90e-02 0.740000    0.9600
## 73            TG(52:5)_[LVL3]; 286    1.60e-02 0.747000    0.9600
## 74  TG(18:1/12:0/18:1) or TG(18:2/    1.46e-02 0.749000    0.9600
## 75    TG(18:1/18:1/16:0)_[LVL2]; 7   -1.71e-02 0.749000    0.9600
## 76           LPC(16:1)_[LVL2]; 258   -1.96e-02 0.757000    0.9600
## 77            TG(53:2)_[LVL2]; 234    1.21e-02 0.760000    0.9600
## 78  TG(18:2/18:1/16:0)_[LVL2]; 500   -1.82e-02 0.761000    0.9600
## 79             TG(54:2)_[LVL3]; 52    1.17e-02 0.765000    0.9600
## 80   TG(14:0/18:1/18:1)_[LVL2]; 25    1.08e-02 0.766000    0.9600
## 81   TG(16:0/18:0/18:1)_[LVL2]; 51    1.29e-02 0.769000    0.9600
## 82            TG(50:2)_[LVL3]; 167    1.12e-02 0.771000    0.9600
## 83              PC(38:4)_[LVL2]; 9   -1.77e-02 0.771000    0.9600
## 84           SM(d33:1)_[LVL2]; 166    1.50e-02 0.783000    0.9600
## 85             PC(38:5)_[LVL2]; 24    1.53e-02 0.787000    0.9600
## 86           SM(d36:2)_[LVL2]; 160   -1.50e-02 0.790000    0.9600
## 87            SM(d38:1)_[LVL2]; 67   -1.18e-02 0.827000    0.9600
## 88            TG(54:3)_[LVL3]; 124   -8.94e-03 0.837000    0.9600
## 89            TG(51:1)_[LVL3]; 249    9.04e-03 0.844000    0.9600
## 90            TG(54:4)_[LVL3]; 129   -9.86e-03 0.845000    0.9600
## 91            TG(49:2)_[LVL3]; 231    9.80e-03 0.847000    0.9600
## 92             SM(42:2)_[LVL2]; 14   -1.00e-02 0.855000    0.9600
## 93            TG(54:6)_[LVL3]; 316    9.63e-03 0.856000    0.9600
## 94            TG(54:5)_[LVL3]; 240    9.20e-03 0.862000    0.9600
## 95  PC(16:0e/18:1(9Z))_[LVL1]; 134    9.64e-03 0.864000    0.9600
## 96            LPC(18:1)_[LVL2]; 34    1.02e-02 0.870000    0.9600
## 97            TG(49:1)_[LVL3]; 187    6.68e-03 0.897000    0.9680
## 98            PC(37:2)_[LVL2]; 350    7.09e-03 0.904000    0.9680
```

```
## 99           TG(46:1)_[LVL3]; 128    6.56e-03 0.904000    0.9680
## 100  TG(18:1/18:1/18:1)_[LVL2]; 15   -4.64e-03 0.917000    0.9680
## 101         SM(d41:2)_[LVL2]; 139   -4.88e-03 0.927000    0.9680
## 102           TG(49:3)_[LVL3]; 218    3.60e-03 0.940000    0.9680
## 103          SM(d40:2)_[LVL2]; 80    3.82e-03 0.940000    0.9680
## 104           TG(52:3)_[LVL3]; 101   -2.40e-03 0.953000    0.9710
## 105           TG(46:2)_[LVL3]; 248    1.81e-03 0.973000    0.9820
## 106           TG(52:4)_[LVL3]; 157   -6.66e-05 0.999000    0.9990
```

#### 7.6 Neuropathy Questionnaire (mnsineuropat)

##### 7.6.1 Crude Model

```
## [1] "Fitting models:"
## [1] "~ mnsineuropat"
## [1] ""
```

###### 7.6.1.1 Heatmap

```
## [1] "heatmap_lipidome_from_limma was created by Tommi Suvitaival"
## [1] ""
## [1] "2019-05-21"
```

```
## Warning: Removed 106 rows containing missing values (geom_point).
```

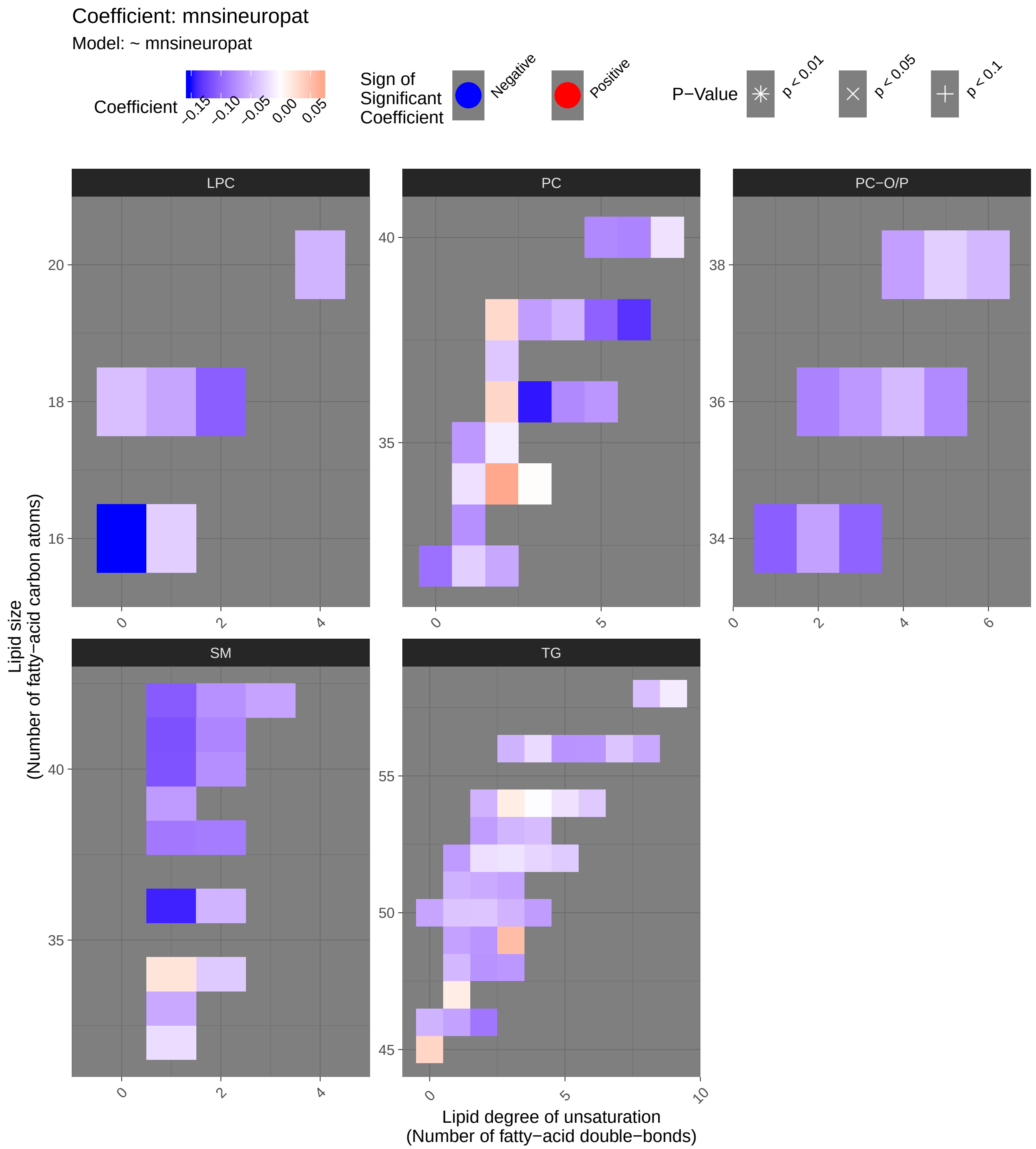

Coefficient: mnsineuropat
Model: ~ mnsineuropat
Coefficient
−0.15
−0.10
−0.05
0.00
0.05
Sign of Significant Coefficient
Negative
Positive
P−Value
p < 0.01
p < 0.05
p < 0.1
LPC
PC
PC−O/P
SM
TG
Lipid size (Number of fatty−acid carbon atoms)
Lipid degree of unsaturation (Number of fatty−acid double−bonds)

##### 7.6.1.2 Tables of Model Coefficients

```
## [1] ""
## [1] "Table: mnsineuropat"
## [1] "  (from model: "
## [1] "   ~ mnsineuropat)"
## [1] ""
##                                Name Coefficient P.Value adj.P.Val
## 1              LPC(16:0)_[LVL1]; 5    -0.15200  0.0124     0.682
## 2              PC(36:3)_[LVL2]; 10    -0.14700  0.0162     0.682
## 3             SM(d36:1)_[LVL2]; 55    -0.14300  0.0193     0.682
## 4               PC(38:6)_[LVL2]; 8    -0.13300  0.0287     0.723
## 5            SM(d41:1)_[LVL2]; 102    -0.11500  0.0601     0.723
## 6             SM(d40:1)_[LVL2]; 39    -0.11400  0.0624     0.723
## 7        SM(d18:1/24:0)_[LVL2]; 61    -0.10800  0.0752     0.723
## 8   PC(16:0e/18:1(9Z))_[LVL1]; 134    -0.10600  0.0819     0.723
## 9             LPC(18:2)_[LVL2]; 33    -0.10600  0.0821     0.723
## 10             PC(38:5)_[LVL2]; 24    -0.10300  0.0897     0.723
## 11          PC(O-34:3)_[LVL2]; 140    -0.10300  0.0904     0.723
## 12             PC(32:0)_[LVL2]; 96    -0.09370  0.1240     0.723
## 13            TG(46:2)_[LVL3]; 248    -0.09050  0.1380     0.723
## 14            SM(d38:1)_[LVL2]; 67    -0.08910  0.1440     0.723
## 15           SM(d38:2)_[LVL2]; 151    -0.08650  0.1560     0.723
## 16          PC(O-36:2)_[LVL2]; 312    -0.08200  0.1780     0.723
## 17             PC(40:6)_[LVL2]; 31    -0.08100  0.1840     0.723
## 18           SM(d41:2)_[LVL2]; 139    -0.08010  0.1890     0.723
## 19              PC(36:4)_[LVL2]; 1    -0.07830  0.1990     0.723
## 20             PC(40:5)_[LVL2]; 95    -0.07790  0.2010     0.723
## 21           PC(O-36:5)_[LVL2]; 92    -0.07730  0.2050     0.723
## 22            SM(d40:2)_[LVL2]; 80    -0.07390  0.2250     0.723
## 23            PC(33:1)_[LVL2]; 177    -0.07310  0.2300     0.723
## 24             SM(42:2)_[LVL2]; 14    -0.07250  0.2350     0.723
## 25  TG(18:1/12:0/18:1) or TG(18:2/    -0.07170  0.2400     0.723
## 26            TG(56:5)_[LVL2]; 230    -0.07090  0.2450     0.723
## 27            TG(49:2)_[LVL3]; 231    -0.06990  0.2520     0.723
## 28            TG(56:6)_[LVL3]; 275    -0.06950  0.2550     0.723
## 29             PC(36:5)_[LVL2]; 23    -0.06920  0.2560     0.723
## 30              PC(34:2)_[LVL2]; 4     0.06860  0.2610     0.723
## 31            TG(48:3)_[LVL3]; 384    -0.06840  0.2620     0.723
## 32            PC(35:1)_[LVL2]; 178    -0.06760  0.2670     0.723
## 33          PC(O-36:3)_[LVL2]; 268    -0.06710  0.2710     0.723
## 34  TG(18:2/18:2/18:2) or TG(18:3/    -0.06650  0.2750     0.723
## 35           SM(d39:1)_[LVL2]; 179    -0.06590  0.2800     0.723
## 36   TG(16:0/18:0/18:1)_[LVL2]; 51    -0.06530  0.2840     0.723
## 37  TG(14:0/18:2/18:2)_[LVL2]; 189    -0.06520  0.2850     0.723
## 38  TG(16:0/22:5/18:1) or TG(20:4/    -0.06510  0.2860     0.723
## 39             PC(38:3)_[LVL2]; 29    -0.06420  0.2920     0.723
## 40            TG(53:2)_[LVL2]; 234    -0.06400  0.2940     0.723
## 41          PC(O-38:4)_[LVL2]; 131    -0.06250  0.3050     0.723
## 42            TG(46:1)_[LVL3]; 128    -0.06180  0.3110     0.723
## 43          PC(O-34:2)_[LVL2]; 171    -0.06170  0.3120     0.723
## 44            TG(49:1)_[LVL3]; 187    -0.06150  0.3130     0.723
## 45   TG(18:2/22:5/16:0)_[LVL2]; 69    -0.06080  0.3180     0.723
## 46            TG(51:3)_[LVL3]; 198    -0.06070  0.3190     0.723
```

```
## 47       SM(d18:2/24:1)_[LVL2]; 40    -0.06030  0.3230     0.723
## 48            LPC(18:1)_[LVL2]; 34    -0.05910  0.3330     0.723
## 49            TG(50:0)_[LVL2]; 159    -0.05880  0.3340     0.723
## 50            PC(32:2)_[LVL2]; 204    -0.05730  0.3470     0.723
## 51           SM(d33:1)_[LVL2]; 166    -0.05680  0.3520     0.723
## 52  TG(16:0/18:2/22:6)_[LVL2]; 117    -0.05630  0.3560     0.723
## 53            TG(51:2)_[LVL2]; 123    -0.05560  0.3620     0.723
## 54            TG(49:3)_[LVL3]; 218     0.05280  0.3870     0.728
## 55            TG(51:1)_[LVL3]; 249    -0.05050  0.4080     0.728
## 56             TG(50:3)_[LVL2]; 47    -0.05030  0.4090     0.728
## 57             TG(54:2)_[LVL3]; 52    -0.05020  0.4100     0.728
## 58            TG(46:0)_[LVL3]; 168    -0.04990  0.4130     0.728
## 59            TG(56:3)_[LVL2]; 290    -0.04960  0.4150     0.728
## 60           LPC(20:4)_[LVL2]; 120    -0.04920  0.4200     0.728
## 61           SM(d36:2)_[LVL2]; 160    -0.04880  0.4230     0.728
## 62            TG(53:3)_[LVL3]; 239    -0.04850  0.4260     0.728
## 63              PC(38:4)_[LVL2]; 9    -0.04760  0.4350     0.731
## 64          PC(O-38:6)_[LVL2]; 236    -0.04640  0.4470     0.731
## 65   TG(14:0/16:0/18:1)_[LVL2]; 54    -0.04620  0.4480     0.731
## 66           PC(O-36:4)_[LVL2]; 71    -0.04500  0.4610     0.738
## 67   TG(14:0/18:1/18:1)_[LVL2]; 25    -0.04380  0.4730     0.738
## 68            TG(53:4)_[LVL3]; 314    -0.04370  0.4740     0.738
## 69  TG(16:0/18:2/18:3)_[LVL2]; 106    -0.04180  0.4930     0.741
## 70  TG(18:1/18:1/22:6)_[LVL2]; 147    -0.04160  0.4950     0.741
## 71            LPC(18:0)_[LVL1]; 22    -0.04150  0.4960     0.741
## 72            TG(56:7)_[LVL3]; 309    -0.03860  0.5270     0.765
## 73             TG(50:1)_[LVL3]; 19    -0.03830  0.5300     0.765
## 74            TG(50:2)_[LVL3]; 167    -0.03730  0.5410     0.765
## 75   TG(16:0/18:2/18:2)_[LVL2]; 27    -0.03680  0.5460     0.765
## 76            PC(37:2)_[LVL2]; 350    -0.03620  0.5520     0.765
## 77            TG(54:6)_[LVL3]; 316    -0.03500  0.5660     0.765
## 78  SM(d16:1/18:1) or SM(d18:2/16:    -0.03460  0.5710     0.765
## 79            TG(52:5)_[LVL3]; 286    -0.03380  0.5790     0.765
## 80             TG(45:0)_[LVL2]; 65     0.03340  0.5830     0.765
## 81           LPC(16:1)_[LVL2]; 258    -0.03180  0.6020     0.765
## 82              PC(36:2)_[LVL2]; 3     0.03170  0.6030     0.765
## 83             PC(32:1)_[LVL2]; 44    -0.03150  0.6050     0.765
## 84           PC(O-38:5)_[LVL2]; 76    -0.03140  0.6060     0.765
## 85            PC(38:2)_[LVL2]; 197     0.02990  0.6230     0.777
## 86            TG(52:4)_[LVL3]; 157    -0.02710  0.6570     0.810
## 87  TG(18:2/18:1/16:0)_[LVL2]; 500     0.02620  0.6680     0.814
## 88  TG(18:0/18:1/20:4)_[LVL2]; 141    -0.02460  0.6860     0.826
## 89            TG(56:4)_[LVL3]; 278    -0.02400  0.6940     0.826
## 90           SM(d32:1)_[LVL2]; 105    -0.02230  0.7140     0.834
## 91   TG(18:1/18:2/18:2)_[LVL2]; 57    -0.02200  0.7180     0.834
## 92            SM(d34:1)_[LVL2]; 26     0.02140  0.7250     0.834
## 93             TG(52:2)_[LVL3]; 97    -0.02080  0.7330     0.834
## 94              PC(34:1)_[LVL2]; 2    -0.01950  0.7490     0.834
## 95            TG(54:5)_[LVL3]; 240    -0.01910  0.7540     0.834
## 96            PC(40:7)_[LVL2]; 165    -0.01900  0.7560     0.834
## 97            TG(52:3)_[LVL3]; 101    -0.01770  0.7720     0.843
## 98            TG(54:3)_[LVL3]; 124     0.01470  0.8090     0.872
## 99            TG(47:1)_[LVL3]; 227     0.01430  0.8140     0.872
## 100           TG(58:9)_[LVL3]; 207    -0.01300  0.8310     0.880
```

```
## 101           PC(35:2)_[LVL2]; 143    -0.01210  0.8430     0.884
## 102  TG(18:1/18:1/18:1)_[LVL2]; 15     0.01030  0.8650     0.899
## 103   TG(18:1/18:1/16:0)_[LVL2]; 7     0.00903  0.8820     0.908
## 104  TG(18:2/18:1/18:1)_[LVL2]; 20    -0.00355  0.9540     0.972
## 105           TG(54:4)_[LVL3]; 129    -0.00224  0.9710     0.975
## 106           PC(34:3)_[LVL2]; 113     0.00194  0.9750     0.975
```

##### 7.6.2 Adjusted Model

```
## [1] "Fitting models:"
## [1] "~ mnsineuropat + Age + bmi + Blood_glucose + Duration_DM + Gender + Hba1c_baseline + log_Blood_
## [1] ""
```

###### 7.6.2.1 Heatmap

```
## [1] "heatmap_lipidome_from_limma was created by Tommi Suvitaival"
## [1] ""
## [1] "2019-05-21"
```

```
## Warning: Removed 106 rows containing missing values (geom_point).
```

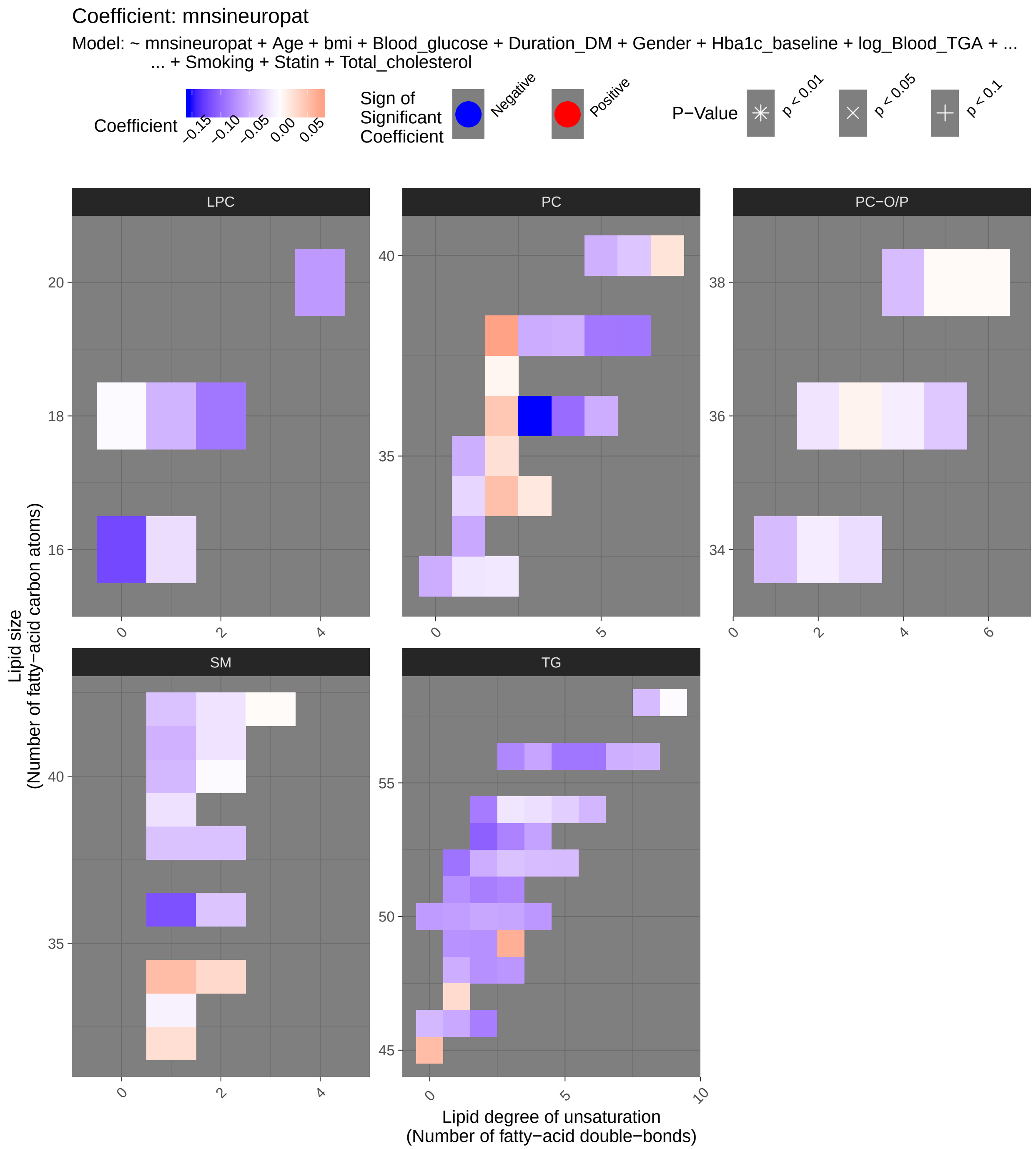

Coefficient: mnsineuropat
Model: ~ mnsineuropat + Age + bmi + Blood_glucose + Duration_DM + Gender + Hba1c_baseline + log_Blood_TGA + ...
... + Smoking + Statin + Total_cholesterol
Coefficient
−0.15
−0.10
−0.05
0.00
0.05
Sign of Significant Coefficient
Negative
Positive
P−Value
p < 0.01
p < 0.05
p < 0.1
LPC
PC
PC−O/P
SM
TG
Lipid size
(Number of fatty−acid carbon atoms)
Lipid degree of unsaturation
(Number of fatty−acid double−bonds)

###### 7.6.2.2 Tables of Model Coefficients

```
## [1] ""
## [1] "Table: mnsineuropat"
## [1] "  (from model: "
## [1] "   ~ mnsineuropat + Age + bmi + Blood_glucose + Duration_DM +"
## [1] "     Gender + Hba1c_baseline + log_Blood_TGA + Smoking + Statin +"
## [1] "     Total_cholesterol)"
## [1] ""
##                               Name Coefficient P.Value adj.P.Val
## 1              TG(53:2)_[LVL2]; 234    -0.10500 0.00460     0.292
## 2               PC(36:3)_[LVL2]; 10    -0.15400 0.00551     0.292
## 3              TG(51:2)_[LVL2]; 123    -0.08550 0.01910     0.434
## 4               TG(54:2)_[LVL3]; 52    -0.08770 0.01920     0.434
## 5              SM(d36:1)_[LVL2]; 55    -0.11600 0.02120     0.434
## 6    TG(16:0/18:0/18:1)_[LVL2]; 51    -0.09320 0.02700     0.434
## 7                LPC(16:0)_[LVL1]; 5    -0.12100 0.03090     0.434
## 8              TG(53:3)_[LVL3]; 239    -0.08280 0.03270     0.434
## 9              TG(56:5)_[LVL2]; 230    -0.09190 0.04110     0.469
## 10             TG(51:3)_[LVL3]; 198    -0.08010 0.04660     0.469
## 11  TG(16:0/22:5/18:1) or TG(20:4/    -0.08940 0.04880     0.469
## 12             TG(56:6)_[LVL3]; 275    -0.09210 0.05570     0.469
## 13               PC(38:6)_[LVL2]; 8    -0.09060 0.07650     0.469
## 14   TG(14:0/18:1/18:1)_[LVL2]; 25    -0.06160 0.07660     0.469
## 15              TG(50:3)_[LVL2]; 47    -0.05980 0.07980     0.469
## 16             TG(46:2)_[LVL3]; 248    -0.08620 0.08680     0.469
## 17              TG(50:1)_[LVL3]; 19    -0.06420 0.08910     0.469
## 18  TG(18:1/12:0/18:1) or TG(18:2/    -0.07350 0.09060     0.469
## 19             TG(51:1)_[LVL3]; 249    -0.07390 0.09160     0.469
## 20              PC(38:5)_[LVL2]; 24    -0.09000 0.09370     0.469
## 21             TG(56:3)_[LVL2]; 290    -0.07970 0.09590     0.469
## 22  TG(14:0/18:2/18:2)_[LVL2]; 189    -0.06870 0.09740     0.469
## 23               PC(36:4)_[LVL2]; 1    -0.09770 0.10500     0.484
## 24             TG(50:2)_[LVL3]; 167    -0.05780 0.11500     0.487
## 25              TG(52:2)_[LVL3]; 97    -0.05410 0.11900     0.487
## 26             LPC(18:2)_[LVL2]; 33    -0.09030 0.11900     0.487
## 27             TG(49:2)_[LVL3]; 231    -0.07370 0.12800     0.488
## 28             TG(48:3)_[LVL3]; 384    -0.06970 0.12900     0.488
## 29             TG(49:1)_[LVL3]; 187    -0.07240 0.14200     0.515
## 30              TG(45:0)_[LVL2]; 65     0.05300 0.14600     0.515
## 31             TG(49:3)_[LVL3]; 218     0.06350 0.16500     0.546
## 32  TG(18:2/18:2/18:2) or TG(18:3/    -0.06870 0.16700     0.546
## 33             TG(53:4)_[LVL3]; 314    -0.06160 0.17100     0.546
## 34             PC(38:2)_[LVL2]; 197     0.07380 0.17600     0.546
## 35             TG(56:4)_[LVL3]; 278    -0.06010 0.18000     0.546
## 36   TG(16:0/18:2/18:2)_[LVL2]; 27    -0.05110 0.19900     0.579
## 37             TG(50:0)_[LVL2]; 159    -0.06650 0.20200     0.579
## 38   TG(14:0/16:0/18:1)_[LVL2]; 54    -0.05380 0.23400     0.642
## 39            LPC(20:4)_[LVL2]; 120    -0.06730 0.24100     0.642
## 40  TG(18:0/18:1/20:4)_[LVL2]; 141    -0.06190 0.26200     0.642
## 41             TG(46:1)_[LVL3]; 128    -0.05730 0.26600     0.642
## 42   TG(18:2/22:5/16:0)_[LVL2]; 69    -0.05650 0.28100     0.642
## 43             PC(33:1)_[LVL2]; 177    -0.05720 0.28900     0.642
## 44              PC(36:5)_[LVL2]; 23    -0.05390 0.29900     0.642
```

```
## 45            SM(d41:1)_[LVL2]; 102    -0.05140 0.30300      0.642
## 46             TG(52:3)_[LVL3]; 101    -0.04040 0.30400      0.642
## 47              PC(40:5)_[LVL2]; 95    -0.05230 0.30500      0.642
## 48             TG(52:4)_[LVL3]; 157    -0.04330 0.30700      0.642
## 49              PC(38:3)_[LVL2]; 29    -0.05490 0.31000      0.642
## 50             SM(d40:1)_[LVL2]; 39    -0.04740 0.31100      0.642
## 51              PC(32:0)_[LVL2]; 96    -0.05410 0.31400      0.642
## 52             TG(56:7)_[LVL3]; 309    -0.05360 0.31700      0.642
## 53             PC(35:1)_[LVL2]; 178    -0.05310 0.32100      0.642
## 54  TG(16:0/18:2/18:3)_[LVL2]; 106    -0.04690 0.32800      0.643
## 55             TG(54:6)_[LVL3]; 316    -0.04820 0.34300      0.643
## 56             TG(52:5)_[LVL3]; 286    -0.04480 0.34500      0.643
## 57             SM(d34:1)_[LVL2]; 26     0.05380 0.34600      0.643
## 58               PC(38:4)_[LVL2]; 9    -0.05230 0.35500      0.643
## 59  TG(16:0/18:2/22:6)_[LVL2]; 117    -0.05030 0.35800      0.643
## 60             TG(46:0)_[LVL3]; 168    -0.04640 0.38400      0.663
## 61        SM(d18:1/24:0)_[LVL2]; 61    -0.04090 0.38700      0.663
## 62             LPC(18:1)_[LVL2]; 34    -0.04950 0.39600      0.663
## 63               PC(34:2)_[LVL2]; 4     0.05070 0.39900      0.663
## 64  PC(16:0e/18:1(9Z))_[LVL1]; 134    -0.04490 0.40100      0.663
## 65            SM(d38:1)_[LVL2]; 67     -0.04020 0.41500      0.677
## 66  TG(18:1/18:1/22:6)_[LVL2]; 147    -0.04420 0.43100      0.683
## 67            SM(d38:2)_[LVL2]; 151    -0.04050 0.43200      0.683
## 68           PC(O-38:4)_[LVL2]; 131    -0.04380 0.45500      0.692
## 69              PC(40:6)_[LVL2]; 31    -0.03760 0.45600      0.692
## 70            SM(d36:2)_[LVL2]; 160    -0.03800 0.45900      0.692
## 71               PC(36:2)_[LVL2]; 3     0.04450 0.46300      0.692
## 72  SM(d16:1/18:1) or SM(d18:2/16:      0.03080 0.48700      0.717
## 73   TG(18:1/18:2/18:2)_[LVL2]; 57    -0.03140 0.52600      0.753
## 74             TG(54:5)_[LVL3]; 240    -0.03210 0.53200      0.753
## 75            PC(O-36:5)_[LVL2]; 92    -0.03590 0.53300      0.753
## 76    TG(18:1/18:1/16:0)_[LVL2]; 7    -0.02780 0.57500      0.802
## 77            SM(d32:1)_[LVL2]; 105     0.02600 0.58900      0.811
## 78   TG(18:1/18:1/18:1)_[LVL2]; 15    -0.02200 0.60900      0.828
## 79             TG(47:1)_[LVL3]; 227     0.02870 0.62700      0.833
## 80   TG(18:2/18:1/18:1)_[LVL2]; 20    -0.02050 0.65600      0.833
## 81           PC(O-34:3)_[LVL2]; 140    -0.02240 0.66400      0.833
## 82               PC(34:1)_[LVL2]; 2    -0.02690 0.66500      0.833
## 83             TG(54:4)_[LVL3]; 129    -0.02090 0.66500      0.833
## 84             PC(35:2)_[LVL2]; 143     0.02460 0.66600      0.833
## 85             PC(40:7)_[LVL2]; 165     0.02210 0.66800      0.833
## 86             TG(54:3)_[LVL3]; 124    -0.01620 0.70300      0.850
## 87            LPC(16:1)_[LVL2]; 258    -0.02200 0.71100      0.850
## 88            SM(d39:1)_[LVL2]; 179    -0.01940 0.71300      0.850
## 89            SM(d41:2)_[LVL2]; 139    -0.01830 0.71400      0.850
## 90              SM(42:2)_[LVL2]; 14    -0.01790 0.72700      0.857
## 91             PC(34:3)_[LVL2]; 113     0.01820 0.74100      0.863
## 92           PC(O-36:2)_[LVL2]; 312    -0.01740 0.75000      0.864
## 93              PC(32:1)_[LVL2]; 44    -0.01640 0.77400      0.883
## 94             PC(32:2)_[LVL2]; 204    -0.01490 0.79000      0.891
## 95  TG(18:2/18:1/16:0)_[LVL2]; 500     0.01320 0.80700      0.900
## 96           PC(O-34:2)_[LVL2]; 171    -0.01170 0.83900      0.924
## 97           PC(O-36:3)_[LVL2]; 268     0.00939 0.85200      0.924
## 98            PC(O-36:4)_[LVL2]; 71    -0.01090 0.85400      0.924
```

```
## 99           SM(d33:1)_[LVL2]; 166    -0.00838 0.86900     0.931
## 100           PC(37:2)_[LVL2]; 350     0.00830 0.88000     0.933
## 101          PC(O-38:5)_[LVL2]; 76     0.00485 0.93500     0.958
## 102         PC(O-38:6)_[LVL2]; 236     0.00435 0.93600     0.958
## 103            SM(d40:2)_[LVL2]; 80    -0.00342 0.94200     0.958
## 104      SM(d18:2/24:1)_[LVL2]; 40     0.00343 0.94600     0.958
## 105           LPC(18:0)_[LVL1]; 22    -0.00369 0.94900     0.958
## 106            TG(58:9)_[LVL3]; 207    -0.00278 0.96200     0.962
```

##### 7.6.3 Fully-Adjusted Model

```
## [1] "Fitting models:"
## [1] "~ mnsineuropat + Age + bmi + Blood_glucose + Duration_DM + Gender + Hba1c_baseline + log_Blood_T
## [1] ""
```

###### 7.6.3.1 Heatmap

```
## [1] "heatmap_lipidome_from_limma was created by Tommi Suvitaival"
## [1] ""
## [1] "2019-05-21"
```

```
## Warning: Removed 106 rows containing missing values (geom_point).
```

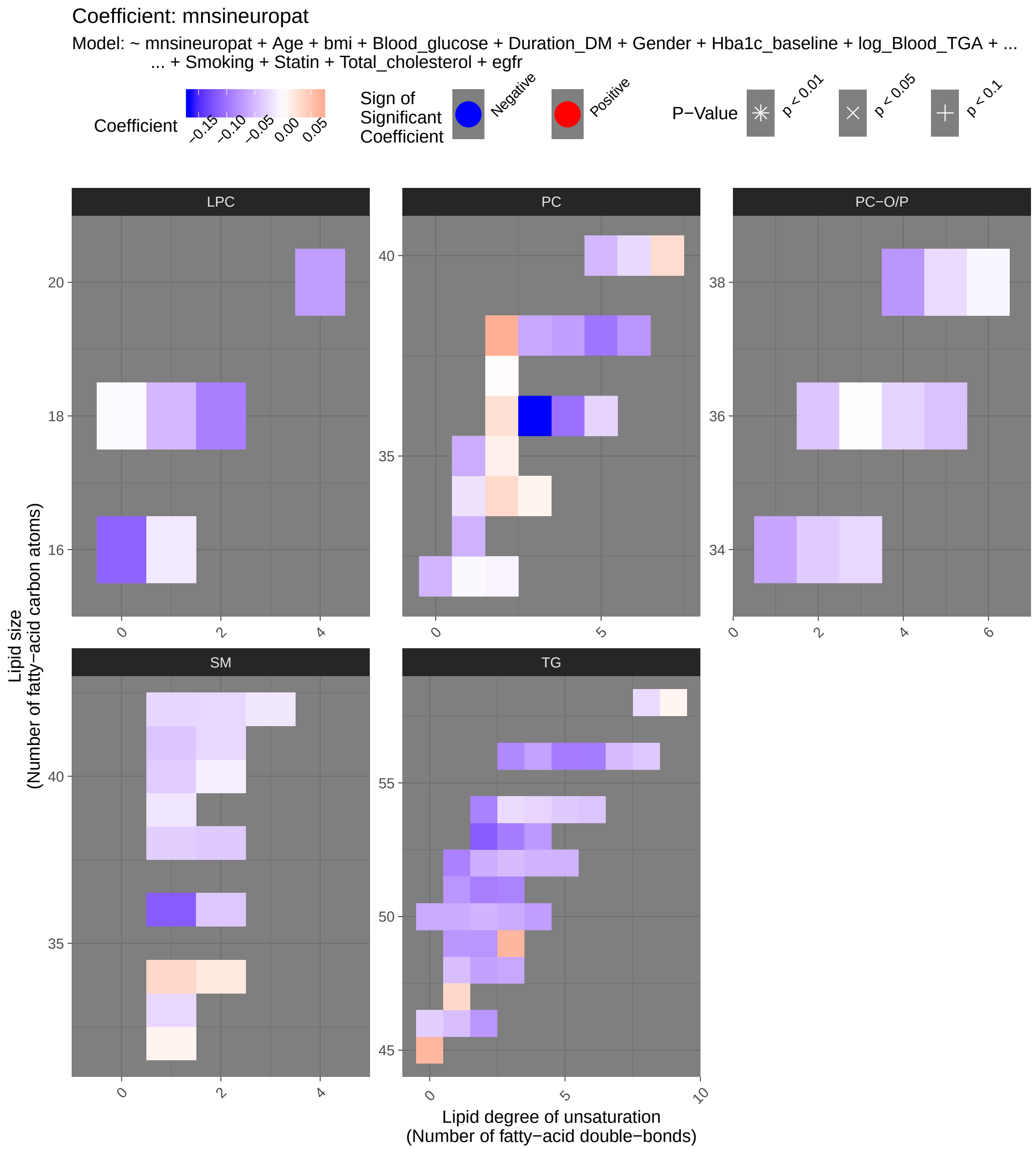
Coefficient: mnsineuropat
Model: ~ mnsineuropat + Age + bmi + Blood_glucose + Duration_DM + Gender + Hba1c_baseline + log_Blood_TGA + ...
... + Smoking + Statin + Total_cholesterol + egfr
Coefficient
−0.15
−0.10
−0.05
0.00
0.05
Sign of Significant Coefficient
Negative
Positive
P−Value
p < 0.01
p < 0.05
p < 0.1
LPC
PC
PC−O/P
SM
TG
Lipid size
(Number of fatty−acid carbon atoms)
Lipid degree of unsaturation
(Number of fatty−acid double−bonds)

###### 7.6.3.2 Tables of Model Coefficients

```
## [1] ""
## [1] "Table: mnsineuropat"
## [1] "  (from model: "
## [1] "   ~ mnsineuropat + Age + bmi + Blood_glucose + Duration_DM +"
## [1] "     Gender + Hba1c_baseline + log_Blood_TGA + Smoking + Statin +"
## [1] "     Total_cholesterol + egfr)"
## [1] ""
##                               Name Coefficient P.Value adj.P.Val
## 1            TG(53:2)_[LVL2]; 234    -0.11700 0.00170     0.160
## 2             PC(36:3)_[LVL2]; 10    -0.16600 0.00302     0.160
## 3            TG(51:2)_[LVL2]; 123    -0.09240 0.01210     0.361
## 4            TG(53:3)_[LVL3]; 239    -0.09420 0.01570     0.361
## 5             TG(54:2)_[LVL3]; 52    -0.09040 0.01700     0.361
## 6            SM(d36:1)_[LVL2]; 55    -0.11800 0.02060     0.364
## 7            TG(51:3)_[LVL3]; 198    -0.08880 0.02880     0.436
## 8   TG(16:0/18:0/18:1)_[LVL2]; 51    -0.09020 0.03420     0.447
## 9            TG(56:5)_[LVL2]; 230    -0.09440 0.03790     0.447
## 10  TG(16:0/22:5/18:1) or TG(20:4/    -0.09100 0.04750     0.454
## 11            LPC(16:0)_[LVL1]; 5    -0.11200 0.04760     0.454
## 12           TG(56:6)_[LVL3]; 275    -0.09490 0.05140     0.454
## 13            PC(38:5)_[LVL2]; 24    -0.09890 0.06810     0.489
## 14           TG(56:3)_[LVL2]; 290    -0.08530 0.07810     0.489
## 15            TG(50:3)_[LVL2]; 47    -0.06000 0.08210     0.489
## 16           TG(51:1)_[LVL3]; 249    -0.07560 0.08810     0.489
## 17            TG(45:0)_[LVL2]; 65     0.06230 0.08920     0.489
## 18             PC(36:4)_[LVL2]; 1    -0.10300 0.09180     0.489
## 19  TG(14:0/18:2/18:2)_[LVL2]; 189    -0.06990 0.09590     0.489
## 20            TG(52:2)_[LVL3]; 97    -0.05830 0.09630     0.489
## 21   TG(14:0/18:1/18:1)_[LVL2]; 25    -0.05840 0.09680     0.489
## 22           TG(53:4)_[LVL3]; 314    -0.07300 0.10700     0.493
## 23            TG(50:1)_[LVL3]; 19    -0.06000 0.11600     0.493
## 24           LPC(18:2)_[LVL2]; 33    -0.09190 0.11700     0.493
## 25           TG(49:2)_[LVL3]; 231    -0.07620 0.12000     0.493
## 26  TG(18:1/12:0/18:1) or TG(18:2/    -0.06700 0.12700     0.493
## 27           TG(49:1)_[LVL3]; 187    -0.07480 0.13300     0.493
## 28           TG(46:2)_[LVL3]; 248    -0.07550 0.13600     0.493
## 29             PC(38:6)_[LVL2]; 8    -0.07560 0.14000     0.493
## 30   TG(16:0/18:2/18:2)_[LVL2]; 27    -0.05860 0.14400     0.493
## 31           TG(56:4)_[LVL3]; 278    -0.06620 0.14500     0.493
## 32           TG(50:2)_[LVL3]; 167    -0.05350 0.14900     0.493
## 33           TG(49:3)_[LVL3]; 218     0.06350 0.17000     0.544
## 34           TG(48:3)_[LVL3]; 384    -0.06280 0.17500     0.547
## 35         PC(O-38:4)_[LVL2]; 131    -0.07480 0.19600     0.566
## 36           TG(52:4)_[LVL3]; 157    -0.05450 0.20000     0.566
## 37           PC(38:2)_[LVL2]; 197     0.06890 0.21100     0.566
## 38             PC(38:4)_[LVL2]; 9    -0.06930 0.22300     0.566
## 39  PC(16:0e/18:1(9Z))_[LVL1]; 134    -0.06490 0.22400     0.566
## 40          LPC(20:4)_[LVL2]; 120    -0.07000 0.22700     0.566
## 41           TG(52:3)_[LVL3]; 101    -0.04760 0.22900     0.566
## 42  TG(16:0/18:2/18:3)_[LVL2]; 106    -0.05770 0.23200     0.566
## 43  TG(18:2/18:2/18:2) or TG(18:3/    -0.05960 0.23400     0.566
## 44  TG(18:0/18:1/20:4)_[LVL2]; 141    -0.06490 0.24500     0.566
```

```
## 45              PC(38:3)_[LVL2]; 29    -0.06300 0.24800      0.566
## 46             TG(52:5)_[LVL3]; 286    -0.05500 0.25000      0.566
## 47             TG(50:0)_[LVL2]; 159    -0.06050 0.25100      0.566
## 48             PC(35:1)_[LVL2]; 178    -0.05930 0.27300      0.602
## 49             PC(33:1)_[LVL2]; 177    -0.05500 0.31300      0.664
## 50   TG(14:0/16:0/18:1)_[LVL2]; 54    -0.04600 0.31300      0.664
## 51              PC(40:5)_[LVL2]; 95    -0.05030 0.32900      0.683
## 52              PC(32:0)_[LVL2]; 96    -0.05240 0.33500      0.683
## 53   TG(18:2/22:5/16:0)_[LVL2]; 69    -0.04900 0.35500      0.710
## 54             TG(56:7)_[LVL3]; 309    -0.04820 0.37200      0.725
## 55             TG(46:1)_[LVL3]; 128    -0.04600 0.37600      0.725
## 56             LPC(18:1)_[LVL2]; 34    -0.05010 0.39600      0.739
## 57            SM(d41:1)_[LVL2]; 102    -0.04080 0.41700      0.739
## 58             TG(54:6)_[LVL3]; 316    -0.04170 0.41700      0.739
## 59            SM(d36:2)_[LVL2]; 160    -0.03970 0.44500      0.739
## 60           PC(O-36:2)_[LVL2]; 312    -0.04130 0.44700      0.739
## 61             SM(d40:1)_[LVL2]; 39    -0.03550 0.45000      0.739
## 62            PC(O-36:5)_[LVL2]; 92    -0.04350 0.45400      0.739
## 63            SM(d38:2)_[LVL2]; 151    -0.03850 0.46000      0.739
## 64             TG(54:5)_[LVL3]; 240    -0.03770 0.46700      0.739
## 65  TG(16:0/18:2/22:6)_[LVL2]; 117    -0.03920 0.47700      0.739
## 66   TG(18:1/18:2/18:2)_[LVL2]; 57    -0.03430 0.49300      0.739
## 67             SM(d38:1)_[LVL2]; 67    -0.03360 0.49900      0.739
## 68    TG(18:1/18:1/16:0)_[LVL2]; 7    -0.03310 0.51000      0.739
## 69             TG(46:0)_[LVL3]; 168    -0.03450 0.52000      0.739
## 70           PC(O-34:2)_[LVL2]; 171    -0.03650 0.52600      0.739
## 71       SM(d18:1/24:0)_[LVL2]; 61    -0.02880 0.54400      0.739
## 72             TG(54:4)_[LVL3]; 129    -0.02940 0.54600      0.739
## 73   TG(18:1/18:1/18:1)_[LVL2]; 15    -0.02600 0.55100      0.739
## 74             SM(d34:1)_[LVL2]; 26     0.03400 0.55100      0.739
## 75              PC(36:5)_[LVL2]; 23    -0.03060 0.55200      0.739
## 76             PC(40:7)_[LVL2]; 165     0.03060 0.55600      0.739
## 77   TG(18:2/18:1/18:1)_[LVL2]; 20    -0.02740 0.55600      0.739
## 78             TG(54:3)_[LVL3]; 124    -0.02450 0.56800      0.739
## 79             TG(47:1)_[LVL3]; 227     0.03360 0.57400      0.739
## 80            SM(d41:2)_[LVL2]; 139    -0.02740 0.58600      0.739
## 81              SM(42:2)_[LVL2]; 14    -0.02820 0.58700      0.739
## 82               PC(34:2)_[LVL2]; 4     0.03270 0.58800      0.739
## 83           PC(O-34:3)_[LVL2]; 140    -0.02810 0.58900      0.739
## 84            SM(d33:1)_[LVL2]; 166    -0.02740 0.59100      0.739
## 85            PC(O-36:4)_[LVL2]; 71    -0.03120 0.59800      0.739
## 86              PC(40:6)_[LVL2]; 31    -0.02660 0.60000      0.739
## 87  TG(18:1/18:1/22:6)_[LVL2]; 147    -0.02630 0.64000      0.780
## 88            PC(O-38:5)_[LVL2]; 76    -0.02530 0.66600      0.795
## 89               PC(36:2)_[LVL2]; 3     0.02610 0.66800      0.795
## 90  SM(d16:1/18:1) or SM(d18:2/16:     0.01860 0.67600      0.796
## 91       SM(d18:2/24:1)_[LVL2]; 40    -0.01680 0.74100      0.851
## 92            SM(d39:1)_[LVL2]; 179    -0.01730 0.74600      0.851
## 93               PC(34:1)_[LVL2]; 2    -0.02030 0.74700      0.851
## 94  TG(18:2/18:1/16:0)_[LVL2]; 500     0.01510 0.78200      0.874
## 95            LPC(16:1)_[LVL2]; 258    -0.01650 0.78400      0.874
## 96             SM(d40:2)_[LVL2]; 80    -0.01130 0.81400      0.898
## 97             PC(35:2)_[LVL2]; 143     0.01300 0.82100      0.898
## 98            SM(d32:1)_[LVL2]; 105     0.00968 0.84100      0.910
```

```
## 99             PC(34:3)_[LVL2]; 113     0.00958 0.86300     0.924
## 100            PC(32:2)_[LVL2]; 204    -0.00833 0.88300     0.935
## 101            TG(58:9)_[LVL3]; 207     0.00809 0.89000     0.935
## 102          PC(O-38:6)_[LVL2]; 236    -0.00639 0.90700     0.942
## 103             PC(32:1)_[LVL2]; 44    -0.00502 0.93100     0.958
## 104            PC(37:2)_[LVL2]; 350     0.00262 0.96200     0.966
## 105            LPC(18:0)_[LVL1]; 22    -0.00254 0.96500     0.966
## 106          PC(O-36:3)_[LVL2]; 268    -0.00213 0.96600     0.966
```

### 8 Appendix

```
## R version 3.6.2 (2019-12-12)
## Platform: x86_64-w64-mingw32/x64 (64-bit)
## Running under: Windows 10 x64 (build 17763)
##
## Matrix products: default
##
## locale:
## [1] LC_COLLATE=English_United States.1252
## [2] LC_CTYPE=English_United States.1252
## [3] LC_MONETARY=English_United States.1252
## [4] LC_NUMERIC=C
## [5] LC_TIME=English_United States.1252
##
## attached base packages:
## [1] stats     graphics  grDevices utils     datasets  methods   base
##
## loaded via a namespace (and not attached):
##  [1] Rcpp_1.0.3       knitr_1.27       magrittr_1.5     hms_0.5.3
##  [5] munsell_0.5.0    tidyselect_1.0.0 colorspace_1.4-1 R6_2.4.1
##  [9] rlang_0.4.6      stringr_1.4.0    dplyr_0.8.3      tools_3.6.2
## [13] grid_3.6.2       gtable_0.3.0     xfun_0.12        htmltools_0.4.0
## [17] ellipsis_0.3.0   lazyeval_0.2.2   yaml_2.2.0       digest_0.6.23
## [21] assertthat_0.2.1 tibble_3.0.1     lifecycle_0.2.0  crayon_1.3.4
## [25] farver_2.0.3     ggplot2_3.2.1    purrr_0.3.3      readr_1.3.1
## [29] tidyr_1.0.0      vctrs_0.2.4      glue_1.3.1       evaluate_0.14
## [33] haven_2.2.0      rmarkdown_2.1    labeling_0.3     limma_3.42.0
## [37] stringi_1.4.4    compiler_3.6.2   pillar_1.4.3     scales_1.1.0
## [41] forcats_0.4.0    pkgconfig_2.0.3
```
