## Supplementary material for "Cardiovascular Autonomic Neuropathy in Type 1 Diabetes is Associated with Several Metabolic Pathways – New Risk Markers on the Horizon": Metabolomics_supplementary_data

### 0033_PROFIL_2017 Neuropathy Metabolomics

Tommi Suvitaival,, Steno Diabetes Center Copenhagen

October 30, 2020

**6 Appendix** **112**

### 1 Settings

### 2 Filter

### 3 CAN Stat

#### 3.1 Crude Model

```
## [1] "Fitting models:"
## [1] "~ CAN_stat"
## [1] ""
```

##### 3.1.1 Tables of Model Coefficients

```
## [1] ""
## [1] "Table: CAN_stat"
## [1] "  (from model: "
## [1] "    ~ CAN_stat)"
## [1] ""
##                               Name Coefficient  P.Value adj.P.Val
## 1  2,4-Dihydroxybutanoic acid; 28     0.34900 2.93e-08  2.20e-06
## 2  3,4-Dihydroxybutanoic acid; 27     0.33300 1.21e-07  4.53e-06
## 3                 Creatinine; 50     0.32000 3.86e-07  9.66e-06
## 4            Citric acid, 4TMS; 6     0.28100 8.11e-06  1.52e-04
## 5               Ribonic acid; 72     0.27200 1.53e-05  2.29e-04
## 6            Myo inositol 6TMS; 1     0.23200 2.34e-04  2.92e-03
## 7          Benzeneacetic acid; 47     0.21200 7.70e-04  8.25e-03
## 8  4-Hydroxybenzeneacetic acid; 4     0.20100 1.42e-03  1.33e-02
## 9          Glyceryl-glycoside; 59     0.19300 2.16e-03  1.71e-02
## 10       4-Deoxytetronic acid; 32     0.19200 2.28e-03  1.71e-02
## 11      Eicosapentaenoic acid; 55    -0.17800 4.81e-03  3.28e-02
## 12                  Glycerol; 58     0.17200 6.44e-03  4.02e-02
## 13       Docosahexaenoic acid; 53    -0.16600 8.48e-03  4.89e-02
## 14       4-Deoxytetronic acid; 33     0.16200 1.01e-02  4.92e-02
## 15           Methionine, 2TMS; 16    -0.16200 1.03e-02  4.92e-02
## 16              Octanoic acid; 68    -0.16100 1.05e-02  4.92e-02
## 17         Succinic acid, 2TMS; 7     0.15000 1.70e-02  7.13e-02
## 18                   Ribitol; 70     0.14900 1.83e-02  7.13e-02
## 19                   Ribitol; 71     0.14800 1.86e-02  7.13e-02
## 20     4-Hydroxybutanoic acid; 43     0.14800 1.90e-02  7.13e-02
## 21           Isoleucine, 2TMS; 18    -0.14600 2.02e-02  7.23e-02
## 22               Valine, 2TMS; 20    -0.13900 2.76e-02  9.42e-02
## 23               Serine, 3TMS; 14    -0.13300 3.53e-02  1.14e-01
## 24             Hydroxyproline; 64     0.13200 3.66e-02  1.14e-01
## 25 4-Hydroxyphenyllactic acid; 44     0.12500 4.65e-02  1.40e-01
## 26             Arachidic acid; 46    -0.11800 5.99e-02  1.73e-01
## 27          Fumaric acid, 2TMS; 9     0.11300 7.15e-02  1.99e-01
## 28              Glycine, 3TMS; 17     0.11200 7.56e-02  2.03e-01
## 29          Nonadecanoic acid; 66    -0.11000 8.09e-02  2.09e-01
## 30                1-Dodecanol; 36     0.10800 8.49e-02  2.12e-01
## 31              Leucine, 2TMS; 19    -0.10600 9.36e-02  2.26e-01
## 32              Decanoic acid; 52    -0.10300 1.01e-01  2.29e-01
## 33            Arabinopyranose; 51     0.10300 1.02e-01  2.29e-01
## 34        3-Indoleacetic acid; 40     0.10200 1.04e-01  2.29e-01
## 35           Malic acid, 3TMS; 11     0.09980 1.13e-01  2.42e-01
## 36           Tridecanoic acid; 74     0.09850 1.18e-01  2.45e-01
## 37              Nonanoic acid; 67     0.08930 1.56e-01  3.16e-01
## 38 2-Hydroxybutyric acid, 2TMS; 2    -0.08410 1.82e-01  3.53e-01
## 39             L-5-Oxoproline; 63     0.08380 1.83e-01  3.53e-01
## 40           Stearic acid, TMS; 2    -0.07590 2.28e-01  4.25e-01
## 41          Pyroglutamic acid; 69     0.07480 2.35e-01  4.25e-01
## 42              Glyceric acid; 30    -0.07430 2.38e-01  4.25e-01
## 43             Tartronic acid; 73    -0.06920 2.71e-01  4.73e-01
## 44                  Tyrosine; 75    -0.06150 3.29e-01  5.38e-01
## 45              Alanine, 2TMS; 25     0.06150 3.29e-01  5.38e-01
## 46           Threonine, 3TMS; 12    -0.06130 3.30e-01  5.38e-01
```

```
## 47                 Glycerol; 57     0.05960 3.44e-01  5.44e-01
## 48              Lactic acid; 29     0.05860 3.52e-01  5.44e-01
## 49           1-Monopalmitin; 37     0.05820 3.55e-01  5.44e-01
## 50     Arachidonic acid, TMS; 24    0.05520 3.81e-01  5.63e-01
## 51          1,3-Propanediol; 34    -0.05500 3.83e-01  5.63e-01
## 52    3-Indolepropionic acid; 41   -0.05100 4.18e-01  6.03e-01
## 53         Glutamic acid, 3TMS; 8   0.04750 4.51e-01  6.38e-01
## 54 alpha-ketoglutaric acid, TMS M   0.04480 4.77e-01  6.63e-01
## 55              Pyruvic acid; 31    0.04120 5.13e-01  6.99e-01
## 56         11-Eicosenoic acid; 35  -0.04000 5.26e-01  7.01e-01
## 57          Palmitic acid, TMS; 5  -0.03860 5.40e-01  7.01e-01
## 58        Heptadecanoic acid; 61   -0.03840 5.42e-01  7.01e-01
## 59             Hydroxylamine; 62   -0.03520 5.76e-01  7.33e-01
## 60           Dodecanoic acid; 54   -0.03020 6.32e-01  7.79e-01
## 61          alpha-Tocopherol; 26   -0.03000 6.34e-01  7.79e-01
## 62       2-Palmitoylglycerol; 39   -0.02720 6.65e-01  8.02e-01
## 63             Proline, 2TMS; 21    0.02650 6.74e-01  8.02e-01
## 64       Phenylalanine, 2TMS; 13   -0.02410 7.02e-01  8.23e-01
## 65 3-Hydroxybutyric acid, 2TMS; 1  -0.02300 7.15e-01  8.24e-01
## 66          Cholesterol, TMS; 23   -0.01960 7.55e-01  8.55e-01
## 67               Bisphenol A; 48    0.01890 7.64e-01  8.55e-01
## 68          Aminomalonic acid; 45  -0.01320 8.34e-01  9.06e-01
## 69               Campesterol; 49    0.01160 8.54e-01  9.06e-01
## 70              Ethanolamine; 56   -0.01040 8.68e-01  9.06e-01
## 71          Myristoleic acid; 65   -0.01030 8.70e-01  9.06e-01
## 72        Heptadecanoic acid; 60   -0.01030 8.70e-01  9.06e-01
## 73            Oleic acid, TMS; 3    0.00788 9.00e-01  9.17e-01
## 74  2-hydroxy Isovaleric acid; 38   0.00753 9.05e-01  9.17e-01
## 75          Linoleic acid, TMS; 4   0.00255 9.68e-01  9.68e-01
```

##### 3.1.2 Forest Plot of Model Coefficients

```
## Warning: Ignoring unknown aesthetics: x
```

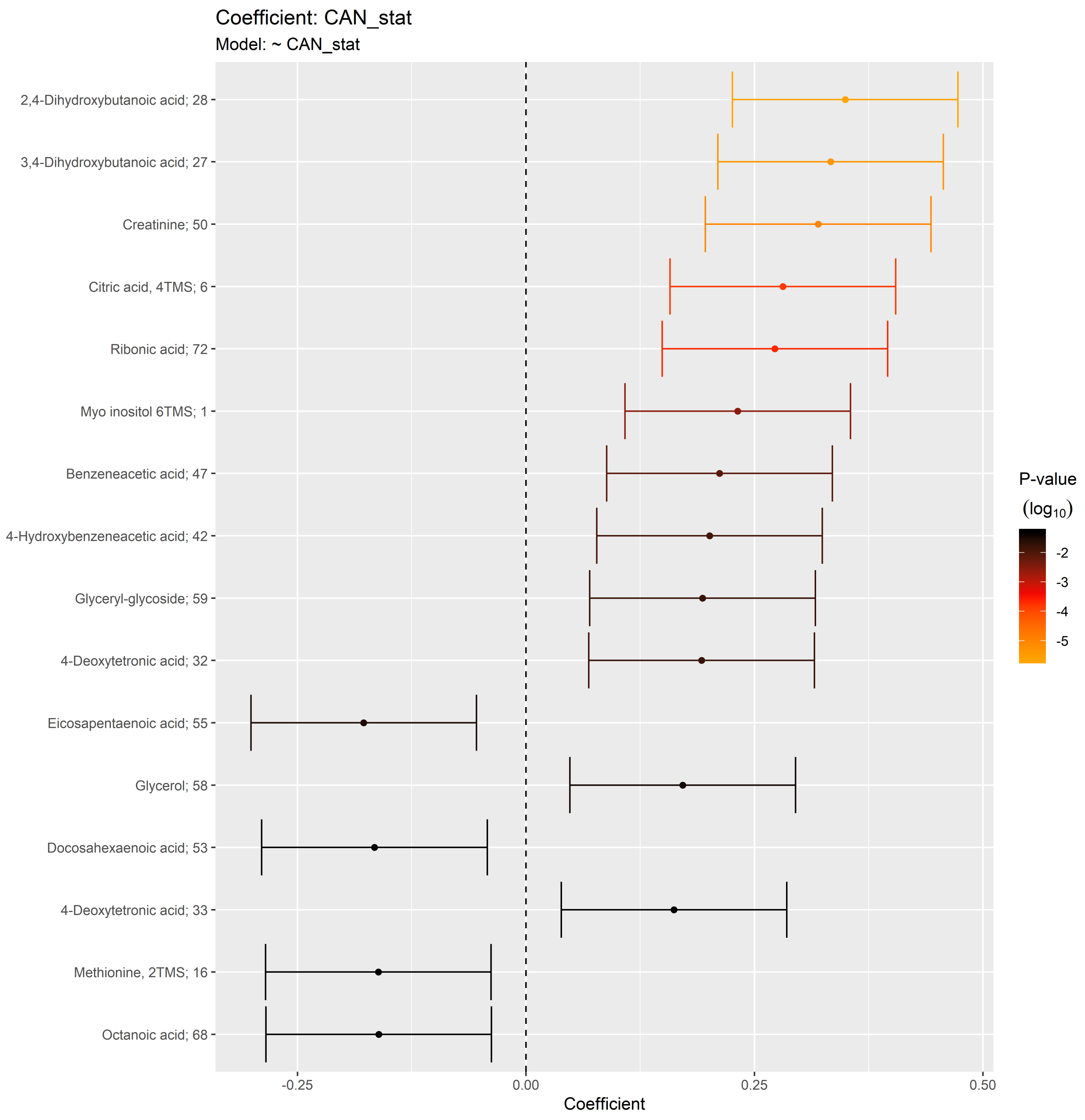

#### 3.2 Adjusted Model

```
## [1] "Fitting models:"
## [1] "~ CAN_stat + Age + bmi + Blood_glucose + Duration_DM + Gender + Hba1c_baseline + log_Blood_TGA +
## [1] ""
```

##### 3.2.1 Tables of Model Coefficients

```
## [1] ""
## [1] "Table: CAN_stat"
## [1] "  (from model: "
## [1] "   ~ CAN_stat + Age + bmi + Blood_glucose + Duration_DM +"
## [1] "     Gender + Hba1c_baseline + log_Blood_TGA + Smoking + Statin +"
## [1] "     Total_cholesterol)"
## [1] ""
##                               Name Coefficient  P.Value adj.P.Val
## 1            Citric acid, 4TMS; 6    0.313000 1.82e-06  0.000101
## 2                  Creatinine; 50    0.307000 2.70e-06  0.000101
## 3  2,4-Dihydroxybutanoic acid; 28    0.297000 5.58e-06  0.000140
## 4  3,4-Dihydroxybutanoic acid; 27    0.261000 6.64e-05  0.001240
## 5          Benzeneacetic acid; 47    0.255000 9.84e-05  0.001480
## 6                Ribonic acid; 72    0.230000 4.46e-04  0.005570
## 7            Isoleucine, 2TMS; 18   -0.197000 2.62e-03  0.027600
## 8        4-Deoxytetronic acid; 32    0.195000 2.95e-03  0.027600
## 9            Myo inositol 6TMS; 1    0.189000 3.84e-03  0.032000
## 10    4-Hydroxybutanoic acid; 43    0.184000 5.00e-03  0.037500
## 11                 Glycerol; 58    0.177000 6.79e-03  0.046300
## 12        Succinic acid, 2TMS; 7    0.175000 7.59e-03  0.047400
## 13 4-Hydroxybenzeneacetic acid; 4    0.168000 1.05e-02  0.060500
## 14          Tridecanoic acid; 74    0.158000 1.61e-02  0.081800
## 15              Valine, 2TMS; 20   -0.157000 1.64e-02  0.081800
## 16             Leucine, 2TMS; 19   -0.147000 2.50e-02  0.109000
## 17             Glycine, 3TMS; 17    0.146000 2.58e-02  0.109000
## 18        Glyceryl-glycoside; 59    0.146000 2.62e-02  0.109000
## 19          Methionine, 2TMS; 16   -0.142000 3.02e-02  0.116000
## 20      Eicosapentaenoic acid; 55   -0.141000 3.10e-02  0.116000
## 21              Serine, 3TMS; 14   -0.137000 3.68e-02  0.128000
## 22             Nonanoic acid; 67    0.136000 3.78e-02  0.128000
## 23             Octanoic acid; 68   -0.135000 3.92e-02  0.128000
## 24            Hydroxyproline; 64    0.132000 4.42e-02  0.138000
## 25            L-5-Oxoproline; 63    0.127000 5.31e-02  0.159000
## 26 2-Hydroxybutyric acid, 2TMS; 2   -0.124000 5.86e-02  0.169000
## 27                  Ribitol; 70    0.121000 6.46e-02  0.176000
## 28 4-Hydroxyphenyllactic acid; 44    0.121000 6.56e-02  0.176000
## 29             Hydroxylamine; 62   -0.119000 6.95e-02  0.180000
## 30       Docosahexaenoic acid; 53   -0.117000 7.30e-02  0.183000
## 31                1-Dodecanol; 36    0.113000 8.52e-02  0.206000
## 32       4-Deoxytetronic acid; 33    0.111000 9.03e-02  0.212000
## 33           Malic acid, 3TMS; 11    0.110000 9.43e-02  0.214000
## 34            Arachidic acid; 46   -0.108000 9.79e-02  0.216000
## 35          Fumaric acid, 2TMS; 9    0.092900 1.56e-01  0.334000
## 36         3-Indoleacetic acid; 40    0.091100 1.64e-01  0.342000
## 37             1-Monopalmitin; 37    0.079400 2.25e-01  0.449000
## 38          Pyroglutamic acid; 69    0.078100 2.33e-01  0.449000
## 39          Nonadecanoic acid; 66   -0.077200 2.38e-01  0.449000
## 40                  Ribitol; 71    0.077100 2.39e-01  0.449000
## 41          Aminomalonic acid; 45    0.068600 2.95e-01  0.540000
## 42           Stearic acid, TMS; 2   -0.067000 3.06e-01  0.547000
## 43              Decanoic acid; 52   -0.051400 4.32e-01  0.754000
## 44 3-Hydroxybutyric acid, 2TMS; 1   -0.049200 4.52e-01  0.770000
```

```
## 45                 Tyrosine; 75  -0.047600 4.68e-01  0.776000
## 46         Cholesterol, TMS; 23   0.046700 4.76e-01  0.776000
## 47        Palmitic acid, TMS; 5  -0.045000 4.92e-01  0.786000
## 48          1,3-Propanediol; 34  -0.041100 5.30e-01  0.818000
## 49     Arachidonic acid, TMS; 24  0.040700 5.35e-01  0.818000
## 50              Campesterol; 49  -0.038300 5.59e-01  0.835000
## 51          Arabinopyranose; 51   0.036100 5.82e-01  0.835000
## 52          Threonine, 3TMS; 12  -0.036100 5.82e-01  0.835000
## 53      2-Palmitoylglycerol; 39  -0.035300 5.90e-01  0.835000
## 54            Alanine, 2TMS; 25   0.031300 6.33e-01  0.839000
## 55              Bisphenol A; 48   0.028900 6.59e-01  0.839000
## 56       Heptadecanoic acid; 60   0.028700 6.61e-01  0.839000
## 57              Lactic acid; 29   0.028700 6.61e-01  0.839000
## 58             Pyruvic acid; 31   0.027800 6.71e-01  0.839000
## 59         alpha-Tocopherol; 26  -0.026500 6.86e-01  0.839000
## 60                 Glycerol; 57   0.026400 6.87e-01  0.839000
## 61           Tartronic acid; 73  -0.026000 6.92e-01  0.839000
## 62       11-Eicosenoic acid; 35  -0.025800 6.94e-01  0.839000
## 63          Dodecanoic acid; 54  -0.023300 7.22e-01  0.845000
## 64             Ethanolamine; 56  -0.022200 7.35e-01  0.845000
## 65            Glyceric acid; 30  -0.022200 7.35e-01  0.845000
## 66       Heptadecanoic acid; 61   0.021400 7.43e-01  0.845000
## 67 alpha-ketoglutaric acid, TMS M   0.020500 7.54e-01  0.845000
## 68  2-hydroxy Isovaleric acid; 38   0.019400 7.67e-01  0.846000
## 69       Glutamic acid, 3TMS; 8   0.017800 7.86e-01  0.854000
## 70        Linoleic acid, TMS; 4   0.009920 8.80e-01  0.939000
## 71           Oleic acid, TMS; 3   0.008350 8.99e-01  0.939000
## 72         Myristoleic acid; 65   0.008090 9.02e-01  0.939000
## 73            Proline, 2TMS; 21   0.006730 9.18e-01  0.943000
## 74    3-Indolepropionic acid; 41   0.002080 9.75e-01  0.988000
## 75      Phenylalanine, 2TMS; 13  -0.000666 9.92e-01  0.992000
```

##### 3.2.2 Forest Plot of Model Coefficients

```
## Warning: Ignoring unknown aesthetics: x

## Warning: Ignoring unknown aesthetics: x

## Warning: Ignoring unknown aesthetics: x

## Warning: Ignoring unknown aesthetics: x

## Warning: Ignoring unknown aesthetics: x

## Warning: Ignoring unknown aesthetics: x

## Warning: Ignoring unknown aesthetics: x
```

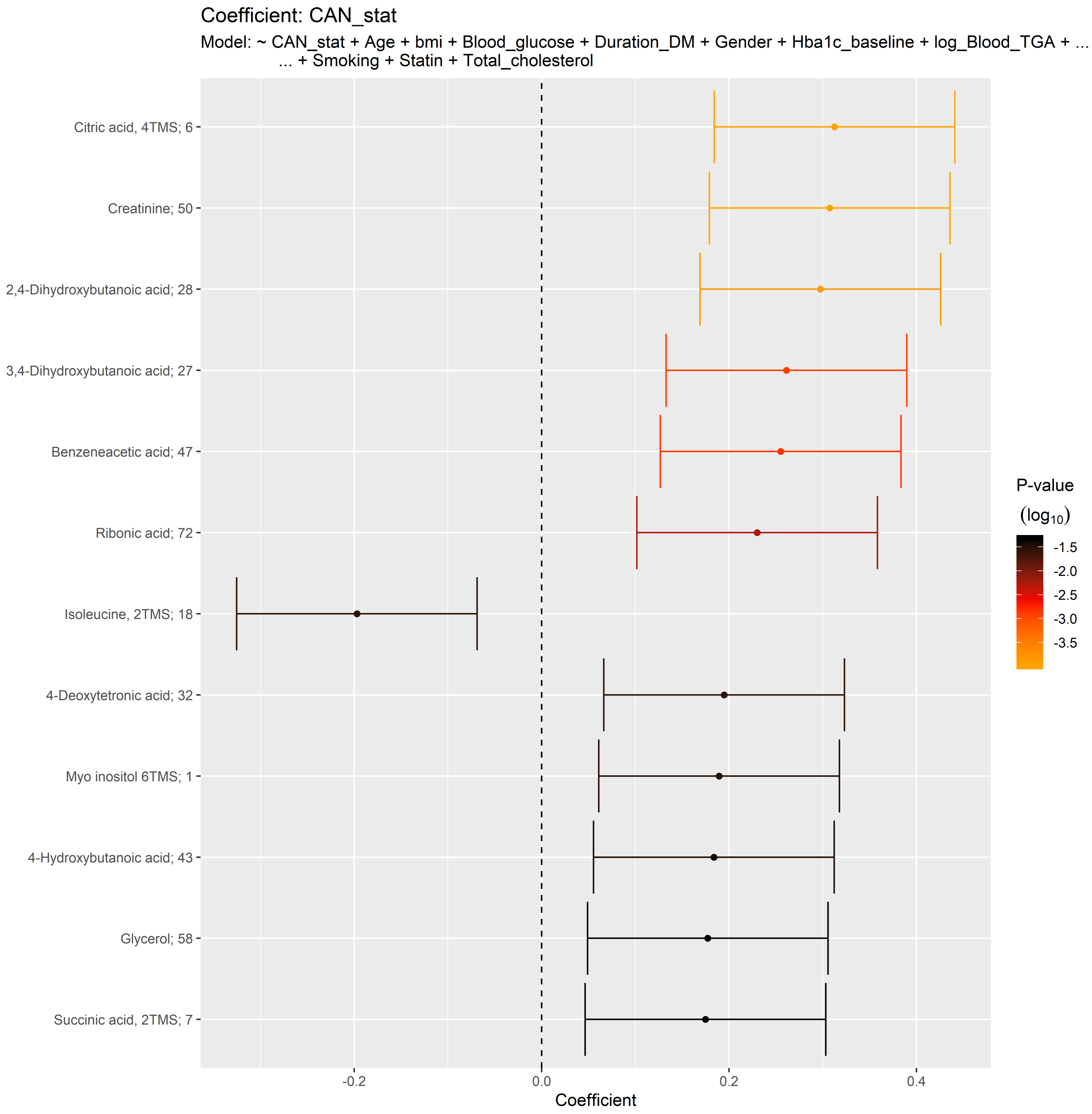
Coefficient: CAN_stat
Model: ~ CAN_stat + Age + bmi + Blood_glucose + Duration_DM + Gender + Hba1c_baseline + log_Blood_TGA + ...
... + Smoking + Statin + Total_cholesterol
Citric acid, 4TMS; 6
Creatinine; 50
2,4-Dihydroxybutanoic acid; 28
3,4-Dihydroxybutanoic acid; 27
Benzeneacetic acid; 47
Ribonic acid; 72
Isoleucine, 2TMS; 18
4-Deoxytetronic acid; 32
Myo inositol 6TMS; 1
4-Hydroxybutanoic acid; 43
Glycerol; 58
Succinic acid, 2TMS; 7
-0.2
0.0
0.2
0.4
Coefficient
P-value
(log10)
-1.5
-2.0
-2.5
-3.0
-3.5

##### 3.2.3 Bipartite Network of Model Coefficients

```
## [1] "bipartite_network_from_limma was created by Tommi Suvitaival"
## [1] ""
## [1] "2019-05-06"

## Warning in if (drop.variables != "none") {: the condition has length > 1 and
## only the first element will be used

## Warning: Removed 4 rows containing missing values (geom_segment).

## Warning: Removed 1 rows containing missing values (geom_text).
```

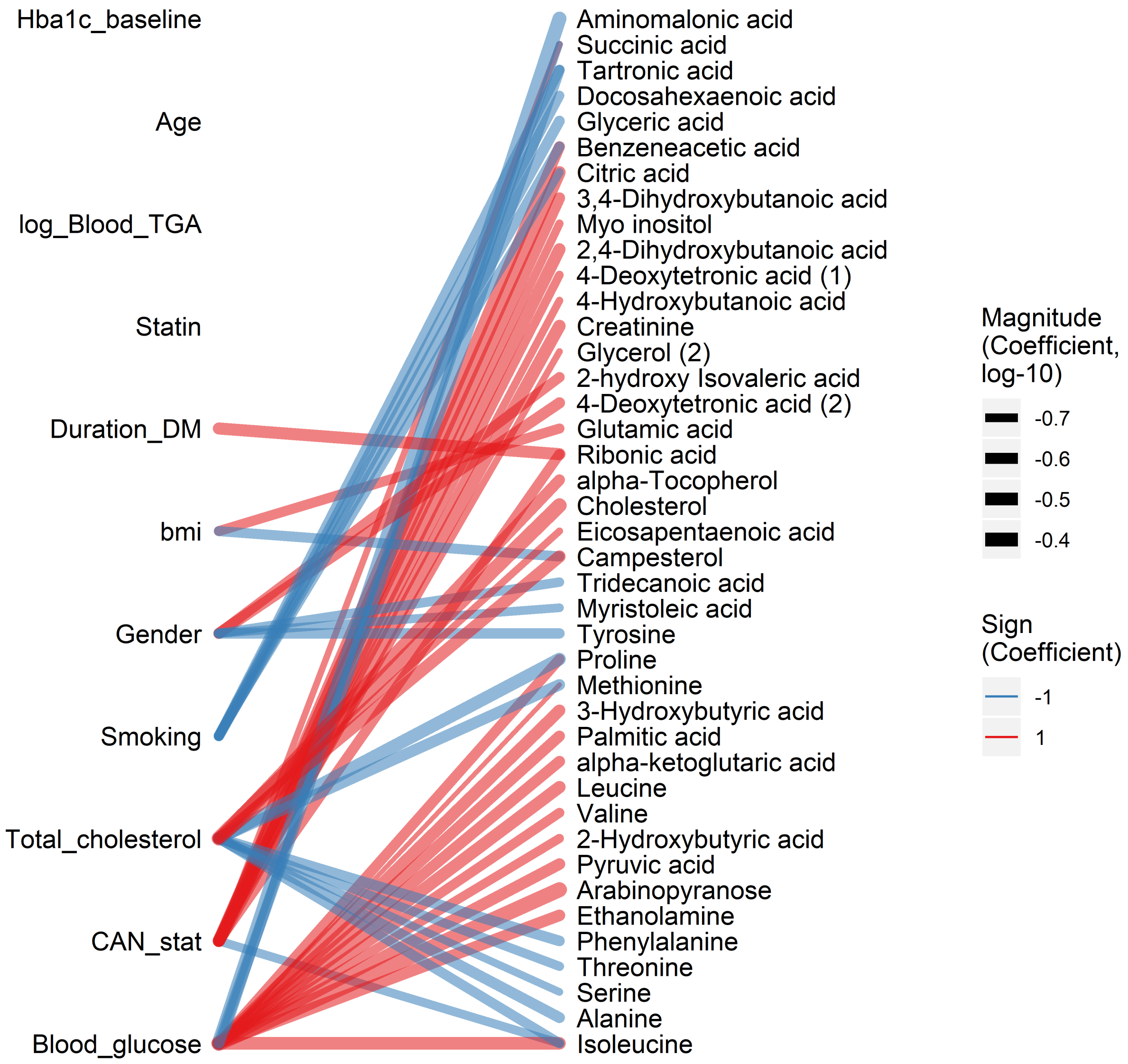

```
## [1] "bipartite_network_from_limma was created by Tommi Suvitaival"
## [1] ""
## [1] "2019-05-06"

## Warning in if (drop.variables != "none") {: the condition has length > 1 and
## only the first element will be used

## Warning: Removed 4 rows containing missing values (geom_segment).

## Warning: Removed 1 rows containing missing values (geom_text).
```

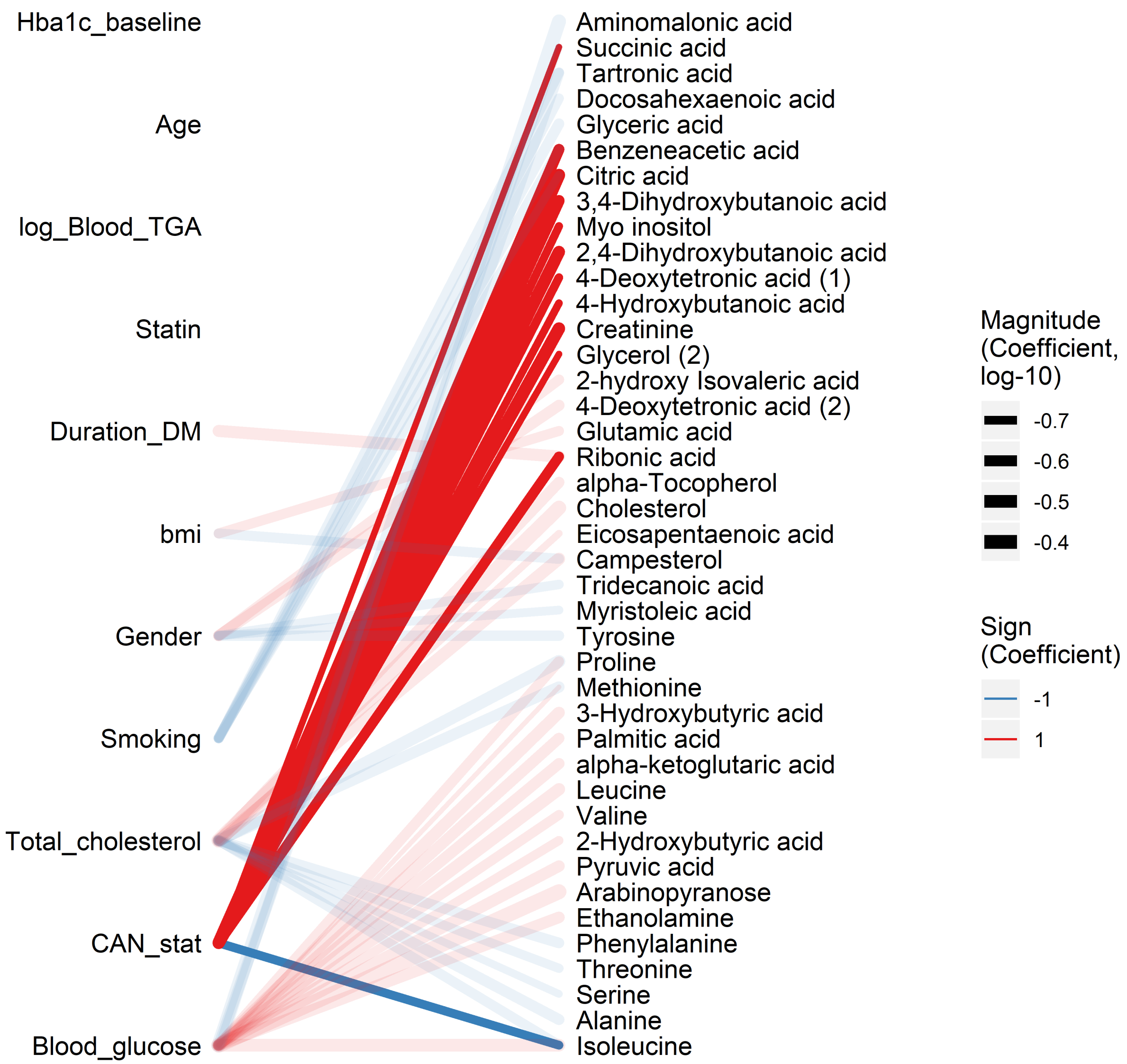

##### 3.3 Fully Adjusted Model

```
## [1] "Fitting models:"
## [1] "~ CAN_stat + Age + bmi + Blood_glucose + Duration_DM + Gender + Hba1c_baseline + log_Blood_TGA +
## [1] ""
```

##### 3.3.1 Tables of Model Coefficients

```
## [1] ""
## [1] "Table: CAN_stat"
## [1] "  (from model: "
## [1] "   ~ CAN_stat + Age + bmi + Blood_glucose + Duration_DM +"
## [1] "     Gender + Hba1c_baseline + log_Blood_TGA + Smoking + Statin +"
## [1] "     Total_cholesterol + egfr)"
## [1] ""
##                               Name Coefficient  P.Value adj.P.Val
## 1            Citric acid, 4TMS; 6     0.25800 0.000137    0.0103
## 2          Benzeneacetic acid; 47     0.23900 0.000425    0.0159
## 3                  Creatinine; 50     0.20900 0.002030    0.0508
## 4      4-Hydroxybutanoic acid; 43     0.18500 0.006790    0.1040
## 5          Succinic acid, 2TMS; 7     0.18400 0.006960    0.1040
## 6  2,4-Dihydroxybutanoic acid; 28     0.16400 0.014500    0.1820
## 7             Tridecanoic acid; 74     0.15900 0.019800    0.1990
## 8                Hydroxylamine; 62    -0.15700 0.021200    0.1990
## 9               L-5-Oxoproline; 63     0.14800 0.030400    0.2520
## 10 3,4-Dihydroxybutanoic acid; 27     0.14300 0.033600    0.2520
## 11                    Glycerol; 58     0.14000 0.039300    0.2680
## 12               Nonanoic acid; 67     0.13200 0.052700    0.3300
## 13       Eicosapentaenoic acid; 55    -0.12400 0.067500    0.3900
## 14            Isoleucine, 2TMS; 18    -0.11900 0.078000    0.3930
## 15                 1-Dodecanol; 36     0.12000 0.078700    0.3930
## 16                Ribonic acid; 72     0.11400 0.091400    0.4280
## 17               Glycine, 3TMS; 17     0.10200 0.133000    0.5160
## 18        Docosahexaenoic acid; 53    -0.10100 0.135000    0.5160
## 19        4-Deoxytetronic acid; 32     0.10000 0.140000    0.5160
## 20              Arachidic acid; 46    -0.09970 0.144000    0.5160
## 21               Leucine, 2TMS; 19    -0.09900 0.145000    0.5160
## 22                     Ribitol; 70     0.09590 0.160000    0.5450
## 23            Cholesterol, TMS; 23     0.08970 0.183000    0.5980
## 24                Valine, 2TMS; 20    -0.08820 0.193000    0.6030
## 25               Octanoic acid; 68    -0.08510 0.211000    0.6180
## 26       Arachidonic acid, TMS; 24     0.08280 0.225000    0.6180
## 27  2-hydroxy Isovaleric acid; 38     0.08180 0.229000    0.6180
## 28              1-Monopalmitin; 37     0.08180 0.231000    0.6180
## 29            Methionine, 2TMS; 16    -0.07950 0.241000    0.6240
## 30                    Glycerol; 57     0.07640 0.263000    0.6560
## 31                     Ribitol; 71    -0.07400 0.272000    0.6580
## 32          Glyceryl-glycoside; 59     0.07130 0.294000    0.6890
## 33           Aminomalonic acid; 45     0.06830 0.313000    0.6950
## 34            Malic acid, 3TMS; 11     0.06810 0.317000    0.6950
## 35                Serine, 3TMS; 14    -0.06700 0.324000    0.6950
## 36                 Lactic acid; 29     0.06600 0.334000    0.6950
## 37           Fumaric acid, 2TMS; 9     0.06180 0.364000    0.7260
## 38 4-Hydroxyphenyllactic acid; 44     0.06120 0.369000    0.7260
## 39          Glutamic acid, 3TMS; 8     0.05980 0.377000    0.7260
## 40              Hydroxyproline; 64     0.05500 0.419000    0.7670
## 41 3-Hydroxybutyric acid, 2TMS; 1    -0.05490 0.419000    0.7670
## 42           Nonadecanoic acid; 66    -0.05220 0.444000    0.7930
## 43 4-Hydroxybenzeneacetic acid; 4     0.05010 0.459000    0.8000
## 44               Decanoic acid; 52    -0.04630 0.496000    0.8450
```

```
## 45           Myo inositol 6TMS; 1     0.04100 0.541000    0.8790
## 46               Pyruvic acid; 31     0.04100 0.547000    0.8790
## 47         Heptadecanoic acid; 61     0.03940 0.563000    0.8790
## 48            1,3-Propanediol; 34    -0.03890 0.569000    0.8790
## 49                Bisphenol A; 48     0.03840 0.575000    0.8790
## 50        2-Palmitoylglycerol; 39    -0.03570 0.602000    0.9030
## 51                Campesterol; 49    -0.03220 0.635000    0.9080
## 52              Alanine, 2TMS; 25     0.02760 0.686000    0.9080
## 53            Arabinopyranose; 51     0.02570 0.703000    0.9080
## 54 alpha-ketoglutaric acid, TMS M     0.02590 0.704000    0.9080
## 55         11-Eicosenoic acid; 35    -0.02540 0.710000    0.9080
## 56          Linoleic acid, TMS; 4     0.02530 0.710000    0.9080
## 57              Glyceric acid; 30     0.02490 0.713000    0.9080
## 58 2-Hydroxybutyric acid, 2TMS; 2    -0.02440 0.718000    0.9080
## 59             Oleic acid, TMS; 3     0.02450 0.719000    0.9080
## 60         Heptadecanoic acid; 60     0.02340 0.731000    0.9080
## 61             Tartronic acid; 73    -0.02260 0.739000    0.9080
## 62           alpha-Tocopherol; 26    -0.01900 0.780000    0.9410
## 63               Ethanolamine; 56     0.01800 0.791000    0.9410
## 64           Stearic acid, TMS; 2    -0.01660 0.808000    0.9420
## 65          Pyroglutamic acid; 69     0.01580 0.817000    0.9420
## 66       4-Deoxytetronic acid; 33     0.01380 0.838000    0.9490
## 67                   Tyrosine; 75     0.01300 0.848000    0.9490
## 68     3-Indolepropionic acid; 41     0.00961 0.888000    0.9690
## 69          Palmitic acid, TMS; 5    -0.00917 0.893000    0.9690
## 70        3-Indoleacetic acid; 40     0.00802 0.906000    0.9690
## 71            Threonine, 3TMS; 12    -0.00589 0.931000    0.9690
## 72              Proline, 2TMS; 21    -0.00441 0.948000    0.9690
## 73            Dodecanoic acid; 54     0.00370 0.957000    0.9690
## 74        Phenylalanine, 2TMS; 13    -0.00281 0.967000    0.9690
## 75           Myristoleic acid; 65     0.00268 0.969000    0.9690
```

##### 3.3.2 Forest Plot of Model Coefficients

```
## Warning: Ignoring unknown aesthetics: x

## Warning: Ignoring unknown aesthetics: x

## Warning: Ignoring unknown aesthetics: x

## Warning: Ignoring unknown aesthetics: x

## Warning: Ignoring unknown aesthetics: x

## Warning: Ignoring unknown aesthetics: x

## Warning: Ignoring unknown aesthetics: x
```

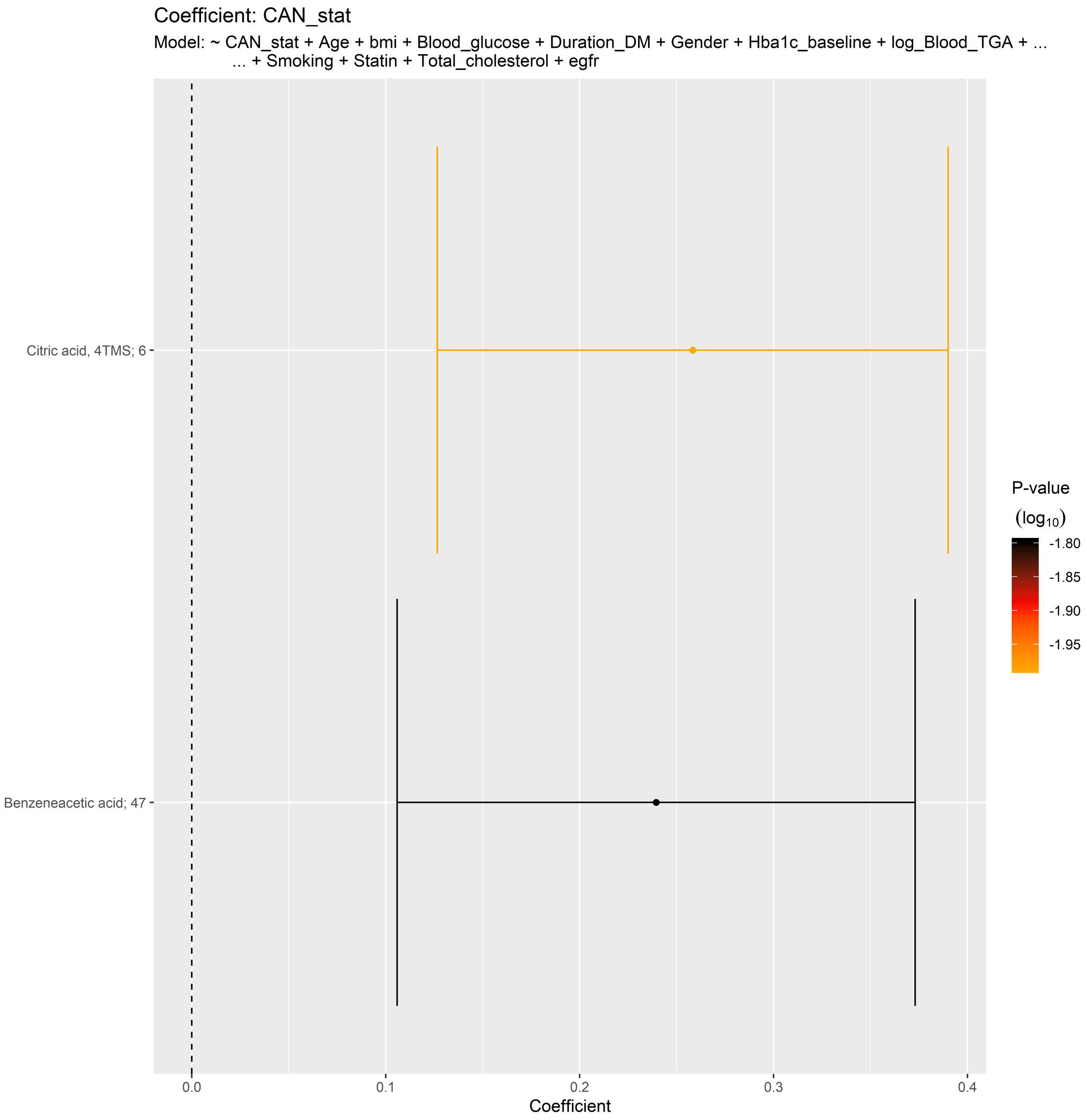
Coefficient: CAN_stat
Model: ~ CAN_stat + Age + bmi + Blood_glucose + Duration_DM + Gender + Hba1c_baseline + log_Blood_TGA + ...
... + Smoking + Statin + Total_cholesterol + egfr
Citric acid, 4TMS; 6
Benzeneacetic acid; 47
0.0
0.1
0.2
0.3
0.4
Coefficient
P-value
(log10)
-1.80
-1.85
-1.90
-1.95

##### 3.3.3 Bipartite Network of Model Coefficients

```
## [1] "bipartite_network_from_limma was created by Tommi Suvitaival"
## [1] ""
## [1] "2019-05-06"

## Warning in if (drop.variables != "none") {: the condition has length > 1 and
## only the first element will be used

## Warning: Removed 5 rows containing missing values (geom_segment).

## Warning: Removed 1 rows containing missing values (geom_text).
```

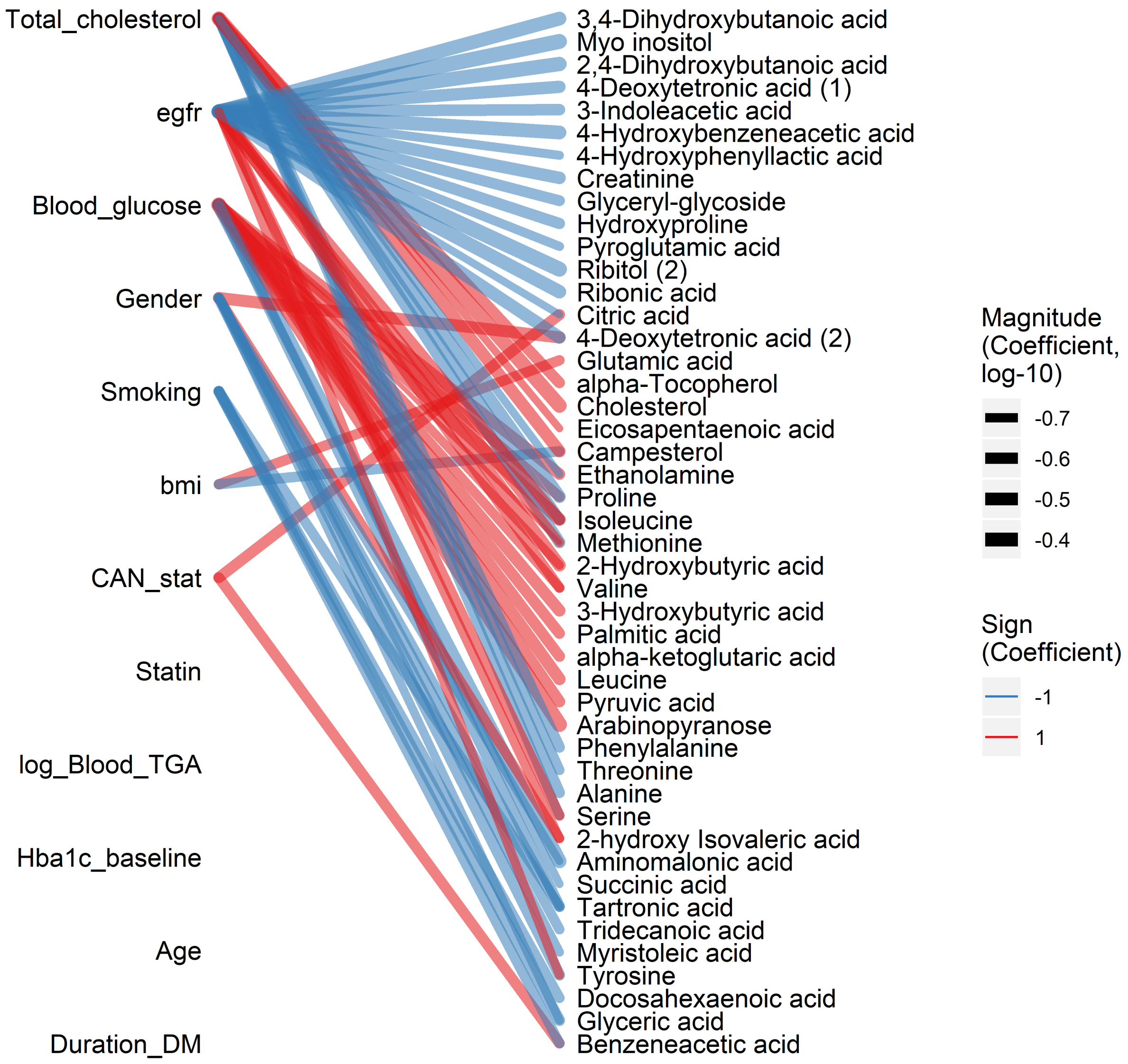

```
## [1] "bipartite_network_from_limma was created by Tommi Suvitaival"
## [1] ""
## [1] "2019-05-06"

## Warning in if (drop.variables != "none") {: the condition has length > 1 and
## only the first element will be used

## Warning: Removed 5 rows containing missing values (geom_segment).

## Warning: Removed 1 rows containing missing values (geom_text).
```

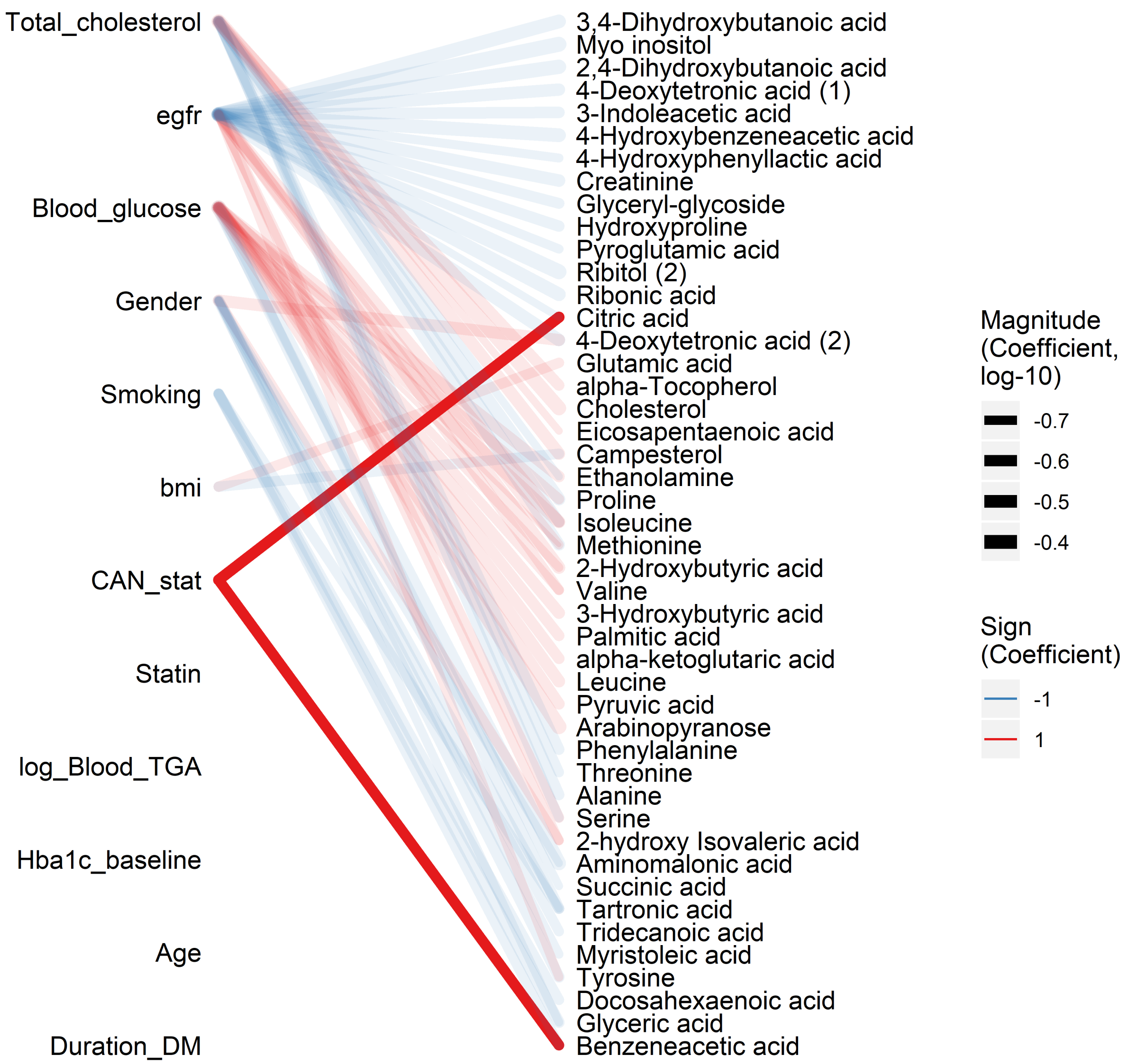

#### 3.4 Combined Forest Plot

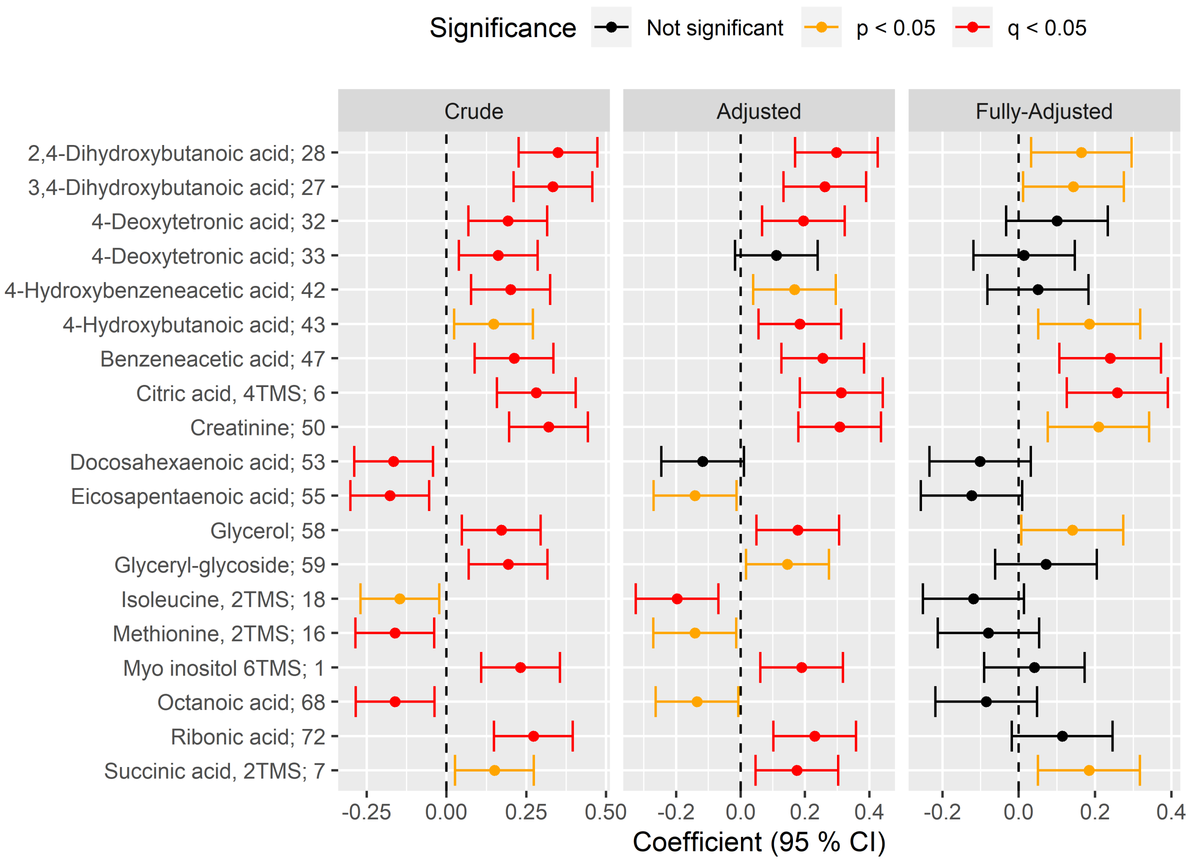

### 4 Vibration Sensation Threshold

#### 4.1 Crude Model

```
## [1] "Fitting models:"
## [1] "~ Vib_pat"
## [1] ""
```

##### 4.1.1 Tables of Model Coefficients

```
## [1] ""
## [1] "Table: Vib_pat"
## [1] "  (from model: "
## [1] "   ~ Vib_pat)"
## [1] ""
##                               Name Coefficient  P.Value adj.P.Val
## 1                      Ribitol; 71     0.23400 6.47e-05   0.00485
## 2  4-Hydroxyphenyllactic acid; 44     0.18700 1.43e-03   0.05360
## 3  3,4-Dihydroxybutanoic acid; 27     0.17100 3.54e-03   0.08840
## 4  2,4-Dihydroxybutanoic acid; 28     0.16400 5.19e-03   0.09720
## 5             Myo inositol 6TMS; 1     0.14700 1.21e-02   0.18200
## 6                      Ribitol; 70     0.13800 1.85e-02   0.23100
## 7  2-Hydroxybutyric acid, 2TMS; 2    -0.13400 2.21e-02   0.23700
## 8  4-Hydroxybenzeneacetic acid; 4     0.12700 3.04e-02   0.24400
## 9             alpha-Tocopherol; 26     0.12500 3.20e-02   0.24400
## 10                Ethanolamine; 56    -0.12500 3.25e-02   0.24400
## 11                L-5-Oxoproline; 63     0.12200 3.65e-02   0.24900
## 12              1,3-Propanediol; 34    -0.12000 3.98e-02   0.24900
## 13           Glyceryl-glycoside; 59     0.11400 5.08e-02   0.29300
## 14                  Creatinine; 50     0.11200 5.64e-02   0.29300
## 15           Nonadecanoic acid; 66    -0.11000 5.94e-02   0.29300
## 16               Hydroxyproline; 64     0.10700 6.62e-02   0.29300
## 17           Fumaric acid, 2TMS; 9     0.10700 6.64e-02   0.29300
## 18        4-Deoxytetronic acid; 32     0.10500 7.21e-02   0.30000
## 19                 Lactic acid; 29    -0.10100 8.54e-02   0.33000
## 20          Heptadecanoic acid; 60    -0.09980 8.79e-02   0.33000
## 21                    Glycerol; 58     0.09720 9.66e-02   0.33300
## 22       Arachidonic acid, TMS; 24    -0.09430 1.07e-01   0.33300
## 23       Eicosapentaenoic acid; 55     0.09330 1.11e-01   0.33300
## 24           Aminomalonic acid; 45    -0.09290 1.12e-01   0.33300
## 25              1-Monopalmitin; 37     0.09250 1.14e-01   0.33300
## 26                Ribonic acid; 72     0.09210 1.15e-01   0.33300
## 27        4-Deoxytetronic acid; 33     0.08350 1.54e-01   0.41900
## 28               Octanoic acid; 68    -0.08230 1.60e-01   0.41900
## 29            Methionine, 2TMS; 16     0.08080 1.67e-01   0.41900
## 30         2-Palmitoylglycerol; 39     0.08060 1.68e-01   0.41900
## 31        Docosahexaenoic acid; 53    -0.07970 1.73e-01   0.41900
## 32               Nonanoic acid; 67    -0.07380 2.07e-01   0.48500
## 33               Decanoic acid; 52     0.07090 2.26e-01   0.48500
## 34             Arachidic acid; 46    -0.07080 2.26e-01   0.48500
## 35             Tartronic acid; 73    -0.06850 2.42e-01   0.48500
## 36               Alanine, 2TMS; 25     0.06750 2.49e-01   0.48500
## 37            Cholesterol, TMS; 23    -0.06550 2.63e-01   0.48500
## 38             Stearic acid, TMS; 2    -0.06480 2.68e-01   0.48500
## 39         3-Indoleacetic acid; 40     0.06480 2.68e-01   0.48500
## 40            Citric acid, 4TMS; 6     0.06470 2.69e-01   0.48500
## 41             Arabinopyranose; 51     0.06440 2.71e-01   0.48500
## 42                    Tyrosine; 75     0.06420 2.72e-01   0.48500
## 43          Heptadecanoic acid; 61    -0.06350 2.78e-01   0.48500
## 44               Glycine, 3TMS; 17     0.06190 2.90e-01   0.49500
## 45                 Bisphenol A; 48    -0.06050 3.01e-01   0.50100
## 46               Glyceric acid; 30    -0.05880 3.15e-01   0.51300
```

```
## 47           Threonine, 3TMS; 12     0.05510 3.46e-01   0.55300
## 48               Pyruvic acid; 31     0.05320 3.63e-01   0.56700
## 49        Benzeneacetic acid; 47     0.04830 4.09e-01   0.62600
## 50              Proline, 2TMS; 21     0.04510 4.41e-01   0.66100
## 51          Pyroglutamic acid; 69     0.04140 4.79e-01   0.70400
## 52                 1-Dodecanol; 36    -0.03940 5.00e-01   0.72200
## 53     4-Hydroxybutanoic acid; 43     0.03560 5.43e-01   0.76500
## 54                   Glycerol; 57    -0.03490 5.51e-01   0.76500
## 55        11-Eicosenoic acid; 35     0.03240 5.80e-01   0.78500
## 56               Serine, 3TMS; 14    -0.03190 5.86e-01   0.78500
## 57            Dodecanoic acid; 54     0.02810 6.30e-01   0.83000
## 58          Linoleic acid, TMS; 4     0.02600 6.57e-01   0.84900
## 59  2-hydroxy Isovaleric acid; 38    -0.02160 7.12e-01   0.90100
## 60            Myristoleic acid; 65     0.02090 7.21e-01   0.90100
## 61 alpha-ketoglutaric acid, TMS M     0.01900 7.45e-01   0.91600
## 62              Hydroxylamine; 62    -0.01620 7.82e-01   0.94600
## 63             Oleic acid, TMS; 3     0.01260 8.29e-01   0.96900
## 64                Campesterol; 49     0.01230 8.34e-01   0.96900
## 65           Tridecanoic acid; 74    -0.01180 8.40e-01   0.96900
## 66           Malic acid, 3TMS; 11     0.00984 8.67e-01   0.97200
## 67        Succinic acid, 2TMS; 7     0.00970 8.68e-01   0.97200
## 68               Valine, 2TMS; 20     0.00762 8.96e-01   0.97200
## 69              Leucine, 2TMS; 19    -0.00682 9.07e-01   0.97200
## 70 3-Hydroxybutyric acid, 2TMS; 1    -0.00633 9.14e-01   0.97200
## 71        Phenylalanine, 2TMS; 13     0.00375 9.49e-01   0.97200
## 72           Isoleucine, 2TMS; 18    -0.00335 9.54e-01   0.97200
## 73        Glutamic acid, 3TMS; 8     0.00330 9.55e-01   0.97200
## 74     3-Indolepropionic acid; 41     0.00299 9.59e-01   0.97200
## 75          Palmitic acid, TMS; 5    -0.00130 9.82e-01   0.98200
```

##### 4.1.2 Forest Plot of Model Coefficients

```
## Warning: Ignoring unknown aesthetics: x
```

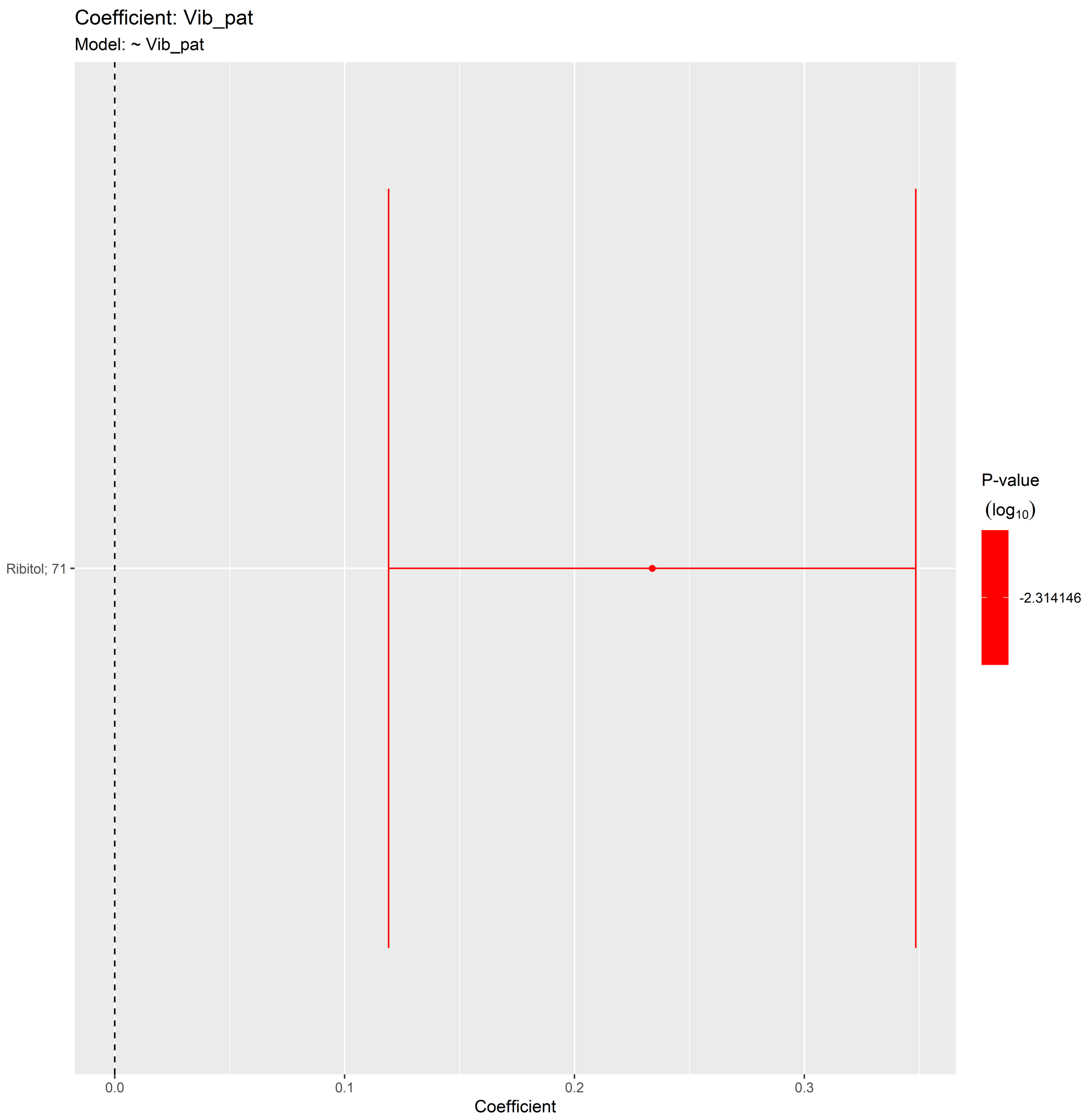

#### 4.2 Adjusted Model

```
## [1] "Fitting models:"
## [1] "~ Vib_pat + Age + bmi + Blood_glucose + Duration_DM + Gender + Hba1c_baseline + log_Blood_TGA +
## [1] ""
```

##### 4.2.1 Tables of Model Coefficients

```
## [1] ""
## [1] "Table: Vib_pat"
## [1] "  (from model: "
## [1] "   ~ Vib_pat + Age + bmi + Blood_glucose + Duration_DM +"
## [1] "     Gender + Hba1c_baseline + log_Blood_TGA + Smoking + Statin +"
## [1] "     Total_cholesterol)"
## [1] ""
##                               Name Coefficient P.Value adj.P.Val
## 1  2-Hydroxybutyric acid, 2TMS; 2    -0.19100 0.00377     0.225
## 2       Arachidonic acid, TMS; 24    -0.17500 0.00764     0.225
## 3                     Ribitol; 71     0.17200 0.00899     0.225
## 4  4-Hydroxyphenyllactic acid; 44     0.15300 0.02030     0.323
## 5        Docosahexaenoic acid; 53    -0.15100 0.02150     0.323
## 6              Hydroxyproline; 64     0.14000 0.03370     0.415
## 7              Arachidic acid; 46    -0.13300 0.04270     0.415
## 8                Ethanolamine; 56    -0.13200 0.04440     0.415
## 9          Nonadecanoic acid; 66    -0.12800 0.05160     0.415
## 10           Stearic acid, TMS; 2    -0.12600 0.05530     0.415
## 11         Heptadecanoic acid; 60    -0.11900 0.06950     0.454
## 12                Lactic acid; 29    -0.11800 0.07390     0.454
## 13            1,3-Propanediol; 34    -0.11500 0.07910     0.454
## 14            Threonine, 3TMS; 12     0.11300 0.08480     0.454
## 15              Octanoic acid; 68    -0.10900 0.09640     0.482
## 16 2,4-Dihydroxybutanoic acid; 28     0.10400 0.11500     0.539
## 17         Heptadecanoic acid; 61    -0.09730 0.13900     0.613
## 18           Myo inositol 6TMS; 1     0.09180 0.16300     0.639
## 19 3-Hydroxybutyric acid, 2TMS; 1    -0.09000 0.17100     0.639
## 20       4-Deoxytetronic acid; 32     0.08950 0.17400     0.639
## 21                 Creatinine; 50     0.08840 0.17900     0.639
## 22             L-5-Oxoproline; 63     0.08090 0.21800     0.651
## 23 alpha-ketoglutaric acid, TMS M    -0.07940 0.22700     0.651
## 24           Aminomalonic acid; 45    -0.07830 0.23400     0.651
## 25         Glyceryl-glycoside; 59     0.07750 0.23800     0.651
## 26                    Ribitol; 70     0.07750 0.23800     0.651
## 27           Isoleucine, 2TMS; 18    -0.07590 0.24800     0.651
## 28                   Glycerol; 58     0.07560 0.25000     0.651
## 29              Glycine, 3TMS; 17     0.07540 0.25200     0.651
## 30        Phenylalanine, 2TMS; 13     0.07030 0.28500     0.690
## 31  2-hydroxy Isovaleric acid; 38    -0.06710 0.30800     0.690
## 32 3,4-Dihydroxybutanoic acid; 27     0.06690 0.30900     0.690
## 33                   Tyrosine; 75     0.06630 0.31300     0.690
## 34           Methionine, 2TMS; 16     0.06570 0.31800     0.690
## 35           Linoleic acid, TMS; 4     0.06350 0.33400     0.690
## 36               Ribonic acid; 72     0.06310 0.33800     0.690
## 37     4-Hydroxybutanoic acid; 43     0.06220 0.34400     0.690
## 38     3-Indolepropionic acid; 41     0.06030 0.35900     0.690
## 39              Leucine, 2TMS; 19    -0.05820 0.37600     0.690
## 40               Pyruvic acid; 31    -0.05800 0.37700     0.690
## 41 4-Hydroxybenzeneacetic acid; 4     0.05710 0.38500     0.690
## 42              Decanoic acid; 52     0.05700 0.38600     0.690
## 43       4-Deoxytetronic acid; 33     0.05300 0.42000     0.703
## 44              Alanine, 2TMS; 25     0.05250 0.42400     0.703
```

```
## 45                 Bisphenol A; 48    -0.05240 0.42600     0.703
## 46             1-Monopalmitin; 37     0.05180 0.43100     0.703
## 47              Glyceric acid; 30    -0.04920 0.45500     0.716
## 48           Citric acid, 4TMS; 6     0.04880 0.45800     0.716
## 49           alpha-Tocopherol; 26     0.04530 0.49100     0.739
## 50                 1-Dodecanol; 36   -0.04510 0.49300     0.739
## 51             Arabinopyranose; 51    0.04280 0.51500     0.753
## 52          Palmitic acid, TMS; 5    -0.04210 0.52200     0.753
## 53        Glutamic acid, 3TMS; 8    -0.03860 0.55800     0.779
## 54          Fumaric acid, 2TMS; 9     0.03830 0.56100     0.779
## 55           Malic acid, 3TMS; 11    -0.03610 0.58300     0.787
## 56           Tridecanoic acid; 74     0.03560 0.58800     0.787
## 57              Nonanoic acid; 67    -0.03470 0.59800     0.787
## 58              Hydroxylamine; 62    -0.03140 0.63300     0.787
## 59                   Glycerol; 57     0.03130 0.63400     0.787
## 60           Cholesterol, TMS; 23    -0.03030 0.64500     0.787
## 61       2-Palmitoylglycerol; 39     0.02970 0.65200     0.787
## 62             Oleic acid, TMS; 3    -0.02950 0.65400     0.787
## 63             Tartronic acid; 73    -0.02880 0.66100     0.787
## 64           Myristoleic acid; 65     0.02750 0.67600     0.792
## 65         Benzeneacetic acid; 47     0.02250 0.73200     0.845
## 66               Valine, 2TMS; 20     0.02010 0.76000     0.863
## 67            Dodecanoic acid; 54     0.01790 0.78600     0.880
## 68          Pyroglutamic acid; 69    -0.01460 0.82400     0.904
## 69        Succinic acid, 2TMS; 7     0.01400 0.83200     0.904
## 70               Serine, 3TMS; 14     0.00880 0.89400     0.945
## 71                Campesterol; 49    -0.00810 0.90200     0.945
## 72       3-Indoleacetic acid; 40    -0.00767 0.90700     0.945
## 73        11-Eicosenoic acid; 35    -0.00562 0.93200     0.948
## 74     Eicosapentaenoic acid; 55    -0.00513 0.93800     0.948
## 75              Proline, 2TMS; 21     0.00427 0.94800     0.948
```

##### 4.2.2 Forest Plot of Model Coefficients

```
## Warning: Ignoring unknown aesthetics: x

## Warning: Ignoring unknown aesthetics: x

## Warning: Ignoring unknown aesthetics: x

## Warning: Ignoring unknown aesthetics: x

## Warning: Ignoring unknown aesthetics: x

## Warning: Ignoring unknown aesthetics: x

## Warning: Ignoring unknown aesthetics: x
```

```
## NULL
```

##### 4.2.3 Bipartite Network of Model Coefficients

```
## [1] "bipartite_network_from_limma was created by Tommi Suvitaival"
## [1] ""
## [1] "2019-05-06"

## Warning in if (drop.variables != "none") {: the condition has length > 1 and
## only the first element will be used

## Warning: Removed 4 rows containing missing values (geom_segment).

## Warning: Removed 1 rows containing missing values (geom_text).
```

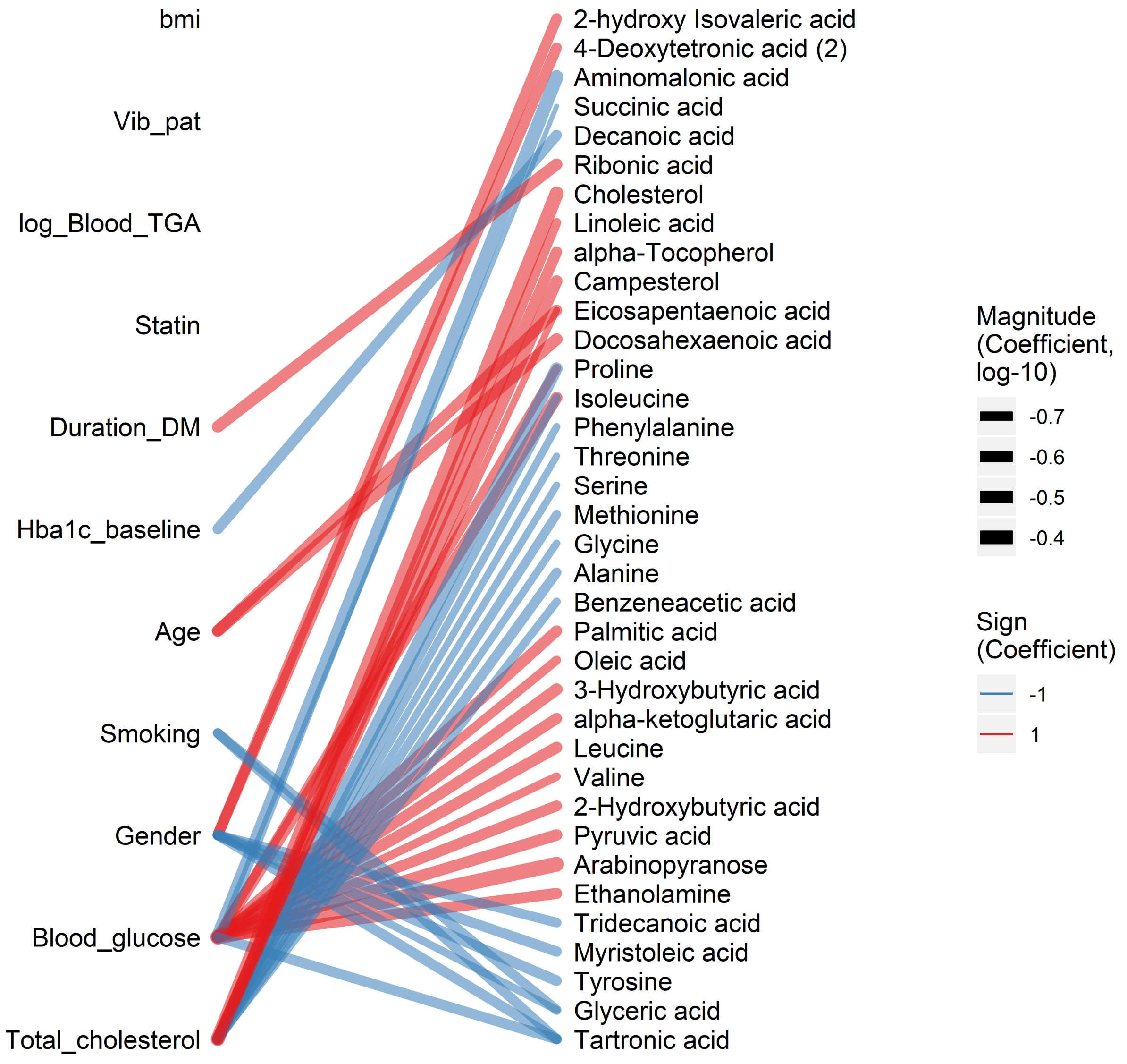

```
## [1] "bipartite_network_from_limma was created by Tommi Suvitaival"
## [1] ""
## [1] "2019-05-06"

## Warning in if (drop.variables != "none") {: the condition has length > 1 and
## only the first element will be used

## Warning: Removed 4 rows containing missing values (geom_segment).

## Warning: Removed 1 rows containing missing values (geom_text).
```

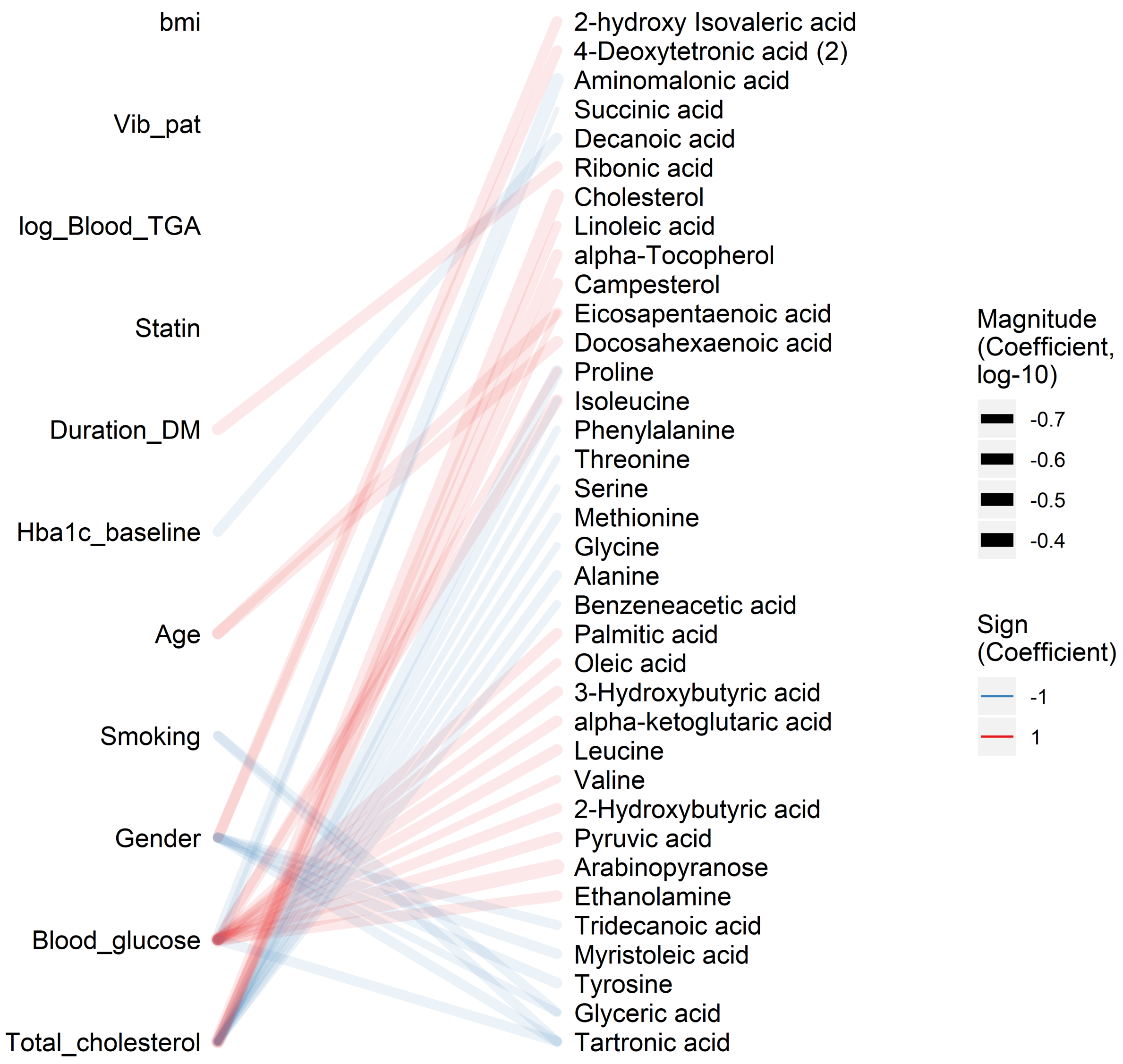

##### 4.3 Fully Adjusted Model

```
## [1] "Fitting models:"
## [1] "~ Vib_pat + Age + bmi + Blood_glucose + Duration_DM + Gender + Hba1c_baseline + log_Blood_TGA +
## [1] ""
```

###### 4.3.1 Tables of Model Coefficients

```
## [1] ""
## [1] "Table: Vib_pat"
## [1] "  (from model: "
## [1] "   ~ Vib_pat + Age + bmi + Blood_glucose + Duration_DM +"
## [1] "     Gender + Hba1c_baseline + log_Blood_TGA + Smoking + Statin +"
## [1] "     Total_cholesterol + egfr)"
## [1] ""
##                               Name Coefficient P.Value adj.P.Val
## 1       Arachidonic acid, TMS; 24    -0.16500  0.0128     0.610
## 2             Threonine, 3TMS; 12     0.13700  0.0385     0.610
## 3       Docosahexaenoic acid; 53    -0.13600  0.0386     0.610
## 4  2-Hydroxybutyric acid, 2TMS; 2    -0.13300  0.0432     0.610
## 5              Arachidic acid; 46    -0.12900  0.0511     0.610
## 6          Heptadecanoic acid; 60    -0.12500  0.0594     0.610
## 7           Nonadecanoic acid; 66    -0.12000  0.0698     0.610
## 8             1,3-Propanediol; 34    -0.11900  0.0709     0.610
## 9                Ethanolamine; 56    -0.11800  0.0732     0.610
## 10 4-Hydroxyphenyllactic acid; 44     0.11500  0.0822     0.616
## 11                Lactic acid; 29    -0.10700  0.1060     0.634
## 12                   Tyrosine; 75     0.10600  0.1090     0.634
## 13           Methionine, 2TMS; 16     0.10500  0.1100     0.634
## 14           Stearic acid, TMS; 2    -0.10200  0.1230     0.661
## 15 3-Hydroxybutyric acid, 2TMS; 1    -0.09160  0.1650     0.750
## 16              Hydroxyproline; 64     0.09040  0.1710     0.750
## 17         Heptadecanoic acid; 61    -0.09030  0.1720     0.750
## 18              L-5-Oxoproline; 63     0.08860  0.1800     0.750
## 19 alpha-ketoglutaric acid, TMS M    -0.08480  0.1990     0.784
## 20          Aminomalonic acid; 45    -0.08140  0.2160     0.811
## 21              Octanoic acid; 68    -0.07690  0.2440     0.849
## 22           Malic acid, 3TMS; 11    -0.07480  0.2570     0.849
## 23                    Ribitol; 71     0.07040  0.2830     0.849
## 24               Valine, 2TMS; 20     0.07070  0.2830     0.849
## 25              Decanoic acid; 52     0.06850  0.2990     0.849
## 26        Phenylalanine, 2TMS; 13     0.06340  0.3370     0.849
## 27     3-Indolepropionic acid; 41     0.06300  0.3400     0.849
## 28               Serine, 3TMS; 14     0.06120  0.3530     0.849
## 29                    Ribitol; 70     0.05960  0.3670     0.849
## 30           Linoleic acid, TMS; 4     0.05870  0.3740     0.849
## 31                   Glycerol; 57     0.05760  0.3830     0.849
## 32        3-Indoleacetic acid; 40    -0.05610  0.3960     0.849
## 33          Pyroglutamic acid; 69    -0.05590  0.3980     0.849
## 34               Pyruvic acid; 31    -0.05400  0.4130     0.849
## 35           alpha-Tocopherol; 26     0.05210  0.4290     0.849
## 36     4-Hydroxybutanoic acid; 43     0.05180  0.4340     0.849
## 37              Glycine, 3TMS; 17     0.04970  0.4520     0.849
## 38              Alanine, 2TMS; 25     0.04850  0.4630     0.849
## 39                 1-Dodecanol; 36    -0.04850  0.4630     0.849
## 40               Hydroxylamine; 62    -0.04750  0.4730     0.849
## 41                 Bisphenol A; 48    -0.04660  0.4810     0.849
## 42                   Glycerol; 58     0.04610  0.4860     0.849
## 43              1-Monopalmitin; 37     0.04600  0.4870     0.849
## 44              Nonanoic acid; 67    -0.04130  0.5330     0.909
```

```
## 45            Arabinopyranose; 51     0.03440  0.6000     0.930
## 46        2-Palmitoylglycerol; 39     0.03370  0.6100     0.930
## 47             Tartronic acid; 73    -0.03170  0.6300     0.930
## 48  2-hydroxy Isovaleric acid; 38    -0.03130  0.6350     0.930
## 49           Tridecanoic acid; 74     0.03050  0.6440     0.930
## 50              Leucine, 2TMS; 19    -0.02790  0.6730     0.930
## 51            Dodecanoic acid; 54     0.02790  0.6730     0.930
## 52           Isoleucine, 2TMS; 18    -0.02730  0.6790     0.930
## 53           Myristoleic acid; 65     0.02520  0.7030     0.930
## 54         Glyceryl-glycoside; 59     0.02490  0.7060     0.930
## 55             Oleic acid, TMS; 3    -0.02480  0.7070     0.930
## 56          Palmitic acid, TMS; 5    -0.02430  0.7120     0.930
## 57 4-Hydroxybenzeneacetic acid; 4    -0.02380  0.7180     0.930
## 58             Glyceric acid; 30    -0.02250  0.7330     0.930
## 59               Ribonic acid; 72    -0.02230  0.7340     0.930
## 60      4-Deoxytetronic acid; 32     0.02150  0.7440     0.930
## 61         Benzeneacetic acid; 47     0.01710  0.7960     0.937
## 62 3,4-Dihydroxybutanoic acid; 27    -0.01690  0.7960     0.937
## 63     Eicosapentaenoic acid; 55     0.01680  0.7980     0.937
## 64      4-Deoxytetronic acid; 33    -0.01670  0.7990     0.937
## 65         11-Eicosenoic acid; 35    -0.01260  0.8490     0.939
## 66           Myo inositol 6TMS; 1    -0.01130  0.8630     0.939
## 67         Glutamic acid, 3TMS; 8    -0.00937  0.8870     0.939
## 68 2,4-Dihydroxybutanoic acid; 28     0.00906  0.8900     0.939
## 69           Citric acid, 4TMS; 6    -0.00805  0.9030     0.939
## 70               Campesterol; 49    -0.00788  0.9050     0.939
## 71         Succinic acid, 2TMS; 7     0.00643  0.9230     0.939
## 72             Proline, 2TMS; 21    -0.00637  0.9230     0.939
## 73                Creatinine; 50     0.00613  0.9260     0.939
## 74          Fumaric acid, 2TMS; 9     0.00578  0.9300     0.939
## 75           Cholesterol, TMS; 23    -0.00498  0.9390     0.939
```

##### 4.3.2 Forest Plot of Model Coefficients

```
## Warning: Ignoring unknown aesthetics: x

## Warning: Ignoring unknown aesthetics: x

## Warning: Ignoring unknown aesthetics: x

## Warning: Ignoring unknown aesthetics: x

## Warning: Ignoring unknown aesthetics: x

## Warning: Ignoring unknown aesthetics: x

## Warning: Ignoring unknown aesthetics: x

## NULL
```

##### 4.3.3 Bipartite Network of Model Coefficients

```
## [1] "bipartite_network_from_limma was created by Tommi Suvitaival"
## [1] ""
## [1] "2019-05-06"

## Warning in if (drop.variables != "none") {: the condition has length > 1 and
## only the first element will be used

## Warning: Removed 5 rows containing missing values (geom_segment).

## Warning: Removed 1 rows containing missing values (geom_text).
```

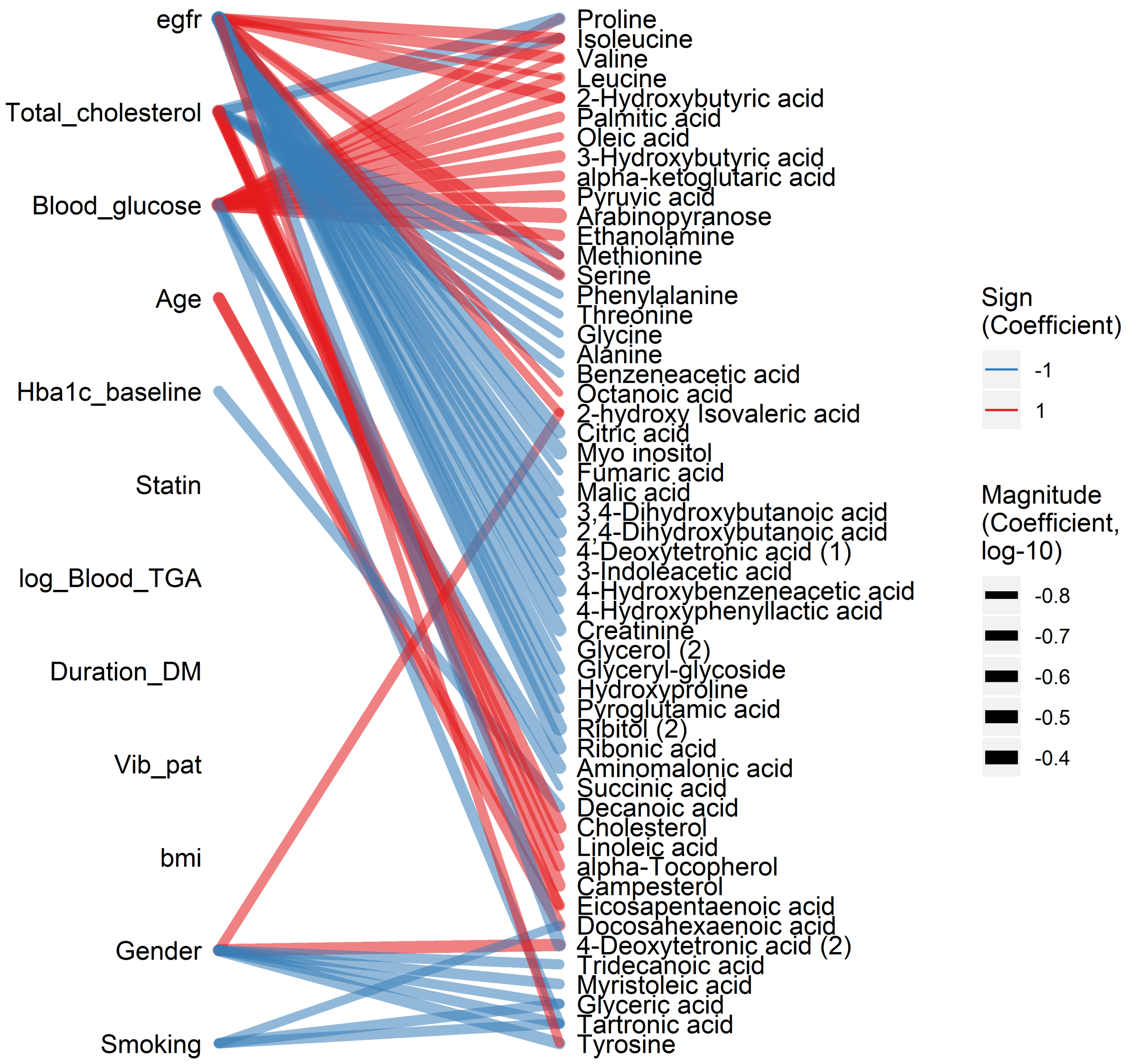

```
## [1] "bipartite_network_from_limma was created by Tommi Suvitaival"
## [1] ""
## [1] "2019-05-06"

## Warning in if (drop.variables != "none") {: the condition has length > 1 and
## only the first element will be used

## Warning: Removed 5 rows containing missing values (geom_segment).

## Warning: Removed 1 rows containing missing values (geom_text).
```

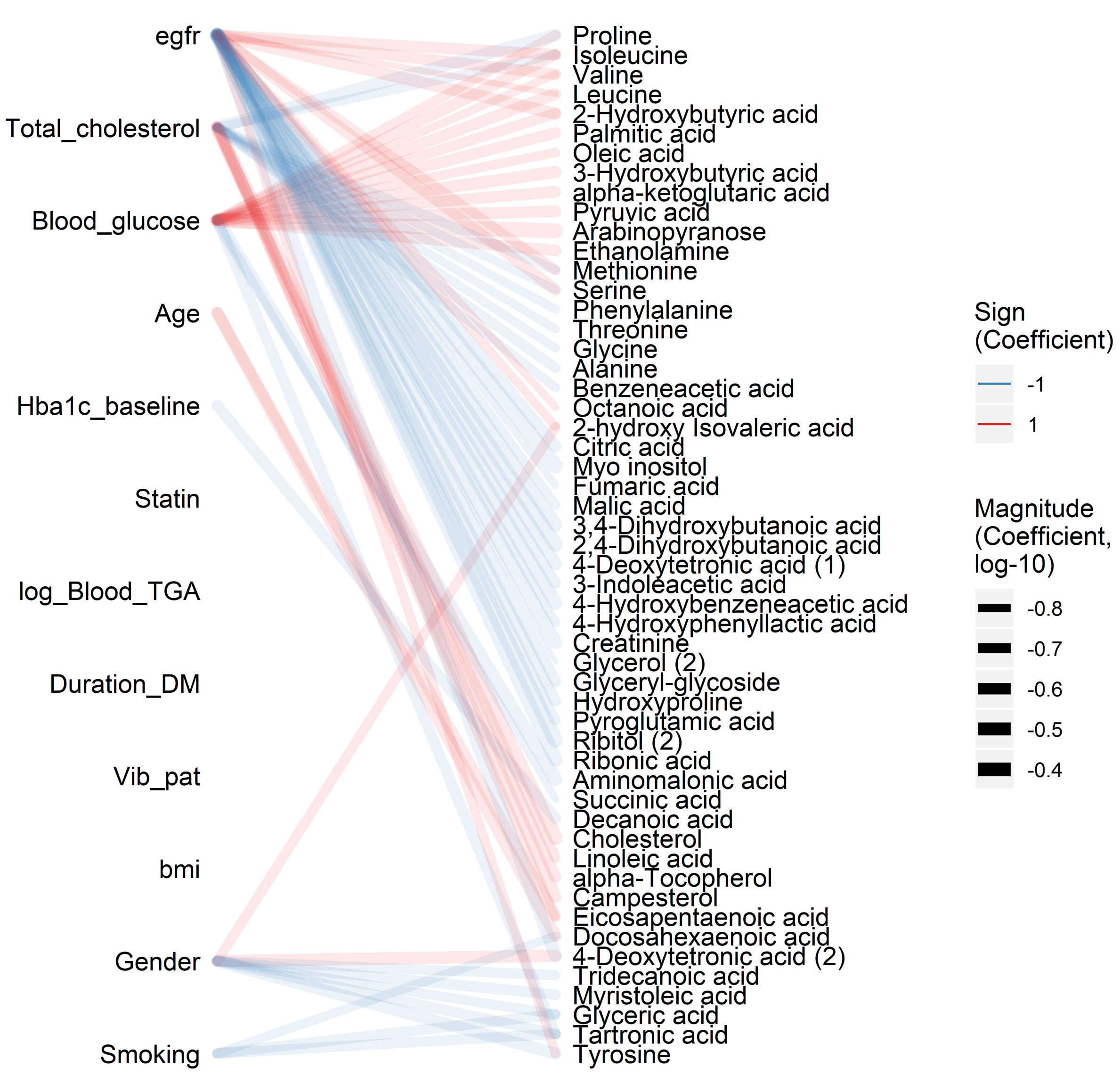

### 5 Secondary Analyses

#### 5.1 Resting HR Vagus

##### 5.1.1 Crude Model

```
## [1] "Fitting models:"
## [1] "~ rest_HR_vag"
## [1] ""
```

###### 5.1.1.1 Tables of Model Coefficients

```
## [1] ""
## [1] "Table: rest_HR_vag"
## [1] "  (from model: "
## [1] "   ~ rest_HR_vag)"
## [1] ""
##                               Name Coefficient  P.Value adj.P.Val
## 1           Methionine, 2TMS; 16    -0.20500 0.000588    0.0441
## 2              Oleic acid, TMS; 3     0.17200 0.003820    0.1040
## 3             Threonine, 3TMS; 12    -0.17100 0.004140    0.1040
## 4  3,4-Dihydroxybutanoic acid; 27     0.15800 0.008000    0.1500
## 5        Docosahexaenoic acid; 53    -0.14900 0.012400    0.1660
## 6                Valine, 2TMS; 20    -0.14800 0.013300    0.1660
## 7  4-Hydroxyphenyllactic acid; 44     0.13800 0.020500    0.1860
## 8  2-Hydroxybutyric acid, 2TMS; 2     0.13600 0.022200    0.1860
## 9      3-Indolepropionic acid; 41    -0.13500 0.022900    0.1860
## 10         Succinic acid, 2TMS; 7     0.13300 0.025900    0.1860
## 11              Tartronic acid; 73    -0.13100 0.027200    0.1860
## 12             1,3-Propanediol; 34    -0.11500 0.053500    0.2840
## 13                     Ribitol; 71     0.11500 0.053600    0.2840
## 14          Palmitic acid, TMS; 5     0.11400 0.055300    0.2840
## 15                 Lactic acid; 29     0.11300 0.056900    0.2840
## 16                    Glycerol; 57     0.11000 0.065800    0.2990
## 17 4-Hydroxybenzeneacetic acid; 4     0.10900 0.067700    0.2990
## 18           Stearic acid, TMS; 2     0.10600 0.074900    0.3120
## 19         11-Eicosenoic acid; 35     0.10400 0.079500    0.3140
## 20          Nonadecanoic acid; 66    -0.09580 0.108000    0.4040
## 21          Fumaric acid, 2TMS; 9     0.08680 0.145000    0.4830
## 22       Phenylalanine, 2TMS; 13    -0.08620 0.148000    0.4830
## 23              Serine, 3TMS; 14    -0.08540 0.151000    0.4830
## 24                Ethanolamine; 56    -0.08310 0.163000    0.4830
## 25              L-5-Oxoproline; 63    -0.08260 0.166000    0.4830
## 26 3-Hydroxybutyric acid, 2TMS; 1     0.08120 0.173000    0.4830
## 27               Leucine, 2TMS; 19    -0.07900 0.185000    0.4830
## 28 2,4-Dihydroxybutanoic acid; 28     0.07600 0.202000    0.4830
## 29      Eicosapentaenoic acid; 55    -0.07560 0.204000    0.4830
## 30           Myristoleic acid; 65     0.07550 0.205000    0.4830
## 31             Dodecanoic acid; 54     0.07490 0.209000    0.4830
## 32                Ribonic acid; 72     0.07390 0.214000    0.4830
## 33       4-Deoxytetronic acid; 33     0.07310 0.219000    0.4830
## 34           alpha-Tocopherol; 26    -0.07170 0.229000    0.4830
## 35         Glutamic acid, 3TMS; 8     0.07070 0.235000    0.4830
## 36           Myo inositol 6TMS; 1     0.07020 0.239000    0.4830
## 37                    Glycerol; 58     0.07020 0.239000    0.4830
## 38              Glyceric acid; 30    -0.06930 0.245000    0.4830
## 39                  Creatinine; 50     0.06760 0.257000    0.4880
## 40                Campesterol; 49     0.06660 0.263000    0.4880
## 41      Arachidonic acid, TMS; 24     0.06620 0.267000    0.4880
## 42            Aminomalonic acid; 45    -0.06320 0.289000    0.5040
## 43         Heptadecanoic acid; 61     0.06310 0.289000    0.5040
## 44                     Ribitol; 70     0.06180 0.300000    0.5110
## 45         Heptadecanoic acid; 60     0.05380 0.366000    0.6060
## 46              Nonanoic acid; 67     0.05320 0.372000    0.6060
```

```
## 47               Bisphenol A; 48    -0.05010 0.400000    0.6390
## 48  2-hydroxy Isovaleric acid; 38     0.04790 0.421000    0.6580
## 49          Malic acid, 3TMS; 11     0.04700 0.430000    0.6590
## 50             Decanoic acid; 52     0.04600 0.440000    0.6600
## 51            Arachidic acid; 46     0.04430 0.457000    0.6720
## 52          Isoleucine, 2TMS; 18    -0.04140 0.487000    0.7020
## 53               1-Dodecanol; 36     0.03880 0.515000    0.7160
## 54                  Tyrosine; 75    -0.03870 0.516000    0.7160
## 55    4-Hydroxybutanoic acid; 43     0.03730 0.531000    0.7250
## 56      4-Deoxytetronic acid; 32    -0.03590 0.547000    0.7320
## 57             Octanoic acid; 68    -0.03420 0.566000    0.7450
## 58          Tridecanoic acid; 74    -0.03250 0.585000    0.7570
## 59             Hydroxylamine; 62    -0.03040 0.610000    0.7760
## 60             Alanine, 2TMS; 25    -0.02550 0.668000    0.8360
## 61        Benzeneacetic acid; 47    -0.02140 0.720000    0.8850
## 62             Proline, 2TMS; 21    -0.01970 0.741000    0.8940
## 63            Hydroxyproline; 64     0.01750 0.769000    0.8940
## 64           Arabinopyranose; 51    -0.01690 0.776000    0.8940
## 65          Citric acid, 4TMS; 6     0.01600 0.788000    0.8940
## 66         Pyroglutamic acid; 69    -0.01550 0.795000    0.8940
## 67             Glycine, 3TMS; 17    -0.01490 0.802000    0.8940
## 68       2-Palmitoylglycerol; 39    -0.01430 0.810000    0.8940
## 69               Pyruvic acid; 31    -0.01170 0.845000    0.9050
## 70          Cholesterol, TMS; 23     0.01110 0.852000    0.9050
## 71 alpha-ketoglutaric acid, TMS M    -0.01010 0.866000    0.9050
## 72        Glyceryl-glycoside; 59     0.00982 0.869000    0.9050
## 73       3-Indoleacetic acid; 40    -0.00800 0.893000    0.9180
## 74         Linoleic acid, TMS; 4    -0.00385 0.948000    0.9610
## 75            1-Monopalmitin; 37     0.00277 0.963000    0.9630
```

###### 5.1.1.2 Forest Plot of Model Coefficients

```
## Warning: Ignoring unknown aesthetics: x
```

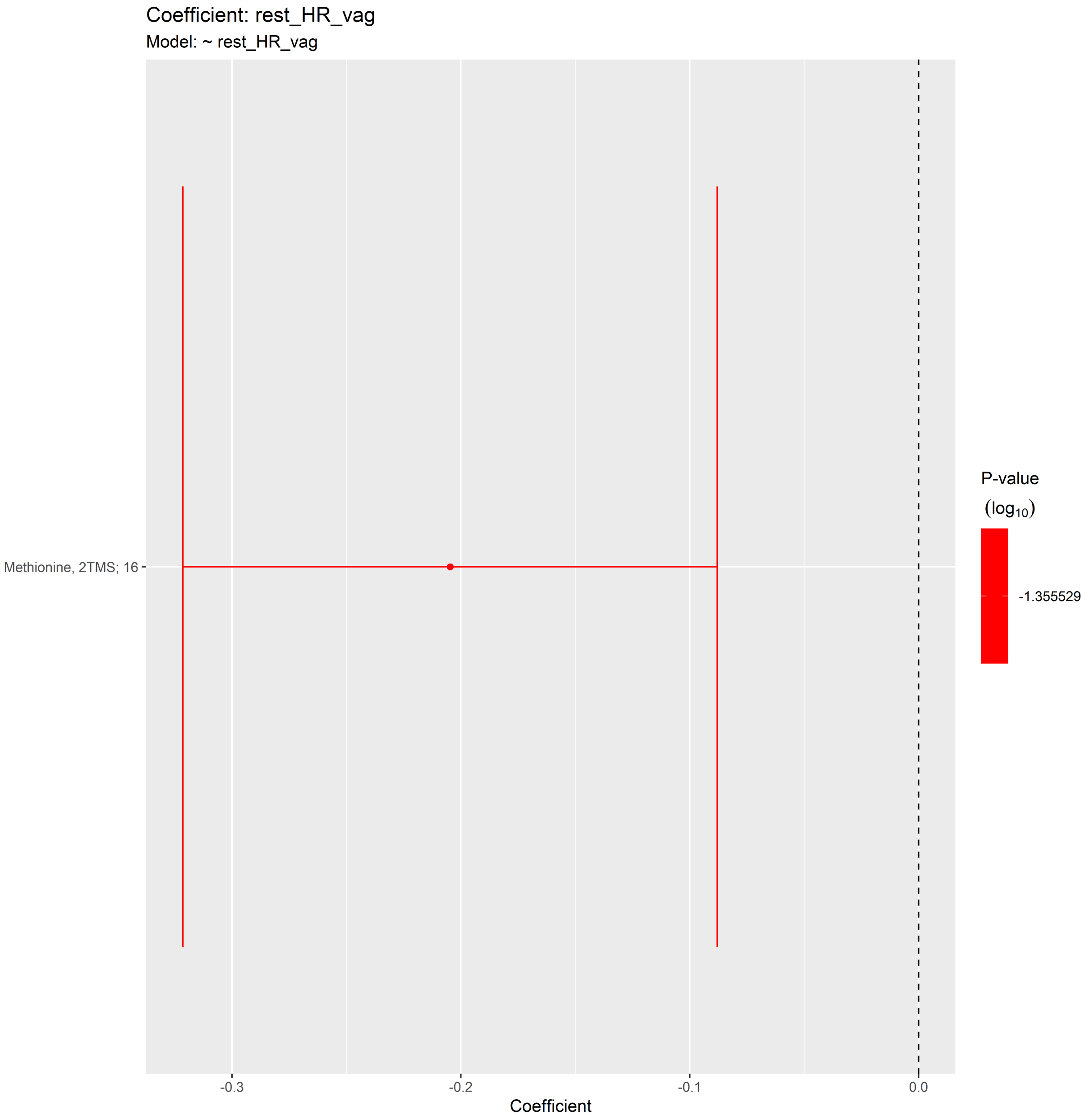

##### 5.1.2 Adjusted Model

```
## [1] "Fitting models:"
## [1] "~ rest_HR_vag + Age + bmi + Blood_glucose + Duration_DM + Gender + Hba1c_baseline + log_Blood_T
## [1] ""
```

###### 5.1.2.1 Tables of Model Coefficients

```
## [1] ""
## [1] "Table: rest_HR_vag"
## [1] "  (from model: "
## [1] "   ~ rest_HR_vag + Age + bmi + Blood_glucose + Duration_DM +"
## [1] "     Gender + Hba1c_baseline + log_Blood_TGA + Smoking + Statin +"
## [1] "     Total_cholesterol)"
## [1] ""
##                              Name Coefficient P.Value adj.P.Val
## 1             Threonine, 3TMS; 12    -0.18600 0.00280     0.102
## 2            Methionine, 2TMS; 16    -0.18200 0.00337     0.102
## 3              Oleic acid, TMS; 3     0.17800 0.00410     0.102
## 4  4-Hydroxyphenyllactic acid; 44     0.16400 0.00853     0.159
## 5                Valine, 2TMS; 20    -0.15500 0.01260     0.159
## 6          Succinic acid, 2TMS; 7     0.15500 0.01270     0.159
## 7           Palmitic acid, TMS; 5     0.13900 0.02550     0.273
## 8                     Glycerol; 58     0.13400 0.03130     0.293
## 9          11-Eicosenoic acid; 35     0.12300 0.04810     0.390
## 10         Heptadecanoic acid; 61     0.11900 0.05550     0.390
## 11 4-Hydroxybenzeneacetic acid; 4     0.11400 0.06620     0.390
## 12           Stearic acid, TMS; 2     0.11100 0.07510     0.390
## 13                Lactic acid; 29     0.11100 0.07530     0.390
## 14              Leucine, 2TMS; 19    -0.11000 0.07640     0.390
## 15 2-Hydroxybutyric acid, 2TMS; 2     0.11000 0.07810     0.390
## 16                    Ribitol; 71     0.10400 0.09300     0.436
## 17 3,4-Dihydroxybutanoic acid; 27     0.10000 0.10700     0.470
## 18  2-hydroxy Isovaleric acid; 38     0.09610 0.12200     0.496
## 19        Phenylalanine, 2TMS; 13    -0.09130 0.14200     0.496
## 20       Docosahexaenoic acid; 53    -0.09050 0.14500     0.496
## 21               Serine, 3TMS; 14    -0.08950 0.15000     0.496
## 22           Malic acid, 3TMS; 11     0.08930 0.15100     0.496
## 23     3-Indolepropionic acid; 41    -0.08900 0.15200     0.496
## 24             Tartronic acid; 73    -0.08590 0.16700     0.512
## 25          Fumaric acid, 2TMS; 9     0.08510 0.17100     0.512
## 26               Ethanolamine; 56    -0.08350 0.17900     0.517
## 27 3-Hydroxybutyric acid, 2TMS; 1     0.07950 0.20100     0.541
## 28           Myristoleic acid; 65     0.07800 0.20900     0.541
## 29              Hydroxylamine; 62    -0.07750 0.21300     0.541
## 30       4-Deoxytetronic acid; 32    -0.07680 0.21600     0.541
## 31              Nonanoic acid; 67     0.07550 0.22500     0.543
## 32            Dodecanoic acid; 54     0.07420 0.23300     0.545
## 33                   Tyrosine; 75    -0.07020 0.25900     0.561
## 34                   Glycerol; 57     0.06930 0.26500     0.561
## 35            1,3-Propanediol; 34    -0.06910 0.26600     0.561
## 36          Nonadecanoic acid; 66    -0.06830 0.27200     0.561
## 37            Arabinopyranose; 51    -0.06580 0.29000     0.561
## 38           alpha-Tocopherol; 26    -0.06550 0.29200     0.561
## 39           Citric acid, 4TMS; 6     0.06460 0.29900     0.561
## 40             Arachidic acid; 46     0.06450 0.29900     0.561
## 41       Arachidonic acid, TMS; 24    0.06210 0.31800     0.582
## 42         Heptadecanoic acid; 60     0.05860 0.34600     0.611
## 43              Decanoic acid; 52     0.05800 0.35000     0.611
## 44           Isoleucine, 2TMS; 18    -0.05450 0.38000     0.632
```

```
## 45           Myo inositol 6TMS; 1     0.05430 0.38200     0.632
## 46             L-5-Oxoproline; 63    -0.05370 0.38800     0.632
## 47                1-Dodecanol; 36     0.05090 0.41300     0.647
## 48               Ribonic acid; 72     0.05080 0.41400     0.647
## 49              Alanine, 2TMS; 25    -0.04490 0.47000     0.719
## 50         Glyceryl-glycoside; 59    -0.03990 0.52100     0.754
## 51 2,4-Dihydroxybutanoic acid; 28     0.03970 0.52300     0.754
## 52                Bisphenol A; 48    -0.03970 0.52300     0.754
## 53         Benzeneacetic acid; 47     0.03830 0.53800     0.758
## 54              Proline, 2TMS; 21    -0.03750 0.54600     0.758
## 55       4-Deoxytetronic acid; 33     0.03630 0.55900     0.763
## 56             1-Monopalmitin; 37     0.03390 0.58500     0.784
## 57                Campesterol; 49     0.03100 0.61800     0.796
## 58    4-Hydroxybutanoic acid; 43      0.03020 0.62700     0.796
## 59                 Creatinine; 50     0.03020 0.62700     0.796
## 60     Eicosapentaenoic acid; 55     -0.02940 0.63700     0.796
## 61              Octanoic acid; 68    -0.02740 0.65900     0.802
## 62           Tridecanoic acid; 74    -0.02710 0.66300     0.802
## 63              Glycine, 3TMS; 17     0.02250 0.71700     0.854
## 64 alpha-ketoglutaric acid, TMS M    -0.02000 0.74800     0.876
## 65              Glyceric acid; 30    -0.01700 0.78500     0.906
## 66          Pyroglutamic acid; 69    -0.01350 0.82800     0.941
## 67        3-Indoleacetic acid; 40    -0.01220 0.84400     0.945
## 68               Pyruvic acid; 31    -0.01070 0.86300     0.952
## 69             Hydroxyproline; 64     0.00942 0.88000     0.956
## 70          Aminomalonic acid; 45    -0.00787 0.89900     0.957
## 71        2-Palmitoylglycerol; 39    -0.00732 0.90600     0.957
## 72         Glutamic acid, 3TMS; 8     0.00611 0.92200     0.960
## 73           Cholesterol, TMS; 23     0.00414 0.94700     0.973
## 74          Linoleic acid, TMS; 4     0.00233 0.97000     0.983
## 75                    Ribitol; 70    -0.00113 0.98500     0.985
```

###### 5.1.2.2 Forest Plot of Model Coefficients

```
## Warning: Ignoring unknown aesthetics: x

## Warning: Ignoring unknown aesthetics: x

## Warning: Ignoring unknown aesthetics: x

## Warning: Ignoring unknown aesthetics: x

## Warning: Ignoring unknown aesthetics: x

## Warning: Ignoring unknown aesthetics: x

## NULL
```

###### 5.1.3 Fully-Adjusted Model

```
## [1] "Fitting models:"
## [1] "~ rest_HR_vag + Age + bmi + Blood_glucose + Duration_DM + Gender + Hba1c_baseline + log_Blood_TG
## [1] ""
```

###### 5.1.3.1 Tables of Model Coefficients

```
## [1] ""
## [1] "Table: rest_HR_vag"
## [1] "  (from model: "
## [1] "   ~ rest_HR_vag + Age + bmi + Blood_glucose + Duration_DM +"
## [1] "     Gender + Hba1c_baseline + log_Blood_TGA + Smoking + Statin +"
## [1] "     Total_cholesterol + egfr)"
## [1] ""
##                               Name Coefficient P.Value adj.P.Val
## 1              Oleic acid, TMS; 3     0.18000 0.00334     0.128
## 2             Threonine, 3TMS; 12    -0.18000 0.00340     0.128
## 3            Methionine, 2TMS; 16    -0.17000 0.00556     0.139
## 4           Succinic acid, 2TMS; 7     0.15400 0.01250     0.221
## 5  4-Hydroxyphenyllactic acid; 44     0.14900 0.01530     0.221
## 6            Palmitic acid, TMS; 5     0.14600 0.01770     0.221
## 7                 Valine, 2TMS; 20    -0.14000 0.02210     0.236
## 8  2-Hydroxybutyric acid, 2TMS; 2     0.13200 0.03100     0.290
## 9                     Glycerol; 58     0.12400 0.04340     0.311
## 10         Heptadecanoic acid; 61     0.12300 0.04560     0.311
## 11          11-Eicosenoic acid; 35     0.12100 0.04850     0.311
## 12            Stearic acid, TMS; 2     0.12100 0.04970     0.311
## 13                 Lactic acid; 29     0.11600 0.05920     0.342
## 14  2-hydroxy Isovaleric acid; 38     0.10900 0.07580     0.406
## 15       4-Deoxytetronic acid; 32    -0.10200 0.09760     0.479
## 16               Leucine, 2TMS; 19    -0.10000 0.10200     0.479
## 17        Phenylalanine, 2TMS; 13    -0.09560 0.12000     0.530
## 18     3-Indolepropionic acid; 41    -0.08920 0.14700     0.540
## 19              Tartronic acid; 73    -0.08720 0.15400     0.540
## 20 4-Hydroxybenzeneacetic acid; 4     0.08630 0.15900     0.540
## 21               Hydroxylamine; 62    -0.08600 0.16200     0.540
## 22       Docosahexaenoic acid; 53    -0.08210 0.18000     0.540
## 23 3-Hydroxybutyric acid, 2TMS; 1     0.08030 0.19100     0.540
## 24           Malic acid, 3TMS; 11     0.07800 0.20500     0.540
## 25                    Glycerol; 57     0.07780 0.20600     0.540
## 26                Ethanolamine; 56    -0.07760 0.20700     0.540
## 27             Dodecanoic acid; 54     0.07670 0.21200     0.540
## 28           Myristoleic acid; 65     0.07640 0.21400     0.540
## 29               Nonanoic acid; 67     0.07470 0.22600     0.540
## 30          Fumaric acid, 2TMS; 9     0.07370 0.23100     0.540
## 31                Serine, 3TMS; 14    -0.07110 0.24600     0.540
## 32 3,4-Dihydroxybutanoic acid; 27     0.07070 0.24700     0.540
## 33             Arabinopyranose; 51    -0.06920 0.25700     0.540
## 34                     Ribitol; 71     0.06850 0.26200     0.540
## 35              Arachidic acid; 46     0.06900 0.26200     0.540
## 36      Arachidonic acid, TMS; 24     0.06860 0.26500     0.540
## 37             1,3-Propanediol; 34    -0.06840 0.26700     0.540
## 38          Nonadecanoic acid; 66    -0.06410 0.29800     0.576
## 39           alpha-Tocopherol; 26    -0.06360 0.29900     0.576
## 40               Decanoic acid; 52     0.06130 0.31700     0.595
## 41                    Tyrosine; 75    -0.05840 0.34200     0.601
## 42         Glyceryl-glycoside; 59    -0.05820 0.34300     0.601
## 43         Heptadecanoic acid; 60     0.05820 0.34500     0.601
## 44                1-Dodecanol; 36     0.05240 0.39500     0.665
```

```
## 45             L-5-Oxoproline; 63    -0.05190 0.39900     0.665
## 46              Alanine, 2TMS; 25    -0.04630 0.45100     0.735
## 47          Citric acid, 4TMS; 6     0.04500 0.46300     0.739
## 48              Proline, 2TMS; 21    -0.04200 0.49300     0.771
## 49           Isoleucine, 2TMS; 18    -0.03800 0.53500     0.807
## 50                Bisphenol A; 48    -0.03780 0.54000     0.807
## 51        Benzeneacetic acid; 47     0.03690 0.54900     0.807
## 52                Campesterol; 49     0.03220 0.59900     0.859
## 53             1-Monopalmitin; 37     0.03160 0.60700     0.859
## 54           Tridecanoic acid; 74    -0.02940 0.63300     0.873
## 55       3-Indoleacetic acid; 40    -0.02870 0.64000     0.873
## 56          Pyroglutamic acid; 69    -0.02640 0.66800     0.895
## 57    4-Hydroxybutanoic acid; 43     0.02540 0.68100     0.895
## 58     Eicosapentaenoic acid; 55    -0.02160 0.72400     0.922
## 59              Ribonic acid; 72     0.02130 0.72700     0.922
## 60 alpha-ketoglutaric acid, TMS M    -0.02060 0.73800     0.922
## 61          Myo inositol 6TMS; 1     0.01750 0.77400     0.942
## 62         Glutamic acid, 3TMS; 8     0.01680 0.78500     0.942
## 63             Octanoic acid; 68    -0.01630 0.79100     0.942
## 64           Cholesterol, TMS; 23     0.01500 0.80600     0.945
## 65              Glycine, 3TMS; 17     0.01380 0.82300     0.949
## 66       4-Deoxytetronic acid; 33     0.01240 0.84000     0.954
## 67               Pyruvic acid; 31    -0.00904 0.88300     0.954
## 68             Hydroxyproline; 64    -0.00891 0.88500     0.954
## 69                   Ribitol; 70    -0.00888 0.88500     0.954
## 70              Glyceric acid; 30    -0.00715 0.90700     0.954
## 71          Aminomalonic acid; 45    -0.00684 0.91100     0.954
## 72       2-Palmitoylglycerol; 39    -0.00628 0.91900     0.954
## 73 2,4-Dihydroxybutanoic acid; 28     0.00548 0.92800     0.954
## 74                Creatinine; 50     0.00234 0.97000     0.983
## 75          Linoleic acid, TMS; 4    -0.00042 0.99500     0.995
```

###### 5.1.3.2 Forest Plot of Model Coefficients

```
## Warning: Ignoring unknown aesthetics: x

## Warning: Ignoring unknown aesthetics: x

## Warning: Ignoring unknown aesthetics: x

## Warning: Ignoring unknown aesthetics: x

## Warning: Ignoring unknown aesthetics: x

## Warning: Ignoring unknown aesthetics: x

## Warning: Ignoring unknown aesthetics: x

## Warning: Ignoring unknown aesthetics: x

## NULL
```

#### 5.2 Deep Breathing (E_I)

##### 5.2.1 Crude Model

```
## [1] "Fitting models:"
## [1] "~ E_I"
## [1] ""
```

###### 5.2.1.1 Tables of Model Coefficients

```
## [1] ""
## [1] "Table: E_I"
## [1] "  (from model: "
## [1] "   ~ E_I)"
## [1] ""
##                                Name Coefficient  P.Value adj.P.Val
## 1  2,4-Dihydroxybutanoic acid; 28    -0.24900 2.85e-05  0.000974
## 2  3,4-Dihydroxybutanoic acid; 27    -0.24700 3.35e-05  0.000974
## 3               Ribonic acid; 72    -0.24500 3.90e-05  0.000974
## 4            Myo inositol 6TMS; 1    -0.22800 1.23e-04  0.002300
## 5            Citric acid, 4TMS; 6    -0.21300 3.46e-04  0.005190
## 6                  Creatinine; 50    -0.20100 7.28e-04  0.009100
## 7        4-Deoxytetronic acid; 32    -0.19600 9.75e-04  0.010400
## 8  4-Hydroxybenzeneacetic acid; 4    -0.18100 2.37e-03  0.022200
## 9                     Ribitol; 71    -0.16800 4.77e-03  0.039800
## 10         Glyceryl-glycoside; 59    -0.16000 7.00e-03  0.052500
## 11 4-Hydroxyphenyllactic acid; 44    -0.13400 2.44e-02  0.166000
## 12           Tridecanoic acid; 74    -0.12700 3.33e-02  0.208000
## 13          Fumaric acid, 2TMS; 9    -0.12300 3.87e-02  0.223000
## 14                   Glycerol; 58    -0.11900 4.61e-02  0.247000
## 15                    Ribitol; 70    -0.11200 5.97e-02  0.298000
## 16              1-Monopalmitin; 37    -0.10800 6.87e-02  0.322000
## 17         Succinic acid, 2TMS; 7    -0.10600 7.49e-02  0.330000
## 18      Arachidonic acid, TMS; 24    -0.10400 7.92e-02  0.330000
## 19           Methionine, 2TMS; 16     0.09770 1.01e-01  0.397000
## 20           alpha-Tocopherol; 26    -0.09360 1.15e-01  0.421000
## 21           Myristoleic acid; 65    -0.09300 1.18e-01  0.421000
## 22             Oleic acid, TMS; 3    -0.08980 1.31e-01  0.447000
## 23                Campesterol; 49    -0.07980 1.80e-01  0.540000
## 24             Hydroxyproline; 64    -0.07970 1.80e-01  0.540000
## 25          Linoleic acid, TMS; 4    -0.07930 1.83e-01  0.540000
## 26             Tartronic acid; 73     0.07850 1.87e-01  0.540000
## 27              Proline, 2TMS; 21    -0.07300 2.20e-01  0.611000
## 28           Malic acid, 3TMS; 11    -0.06930 2.44e-01  0.651000
## 29              Octanoic acid; 68     0.06780 2.54e-01  0.651000
## 30         Heptadecanoic acid; 61    -0.06670 2.62e-01  0.651000
## 31     3-Indolepropionic acid; 41     0.06460 2.77e-01  0.651000
## 32     4-Hydroxybutanoic acid; 43    -0.06170 2.99e-01  0.651000
## 33            Arabinopyranose; 51    -0.06090 3.06e-01  0.651000
## 34             L-5-Oxoproline; 63    -0.06020 3.12e-01  0.651000
## 35          Palmitic acid, TMS; 5    -0.05930 3.19e-01  0.651000
## 36 2-Hydroxybutyric acid, 2TMS; 2    -0.05910 3.21e-01  0.651000
## 37              Glycine, 3TMS; 17    -0.05900 3.21e-01  0.651000
## 38        Glutamic acid, 3TMS; 8    -0.05800 3.30e-01  0.651000
## 39                1-Dodecanol; 36    -0.05610 3.45e-01  0.664000
## 40               Serine, 3TMS; 14     0.05430 3.61e-01  0.670000
## 41 3-Hydroxybutyric acid, 2TMS; 1    -0.05370 3.67e-01  0.670000
## 42          Pyroglutamic acid; 69    -0.04990 4.01e-01  0.677000
## 43                   Tyrosine; 75     0.04890 4.11e-01  0.677000
## 44             1,3-Propanediol; 34     0.04870 4.13e-01  0.677000
## 45         Benzeneacetic acid; 47    -0.04860 4.14e-01  0.677000
## 46       Docosahexaenoic acid; 53     0.04840 4.15e-01  0.677000
```

```
## 47                Bisphenol A; 48    -0.04340 4.66e-01  0.743000
## 48      4-Deoxytetronic acid; 33    -0.03850 5.18e-01  0.809000
## 49           Threonine, 3TMS; 12    -0.03720 5.32e-01  0.815000
## 50              Valine, 2TMS; 20     0.03550 5.50e-01  0.824000
## 51              Ethanolamine; 56     0.03460 5.61e-01  0.824000
## 52          Cholesterol, TMS; 23    -0.03170 5.94e-01  0.852000
## 53         Nonadecanoic acid; 66     0.03100 6.03e-01  0.852000
## 54           Dodecanoic acid; 54    -0.02980 6.16e-01  0.852000
## 55  2-hydroxy Isovaleric acid; 38    -0.02730 6.46e-01  0.852000
## 56            Arachidic acid; 46     0.02660 6.55e-01  0.852000
## 57          Stearic acid, TMS; 2    -0.02580 6.64e-01  0.852000
## 58             Nonanoic acid; 67    -0.02460 6.79e-01  0.852000
## 59             Glyceric acid; 30     0.02410 6.85e-01  0.852000
## 60     Eicosapentaenoic acid; 55     0.02220 7.09e-01  0.852000
## 61        Phenylalanine, 2TMS; 13     0.02200 7.11e-01  0.852000
## 62               Lactic acid; 29     0.02160 7.16e-01  0.852000
## 63       3-Indoleacetic acid; 40    -0.02040 7.32e-01  0.852000
## 64             Leucine, 2TMS; 19    -0.02010 7.35e-01  0.852000
## 65                  Glycerol; 57     0.01980 7.39e-01  0.852000
## 66       2-Palmitoylglycerol; 39    -0.01560 7.93e-01  0.895000
## 67              Pyruvic acid; 31     0.01510 8.00e-01  0.895000
## 68        11-Eicosenoic acid; 35     0.01250 8.33e-01  0.911000
## 69 alpha-ketoglutaric acid, TMS M    -0.01140 8.48e-01  0.911000
## 70              Hydroxylamine; 62     0.01120 8.50e-01  0.911000
## 71             Decanoic acid; 52    -0.01020 8.64e-01  0.912000
## 72        Heptadecanoic acid; 60     0.00782 8.95e-01  0.927000
## 73             Alanine, 2TMS; 25     0.00729 9.02e-01  0.927000
## 74         Aminomalonic acid; 45     0.00393 9.47e-01  0.960000
## 75          Isoleucine, 2TMS; 18    -0.00151 9.80e-01  0.980000
```

###### 5.2.1.2 Forest Plot of Model Coefficients

```
## Warning: Ignoring unknown aesthetics: x
```

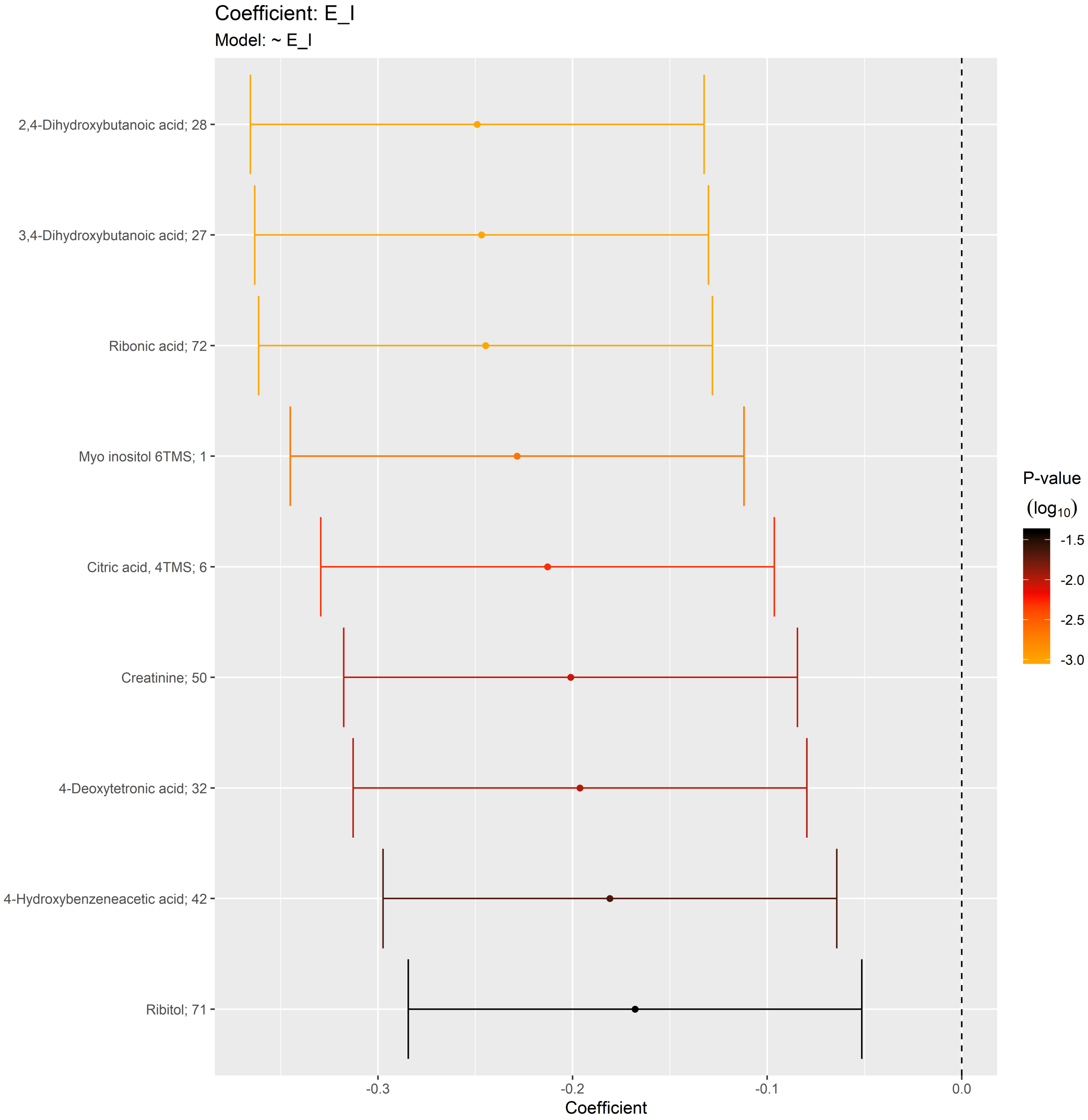

##### 5.2.2 Adjusted Model

```
## [1] "Fitting models:"
## [1] "~ E_I + Age + bmi + Blood_glucose + Duration_DM + Gender + Hba1c_baseline + log_Blood_TGA + Smok
## [1] ""
```

###### 5.2.2.1 Tables of Model Coefficients

```
## [1] ""
## [1] "Table: E_I"
## [1] "  (from model: "
## [1] "    ~ E_I + Age + bmi + Blood_glucose + Duration_DM + Gender"
## [1] "      + Hba1c_baseline + log_Blood_TGA + Smoking + Statin +"
## [1] "      Total_cholesterol)"
## [1] ""
##                               Name Coefficient  P.Value adj.P.Val
## 1               Ribonic acid; 72   -0.219000 0.000682    0.0486
## 2            Citric acid, 4TMS; 6   -0.207000 0.001300    0.0486
## 3  2,4-Dihydroxybutanoic acid; 28   -0.198000 0.002100    0.0526
## 4                 Creatinine; 50   -0.176000 0.006270    0.1170
## 5            Myo inositol 6TMS; 1   -0.160000 0.012700    0.1500
## 6            Tridecanoic acid; 74   -0.159000 0.013500    0.1500
## 7        4-Deoxytetronic acid; 32   -0.158000 0.014000    0.1500
## 8  3,4-Dihydroxybutanoic acid; 27   -0.153000 0.017400    0.1630
## 9           Linoleic acid, TMS; 4   -0.123000 0.055200    0.4600
## 10     4-Hydroxybutanoic acid; 43   -0.115000 0.074200    0.5160
## 11         Glyceryl-glycoside; 59   -0.114000 0.075700    0.5160
## 12            Threonine, 3TMS; 12   -0.108000 0.092200    0.5560
## 13             1-Monopalmitin; 37   -0.107000 0.096700    0.5560
## 14 4-Hydroxybenzeneacetic acid; 4   -0.102000 0.113000    0.5560
## 15         Succinic acid, 2TMS; 7   -0.101000 0.115000    0.5560
## 16                Bisphenol A; 48   -0.099400 0.122000    0.5560
## 17             Hydroxyproline; 64   -0.096600 0.133000    0.5560
## 18                   Glycerol; 58   -0.094100 0.144000    0.5560
## 19        Heptadecanoic acid; 61   -0.090300 0.161000    0.5560
## 20           Methionine, 2TMS; 16    0.088400 0.170000    0.5560
## 21             Tartronic acid; 73    0.085200 0.186000    0.5560
## 22      Docosahexaenoic acid; 53    0.082900 0.198000    0.5560
## 23                1-Dodecanol; 36   -0.081600 0.205000    0.5560
## 24           Myristoleic acid; 65   -0.081300 0.207000    0.5560
## 25             Oleic acid, TMS; 3   -0.081200 0.207000    0.5560
## 26                   Tyrosine; 75    0.080600 0.211000    0.5560
## 27           Cholesterol, TMS; 23   -0.079900 0.215000    0.5560
## 28              Nonanoic acid; 67   -0.078900 0.220000    0.5560
## 29     Eicosapentaenoic acid; 55    0.078500 0.223000    0.5560
## 30     Arachidonic acid, TMS; 24   -0.078200 0.225000    0.5560
## 31          Fumaric acid, 2TMS; 9   -0.077300 0.230000    0.5560
## 32              Glycine, 3TMS; 17   -0.074000 0.251000    0.5790
## 33          Palmitic acid, TMS; 5   -0.073200 0.255000    0.5790
## 34 4-Hydroxyphenyllactic acid; 44   -0.072100 0.262000    0.5790
## 35             L-5-Oxoproline; 63   -0.070900 0.271000    0.5800
## 36                    Ribitol; 71   -0.067900 0.292000    0.6070
## 37           Malic acid, 3TMS; 11   -0.061900 0.336000    0.6820
## 38              Proline, 2TMS; 21   -0.060600 0.347000    0.6840
## 39  2-hydroxy Isovaleric acid; 38   -0.058900 0.360000    0.6920
## 40         11-Eicosenoic acid; 35    0.057000 0.376000    0.7040
## 41              Hydroxylamine; 62    0.055900 0.385000    0.7040
## 42                Campesterol; 49   -0.050800 0.430000    0.7680
## 43              Octanoic acid; 68    0.046600 0.469000    0.7900
## 44            Arabinopyranose; 51   -0.046200 0.473000    0.7900
```

```
## 45              Pyruvic acid; 31    0.045100 0.483000    0.7900
## 46                   Ribitol; 70   -0.043200 0.503000    0.7900
## 47        3-Indoleacetic acid; 40    0.040300 0.532000    0.7900
## 48          Aminomalonic acid; 45   -0.040100 0.534000    0.7900
## 49         Glutamic acid, 3TMS; 8   -0.039600 0.539000    0.7900
## 50       4-Deoxytetronic acid; 33   -0.039500 0.539000    0.7900
## 51              Alanine, 2TMS; 25    0.039000 0.544000    0.7900
## 52 2-Hydroxybutyric acid, 2TMS; 2   -0.037300 0.562000    0.7900
## 53     3-Indolepropionic acid; 41    0.037300 0.562000    0.7900
## 54         Benzeneacetic acid; 47   -0.036700 0.569000    0.7900
## 55 alpha-ketoglutaric acid, TMS M    0.034500 0.592000    0.7950
## 56                Lactic acid; 29    0.034300 0.594000    0.7950
## 57 3-Hydroxybutyric acid, 2TMS; 1   -0.033300 0.605000    0.7960
## 58              Decanoic acid; 52   -0.031300 0.627000    0.8100
## 59          Nonadecanoic acid; 66    0.028800 0.655000    0.8240
## 60          Pyroglutamic acid; 69   -0.028400 0.659000    0.8240
## 61            1,3-Propanediol; 34    0.023700 0.713000    0.8760
## 62           Stearic acid, TMS; 2   -0.021000 0.744000    0.9000
## 63         Heptadecanoic acid; 60   -0.016800 0.794000    0.9270
## 64              Glyceric acid; 30    0.015800 0.806000    0.9270
## 65               Valine, 2TMS; 20    0.015000 0.816000    0.9270
## 66             Arachidic acid; 46    0.014300 0.824000    0.9270
## 67                   Glycerol; 57    0.014000 0.828000    0.9270
## 68           alpha-Tocopherol; 26   -0.012700 0.843000    0.9300
## 69           Isoleucine, 2TMS; 18    0.010200 0.874000    0.9350
## 70              Leucine, 2TMS; 19   -0.009470 0.883000    0.9350
## 71                Ethanolamine; 56    0.009310 0.885000    0.9350
## 72            Dodecanoic acid; 54   -0.007530 0.907000    0.9450
## 73        Phenylalanine, 2TMS; 13   -0.001640 0.980000    0.9970
## 74               Serine, 3TMS; 14    0.001230 0.985000    0.9970
## 75        2-Palmitoylglycerol; 39    0.000248 0.997000    0.9970
```

###### 5.2.2.2 Forest Plot of Model Coefficients

```
## Warning: Ignoring unknown aesthetics: x

## Warning: Ignoring unknown aesthetics: x

## Warning: Ignoring unknown aesthetics: x

## Warning: Ignoring unknown aesthetics: x

## Warning: Ignoring unknown aesthetics: x

## Warning: Ignoring unknown aesthetics: x

## Warning: Ignoring unknown aesthetics: x

## Warning: Ignoring unknown aesthetics: x
```

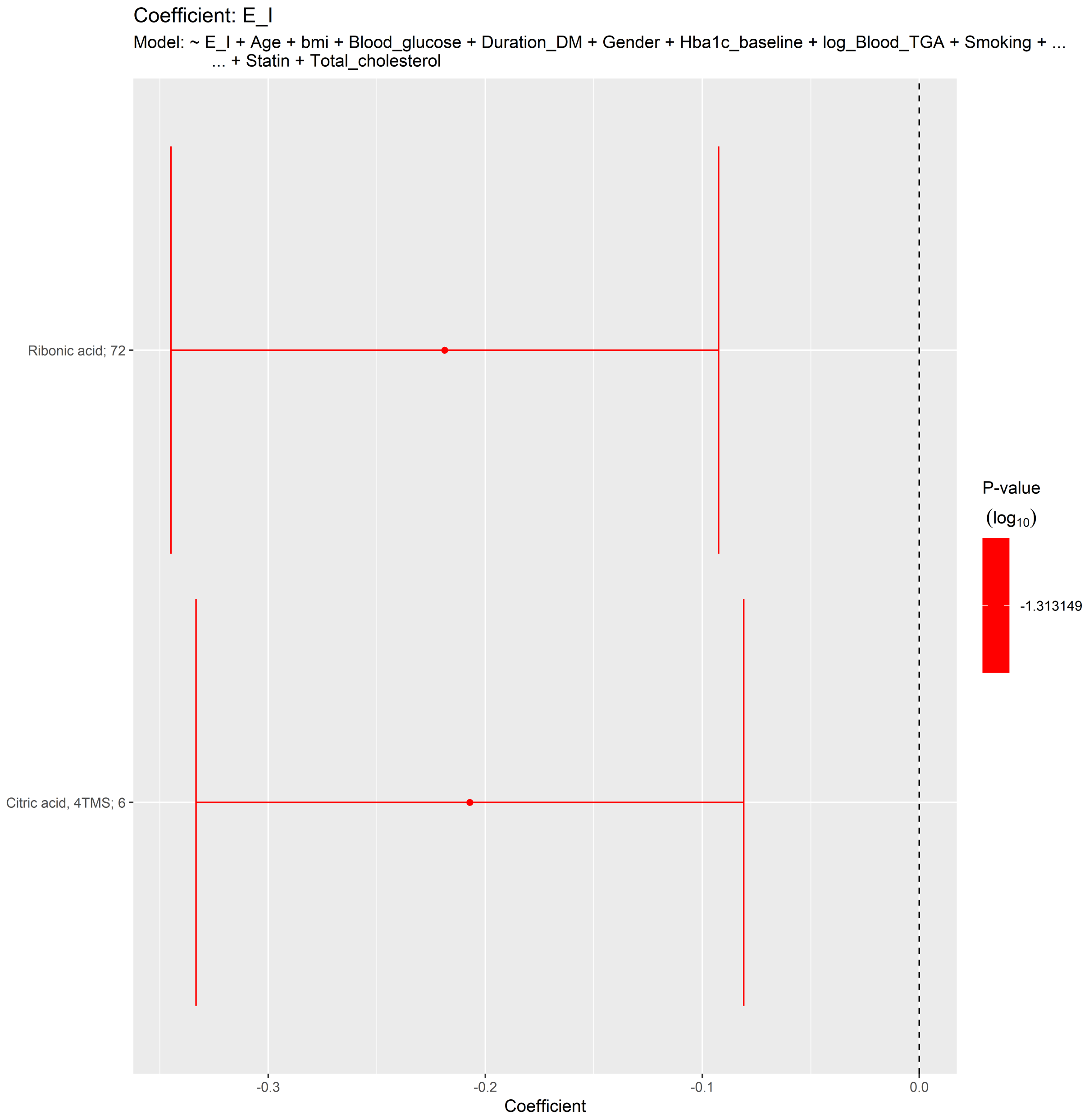

Coefficient: E_I
Model: ~ E_I + Age + bmi + Blood_glucose + Duration_DM + Gender + Hba1c_baseline + log_Blood_TGA + Smoking + ...
... + Statin + Total_cholesterol
Ribonic acid; 72
Citric acid, 4TMS; 6
P-value
(log10)
-1.313149
-0.3
-0.2
-0.1
0.0
Coefficient

##### 5.2.3 Fully-Adjusted Model

```
## [1] "Fitting models:"
## [1] "~ E_I + Age + bmi + Blood_glucose + Duration_DM + Gender + Hba1c_baseline + log_Blood_TGA + Smok
## [1] ""
```

###### 5.2.3.1 Tables of Model Coefficients

```
## [1] ""
## [1] "Table: E_I"
## [1] "  (from model: "
## [1] "   ~ E_I + Age + bmi + Blood_glucose + Duration_DM + Gender"
## [1] "     + Hba1c_baseline + log_Blood_TGA + Smoking + Statin +"
## [1] "     Total_cholesterol + egfr)"
## [1] ""
##                               Name Coefficient P.Value adj.P.Val
## 1             Citric acid, 4TMS; 6    -0.15900  0.0143     0.526
## 2             Tridecanoic acid; 74    -0.15900  0.0149     0.526
## 3            Linoleic acid, TMS; 4    -0.14700  0.0240     0.526
## 4              Threonine, 3TMS; 12    -0.13800  0.0339     0.526
## 5                 Ribonic acid; 72    -0.13200  0.0402     0.526
## 6  2-Hydroxybutyric acid, 2TMS; 2    -0.11800  0.0674     0.526
## 7            Cholesterol, TMS; 23    -0.11300  0.0790     0.526
## 8                 Bisphenol A; 48    -0.11400  0.0827     0.526
## 9   2-hydroxy Isovaleric acid; 38    -0.11000  0.0904     0.526
## 10         Heptadecanoic acid; 61    -0.10900  0.0942     0.526
## 11           Palmitic acid, TMS; 5    -0.10700  0.0979     0.526
## 12              1-Monopalmitin; 37    -0.10700  0.1000     0.526
## 13        3-Indoleacetic acid; 40     0.10600  0.1030     0.526
## 14     4-Hydroxybutanoic acid; 43    -0.10600  0.1050     0.526
## 15      Arachidonic acid, TMS; 24    -0.10600  0.1050     0.526
## 16         Succinic acid, 2TMS; 7    -0.10200  0.1190     0.560
## 17 2,4-Dihydroxybutanoic acid; 28    -0.09640  0.1330     0.565
## 18              Oleic acid, TMS; 3    -0.09700  0.1360     0.565
## 19              Tartronic acid; 73     0.08800  0.1730     0.635
## 20                  Creatinine; 50    -0.08790  0.1750     0.635
## 21       4-Deoxytetronic acid; 32    -0.08560  0.1870     0.635
## 22           Myristoleic acid; 65    -0.08410  0.1960     0.635
## 23                 1-Dodecanol; 36    -0.08240  0.2080     0.635
## 24              L-5-Oxoproline; 63    -0.08210  0.2080     0.635
## 25               Hydroxylamine; 62     0.08150  0.2120     0.635
## 26         Glutamic acid, 3TMS; 8    -0.07490  0.2480     0.691
## 27               Nonanoic acid; 67    -0.07550  0.2490     0.691
## 28         11-Eicosenoic acid; 35     0.07090  0.2770     0.741
## 29       Docosahexaenoic acid; 53     0.06850  0.2900     0.750
## 30            Stearic acid, TMS; 2    -0.06340  0.3300     0.782
## 31      Eicosapentaenoic acid; 55     0.06280  0.3320     0.782
## 32                Serine, 3TMS; 14    -0.05930  0.3610     0.782
## 33 3,4-Dihydroxybutanoic acid; 27    -0.05880  0.3610     0.782
## 34                 Campesterol; 49    -0.05890  0.3640     0.782
## 35                    Glycerol; 58    -0.05900  0.3660     0.782
## 36         Glyceryl-glycoside; 59    -0.05480  0.3990     0.782
## 37            Isoleucine, 2TMS; 18    -0.05220  0.4190     0.782
## 38            Myo inositol 6TMS; 1    -0.05160  0.4210     0.782
## 39               Alanine, 2TMS; 25     0.05100  0.4330     0.782
## 40               Proline, 2TMS; 21    -0.05000  0.4410     0.782
## 41               Leucine, 2TMS; 19    -0.04950  0.4460     0.782
## 42            Methionine, 2TMS; 16     0.04930  0.4480     0.782
## 43                     Ribitol; 71     0.04610  0.4720     0.782
## 44                Pyruvic acid; 31     0.04660  0.4740     0.782
```

```
## 45                  Tyrosine; 75     0.04490  0.4900     0.782
## 46          Fumaric acid, 2TMS; 9    -0.04300  0.5100     0.782
## 47              Decanoic acid; 52    -0.04280  0.5100     0.782
## 48              Glycine, 3TMS; 17    -0.04250  0.5140     0.782
## 49               Valine, 2TMS; 20    -0.04150  0.5220     0.782
## 50       4-Deoxytetronic acid; 33     0.04120  0.5250     0.782
## 51            Arabinopyranose; 51    -0.03960  0.5380     0.782
## 52 alpha-ketoglutaric acid, TMS M     0.03970  0.5420     0.782
## 53          Aminomalonic acid; 45    -0.03810  0.5560     0.782
## 54 3-Hydroxybutyric acid, 2TMS; 1    -0.03730  0.5670     0.782
## 55     3-Indolepropionic acid; 41     0.03670  0.5740     0.782
## 56             Hydroxyproline; 64    -0.03040  0.6410     0.858
## 57          Benzeneacetic acid; 47    -0.02550  0.6960     0.873
## 58            1,3-Propanediol; 34     0.02500  0.7020     0.873
## 59 4-Hydroxyphenyllactic acid; 44    -0.02370  0.7160     0.873
## 60            Malic acid, 3TMS; 11    -0.02340  0.7200     0.873
## 61             Dodecanoic acid; 54    -0.02340  0.7200     0.873
## 62            alpha-Tocopherol; 26    -0.02210  0.7330     0.873
## 63                    Glycerol; 57    -0.02050  0.7530     0.873
## 64                     Ribitol; 70    -0.01980  0.7610     0.873
## 65          Heptadecanoic acid; 60    -0.01930  0.7670     0.873
## 66           Pyroglutamic acid; 69     0.01920  0.7680     0.873
## 67                 Lactic acid; 29     0.01710  0.7930     0.888
## 68               Glyceric acid; 30    -0.01550  0.8110     0.891
## 69           Nonadecanoic acid; 66     0.01480  0.8200     0.891
## 70                Ethanolamine; 56    -0.01380  0.8320     0.892
## 71 4-Hydroxybenzeneacetic acid; 4    -0.00965  0.8810     0.925
## 72         2-Palmitoylglycerol; 39    -0.00874  0.8940     0.925
## 73               Octanoic acid; 68     0.00815  0.9000     0.925
## 74         Phenylalanine, 2TMS; 13     0.00370  0.9550     0.956
## 75              Arachidic acid; 46     0.00359  0.9560     0.956
```

###### 5.2.3.2 Forest Plot of Model Coefficients

```
## Warning: Ignoring unknown aesthetics: x

## Warning: Ignoring unknown aesthetics: x

## Warning: Ignoring unknown aesthetics: x

## Warning: Ignoring unknown aesthetics: x

## Warning: Ignoring unknown aesthetics: x

## Warning: Ignoring unknown aesthetics: x

## Warning: Ignoring unknown aesthetics: x

## NULL
```

#### 5.3 Lying to Standing Test (lig_staa)

##### 5.3.1 Crude Model

```
## [1] "Fitting models:"
## [1] "~ lig_staa"
## [1] ""
```

###### 5.3.1.1 Tables of Model Coefficients

```
## [1] ""
## [1] "Table: lig_staa"
## [1] "  (from model: "
## [1] "   ~ lig_staa)"
## [1] ""
##                               Name Coefficient  P.Value adj.P.Val
## 1  3,4-Dihydroxybutanoic acid; 27   -0.206000 0.000663    0.0394
## 2  2,4-Dihydroxybutanoic acid; 28   -0.198000 0.001050    0.0394
## 3                 Creatinine; 50   -0.155000 0.010200    0.1670
## 4            Myo inositol 6TMS; 1   -0.153000 0.011100    0.1670
## 5               Ethanolamine; 56    0.153000 0.011200    0.1670
## 6  4-Hydroxybenzeneacetic acid; 4   -0.148000 0.014400    0.1800
## 7                    Ribitol; 71   -0.117000 0.052500    0.5620
## 8       Arachidonic acid, TMS; 24   -0.109000 0.070200    0.6430
## 9               Ribonic acid; 72   -0.107000 0.077200    0.6430
## 10             Nonanoic acid; 67   -0.100000 0.097500    0.7300
## 11            L-5-Oxoproline; 63   -0.097300 0.107000    0.7300
## 12             Hydroxylamine; 62    0.088500 0.143000    0.8650
## 13                  Glycerol; 58   -0.083500 0.167000    0.8650
## 14             Octanoic acid; 68    0.080900 0.180000    0.8650
## 15          Isoleucine, 2TMS; 18    0.077100 0.202000    0.8650
## 16          Methionine, 2TMS; 16    0.076600 0.205000    0.8650
## 17             Glycine, 3TMS; 17   -0.075500 0.211000    0.8650
## 18         Pyroglutamic acid; 69   -0.071200 0.238000    0.8650
## 19                  Glycerol; 57   -0.068900 0.254000    0.8650
## 20      4-Deoxytetronic acid; 32   -0.068500 0.257000    0.8650
## 21          Myristoleic acid; 65   -0.066200 0.273000    0.8650
## 22        2-Palmitoylglycerol; 39    0.064900 0.282000    0.8650
## 23            Oleic acid, TMS; 3   -0.061000 0.313000    0.8650
## 24         Nonadecanoic acid; 66    0.060900 0.313000    0.8650
## 25            Arachidic acid; 46    0.059000 0.328000    0.8650
## 26               Valine, 2TMS; 20    0.057400 0.342000    0.8650
## 27             Glyceric acid; 30    0.056900 0.346000    0.8650
## 28         Succinic acid, 2TMS; 7   -0.056900 0.346000    0.8650
## 29 4-Hydroxyphenyllactic acid; 44   -0.055300 0.360000    0.8650
## 30         Glyceryl-glycoside; 59   -0.054700 0.365000    0.8650
## 31 alpha-ketoglutaric acid, TMS M   -0.051900 0.390000    0.8650
## 32             Hydroxyproline; 64   -0.050900 0.399000    0.8650
## 33          Fumaric acid, 2TMS; 9   -0.049900 0.409000    0.8650
## 34          Aminomalonic acid; 45   -0.047100 0.436000    0.8650
## 35         Glutamic acid, 3TMS; 8   -0.045800 0.448000    0.8650
## 36               Serine, 3TMS; 14    0.042700 0.480000    0.8650
## 37     3-Indolepropionic acid; 41    0.042600 0.481000    0.8650
## 38                   Ribitol; 70   -0.042100 0.486000    0.8650
## 39          Linoleic acid, TMS; 4    0.041400 0.493000    0.8650
## 40       4-Deoxytetronic acid; 33   -0.040400 0.503000    0.8650
## 41       Docosahexaenoic acid; 53    0.039800 0.510000    0.8650
## 42           alpha-Tocopherol; 26    0.038300 0.526000    0.8650
## 43           Citric acid, 4TMS; 6   -0.036200 0.548000    0.8650
## 44           Tridecanoic acid; 74   -0.035800 0.554000    0.8650
## 45              Leucine, 2TMS; 19    0.035700 0.554000    0.8650
## 46 3-Hydroxybutyric acid, 2TMS; 1   -0.033800 0.576000    0.8650
```

```
## 47            1-Monopalmitin; 37   -0.033400 0.580000    0.8650
## 48  2-hydroxy Isovaleric acid; 38   -0.031200 0.606000    0.8650
## 49              Proline, 2TMS; 21   -0.030500 0.614000    0.8650
## 50              Alanine, 2TMS; 25   -0.029800 0.621000    0.8650
## 51 2-Hydroxybutyric acid, 2TMS; 2    0.029300 0.628000    0.8650
## 52                1-Dodecanol; 36   -0.028900 0.633000    0.8650
## 53               Pyruvic acid; 31   -0.028000 0.643000    0.8650
## 54          Benzeneacetic acid; 47   -0.027700 0.646000    0.8650
## 55        Phenylalanine, 2TMS; 13    0.027500 0.649000    0.8650
## 56      Eicosapentaenoic acid; 55    0.026400 0.662000    0.8650
## 57        3-Indoleacetic acid; 40   -0.026100 0.665000    0.8650
## 58             Tartronic acid; 73    0.025000 0.679000    0.8650
## 59                Lactic acid; 29    0.024700 0.683000    0.8650
## 60                Bisphenol A; 48    0.023900 0.692000    0.8650
## 61                   Tyrosine; 75   -0.022600 0.709000    0.8710
## 62            Dodecanoic acid; 54   -0.018900 0.755000    0.9130
## 63           Cholesterol, TMS; 23   -0.017800 0.768000    0.9150
## 64           Malic acid, 3TMS; 11    0.015200 0.801000    0.9220
## 65         Heptadecanoic acid; 60   -0.014500 0.810000    0.9220
## 66            Threonine, 3TMS; 12   -0.014200 0.813000    0.9220
## 67                Campesterol; 49   -0.012700 0.834000    0.9220
## 68          Palmitic acid, TMS; 5   -0.012500 0.836000    0.9220
## 69              Decanoic acid; 52   -0.010700 0.859000    0.9340
## 70         11-Eicosenoic acid; 35   -0.008240 0.891000    0.9550
## 71           1,3-Propanediol; 34    0.005340 0.930000    0.9710
## 72            Arabinopyranose; 51   -0.005140 0.932000    0.9710
## 73           Stearic acid, TMS; 2    0.003060 0.960000    0.9860
## 74         Heptadecanoic acid; 61    0.001390 0.982000    0.9880
## 75     4-Hydroxybutanoic acid; 43   -0.000927 0.988000    0.9880
```

###### 5.3.1.2 Forest Plot of Model Coefficients

```
## Warning: Ignoring unknown aesthetics: x
```

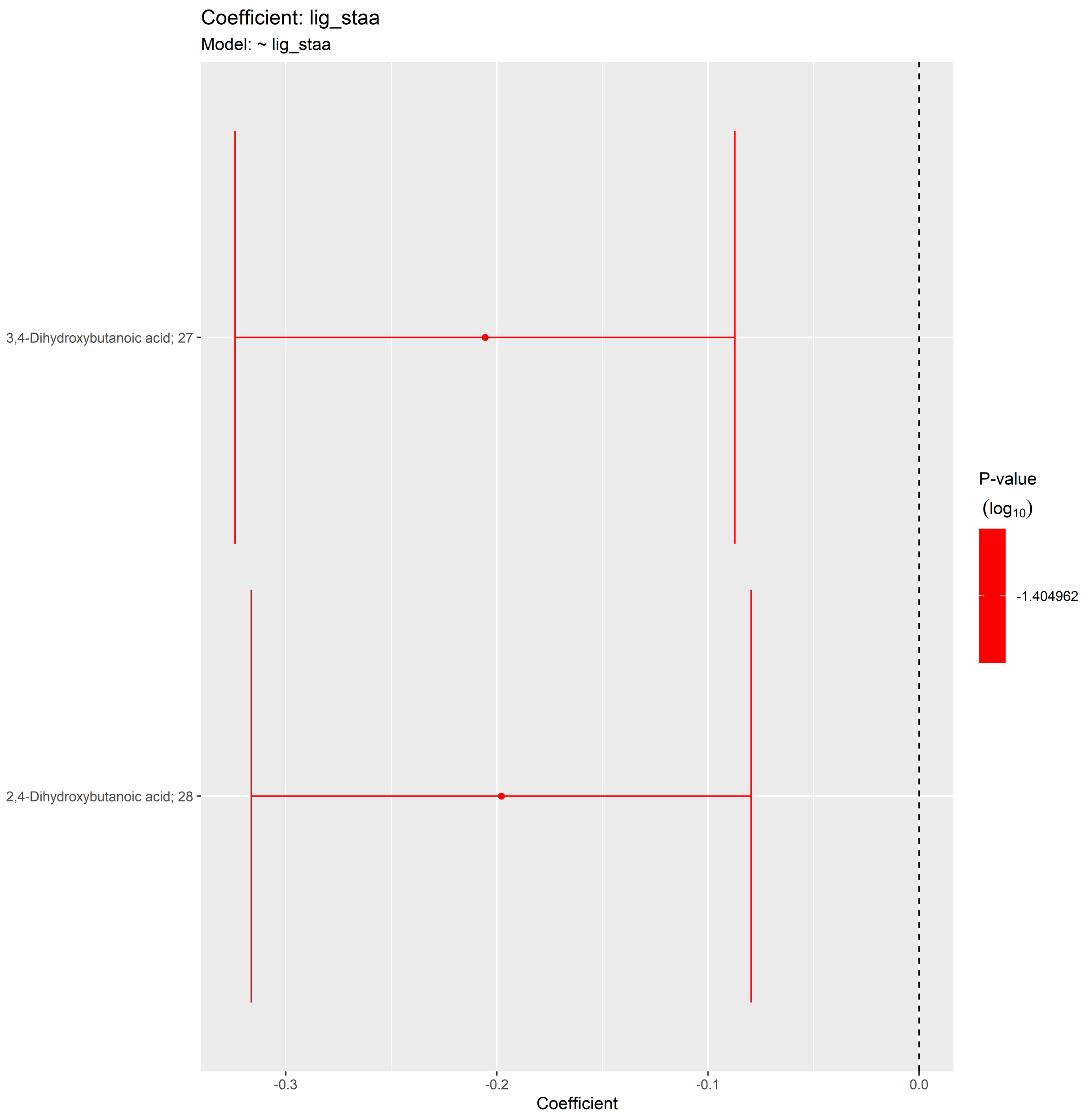

##### 5.3.2 Adjusted Model

```
## [1] "Fitting models:"
## [1] "~ lig_staa + Age + bmi + Blood_glucose + Duration_DM + Gender + Hba1c_baseline + log_Blood_TGA +
## [1] ""
```

###### 5.3.2.1 Tables of Model Coefficients

```
## [1] ""
## [1] "Table: lig_staa"
## [1] "  (from model: "
## [1] "   ~ lig_staa + Age + bmi + Blood_glucose + Duration_DM +"
## [1] "     Gender + Hba1c_baseline + log_Blood_TGA + Smoking + Statin +"
## [1] "     Total_cholesterol)"
## [1] ""
##                               Name Coefficient P.Value adj.P.Val
## 1  2,4-Dihydroxybutanoic acid; 28   -0.147000  0.0157     0.469
## 2                Ethanolamine; 56    0.138000  0.0231     0.469
## 3            Myo inositol 6TMS; 1   -0.135000  0.0259     0.469
## 4                  Creatinine; 50   -0.135000  0.0261     0.469
## 5               Nonanoic acid; 67   -0.131000  0.0314     0.469
## 6  3,4-Dihydroxybutanoic acid; 27   -0.126000  0.0375     0.469
## 7       Arachidonic acid, TMS; 24   -0.103000  0.0902     0.827
## 8               L-5-Oxoproline; 63   -0.101000  0.0950     0.827
## 9                Hydroxylamine; 62    0.100000  0.0992     0.827
## 10 4-Hydroxybenzeneacetic acid; 4   -0.091800  0.1310     0.980
## 11                Ribonic acid; 72   -0.084400  0.1650     0.980
## 12            Isoleucine, 2TMS; 18    0.082600  0.1740     0.980
## 13                    Glycerol; 58   -0.080500  0.1850     0.980
## 14                    Glycerol; 57   -0.076800  0.2060     0.980
## 15               Glycine, 3TMS; 17   -0.074300  0.2220     0.980
## 16            Methionine, 2TMS; 16    0.072900  0.2300     0.980
## 17         2-Palmitoylglycerol; 39    0.072700  0.2320     0.980
## 18           Nonadecanoic acid; 66    0.065900  0.2780     0.980
## 19            Cholesterol, TMS; 23   -0.063800  0.2940     0.980
## 20                     Ribitol; 71   -0.063700  0.2950     0.980
## 21              Oleic acid, TMS; 3   -0.059800  0.3250     0.980
## 22              Arachidic acid; 46    0.058300  0.3380     0.980
## 23  2-hydroxy Isovaleric acid; 38   -0.058200  0.3380     0.980
## 24              Hydroxyproline; 64   -0.057900  0.3410     0.980
## 25       4-Deoxytetronic acid; 32   -0.053000  0.3830     0.980
## 26            alpha-Tocopherol; 26    0.053000  0.3840     0.980
## 27                 1-Dodecanol; 36   -0.052800  0.3850     0.980
## 28          Succinic acid, 2TMS; 7   -0.051500  0.3970     0.980
## 29           Aminomalonic acid; 45   -0.051200  0.3990     0.980
## 30               Octanoic acid; 68    0.049600  0.4150     0.980
## 31             Threonine, 3TMS; 12   -0.045400  0.4550     0.980
## 32            Myristoleic acid; 65   -0.045300  0.4560     0.980
## 33            Tridecanoic acid; 74   -0.040200  0.5090     0.980
## 34           Pyroglutamic acid; 69   -0.038000  0.5320     0.980
## 35                 Lactic acid; 29    0.037900  0.5330     0.980
## 36        Phenylalanine, 2TMS; 13    0.033500  0.5810     0.980
## 37      Eicosapentaenoic acid; 55    0.032100  0.5970     0.980
## 38               Leucine, 2TMS; 19    0.031900  0.5990     0.980
## 39 alpha-ketoglutaric acid, TMS M   -0.030900  0.6110     0.980
## 40       Docosahexaenoic acid; 53    0.030700  0.6130     0.980
## 41          Glutamic acid, 3TMS; 8   -0.030400  0.6170     0.980
## 42               Alanine, 2TMS; 25   -0.030400  0.6170     0.980
## 43           Fumaric acid, 2TMS; 9   -0.029000  0.6340     0.980
## 44 2-Hydroxybutyric acid, 2TMS; 2    0.028600  0.6380     0.980
```

```
## 45           Citric acid, 4TMS; 6   -0.027200  0.6540     0.980
## 46 4-Hydroxyphenyllactic acid; 44   -0.027100  0.6550     0.980
## 47         Heptadecanoic acid; 60   -0.027100  0.6560     0.980
## 48         Glyceryl-glycoside; 59   -0.027000  0.6570     0.980
## 49              Proline, 2TMS; 21   -0.026800  0.6590     0.980
## 50               Valine, 2TMS; 20    0.025900  0.6700     0.980
## 51          Palmitic acid, TMS; 5   -0.025300  0.6770     0.980
## 52              Glyceric acid; 30    0.024100  0.6910     0.980
## 53       4-Deoxytetronic acid; 33   -0.022700  0.7090     0.980
## 54 3-Hydroxybutyric acid, 2TMS; 1   -0.021900  0.7190     0.980
## 55          Linoleic acid, TMS; 4    0.020400  0.7380     0.980
## 56     3-Indolepropionic acid; 41    0.019100  0.7530     0.980
## 57         Benzeneacetic acid; 47   -0.017200  0.7780     0.980
## 58             1-Monopalmitin; 37   -0.016500  0.7860     0.980
## 59               Serine, 3TMS; 14    0.015000  0.8060     0.980
## 60        3-Indoleacetic acid; 40   -0.014500  0.8110     0.980
## 61               Pyruvic acid; 31   -0.013000  0.8310     0.980
## 62            1,3-Propanediol; 34   -0.011300  0.8520     0.980
## 63           Malic acid, 3TMS; 11    0.011200  0.8530     0.980
## 64             Tartronic acid; 73    0.010500  0.8630     0.980
## 65           Stearic acid, TMS; 2    0.010400  0.8640     0.980
## 66              Decanoic acid; 52    0.007920  0.8960     0.980
## 67         Heptadecanoic acid; 61   -0.006740  0.9120     0.980
## 68            Arabinopyranose; 51   -0.006370  0.9170     0.980
## 69                    Ribitol; 70   -0.006320  0.9170     0.980
## 70     4-Hydroxybutanoic acid; 43   -0.005550  0.9270     0.980
## 71                Bisphenol A; 48    0.005540  0.9270     0.980
## 72        11-Eicosenoic acid; 35    0.002860  0.9620     0.992
## 73            Dodecanoic acid; 54   -0.001370  0.9820     0.992
## 74                   Tyrosine; 75   -0.000878  0.9880     0.992
## 75                Campesterol; 49    0.000615  0.9920     0.992
```

###### 5.3.2.2 Forest Plot of Model Coefficients

```
## Warning: Ignoring unknown aesthetics: x

## Warning: Ignoring unknown aesthetics: x

## Warning: Ignoring unknown aesthetics: x

## Warning: Ignoring unknown aesthetics: x

## Warning: Ignoring unknown aesthetics: x

## NULL
```

##### 5.3.3 Fully-Adjusted Model

```
## [1] "Fitting models:"
## [1] "~ lig_staa + Age + bmi + Blood_glucose + Duration_DM + Gender + Hba1c_baseline + log_Blood_TGA +
## [1] ""
```

###### 5.3.3.1 Tables of Model Coefficients

```
## [1] ""
## [1] "Table: lig_staa"
## [1] "  (from model: "
## [1] "   ~ lig_staa + Age + bmi + Blood_glucose + Duration_DM +"
## [1] "     Gender + Hba1c_baseline + log_Blood_TGA + Smoking + Statin +"
## [1] "     Total_cholesterol + egfr)"
## [1] ""
##                               Name Coefficient P.Value adj.P.Val
## 1               Nonanoic acid; 67   -0.130000  0.0322     0.988
## 2                Ethanolamine; 56    0.128000  0.0344     0.988
## 3        Arachidonic acid, TMS; 24   -0.119000  0.0495     0.988
## 4               Hydroxylamine; 62    0.113000  0.0633     0.988
## 5              L-5-Oxoproline; 63   -0.105000  0.0840     0.988
## 6                    Glycerol; 57   -0.096900  0.1110     0.988
## 7    2-hydroxy Isovaleric acid; 38   -0.085000  0.1610     0.988
## 8            Cholesterol, TMS; 23   -0.084200  0.1620     0.988
## 9  2,4-Dihydroxybutanoic acid; 28   -0.075700  0.2080     0.988
## 10                 Creatinine; 50   -0.076100  0.2080     0.988
## 11        2-Palmitoylglycerol; 39    0.076000  0.2110     0.988
## 12              Oleic acid, TMS; 3   -0.067000  0.2680     0.988
## 13 3,4-Dihydroxybutanoic acid; 27   -0.066300  0.2710     0.988
## 14                   Glycerol; 58   -0.059100  0.3310     0.988
## 15           Myo inositol 6TMS; 1   -0.057900  0.3350     0.988
## 16          Nonadecanoic acid; 66    0.058500  0.3350     0.988
## 17            Threonine, 3TMS; 12   -0.058600  0.3350     0.988
## 18          Glutamic acid, 3TMS; 8   -0.055700  0.3570     0.988
## 19           alpha-Tocopherol; 26    0.054200  0.3700     0.988
## 20              Arachidic acid; 46    0.052000  0.3920     0.988
## 21          Aminomalonic acid; 45   -0.050900  0.4000     0.988
## 22                1-Dodecanol; 36   -0.049500  0.4150     0.988
## 23              Glycine, 3TMS; 17   -0.048700  0.4210     0.988
## 24           Myristoleic acid; 65   -0.048700  0.4210     0.988
## 25         Succinic acid, 2TMS; 7   -0.048200  0.4280     0.988
## 26           Methionine, 2TMS; 16    0.048000  0.4290     0.988
## 27           Isoleucine, 2TMS; 18    0.046900  0.4380     0.988
## 28          Palmitic acid, TMS; 5   -0.043200  0.4760     0.988
## 29        Phenylalanine, 2TMS; 13    0.038000  0.5320     0.988
## 30           Malic acid, 3TMS; 11    0.037700  0.5350     0.988
## 31           Tridecanoic acid; 74   -0.037000  0.5420     0.988
## 32 4-Hydroxybenzeneacetic acid; 4   -0.035400  0.5580     0.988
## 33                   Tyrosine; 75   -0.030900  0.6110     0.988
## 34 alpha-ketoglutaric acid, TMS M   -0.030900  0.6110     0.988
## 35      4-Deoxytetronic acid; 33    0.030400  0.6150     0.988
## 36          Linoleic acid, TMS; 4    0.029500  0.6270     0.988
## 37         Heptadecanoic acid; 60   -0.027300  0.6530     0.988
## 38              Alanine, 2TMS; 25   -0.027200  0.6540     0.988
## 39                Lactic acid; 29    0.026500  0.6630     0.988
## 40               Ribonic acid; 72   -0.024200  0.6880     0.988
## 41     3-Indolepropionic acid; 41    0.021800  0.7190     0.988
## 42               Serine, 3TMS; 14   -0.021400  0.7230     0.988
## 43        3-Indoleacetic acid; 40    0.021200  0.7260     0.988
## 44              Octanoic acid; 68    0.021200  0.7270     0.988
```

```
## 45                Proline, 2TMS; 21   -0.019900  0.7430     0.988
## 46             Citric acid, 4TMS; 6    0.019300  0.7500     0.988
## 47 3-Hydroxybutyric acid, 2TMS; 1   -0.019100  0.7530     0.988
## 48                 Pyruvic acid; 31   -0.018400  0.7610     0.988
## 49      Eicosapentaenoic acid; 55    0.018100  0.7650     0.988
## 50               Hydroxyproline; 64   -0.017700  0.7700     0.988
## 51       Docosahexaenoic acid; 53    0.017400  0.7730     0.988
## 52 2-Hydroxybutyric acid, 2TMS; 2   -0.016700  0.7820     0.988
## 53                      Ribitol; 71    0.015300  0.7980     0.988
## 54          Heptadecanoic acid; 61   -0.015100  0.8040     0.988
## 55          Glyceryl-glycoside; 59    0.013900  0.8190     0.988
## 56             Stearic acid, TMS; 2   -0.012300  0.8400     0.988
## 57              Dodecanoic acid; 54   -0.011700  0.8470     0.988
## 58          Benzeneacetic acid; 47   -0.011300  0.8520     0.988
## 59               Tartronic acid; 73    0.009970  0.8690     0.988
## 60                      Ribitol; 70    0.010000  0.8690     0.988
## 61                 Valine, 2TMS; 20   -0.008790  0.8850     0.988
## 62                Leucine, 2TMS; 19    0.008770  0.8850     0.988
## 63               1-Monopalmitin; 37   -0.008760  0.8850     0.988
## 64              1,3-Propanediol; 34   -0.008710  0.8860     0.988
## 65           Pyroglutamic acid; 69   -0.007380  0.9030     0.988
## 66          11-Eicosenoic acid; 35    0.006590  0.9140     0.988
## 67 4-Hydroxyphenyllactic acid; 44    0.005900  0.9230     0.988
## 68           Fumaric acid, 2TMS; 9   -0.005700  0.9250     0.988
## 69                  Bisphenol A; 48    0.004570  0.9400     0.988
## 70     4-Hydroxybutanoic acid; 43    0.003780  0.9500     0.988
## 71                Glyceric acid; 30    0.003320  0.9560     0.988
## 72                  Campesterol; 49    0.003290  0.9570     0.988
## 73             Arabinopyranose; 51    0.002010  0.9730     0.988
## 74        4-Deoxytetronic acid; 32   -0.001920  0.9750     0.988
## 75                Decanoic acid; 52    0.000269  0.9960     0.996
```

###### 5.3.3.2 Forest Plot of Model Coefficients

```
## Warning: Ignoring unknown aesthetics: x

## Warning: Ignoring unknown aesthetics: x

## Warning: Ignoring unknown aesthetics: x

## Warning: Ignoring unknown aesthetics: x

## Warning: Ignoring unknown aesthetics: x

## Warning: Ignoring unknown aesthetics: x

## NULL
```

#### 5.4 Valsalva Maneuver (Valsal)

##### 5.4.1 Crude Model

```
## [1] "Fitting models:"
## [1] "~ Valsal"
## [1] ""
```

###### 5.4.1.1 Tables of Model Coefficients

```
## [1] ""
## [1] "Table: Valsal"
## [1] "  (from model: "
## [1] "   ~ Valsal)"
## [1] ""
##                               Name Coefficient  P.Value adj.P.Val
## 1  3,4-Dihydroxybutanoic acid; 27   -3.05e-01 2.51e-05   0.00188
## 2  2,4-Dihydroxybutanoic acid; 28   -2.74e-01 1.59e-04   0.00417
## 3  4-Hydroxybenzeneacetic acid; 4   -2.73e-01 1.67e-04   0.00417
## 4                     Glycerol; 58   -2.37e-01 1.06e-03   0.01790
## 5                      Ribitol; 70   -2.35e-01 1.20e-03   0.01790
## 6                      Ribitol; 71   -2.29e-01 1.54e-03   0.01930
## 7              Citric acid, 4TMS; 6   -2.20e-01 2.43e-03   0.02610
## 8              Myo inositol 6TMS; 1   -2.14e-01 3.16e-03   0.02960
## 9                 Ribonic acid; 72   -2.04e-01 4.90e-03   0.04080
## 10                Valine, 2TMS; 20    1.74e-01 1.63e-02   0.12200
## 11       4-Deoxytetronic acid; 32   -1.71e-01 1.84e-02   0.12600
## 12           Fumaric acid, 2TMS; 9   -1.68e-01 2.02e-02   0.12600
## 13              Oleic acid, TMS; 3   -1.57e-01 3.05e-02   0.17200
## 14                Ethanolamine; 56    1.55e-01 3.21e-02   0.17200
## 15              Tartronic acid; 73    1.34e-01 6.51e-02   0.32100
## 16               Glyceric acid; 30    1.31e-01 7.15e-02   0.32100
## 17                  Creatinine; 50   -1.30e-01 7.31e-02   0.32100
## 18            Methionine, 2TMS; 16    1.27e-01 8.06e-02   0.32100
## 19          Benzeneacetic acid; 47   -1.25e-01 8.40e-02   0.32100
## 20               Leucine, 2TMS; 19    1.24e-01 8.69e-02   0.32100
## 21            Isoleucine, 2TMS; 18    1.23e-01 8.99e-02   0.32100
## 22               Alanine, 2TMS; 25    1.07e-01 1.41e-01   0.48200
## 23              Hydroxyproline; 64   -1.05e-01 1.48e-01   0.48400
## 24     3-Indolepropionic acid; 41    1.01e-01 1.64e-01   0.51100
## 25                Serine, 3TMS; 14    9.81e-02 1.76e-01   0.52000
## 26      Eicosapentaenoic acid; 55   -9.71e-02 1.80e-01   0.52000
## 27  2-hydroxy Isovaleric acid; 38    8.26e-02 2.55e-01   0.64100
## 28 4-Hydroxyphenyllactic acid; 44   -8.21e-02 2.57e-01   0.64100
## 29             1,3-Propanediol; 34    8.12e-02 2.62e-01   0.64100
## 30          Glyceryl-glycoside; 59   -8.06e-02 2.66e-01   0.64100
## 31          Succinic acid, 2TMS; 7   -7.98e-02 2.71e-01   0.64100
## 32               Glycine, 3TMS; 17   -7.82e-02 2.81e-01   0.64100
## 33           Cholesterol, TMS; 23    7.66e-02 2.90e-01   0.64100
## 34             Threonine, 3TMS; 12    7.66e-02 2.90e-01   0.64100
## 35          11-Eicosenoic acid; 35   -7.53e-02 2.99e-01   0.64100
## 36            Pyroglutamic acid; 69   -7.37e-02 3.09e-01   0.64300
## 37             Dodecanoic acid; 54   -6.78e-02 3.49e-01   0.69600
## 38                     Glycerol; 57   -6.68e-02 3.57e-01   0.69600
## 39            Myristoleic acid; 65   -6.61e-02 3.62e-01   0.69600
## 40      Arachidonic acid, TMS; 24   -6.21e-02 3.92e-01   0.73400
## 41         Phenylalanine, 2TMS; 13    6.02e-02 4.06e-01   0.74300
## 42                L-5-Oxoproline; 63   -5.61e-02 4.39e-01   0.77700
## 43               Octanoic acid; 68    5.42e-02 4.54e-01   0.77700
## 44 3-Hydroxybutyric acid, 2TMS; 1   -5.38e-02 4.57e-01   0.77700
## 45              1-Monopalmitin; 37   -5.23e-02 4.70e-01   0.77700
## 46               Arachidic acid; 46    5.05e-02 4.86e-01   0.77700
```

```
## 47           Malic acid, 3TMS; 11   -5.03e-02 4.87e-01   0.77700
## 48           alpha-Tocopherol; 26    4.90e-02 4.99e-01   0.77900
## 49              Nonanoic acid; 67    4.76e-02 5.12e-01   0.78300
## 50              Decanoic acid; 52   -4.53e-02 5.32e-01   0.79800
## 51        Heptadecanoic acid; 61   -4.28e-02 5.55e-01   0.80900
## 52                Campesterol; 49   -4.22e-02 5.61e-01   0.80900
## 53           Tridecanoic acid; 74   -3.70e-02 6.10e-01   0.85700
## 54                Lactic acid; 29    3.53e-02 6.26e-01   0.85700
## 55       2-Palmitoylglycerol; 39   -3.50e-02 6.29e-01   0.85700
## 56          Nonadecanoic acid; 66   -3.27e-02 6.52e-01   0.87300
## 57              Hydroxylamine; 62    2.95e-02 6.84e-01   0.89900
## 58               Pyruvic acid; 31    2.76e-02 7.03e-01   0.89900
## 59                Bisphenol A; 48    2.66e-02 7.14e-01   0.89900
## 60        Glutamic acid, 3TMS; 8   -2.49e-02 7.31e-01   0.89900
## 61          Aminomalonic acid; 45    2.42e-02 7.39e-01   0.89900
## 62      4-Deoxytetronic acid; 33   -2.37e-02 7.44e-01   0.89900
## 63    4-Hydroxybutanoic acid; 43   -1.91e-02 7.93e-01   0.93900
## 64 alpha-ketoglutaric acid, TMS M   -1.79e-02 8.05e-01   0.93900
## 65            Arabinopyranose; 51   -1.62e-02 8.23e-01   0.93900
## 66                1-Dodecanol; 36    1.59e-02 8.26e-01   0.93900
## 67              Proline, 2TMS; 21    1.24e-02 8.64e-01   0.96800
## 68      Docosahexaenoic acid; 53    9.69e-03 8.94e-01   0.98600
## 69          Linoleic acid, TMS; 4   -6.11e-03 9.33e-01   1.00000
## 70        Heptadecanoic acid; 60   -5.25e-03 9.42e-01   1.00000
## 71          Palmitic acid, TMS; 5    2.92e-03 9.68e-01   1.00000
## 72 2-Hydroxybutyric acid, 2TMS; 2   -8.25e-04 9.91e-01   1.00000
## 73       3-Indoleacetic acid; 40   -7.72e-04 9.91e-01   1.00000
## 74           Stearic acid, TMS; 2    2.88e-04 9.97e-01   1.00000
## 75                   Tyrosine; 75   -3.16e-05 1.00e+00   1.00000
```

###### 5.4.1.2 Forest Plot of Model Coefficients

```
## Warning: Ignoring unknown aesthetics: x
```

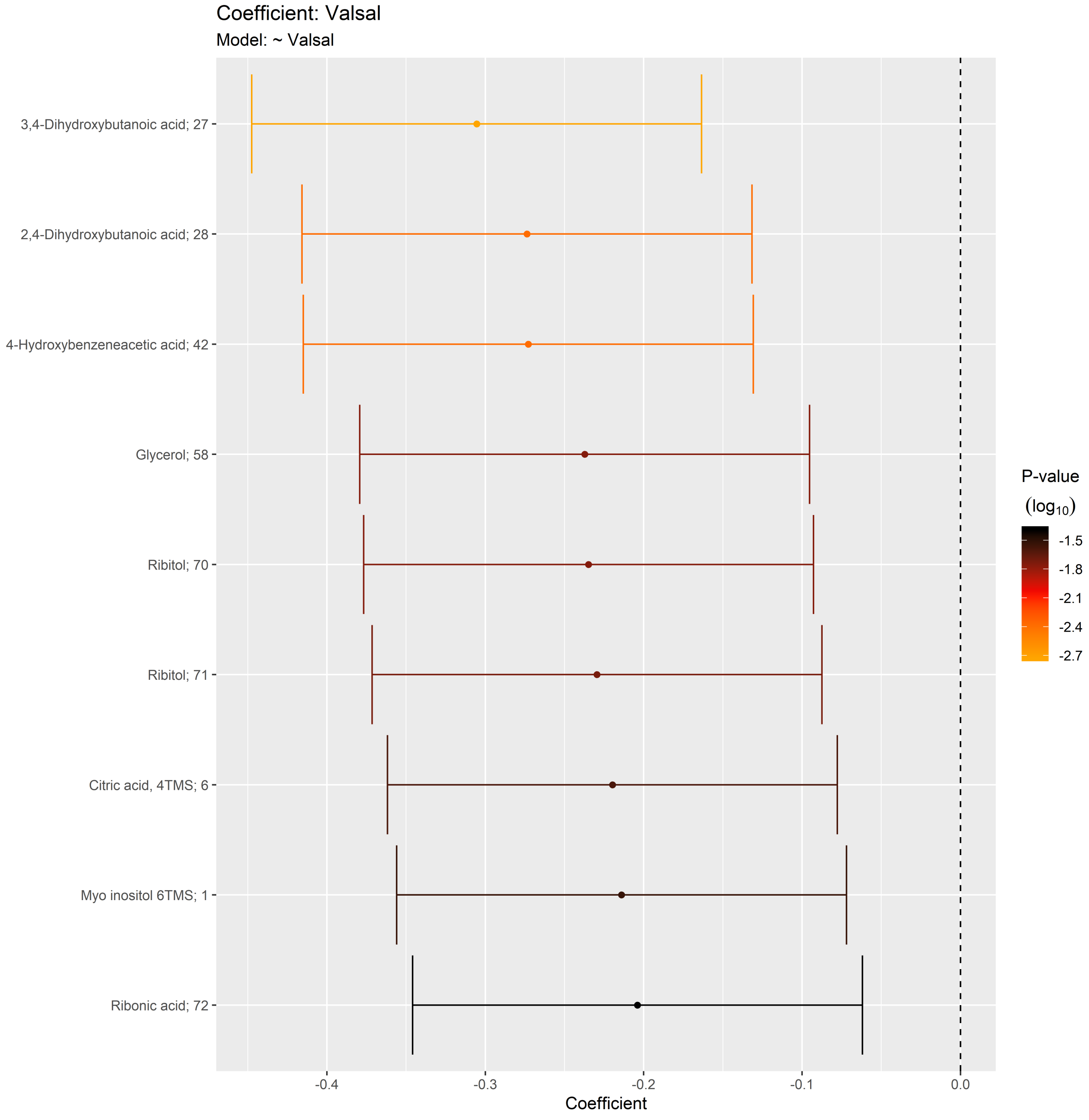

##### 5.4.2 Adjusted Model

```
## [1] "Fitting models:"
## [1] "~ Valsal + Age + bmi + Blood_glucose + Duration_DM + Gender + Hba1c_baseline + log_Blood_TGA + S
## [1] ""
```

###### 5.4.2.1 Tables of Model Coefficients

```
## [1] ""
## [1] "Table: Valsal"
## [1] "  (from model: "
## [1] "   ~ Valsal + Age + bmi + Blood_glucose + Duration_DM + Gender"
## [1] "     + Hba1c_baseline + log_Blood_TGA + Smoking + Statin +"
## [1] "     Total_cholesterol)"
## [1] ""
##                                Name Coefficient P.Value adj.P.Val
## 1  4-Hydroxybenzeneacetic acid; 4    -0.23400 0.00308    0.0779
## 2                    Glycerol; 58    -0.22900 0.00379    0.0779
## 3                     Ribitol; 70    -0.22700 0.00408    0.0779
## 4             Citric acid, 4TMS; 6    -0.22700 0.00416    0.0779
## 5                 Ribonic acid; 72    -0.21700 0.00619    0.0929
## 6  2,4-Dihydroxybutanoic acid; 28    -0.21100 0.00775    0.0969
## 7  3,4-Dihydroxybutanoic acid; 27    -0.18900 0.01720    0.1650
## 8             Myo inositol 6TMS; 1    -0.18800 0.01760    0.1650
## 9               Oleic acid, TMS; 3    -0.16300 0.03880    0.3210
## 10                    Ribitol; 71    -0.15600 0.04840    0.3210
## 11            Isoleucine, 2TMS; 18     0.15200 0.05450    0.3210
## 12              Tartronic acid; 73     0.14800 0.06100    0.3210
## 13                Valine, 2TMS; 20     0.14700 0.06220    0.3210
## 14           Fumaric acid, 2TMS; 9    -0.14700 0.06340    0.3210
## 15               Leucine, 2TMS; 19     0.14600 0.06480    0.3210
## 16          Benzeneacetic acid; 47    -0.14400 0.06850    0.3210
## 17                   Glycerol; 57    -0.14100 0.07560    0.3330
## 18            alpha-Tocopherol; 26     0.13200 0.09510    0.3960
## 19            Methionine, 2TMS; 16     0.12700 0.10800    0.4020
## 20                Ethanolamine; 56     0.12600 0.11100    0.4020
## 21               Alanine, 2TMS; 25     0.12500 0.11500    0.4020
## 22               Glycine, 3TMS; 17    -0.12400 0.11800    0.4020
## 23                  Creatinine; 50    -0.11500 0.14800    0.4790
## 24              Hydroxyproline; 64    -0.11300 0.15300    0.4790
## 25          Succinic acid, 2TMS; 7    -0.10300 0.19200    0.5760
## 26               Glyceric acid; 30     0.09940 0.20900    0.6020
## 27                 Serine, 3TMS; 14     0.07750 0.32700    0.8750
## 28      3-Indolepropionic acid; 41     0.07440 0.34700    0.8750
## 29            Tridecanoic acid; 74    -0.07310 0.35500    0.8750
## 30        4-Deoxytetronic acid; 32    -0.07250 0.36000    0.8750
## 31              Threonine, 3TMS; 12     0.07050 0.37300    0.8750
## 32       Arachidonic acid, TMS; 24    -0.07040 0.37300    0.8750
## 33          11-Eicosenoic acid; 35    -0.06700 0.39700    0.9010
## 34              Arachidic acid; 46     0.06430 0.41600    0.9010
## 35             Malic acid, 3TMS; 11    -0.06140 0.43800    0.9010
## 36              1-Monopalmitin; 37    -0.06130 0.43800    0.9010
## 37              1,3-Propanediol; 34     0.05930 0.45400    0.9010
## 38                Pyruvic acid; 31     0.05630 0.47700    0.9010
## 39               Hydroxylamine; 62     0.05290 0.50300    0.9010
## 40       Eicosapentaenoic acid; 55    -0.05270 0.50500    0.9010
## 41      4-Hydroxybutanoic acid; 43    -0.05040 0.52400    0.9010
## 42  2-hydroxy Isovaleric acid; 38     0.05000 0.52800    0.9010
## 43          Heptadecanoic acid; 61    -0.04780 0.54500    0.9010
## 44              Arabinopyranose; 51    -0.04760 0.54700    0.9010
```

```
## 45        3-Indoleacetic acid; 40     0.04650 0.55600    0.9010
## 46 4-Hydroxyphenyllactic acid; 44    -0.04500 0.57000    0.9010
## 47             L-5-Oxoproline; 63    -0.04430 0.57600    0.9010
## 48           Stearic acid, TMS; 2     0.04280 0.58900    0.9010
## 49                Campesterol; 49    -0.04270 0.58900    0.9010
## 50          Linoleic acid, TMS; 4    -0.03990 0.61400    0.9100
## 51                 1-Dodecanol; 36    -0.03930 0.61900    0.9100
## 52           Myristoleic acid; 65    -0.03740 0.63600    0.9180
## 53                   Tyrosine; 75     0.03390 0.66800    0.9300
## 54            Dodecanoic acid; 54    -0.03380 0.67000    0.9300
## 55         Heptadecanoic acid; 60    -0.03070 0.69800    0.9480
## 56       Docosahexaenoic acid; 53     0.02890 0.71500    0.9480
## 57        Phenylalanine, 2TMS; 13     0.02570 0.74500    0.9480
## 58              Proline, 2TMS; 21     0.02460 0.75600    0.9480
## 59              Nonanoic acid; 67     0.02440 0.75700    0.9480
## 60           Cholesterol, TMS; 23     0.02320 0.76900    0.9480
## 61 2-Hydroxybutyric acid, 2TMS; 2    -0.02300 0.77100    0.9480
## 62          Nonadecanoic acid; 66    -0.02030 0.79800    0.9540
## 63          Aminomalonic acid; 45    -0.01910 0.80900    0.9540
## 64 alpha-ketoglutaric acid, TMS M     0.01840 0.81600    0.9540
## 65          Pyroglutamic acid; 69    -0.01600 0.84000    0.9540
## 66                Bisphenol A; 48    -0.01400 0.85900    0.9540
## 67              Octanoic acid; 68     0.01340 0.86600    0.9540
## 68         Glyceryl-glycoside; 59    -0.01340 0.86600    0.9540
## 69              Decanoic acid; 52     0.01080 0.89200    0.9540
## 70        2-Palmitoylglycerol; 39    -0.00955 0.90400    0.9540
## 71 3-Hydroxybutyric acid, 2TMS; 1     0.00888 0.91100    0.9540
## 72         Palmitic acid, TMS; 5     -0.00837 0.91600    0.9540
## 73                Lactic acid; 29    -0.00464 0.95300    0.9730
## 74         Glutamic acid, 3TMS; 8     0.00392 0.96100    0.9730
## 75       4-Deoxytetronic acid; 33    -0.00199 0.98000    0.9800
```

###### 5.4.2.2 Forest Plot of Model Coefficients

```
## Warning: Ignoring unknown aesthetics: x

## Warning: Ignoring unknown aesthetics: x

## Warning: Ignoring unknown aesthetics: x

## Warning: Ignoring unknown aesthetics: x

## Warning: Ignoring unknown aesthetics: x

## NULL
```

##### 5.4.3 Fully-Adjusted Model

```
## [1] "Fitting models:"
## [1] "~ Valsal + Age + bmi + Blood_glucose + Duration_DM + Gender + Hba1c_baseline + log_Blood_TGA + S
## [1] ""
```

###### 5.4.3.1 Tables of Model Coefficients

```
## [1] ""
## [1] "Table: Valsal"
## [1] "  (from model: "
## [1] "   ~ Valsal + Age + bmi + Blood_glucose + Duration_DM + Gender"
## [1] "     + Hba1c_baseline + log_Blood_TGA + Smoking + Statin +"
## [1] "     Total_cholesterol + egfr)"
## [1] ""
##                              Name Coefficient P.Value adj.P.Val
## 1                    Glycerol; 58   -2.23e-01 0.00548     0.219
## 2                     Ribitol; 70   -2.22e-01 0.00585     0.219
## 3              Oleic acid, TMS; 3   -1.94e-01 0.01590     0.398
## 4                    Glycerol; 57   -1.74e-01 0.03000     0.550
## 5             Citric acid, 4TMS; 6   -1.67e-01 0.03670     0.550
## 6              Tartronic acid; 73    1.56e-01 0.05070     0.580
## 7             alpha-Tocopherol; 26    1.53e-01 0.05570     0.580
## 8  4-Hydroxybenzeneacetic acid; 4   -1.49e-01 0.06180     0.580
## 9           Benzeneacetic acid; 47   -1.31e-01 0.10400     0.865
## 10               Alanine, 2TMS; 25    1.24e-01 0.12200     0.880
## 11         3-Indoleacetic acid; 40    1.18e-01 0.14300     0.880
## 12           Fumaric acid, 2TMS; 9   -1.17e-01 0.14400     0.880
## 13                Ethanolamine; 56    1.13e-01 0.15900     0.880
## 14 2,4-Dihydroxybutanoic acid; 28   -1.06e-01 0.18400     0.880
## 15                Ribonic acid; 72   -1.05e-01 0.18900     0.880
## 16 3,4-Dihydroxybutanoic acid; 27   -1.03e-01 0.19700     0.880
## 17 2-Hydroxybutyric acid, 2TMS; 2   -1.01e-01 0.20800     0.880
## 18         Succinic acid, 2TMS; 7   -9.82e-02 0.22200     0.880
## 19               Leucine, 2TMS; 19    9.66e-02 0.22800     0.880
## 20       Arachidonic acid, TMS; 24   -9.53e-02 0.23500     0.880
## 21               Glycine, 3TMS; 17   -8.34e-02 0.29800     0.894
## 22            Isoleucine, 2TMS; 18    8.23e-02 0.30400     0.894
## 23                Valine, 2TMS; 20    8.15e-02 0.30800     0.894
## 24         11-Eicosenoic acid; 35   -7.98e-02 0.32000     0.894
## 25      Eicosapentaenoic acid; 55   -7.55e-02 0.34600     0.894
## 26     3-Indolepropionic acid; 41    7.48e-02 0.35200     0.894
## 27            Myo inositol 6TMS; 1   -7.39e-02 0.35400     0.894
## 28       4-Deoxytetronic acid; 33    7.26e-02 0.36500     0.894
## 29         Heptadecanoic acid; 61   -7.15e-02 0.37300     0.894
## 30            Methionine, 2TMS; 16    7.12e-02 0.37400     0.894
## 31              L-5-Oxoproline; 63   -6.78e-02 0.39900     0.894
## 32               Glyceric acid; 30    6.58e-02 0.41100     0.894
## 33             1,3-Propanediol; 34    6.57e-02 0.41300     0.894
## 34           Linoleic acid, TMS; 4   -5.98e-02 0.45700     0.894
## 35              Hydroxyproline; 64   -5.95e-02 0.45900     0.894
## 36             Dodecanoic acid; 54   -5.81e-02 0.47000     0.894
## 37            Myristoleic acid; 65   -5.55e-02 0.48900     0.894
## 38            Tridecanoic acid; 74   -5.50e-02 0.49400     0.894
## 39                 Campesterol; 49   -5.48e-02 0.49400     0.894
## 40               Hydroxylamine; 62    5.37e-02 0.50400     0.894
## 41                Pyruvic acid; 31    5.29e-02 0.51000     0.894
## 42              1-Monopalmitin; 37   -5.29e-02 0.51000     0.894
## 43               Octanoic acid; 68   -5.21e-02 0.51600     0.894
## 44           Nonadecanoic acid; 66   -5.11e-02 0.52500     0.894
```

```
## 45          Palmitic acid, TMS; 5   -4.83e-02 0.54700     0.895
## 46     4-Hydroxybutanoic acid; 43   -4.82e-02 0.54900     0.895
## 47              Arachidic acid; 46    4.58e-02 0.56900     0.907
## 48                     Ribitol; 71   -4.28e-02 0.59200     0.913
## 49          Glyceryl-glycoside; 59    4.25e-02 0.59600     0.913
## 50             Threonine, 3TMS; 12    3.87e-02 0.63000     0.945
## 51                 1-Dodecanol; 36   -3.67e-02 0.64800     0.953
## 52             Arabinopyranose; 51   -3.48e-02 0.66300     0.957
## 53                  Creatinine; 50   -3.06e-02 0.70200     0.962
## 54          Glutamic acid, 3TMS; 8   -2.96e-02 0.71100     0.962
## 55          Heptadecanoic acid; 60   -2.93e-02 0.71500     0.962
## 56           Pyroglutamic acid; 69    2.84e-02 0.72400     0.962
## 57                 Bisphenol A; 48   -2.34e-02 0.77200     0.962
## 58               Proline, 2TMS; 21    2.31e-02 0.77300     0.962
## 59                    Tyrosine; 75   -2.15e-02 0.78900     0.962
## 60                 Lactic acid; 29   -2.12e-02 0.79200     0.962
## 61               Nonanoic acid; 67    2.09e-02 0.79500     0.962
## 62            Malic acid, 3TMS; 11   -2.08e-02 0.79600     0.962
## 63 4-Hydroxyphenyllactic acid; 44   -1.68e-02 0.83400     0.990
## 64 alpha-ketoglutaric acid, TMS M    1.52e-02 0.85000     0.990
## 65         Phenylalanine, 2TMS; 13    9.47e-03 0.90600     0.990
## 66        Docosahexaenoic acid; 53    9.11e-03 0.90900     0.990
## 67  2-hydroxy Isovaleric acid; 38    8.38e-03 0.91700     0.990
## 68         2-Palmitoylglycerol; 39   -6.68e-03 0.93400     0.990
## 69           Aminomalonic acid; 45   -4.64e-03 0.95400     0.990
## 70        4-Deoxytetronic acid; 32    4.61e-03 0.95400     0.990
## 71            Stearic acid, TMS; 2   -3.03e-03 0.97000     0.990
## 72 3-Hydroxybutyric acid, 2TMS; 1    2.76e-03 0.97300     0.990
## 73                Serine, 3TMS; 14    2.65e-03 0.97400     0.990
## 74            Cholesterol, TMS; 23   -2.35e-03 0.97700     0.990
## 75               Decanoic acid; 52   -1.57e-05 1.00000     1.000
```

###### 5.4.3.2 Forest Plot of Model Coefficients

```
## Warning: Ignoring unknown aesthetics: x

## Warning: Ignoring unknown aesthetics: x

## Warning: Ignoring unknown aesthetics: x

## Warning: Ignoring unknown aesthetics: x

## Warning: Ignoring unknown aesthetics: x

## NULL
```

#### 5.5 Heart Rate Variablity (SDNN)

##### 5.5.1 Crude Model

```
## [1] "Fitting models:"
## [1] "~ SDNN + rest_HR_vag"
## [1] ""
```

###### 5.5.1.1 Tables of Model Coefficients

```
## [1] ""
## [1] "Table: SDNN"
## [1] "  (from model: "
## [1] "    ~ SDNN + rest_HR_vag)"
## [1] ""
##                               Name Coefficient P.Value adj.P.Val
## 1        4-Deoxytetronic acid; 32    -0.17900 0.00562     0.243
## 2            Myo inositol 6TMS; 1    -0.16600 0.01030     0.243
## 3  2,4-Dihydroxybutanoic acid; 28    -0.15900 0.01410     0.243
## 4  3,4-Dihydroxybutanoic acid; 27    -0.15700 0.01530     0.243
## 5                Valine, 2TMS; 20    -0.15500 0.01620     0.243
## 6                  Creatinine; 50    -0.14400 0.02590     0.322
## 7               Leucine, 2TMS; 19    -0.14000 0.03010     0.322
## 8             Arabinopyranose; 51    -0.13100 0.04300     0.403
## 9                 Lactic acid; 29     0.12200 0.05870     0.448
## 10            Tridecanoic acid; 74    -0.12200 0.05980     0.448
## 11                     Ribitol; 70    -0.11800 0.06720     0.458
## 12                Ribonic acid; 72    -0.11300 0.08140     0.474
## 13            Threonine, 3TMS; 12    -0.11200 0.08430     0.474
## 14        Phenylalanine, 2TMS; 13    -0.11000 0.08840     0.474
## 15           Isoleucine, 2TMS; 18    -0.09890 0.12600     0.629
## 16            Dodecanoic acid; 54     0.09200 0.15400     0.638
## 17         Glutamic acid, 3TMS; 8    -0.08810 0.17200     0.638
## 18              Octanoic acid; 68     0.08510 0.18800     0.638
## 19 4-Hydroxybenzeneacetic acid; 4    -0.08450 0.19100     0.638
## 20        3-Indoleacetic acid; 40    -0.08430 0.19200     0.638
## 21             L-5-Oxoproline; 63    -0.08280 0.20000     0.638
## 22              Nonanoic acid; 67    -0.08260 0.20100     0.638
## 23             Oleic acid, TMS; 3    -0.08200 0.20400     0.638
## 24     3-Indolepropionic acid; 41     0.07970 0.21700     0.638
## 25              Proline, 2TMS; 21    -0.07700 0.23300     0.638
## 26         Glyceryl-glycoside; 59    -0.07690 0.23400     0.638
## 27             Arachidic acid; 46     0.07590 0.24000     0.638
## 28            Citric acid, 4TMS; 6    -0.07500 0.24600     0.638
## 29        2-Palmitoylglycerol; 39     0.07480 0.24700     0.638
## 30              Glycine, 3TMS; 17    -0.06700 0.30000     0.730
## 31             Hydroxyproline; 64    -0.06550 0.31000     0.730
## 32                    Ribitol; 71    -0.06460 0.31700     0.730
## 33               Serine, 3TMS; 14    -0.06160 0.34100     0.730
## 34 3-Hydroxybutyric acid, 2TMS; 1    -0.05840 0.36600     0.730
## 35 2-Hydroxybutyric acid, 2TMS; 2    -0.05830 0.36700     0.730
## 36                   Glycerol; 57    -0.05800 0.36900     0.730
## 37     4-Hydroxybutanoic acid; 43    -0.05780 0.37100     0.730
## 38         Benzeneacetic acid; 47    -0.05680 0.37900     0.730
## 39           alpha-Tocopherol; 26    -0.05680 0.37900     0.730
## 40           Pyroglutamic acid; 69   -0.05550 0.39100     0.732
## 41           Methionine, 2TMS; 16    -0.05390 0.40400     0.740
## 42            Myristoleic acid; 65   -0.05110 0.42900     0.758
## 43      Arachidonic acid, TMS; 24    -0.05010 0.43800     0.758
## 44                Bisphenol A; 48    -0.04860 0.45200     0.758
## 45           Malic acid, 3TMS; 11     0.04830 0.45500     0.758
## 46              Hydroxylamine; 62     0.04470 0.48900     0.797
```

```
## 47                    Tyrosine; 75    -0.04230 0.51200     0.817
## 48             1-Monopalmitin; 37    -0.04070 0.52800     0.825
## 49              Glyceric acid; 30     0.03800 0.55600     0.851
## 50         Heptadecanoic acid; 60     0.03660 0.57100     0.856
## 51  2-hydroxy Isovaleric acid; 38    -0.03260 0.61400     0.860
## 52              Decanoic acid; 52     0.03250 0.61500     0.860
## 53          Fumaric acid, 2TMS; 9    -0.02940 0.64900     0.860
## 54          Palmitic acid, TMS; 5    -0.02930 0.65000     0.860
## 55               Ethanolamine; 56     0.02900 0.65300     0.860
## 56              Alanine, 2TMS; 25    -0.02760 0.66900     0.860
## 57          Aminomalonic acid; 45    -0.02670 0.67900     0.860
## 58       4-Deoxytetronic acid; 33     0.02620 0.68500     0.860
## 59          Linoleic acid, TMS; 4    -0.02600 0.68700     0.860
## 60     Eicosapentaenoic acid; 55    -0.02490 0.70000     0.860
## 61            Tartronic acid; 73     0.02260 0.72700     0.860
## 62               Campesterol; 49    -0.02180 0.73600     0.860
## 63              Pyruvic acid; 31    -0.02120 0.74200     0.860
## 64         Heptadecanoic acid; 61    -0.02080 0.74700     0.860
## 65            1,3-Propanediol; 34     0.02060 0.75000     0.860
## 66                  Glycerol; 58    -0.02000 0.75700     0.860
## 67 alpha-ketoglutaric acid, TMS M     0.01710 0.79100     0.886
## 68 4-Hydroxyphenyllactic acid; 44     0.01550 0.81000     0.894
## 69          Nonadecanoic acid; 66    -0.01330 0.83700     0.896
## 70           Cholesterol, TMS; 23    -0.01280 0.84200     0.896
## 71         Succinic acid, 2TMS; 7     0.01240 0.84800     0.896
## 72           Stearic acid, TMS; 2     0.00640 0.92100     0.959
## 73         11-Eicosenoic acid; 35     0.00458 0.94300     0.965
## 74               1-Dodecanol; 36    -0.00305 0.96200     0.965
## 75      Docosahexaenoic acid; 53     0.00281 0.96500     0.965
```

###### 5.5.1.2 Forest Plot of Model Coefficients

```
## Warning: Ignoring unknown aesthetics: x
```

```
## NULL
```

##### 5.5.2 Adjusted Model

```
## [1] "Fitting models:"
## [1] "~ SDNN + rest_HR_vag + Age + bmi + Blood_glucose + Duration_DM + Gender + Hba1c_baseline + log_
## [1] ""
```

###### 5.5.2.1 Tables of Model Coefficients

```
## [1] ""
## [1] "Table: SDNN"
## [1] "  (from model: "
## [1] "   ~ SDNN + rest_HR_vag + Age + bmi + Blood_glucose +"
## [1] "     Duration_DM + Gender + Hba1c_baseline + log_Blood_TGA +"
## [1] "     Smoking + Statin + Total_cholesterol)"
## [1] ""
##                               Name Coefficient P.Value adj.P.Val
## 1                Valine, 2TMS; 20    -0.18100 0.00636     0.477
## 2        4-Deoxytetronic acid; 32    -0.15600 0.01830     0.547
## 3             Threonine, 3TMS; 12    -0.13500 0.04080     0.547
## 4               Leucine, 2TMS; 19    -0.13200 0.04540     0.547
## 5         Phenylalanine, 2TMS; 13    -0.13000 0.05010     0.547
## 6            Tridecanoic acid; 74    -0.12800 0.05320     0.547
## 7                  Creatinine; 50    -0.12400 0.06130     0.547
## 8  2,4-Dihydroxybutanoic acid; 28    -0.12100 0.06730     0.547
## 9             Dodecanoic acid; 54     0.11700 0.07700     0.547
## 10             Nonanoic acid; 67    -0.11700 0.07790     0.547
## 11          Myo inositol 6TMS; 1    -0.11400 0.08390     0.547
## 12        2-Palmitoylglycerol; 39     0.11200 0.09100     0.547
## 13               Lactic acid; 29     0.11100 0.09490     0.547
## 14              Ribonic acid; 72    -0.10000 0.13000     0.670
## 15         Glutamic acid, 3TMS; 8    -0.09710 0.14200     0.670
## 16 3,4-Dihydroxybutanoic acid; 27    -0.09490 0.15200     0.670
## 17  2-hydroxy Isovaleric acid; 38    -0.09300 0.16000     0.670
## 18           Isoleucine, 2TMS; 18    -0.08930 0.17700     0.670
## 19              Hydroxylamine; 62     0.08820 0.18200     0.670
## 20     4-Hydroxybutanoic acid; 43    -0.08600 0.19400     0.670
## 21          Aminomalonic acid; 45    -0.08510 0.19800     0.670
## 22                    Ribitol; 70    -0.08210 0.21500     0.670
## 23              L-5-Oxoproline; 63    -0.08170 0.21700     0.670
## 24             Arabinopyranose; 51    -0.08130 0.22000     0.670
## 25               Glycine, 3TMS; 17    -0.08060 0.22300     0.670
## 26                 Bisphenol A; 48    -0.07820 0.23700     0.685
## 27              Oleic acid, TMS; 3    -0.07420 0.26200     0.703
## 28              Hydroxyproline; 64    -0.07410 0.26300     0.703
## 29              Arachidic acid; 46     0.07220 0.27500     0.712
## 30                Serine, 3TMS; 14    -0.07080 0.28500     0.712
## 31         3-Indoleacetic acid; 40    -0.06880 0.29900     0.718
## 32           Malic acid, 3TMS; 11     0.06630 0.31700     0.718
## 33                   Glycerol; 57    -0.06540 0.32300     0.718
## 34 alpha-ketoglutaric acid, TMS M     0.06510 0.32600     0.718
## 35     3-Indolepropionic acid; 41     0.06120 0.35500     0.759
## 36               Proline, 2TMS; 21    -0.06010 0.36400     0.759
## 37 2-Hydroxybutyric acid, 2TMS; 2    -0.05480 0.40700     0.809
## 38           Cholesterol, TMS; 23    -0.05450 0.41000     0.809
## 39              Octanoic acid; 68     0.05290 0.42400     0.816
## 40      Arachidonic acid, TMS; 24    -0.05070 0.44400     0.832
## 41           Methionine, 2TMS; 16    -0.04940 0.45600     0.833
## 42         Heptadecanoic acid; 61    -0.04650 0.48200     0.861
## 43 4-Hydroxybenzeneacetic acid; 4    -0.04320 0.51400     0.881
## 44        Docosahexaenoic acid; 53     0.04130 0.53200     0.881
```

```
## 45         Benzeneacetic acid; 47    -0.04070 0.53900     0.881
## 46          Palmitic acid, TMS; 5    -0.03830 0.56300     0.881
## 47 4-Hydroxyphenyllactic acid; 44     0.03820 0.56400     0.881
## 48          Linoleic acid, TMS; 4    -0.03720 0.57400     0.881
## 49         Glyceryl-glycoside; 59    -0.03690 0.57800     0.881
## 50              Decanoic acid; 52     0.03580 0.58900     0.881
## 51           Myristoleic acid; 65    -0.03480 0.59900     0.881
## 52              Glyceric acid; 30     0.03010 0.64900     0.931
## 53 3-Hydroxybutyric acid, 2TMS; 1    -0.02910 0.66100     0.931
## 54         11-Eicosenoic acid; 35     0.02630 0.69100     0.931
## 55           Citric acid, 4TMS; 6    -0.02570 0.69800     0.931
## 56          Pyroglutamic acid; 69    -0.02480 0.70800     0.931
## 57               Ethanolamine; 56     0.02440 0.71200     0.931
## 58           Stearic acid, TMS; 2     0.02370 0.72000     0.931
## 59                   Tyrosine; 75    -0.02010 0.76200     0.952
## 60      Eicosapentaenoic acid; 55     0.01840 0.78100     0.952
## 61                1-Dodecanol; 36    -0.01700 0.79700     0.952
## 62             Tartronic acid; 73     0.01680 0.79900     0.952
## 63          Nonadecanoic acid; 66    -0.01510 0.81900     0.952
## 64                   Glycerol; 58    -0.01440 0.82700     0.952
## 65               Pyruvic acid; 31     0.01390 0.83400     0.952
## 66             1-Monopalmitin; 37    -0.01330 0.84100     0.952
## 67                Campesterol; 49     0.01150 0.86300     0.952
## 68           alpha-Tocopherol; 26    -0.01120 0.86600     0.952
## 69         Succinic acid, 2TMS; 7    -0.01040 0.87600     0.952
## 70            1,3-Propanediol; 34     0.00917 0.89000     0.953
## 71         Heptadecanoic acid; 60    -0.00461 0.94400     0.974
## 72                    Ribitol; 71     0.00392 0.95300     0.974
## 73       4-Deoxytetronic acid; 33    -0.00294 0.96500     0.974
## 74              Alanine, 2TMS; 25    -0.00288 0.96500     0.974
## 75          Fumaric acid, 2TMS; 9    -0.00215 0.97400     0.974
```

###### 5.5.2.2 Forest Plot of Model Coefficients

```
## Warning: Ignoring unknown aesthetics: x

## Warning: Ignoring unknown aesthetics: x

## Warning: Ignoring unknown aesthetics: x

## Warning: Ignoring unknown aesthetics: x

## Warning: Ignoring unknown aesthetics: x

## Warning: Ignoring unknown aesthetics: x

## Warning: Ignoring unknown aesthetics: x

## NULL
```

###### 5.5.3 Fully-Adjusted Model

```
## [1] "Fitting models:"
## [1] "~ SDNN + rest_HR_vag + Age + bmi + Blood_glucose + Duration_DM + Gender + Hba1c_baseline + log_]
## [1] ""
```

###### 5.5.3.1 Tables of Model Coefficients

```
## [1] ""
## [1] "Table: SDNN"
## [1] "  (from model: "
## [1] "   ~ SDNN + rest_HR_vag + Age + bmi + Blood_glucose +"
## [1] "     Duration_DM + Gender + Hba1c_baseline + log_Blood_TGA +"
## [1] "     Smoking + Statin + Total_cholesterol + egfr)"
## [1] ""
##                                Name Coefficient P.Value adj.P.Val
## 1                Valine, 2TMS; 20   -0.210000 0.00137     0.103
## 2               Leucine, 2TMS; 19   -0.152000 0.02060     0.563
## 3             Threonine, 3TMS; 12   -0.147000 0.02510     0.563
## 4            Tridecanoic acid; 74   -0.126000 0.05590     0.563
## 5         Phenylalanine, 2TMS; 13   -0.124000 0.05990     0.563
## 6            Isoleucine, 2TMS; 18   -0.120000 0.06740     0.563
## 7          Glutamic acid, 3TMS; 8   -0.117000 0.07340     0.563
## 8   2-hydroxy Isovaleric acid; 38   -0.117000 0.07400     0.563
## 9               Nonanoic acid; 67   -0.117000 0.07610     0.563
## 10       4-Deoxytetronic acid; 32   -0.114000 0.08120     0.563
## 11            Dodecanoic acid; 54    0.114000 0.08260     0.563
## 12        2-Palmitoylglycerol; 39    0.112000 0.09030     0.564
## 13               Hydroxylamine; 62    0.105000 0.11200     0.601
## 14               Serine, 3TMS; 14   -0.104000 0.11200     0.601
## 15                Lactic acid; 29    0.102000 0.12000     0.601
## 16 2-Hydroxybutyric acid, 2TMS; 2   -0.095600 0.14400     0.676
## 17          Aminomalonic acid; 45   -0.088200 0.17800     0.714
## 18           Malic acid, 3TMS; 11    0.087200 0.18500     0.714
## 19             L-5-Oxoproline; 63   -0.086100 0.19100     0.714
## 20                Bisphenol A; 48   -0.082800 0.20900     0.714
## 21                   Glycerol; 57   -0.081500 0.21500     0.714
## 22     4-Hydroxybutanoic acid; 43   -0.078600 0.23300     0.714
## 23              Oleic acid, TMS; 3  -0.078200 0.23300     0.714
## 24            Arabinopyranose; 51   -0.076400 0.24200     0.714
## 25                 Creatinine; 50   -0.076300 0.24500     0.714
## 26           Cholesterol, TMS; 23   -0.074500 0.25400     0.714
## 27           Methionine, 2TMS; 16   -0.071400 0.27700     0.714
## 28                    Ribitol; 70   -0.069600 0.29000     0.714
## 29                    Ribitol; 71    0.067600 0.30000     0.714
## 30 alpha-ketoglutaric acid, TMS M    0.067100 0.30800     0.714
## 31              Glycine, 3TMS; 17   -0.066300 0.31400     0.714
## 32             Arachidic acid; 46    0.065300 0.32100     0.714
## 33 4-Hydroxyphenyllactic acid; 44    0.064300 0.32800     0.714
## 34      Arachidonic acid, TMS; 24   -0.063100 0.33800     0.714
## 35 2,4-Dihydroxybutanoic acid; 28   -0.062200 0.34000     0.714
## 36     3-Indolepropionic acid; 41    0.062400 0.34300     0.714
## 37          Heptadecanoic acid; 61   -0.054200 0.41000     0.822
## 38              Proline, 2TMS; 21   -0.053000 0.41900     0.822
## 39           Myo inositol 6TMS; 1   -0.050900 0.43400     0.822
## 40          Palmitic acid, TMS; 5   -0.050800 0.43900     0.822
## 41               Ribonic acid; 72   -0.049400 0.44900     0.822
## 42 3,4-Dihydroxybutanoic acid; 27   -0.043900 0.50100     0.886
## 43             Hydroxyproline; 64   -0.042700 0.51600     0.886
## 44                   Tyrosine; 75   -0.041200 0.53000     0.886
```

```
## 45        3-Indoleacetic acid; 40   -0.040500 0.53800     0.886
## 46      4-Deoxytetronic acid; 33    0.039300 0.54800     0.886
## 47         Benzeneacetic acid; 47   -0.038800 0.55500     0.886
## 48              Octanoic acid; 68    0.034000 0.60500     0.927
## 49           Linoleic acid, TMS; 4   -0.032900 0.61700     0.927
## 50            Myristoleic acid; 65   -0.032400 0.62200     0.927
## 51 3-Hydroxybutyric acid, 2TMS; 1   -0.031000 0.63700     0.927
## 52              Decanoic acid; 52    0.030500 0.64300     0.927
## 53         11-Eicosenoic acid; 35    0.029400 0.65500     0.927
## 54      Docosahexaenoic acid; 53    0.027000 0.68000     0.945
## 55          Nonadecanoic acid; 66   -0.022800 0.72900     0.985
## 56                1-Dodecanol; 36   -0.019900 0.76200     0.985
## 57             Tartronic acid; 73    0.019400 0.76700     0.985
## 58          Fumaric acid, 2TMS; 9    0.018100 0.78400     0.985
## 59            alpha-Tocopherol; 26   -0.014700 0.82200     0.985
## 60               Ethanolamine; 56    0.014300 0.82800     0.985
## 61              Glyceric acid; 30    0.013200 0.84100     0.985
## 62               Pyruvic acid; 31    0.011100 0.86600     0.985
## 63                Campesterol; 49    0.009490 0.88500     0.985
## 64             1-Monopalmitin; 37   -0.009430 0.88600     0.985
## 65           Citric acid, 4TMS; 6    0.008590 0.89600     0.985
## 66         Succinic acid, 2TMS; 7   -0.008460 0.89800     0.985
## 67            1,3-Propanediol; 34    0.008190 0.90100     0.985
## 68           Stearic acid, TMS; 2    0.006220 0.92500     0.985
## 69 4-Hydroxybenzeneacetic acid; 4    0.005520 0.93300     0.985
## 70     Eicosapentaenoic acid; 55    0.004930 0.94000     0.985
## 71         Glyceryl-glycoside; 59   -0.004840 0.94100     0.985
## 72         Heptadecanoic acid; 60   -0.003960 0.95200     0.985
## 73                   Glycerol; 58    0.002490 0.97000     0.985
## 74           Pyroglutamic acid; 69   -0.002310 0.97200     0.985
## 75              Alanine, 2TMS; 25   -0.000462 0.99400     0.994
```

###### 5.5.3.2 Forest Plot of Model Coefficients

```
## Warning: Ignoring unknown aesthetics: x

## Warning: Ignoring unknown aesthetics: x

## Warning: Ignoring unknown aesthetics: x

## Warning: Ignoring unknown aesthetics: x

## Warning: Ignoring unknown aesthetics: x

## Warning: Ignoring unknown aesthetics: x

## Warning: Ignoring unknown aesthetics: x

## NULL
```

#### 5.6 Neuropathy Questionnaire (mnsineuropat)

##### 5.6.1 Crude Model

```
## [1] "Fitting models:"
## [1] "~ mnsineuropat"
## [1] ""
```

###### 5.6.1.1 Tables of Model Coefficients

```
## [1] ""
## [1] "Table: mnsineuropat"
## [1] "  (from model: "
## [1] "   ~ mnsineuropat)"
## [1] ""
## [1] "No significant associations at p.adj < 0.05"
```

###### 5.6.1.2 Forest Plot of Model Coefficients

```
## NULL
```

#### 5.7 Adjusted Model

```
## [1] "Fitting models:"
## [1] "~ mnsineuropat + Age + bmi + Blood_glucose + Duration_DM + Gender + Hba1c_baseline + log_Blood_
## [1] ""
```

###### 5.7.0.1 Tables of Model Coefficients

```
## [1] ""
## [1] "Table: mnsineuropat"
## [1] "  (from model: "
## [1] "   ~ mnsineuropat + Age + bmi + Blood_glucose + Duration_DM +"
## [1] "     Gender + Hba1c_baseline + log_Blood_TGA + Smoking + Statin +"
## [1] "     Total_cholesterol)"
## [1] ""
##                               Name Coefficient P.Value adj.P.Val
## 1                   Creatinine; 50     0.15300  0.0132     0.414
## 2             Myo inositol 6TMS; 1     0.14900  0.0154     0.414
## 3                Glycine, 3TMS; 17     0.14800  0.0166     0.414
## 4             Tridecanoic acid; 74    -0.13300  0.0305     0.509
## 5            Fumaric acid, 2TMS; 9     0.12500  0.0424     0.509
## 6                     Glycerol; 58     0.12100  0.0504     0.509
## 7                      Ribitol; 70     0.11500  0.0617     0.509
## 8             Malic acid, 3TMS; 11     0.11400  0.0643     0.509
## 9                  Bisphenol A; 48     0.11100  0.0717     0.509
## 10 4-Hydroxyphenyllactic acid; 44     0.11000  0.0735     0.509
## 11                Ribonic acid; 72     0.11000  0.0747     0.509
## 12               Decanoic acid; 52    -0.10500  0.0892     0.551
## 13       4-Deoxytetronic acid; 32     0.10300  0.0955     0.551
## 14              Oleic acid, TMS; 3     0.10000  0.1030     0.554
## 15            Stearic acid, TMS; 2     0.08600  0.1630     0.711
## 16              Tartronic acid; 73    -0.08240  0.1810     0.711
## 17               Proline, 2TMS; 21    -0.08170  0.1850     0.711
## 18             1,3-Propanediol; 34     0.07890  0.2000     0.711
## 19          Palmitic acid, TMS; 5     0.07830  0.2040     0.711
## 20 2,4-Dihydroxybutanoic acid; 28     0.07750  0.2080     0.711
## 21  2-hydroxy Isovaleric acid; 38     0.07750  0.2080     0.711
## 22          Glyceryl-glycoside; 59     0.07560  0.2200     0.711
## 23           Linoleic acid, TMS; 4     0.07420  0.2290     0.711
## 24 4-Hydroxybenzeneacetic acid; 4     0.07180  0.2440     0.711
## 25               Octanoic acid; 68    -0.07050  0.2520     0.711
## 26 3-Hydroxybutyric acid, 2TMS; 1     0.06980  0.2570     0.711
## 27 3,4-Dihydroxybutanoic acid; 27     0.06810  0.2690     0.711
## 28                     Ribitol; 71     0.06730  0.2740     0.711
## 29          Succinic acid, 2TMS; 7     0.06650  0.2810     0.711
## 30            alpha-Tocopherol; 26    -0.06550  0.2880     0.711
## 31            Citric acid, 4TMS; 6     0.06460  0.2940     0.711
## 32           11-Eicosenoic acid; 35     0.06230  0.3120     0.711
## 33          Heptadecanoic acid; 61     0.06050  0.3260     0.711
## 34            Isoleucine, 2TMS; 18    -0.05980  0.3320     0.711
## 35             Arabinopyranose; 51     0.05570  0.3660     0.711
## 36          Heptadecanoic acid; 60    -0.05530  0.3690     0.711
## 37            Methionine, 2TMS; 16    -0.05480  0.3730     0.711
## 38             Threonine, 3TMS; 12    -0.05420  0.3790     0.711
## 39           Nonadecanoic acid; 66     0.05370  0.3840     0.711
## 40          3-Indoleacetic acid; 40     0.05290  0.3900     0.711
## 41                Pyruvic acid; 31    -0.05220  0.3970     0.711
## 42      3-Indolepropionic acid; 41     0.05200  0.3980     0.711
## 43           Aminomalonic acid; 45     0.04770  0.4380     0.755
## 44               Alanine, 2TMS; 25     0.04730  0.4430     0.755
```

```
## 45          Myristoleic acid; 65     0.04540  0.4610     0.768
## 46               1-Dodecanol; 36    -0.04410  0.4740     0.768
## 47             Hydroxylamine; 62    -0.04340  0.4810     0.768
## 48             Glyceric acid; 30    -0.04230  0.4930     0.770
## 49             Leucine, 2TMS; 19    -0.03990  0.5170     0.791
## 50      Eicosapentaenoic acid; 55     0.03860  0.5300     0.796
## 51              Ethanolamine; 56    -0.03730  0.5450     0.799
## 52              Valine, 2TMS; 20    -0.03600  0.5590     0.799
## 53                  Tyrosine; 75    -0.03550  0.5650     0.799
## 54              Serine, 3TMS; 14     0.03390  0.5820     0.809
## 55            1-Monopalmitin; 37    -0.03270  0.5960     0.813
## 56 alpha-ketoglutaric acid, TMS M    -0.03140  0.6100     0.817
## 57          Pyroglutamic acid; 69     0.02790  0.6500     0.837
## 58         Benzeneacetic acid; 47     0.02740  0.6570     0.837
## 59        2-Palmitoylglycerol; 39    -0.02720  0.6590     0.837
## 60            Arachidic acid; 46     0.02600  0.6740     0.842
## 61        Phenylalanine, 2TMS; 13    -0.02380  0.6990     0.847
## 62                  Glycerol; 57    -0.02350  0.7030     0.847
## 63            Hydroxyproline; 64    -0.02230  0.7170     0.847
## 64 2-Hydroxybutyric acid, 2TMS; 2     0.02180  0.7230     0.847
## 65      4-Deoxytetronic acid; 33    -0.02070  0.7370     0.850
## 66             Nonanoic acid; 67     0.01550  0.8020     0.911
## 67    4-Hydroxybutanoic acid; 43     0.01320  0.8300     0.922
## 68      Arachidonic acid, TMS; 24    -0.01270  0.8360     0.922
## 69               Lactic acid; 29    -0.01150  0.8520     0.926
## 70         Glutamic acid, 3TMS; 8     0.01020  0.8690     0.931
## 71            L-5-Oxoproline; 63    -0.00845  0.8910     0.941
## 72          Cholesterol, TMS; 23     0.00735  0.9050     0.943
## 73       Docosahexaenoic acid; 53    -0.00380  0.9510     0.962
## 74               Campesterol; 49    -0.00362  0.9530     0.962
## 75            Dodecanoic acid; 54     0.00295  0.9620     0.962
```

###### 5.7.0.2 Forest Plot of Model Coefficients

```
## Warning: Ignoring unknown aesthetics: x

## Warning: Ignoring unknown aesthetics: x

## Warning: Ignoring unknown aesthetics: x

## Warning: Ignoring unknown aesthetics: x

## NULL
```

#### 5.8 Fully-Adjusted Model

```
## [1] "Fitting models:"
## [1] "~ mnsineuropat + Age + bmi + Blood_glucose + Duration_DM + Gender + Hba1c_baseline + log_Blood_
## [1] ""
```

###### 5.8.0.1 Tables of Model Coefficients

```
## [1] ""
## [1] "Table: mnsineuropat"
## [1] "  (from model: "
## [1] "   ~ mnsineuropat + Age + bmi + Blood_glucose + Duration_DM +"
## [1] "     Gender + Hba1c_baseline + log_Blood_TGA + Smoking + Statin +"
## [1] "     Total_cholesterol + egfr)"
## [1] ""
##                               Name Coefficient P.Value adj.P.Val
## 1            Tridecanoic acid; 74   -1.41e-01  0.0211     0.654
## 2               Glycine, 3TMS; 17    1.33e-01  0.0295     0.654
## 3                 Bisphenol A; 48    1.11e-01  0.0698     0.654
## 4   2-hydroxy Isovaleric acid; 38    1.07e-01  0.0809     0.654
## 5                  Creatinine; 50    1.05e-01  0.0856     0.654
## 6                    Glycerol; 58    1.05e-01  0.0861     0.654
## 7             Stearic acid, TMS; 2    1.05e-01  0.0877     0.654
## 8               Oleic acid, TMS; 3    1.05e-01  0.0882     0.654
## 9               Decanoic acid; 52   -1.03e-01  0.0924     0.654
## 10                    Ribitol; 70    1.03e-01  0.0943     0.654
## 11          Fumaric acid, 2TMS; 9    1.02e-01  0.0960     0.654
## 12           Malic acid, 3TMS; 11    9.49e-02  0.1220     0.704
## 13              Proline, 2TMS; 21   -9.27e-02  0.1310     0.704
## 14          Palmitic acid, TMS; 5    9.25e-02  0.1310     0.704
## 15            Myo inositol 6TMS; 1    8.17e-02  0.1830     0.800
## 16              Tartronic acid; 73   -8.05e-02  0.1890     0.800
## 17 4-Hydroxyphenyllactic acid; 44    8.02e-02  0.1910     0.800
## 18             1,3-Propanediol; 34    7.84e-02  0.2010     0.800
## 19 3-Hydroxybutyric acid, 2TMS; 1    7.28e-02  0.2350     0.800
## 20       4-Deoxytetronic acid; 33   -6.93e-02  0.2580     0.800
## 21          Linoleic acid, TMS; 4    6.89e-02  0.2610     0.800
## 22                Serine, 3TMS; 14    6.84e-02  0.2650     0.800
## 23 2-Hydroxybutyric acid, 2TMS; 2    6.73e-02  0.2720     0.800
## 24         Heptadecanoic acid; 61    6.72e-02  0.2730     0.800
## 25         Succinic acid, 2TMS; 7    6.50e-02  0.2890     0.800
## 26            alpha-Tocopherol; 26   -6.12e-02  0.3190     0.800
## 27              Hydroxyproline; 64   -6.09e-02  0.3200     0.800
## 28          Nonadecanoic acid; 66    6.00e-02  0.3280     0.800
## 29         11-Eicosenoic acid; 35    5.91e-02  0.3350     0.800
## 30       4-Deoxytetronic acid; 32    5.87e-02  0.3390     0.800
## 31         Heptadecanoic acid; 60   -5.67e-02  0.3550     0.800
## 32               Ribonic acid; 72    5.66e-02  0.3560     0.800
## 33               Hydroxylamine; 62   -5.66e-02  0.3560     0.800
## 34      Eicosapentaenoic acid; 55    5.58e-02  0.3630     0.800
## 35     3-Indolepropionic acid; 41    5.42e-02  0.3770     0.807
## 36               Pyruvic acid; 31   -5.03e-02  0.4120     0.809
## 37          Aminomalonic acid; 45    4.98e-02  0.4170     0.809
## 38            Arabinopyranose; 51    4.94e-02  0.4210     0.809
## 39              Octanoic acid; 68   -4.83e-02  0.4310     0.809
## 40            Threonine, 3TMS; 12   -4.83e-02  0.4310     0.809
## 41           Myristoleic acid; 65    4.44e-02  0.4690     0.853
## 42                 1-Dodecanol; 36   -4.35e-02  0.4780     0.853
## 43               Alanine, 2TMS; 25    4.13e-02  0.5000     0.856
## 44         Glyceryl-glycoside; 59    4.07e-02  0.5070     0.856
```

```
## 45           1-Monopalmitin; 37   -4.01e-02  0.5140     0.856
## 46         Methionine, 2TMS; 16   -3.41e-02  0.5780     0.906
## 47      Phenylalanine, 2TMS; 13   -3.32e-02  0.5890     0.906
## 48 alpha-ketoglutaric acid, TMS M   -3.21e-02  0.6010     0.906
## 49           Arachidic acid; 46    3.08e-02  0.6150     0.906
## 50        Glutamic acid, 3TMS; 8    3.01e-02  0.6230     0.906
## 51          Citric acid, 4TMS; 6    2.94e-02  0.6310     0.906
## 52         Isoleucine, 2TMS; 18   -2.88e-02  0.6390     0.906
## 53        Cholesterol, TMS; 23    2.83e-02  0.6440     0.906
## 54      2-Palmitoylglycerol; 39   -2.76e-02  0.6520     0.906
## 55            Glyceric acid; 30   -2.50e-02  0.6830     0.926
## 56        Benzeneacetic acid; 47    2.38e-02  0.6980     0.926
## 57            Leucine, 2TMS; 19   -2.33e-02  0.7040     0.926
## 58             Ethanolamine; 56   -2.16e-02  0.7240     0.937
## 59       3-Indoleacetic acid; 40    2.04e-02  0.7400     0.941
## 60 4-Hydroxybenzeneacetic acid; 4    1.89e-02  0.7590     0.948
## 61                 Tyrosine; 75   -1.66e-02  0.7870     0.967
## 62 2,4-Dihydroxybutanoic acid; 28    1.55e-02  0.8000     0.968
## 63 3,4-Dihydroxybutanoic acid; 27    1.45e-02  0.8130     0.968
## 64            Nonanoic acid; 67    1.32e-02  0.8300     0.973
## 65      Docosahexaenoic acid; 53    1.06e-02  0.8630     0.982
## 66             Valine, 2TMS; 20   -1.05e-02  0.8640     0.982
## 67           Dodecanoic acid; 54    7.29e-03  0.9050     0.999
## 68                 Glycerol; 57   -5.06e-03  0.9340     0.999
## 69     4-Hydroxybutanoic acid; 43    4.76e-03  0.9380     0.999
## 70          Pyroglutamic acid; 69    4.42e-03  0.9430     0.999
## 71              Campesterol; 49   -3.91e-03  0.9490     0.999
## 72            L-5-Oxoproline; 63   -3.11e-03  0.9600     0.999
## 73                  Ribitol; 71   -6.43e-04  0.9920     0.999
## 74              Lactic acid; 29    5.49e-04  0.9930     0.999
## 75      Arachidonic acid, TMS; 24    5.95e-05  0.9990     0.999
```

###### 5.8.0.2 Forest Plot of Model Coefficients

```
## Warning: Ignoring unknown aesthetics: x

## Warning: Ignoring unknown aesthetics: x

## Warning: Ignoring unknown aesthetics: x

## Warning: Ignoring unknown aesthetics: x

## Warning: Ignoring unknown aesthetics: x

## Warning: Ignoring unknown aesthetics: x

## NULL
```

### 6 Appendix

```
## R version 3.6.2 (2019-12-12)
## Platform: x86_64-w64-mingw32/x64 (64-bit)
## Running under: Windows 10 x64 (build 17763)
##
## Matrix products: default
##
## locale:
## [1] LC_COLLATE=English_United States.1252
## [2] LC_CTYPE=English_United States.1252
## [3] LC_MONETARY=English_United States.1252
## [4] LC_NUMERIC=C
## [5] LC_TIME=English_United States.1252
##
## attached base packages:
## [1] stats     graphics  grDevices utils     datasets  methods   base
##
## loaded via a namespace (and not attached):
##  [1] Rcpp_1.0.3         plyr_1.8.5         pillar_1.4.3       compiler_3.6.2
##  [5] RColorBrewer_1.1-2 forcats_0.4.0      tools_3.6.2        digest_0.6.23
##  [9] evaluate_0.14      lifecycle_0.2.0    tibble_3.0.1       gtable_0.3.0
## [13] pkgconfig_2.0.3    rlang_0.4.6        yaml_2.2.0         haven_2.2.0
## [17] xfun_0.12          stringr_1.4.0      dplyr_0.8.3        knitr_1.27
## [21] vctrs_0.2.4        hms_0.5.3          grid_3.6.2         tidyselect_1.0.0
## [25] glue_1.3.1         R6_2.4.1           rmarkdown_2.1      limma_3.42.0
## [29] reshape2_1.4.3     ggplot2_3.2.1      readr_1.3.1        purrr_0.3.3
## [33] farver_2.0.3       tidyr_1.0.0        magrittr_1.5       scales_1.1.0
## [37] ellipsis_0.3.0     htmltools_0.4.0    assertthat_0.2.1   colorspace_1.4-1
## [41] labeling_0.3       stringi_1.4.4      lazyeval_0.2.2     munsell_0.5.0
## [45] crayon_1.3.4
```
